# The winding roads to adulthood: a twin study

**DOI:** 10.1101/2021.02.16.431456

**Authors:** Kaili Rimfeld, Margherita Malancini, Amy E. Packer, Agnieszka Gidziela, Andrea G. Allegrini, Ziada Ayorech, Emily Smith-Woolley, Andrew McMillan, Rachel Ogden, Philip S. Dale, Thalia C. Eley, Robert Plomin

## Abstract

In the 21^st^ century, emerging adulthood has stretched from the late teens through the twenties. Although this extended transition to adulthood can create stress, it can also offer opportunities to explore vocations and relationships that provide a better fit to individuals’ proclivities, including their genetic propensities.

Here we report the results of the first systematic investigation of genetic and environmental influences on 57 psychological traits covering major issues in emerging adulthood such as aspirations, thoughts and attitudes, relationships and personality. We also investigate how these traits relate to physical and mental health, educational attainment and wellbeing using a sample of nearly 5000 pairs of UK twins aged 21-25 from the Twins Early Development Study.

All 57 traits showed significant genetic influence, with an average heritability of 34% (SNP heritability ∼10%). Most of the variance (59% on average) was explained by non-shared environmental influences. These diverse traits were associated with mental health (average correlation .20), wellbeing (.16), physical health (.12) and educational attainment (.06). Shared genetic factors explained the majority of these correlations (∼50%). Together, these emerging adulthood traits explained on average 30% of the variance in the outcomes (range = 8 to 69%), suggesting that these traits relate to the outcomes additively.

We conclude that the environmental uncertainties of emerging adulthood in the 21^st^ century do not diminish the importance of genetics. As adolescents travel down long and winding roads to adulthood, their trip is substantially influenced by genetic proclivities that nudge them down different paths leading to different destinations.

## Introduction

The winding road to adulthood is the subtitle of the field-defining book on emerging adulthood by Jeffrey Arnett (Arnett, 2004, 2015). The book documents the recent societal changes – for example, insecure employment, unstable relationships, difficulties in owning a home -- that have prolonged the transition from adolescence to adulthood, a process which he called *emerging adulthood*. The transition from adolescence to adulthood is marked by instability, self-focus, feeling in-between, exploring identity in work and love, and an optimistic sense of possibilities.

An enormous menu of experiences is on offer during emerging adulthood for exploring self, relationships, aspirations, attitudes, autonomy and occupations. The bespoke nature of these experiences is amplified by the internet and social media. The current generation of 20+-year-olds is the first generation of digital natives. Twenty years ago, the internet, e-mail, and mobile phones were still a novelty. Even in 2004, when the first edition of Arnett’s book was published, Facebook, Twitter, and Instagram did not exist or were not so widely used.

More than 6000 papers, primarily in psychology and psychiatry, have investigated emerging adulthood. Most of these are normative, describing average changes that take place during the twenties (Arnett, 2016). However, the paths individuals take can vary widely. Much less is known about individual differences, which are a key feature of emerging adulthood as the transition to adulthood is less lock-stepped than in the past.

There is even variability in the extent to which young people *want* to emerge into adulthood. According to Arnett, this involves accepting responsibility for oneself, making independent decisions and becoming financially independent. In a US survey of 1029 individuals aged 18-29, a third agreed with the statement, ‘If I could have my way, I would never become an adult’ (Arnett & Schwab, 2012). The slogan of the three protagonists wandering through their twenties together in *Generation X* is ‘dead at 30, buried at 70’ (Coupland, 1991).

Genetic and environmental analyses of individual differences in emerging adulthood have been reported for domains that are assessed throughout the life course (Bergen, Gardner, & Kendler, 2007). These include psychopathology (Polderman et al., 2015), personality (Turkheimer, Pettersson, & Horn, 2014) and cognitive abilities (Briley & Tucker-Drob, 2013). However, much less is known about the genetic and environmental aetiology of individual differences in traits especially salient in emerging adulthood, such as identity, aspirations, and relationships. Two exceptions are political orientation and participation (Hufer, Kornadt, Kandler, & Riemann, 2020; Kornadt, Hufer, Kandler, & Riemann, 2018) and religiousness (Koenig, McGue, & Iacono, 2008). Interestingly, these studies generally suggest greater genetic influence and less shared environmental influence in emerging adulthood than in adolescence, especially for political ideology (Hatemi & McDermott, 2012).

The great heterogeneity of experience during emerging adulthood suggests the hypothesis that environmental differences are especially important. A refinement of this hypothesis of heightened environmental influence is that any residual effect of family environment is overwhelmed as emerging adults make their own way in the world. The instability and uncertainty of emerging adulthood produces idiosyncratic experiences that nudge individuals down different paths. Such experiences are not likely to be shared by siblings, which in behavioural genetics is called non-shared environment.

A less obvious possibility is that, despite the heterogeneity of environmental experiences in emerging adulthood, genetic influence on individual differences remains substantial, which is the trend found in the few relevant twin studies of emerging adulthood (Hufer et al., 2020; Koenig et al., 2008; Kornadt et al., 2018). One major mechanism by which genotypes become phenotypes is by affecting how individuals select, modify and create experiences in part on the basis of their genetic propensities. This is called gene-environment correlation. For this reason, it is possible that the instability and uncertainties of emerging adulthood create more opportunities for individuals to make choices and select increasingly diverse life experiences that are correlated with their genotypes.

The specific choices, aspirations, thoughts and feelings of emerging adults are likely to be related to outcomes such as physical and mental health, wellbeing, and socioeconomic status (Bonnie, Stroud, & Breiner, 2015; Johnson, Crosnoe, & Elder, 2011). The extended transition from adolescence to adulthood is associated with a prolonged period of risk-taking, decision making and self-regulation with markedly less parental control and parental influence on lifestyle, exercise and diet (Bonnie et al., 2015). These factors are all likely to be associated with physical and mental health and wellbeing. Emerging adults, for the first time, can make decisions about their daily life, sleeping, eating and exercise patterns. These decisions are likely to be linked to their personality and life goals, relationships and aspirations for life (Harris, Gordon-Larsen, Chantala, & Udry, 2006).

On average the physical health of emerging adults tends to be better than that of older adults. For example, 96% of the US emerging adults report being in excellent, very good or good health (Park, Paul Mulye, Adams, Brindis, & Irwin, 2006). Nonetheless, emerging adults differ and poor health during this period predicts early onset of chronic conditions such as cardiovascular problems that increase in general later in life (Tanner, 2016; Yaffe et al., 2014).

Unlike physical illness, mental illness is highly prevalent during emerging adulthood, as most mental illness begins to emerge by adolescence (Kessler et al., 2005; Tanner, 2016). The US National Survey of Comorbidity reports that up to 44% of young adults meet the criteria for a mental health disorder in a given year. Wellbeing, which is more than the absence of mental illness, is also important, and likely related to many traits especially relevant to emerging adulthood traits (Ong & Bergeman, 2004; Ong, Bergeman, Bisconti, & Wallace, 2006; van de Weijer, de Vries, & Bartels, 2020; Veenhoven, 2008). However, little is known about the causes and correlates of wellbeing or mental health and illness during emerging adulthood.

Another key outcome in emerging adulthood is educational attainment for those who continue their education beyond compulsory schooling. This is especially important since educational outcomes enable young adults to pursue different lifelong trajectories. Importantly, educational attainment is related to physical and mental health across the lifespan (Cutler & Lleras-Muney, 2012). Educational achievement during compulsory schooling can be explained by a package of heritable cognitive and non-cognitive traits, such as personality, behavioural problems, self-esteem (Krapohl et al., 2014), but less is known about the correlates of educational attainment during emerging adulthood.

The current study investigates the genetic and environmental aetiology of individual differences in traits especially relevant to emerging adulthood, including life goals, purpose in life, relationships, and attitudes towards marriage, money and occupation. We refer to these traits as emerging adulthood (EA) traits. This is the first report of an emerging adulthood assessment of twins in the Twins Early Development Study when the twins were 21 to 25 years old (Rimfeld et al., 2019).

Our second objective is to examine the genetic and environmental aetiology of the extent to which these emerging adulthood traits are associated with physical health, mental health and illness, wellbeing and educational outcomes and aspirations.

These exploratory analyses, together with specific hypotheses, were preregistered in Open Science Framework (osf.io/m6vwj) prior to accessing the data.

## Methods

### Sample

Our sample was drawn from the Twins Early Development Study (TEDS), a longitudinal study that recruited through national birth records over 16,000 twin pairs born between 1994 and 1996 in England and Wales. Although there has been some attrition, more than 8,000 twin pairs are still engaged in the study, and TEDS remains reasonably representative of the population in England and Wales in terms of ethnicity and family socioeconomic factors, including data from the most recent data collection used in the present study (Rimfeld et al., 2019). Zygosity was assessed using a parent-reported questionnaire of physical similarity, which has been shown to be over 95% accurate when compared to DNA testing (Price et al., 2000). When zygosity was not clear, DNA testing was used to resolve zygosity.

Here we used the subsample of twins who had contributed to the TEDS-21 data collection. At the start of TEDS-21 data collection, twin ages ranged from 20.56 to 25.59 years (mean= 22.29, SD=.92). All data were collected using questionnaires. Due to the length and number of measures included, data collection was completed in two phases (Phase 1 started in June 2017, phase 2 started in February 2018, and both continued until January or February 2019). In both phases of TEDS-21 questionnaire data collection, participants could complete the assessment via a smartphone app, via the web, or on paper. Incentives included a £10 voucher on completion of each phase. Twins were also offered entries in prize draws at intervals during the data collections of phases 1 and 2. Ethical approval was provided by the King’s College London Research Ethics Committee (reference number: PNM/09/10-104) and informed written consent was received from all participants. No exclusions were applied. All available data was used (N = 10,614, twins who had completed at least one phase of TEDS-21 data collection), therefore the sample size varied between measures. Sample size by zygosity for each measure is presented in Supplementary Table S2 and Table S3 (one twin randomly selected from each pair so that the data points are independent).

### Measures

The Twins Early Development Study (TEDS) measures have been summarised previously (Rimfeld et al., 2019). Figure 1a lists the measures included in this study, which were administered during TEDS-21 data collection; a more detailed description of measures can be found in Supplementary Table S1. We included 57 measures of traits selected to represent issues in emerging adulthood (EA) such as aspirations, thoughts and attitudes, life events, relationships, sexual and health behaviour and personality. We also included measures related to what are often considered to be the core functional outcomes even though here we refer to the data collected at the same time: adverse physical health (e.g., over the counter painkillers taken, hospitalisations, self-reported health), adverse mental health (e.g., variables covering diverse forms of psychopathology), wellbeing (e.g., financial wellbeing, satisfaction with relationships and community), and education. These variables were used to create factors representing wellbeing, adverse mental health, and adverse physical health using principal component analyses (PCA). These composite measures, including their factor loadings, are presented in Figure 1b. Two items were used that assessed the highest educational level achieved at the time of data collection and the highest planned educational level for the future.

**Figure 1a.**
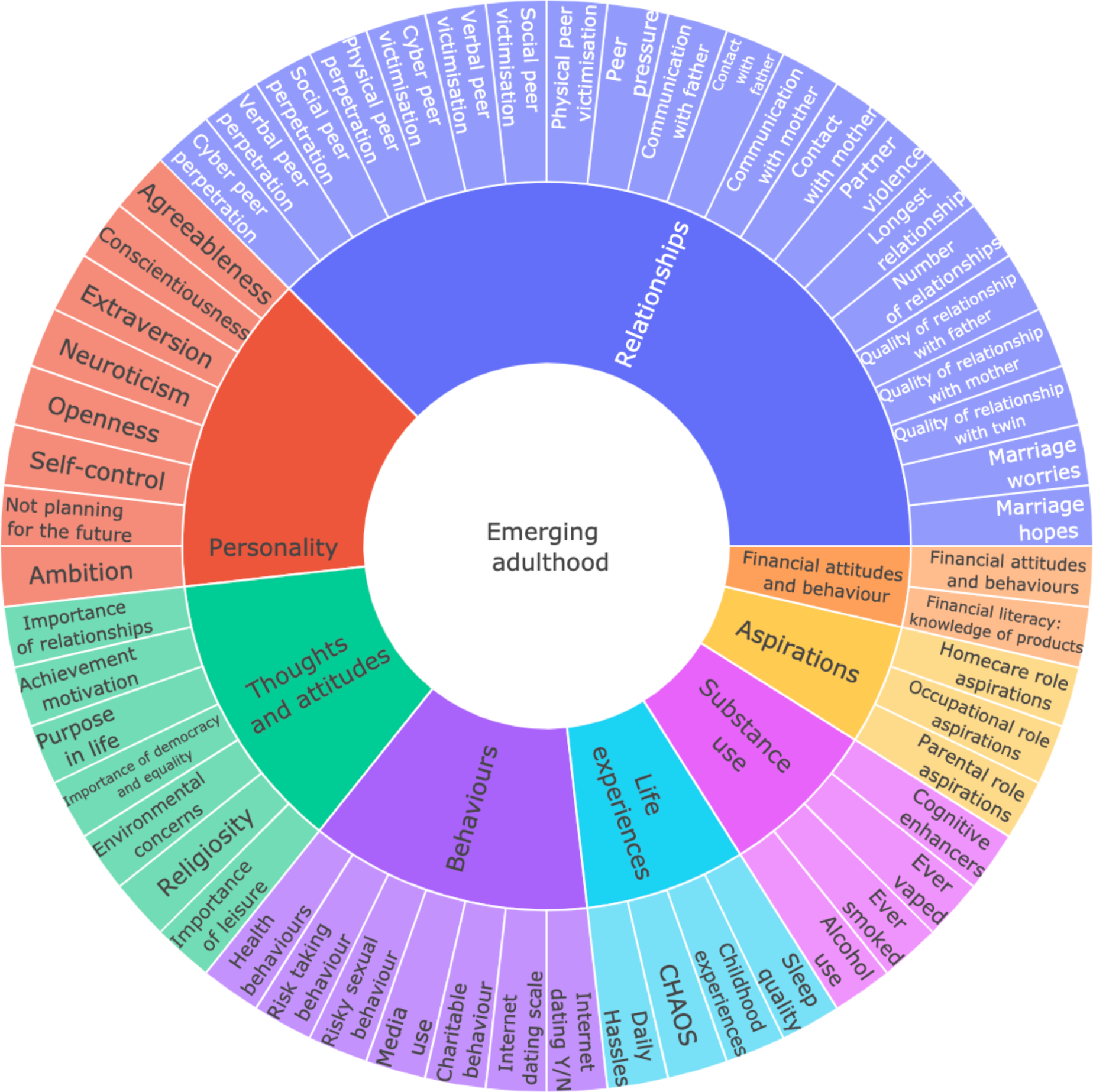
Summary of emerging adulthood measures

**Figure 1b.**
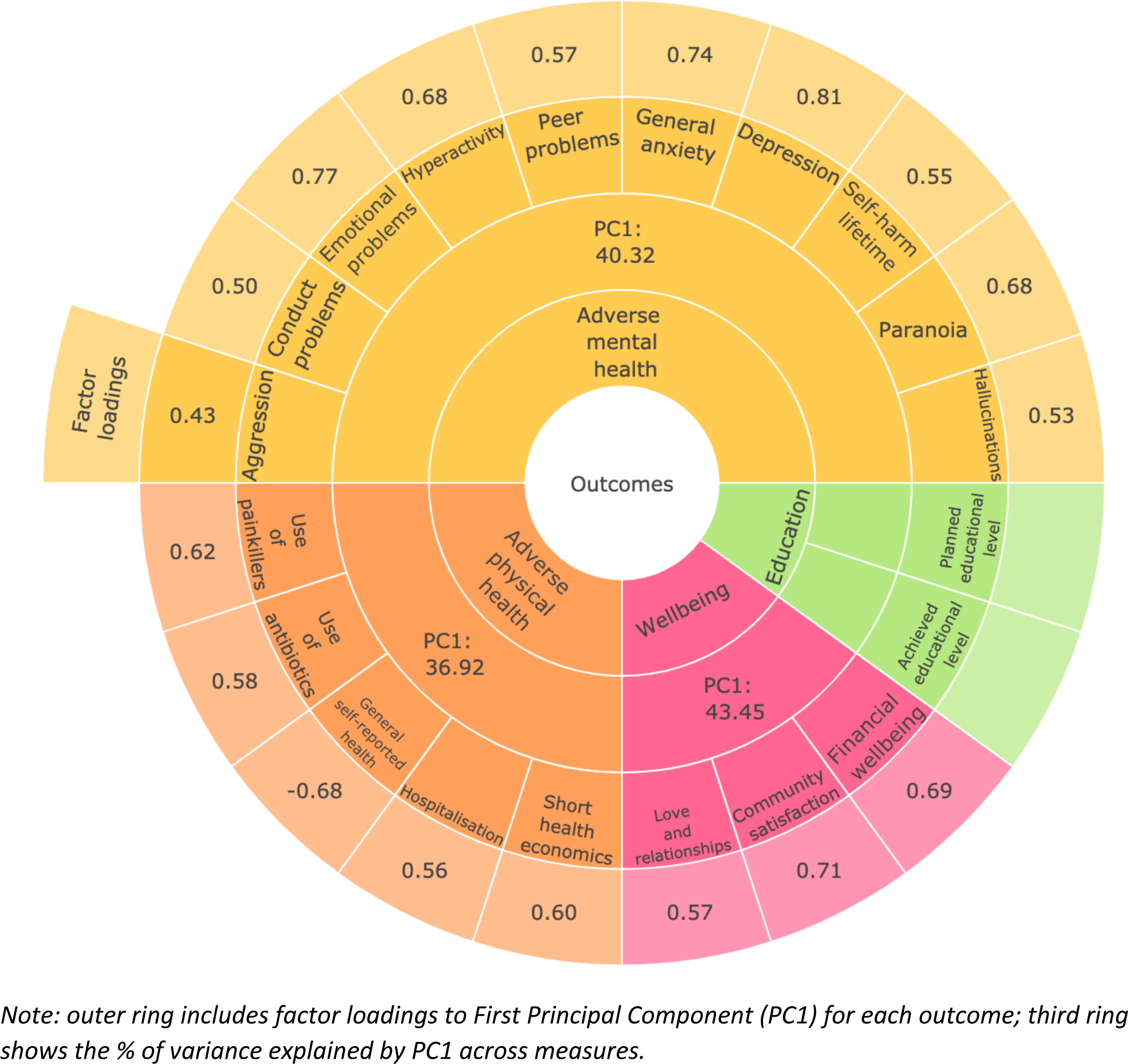
Summary of functional outcomes; physical and mental health, wellbeing, achievement and planned educational level.

DNA samples have been obtained from 12,500 individuals and genotyped on one of two DNA microarrays (Affymetrix GeneChip 6.0 or Illumina HumanOmniExpressExome chips). After stringent quality control, the total sample size available for genomic analyses is 10,346 (including 7,026 unrelated individuals and 3,320 additional DZ co-twins). Of these, 7,289 individuals were genotyped on Illumina arrays, and 3,057 individuals and were genotyped on Affymetrix arrays ((Rimfeld et al., 2019); see (Selzam et al., 2018) for a detailed description of genotyping and quality control). DNA data was used for genome-based restricted maximum likelihood methods using genome-wide complex trait analyses (see genetic analyses). The sample size for DNA-based analyses varied between measures (see Supplementary Table S14).

### Quality control

Data cleaning (see, https://www.teds.ac.uk/datadictionary/studies/rawdata/web_data_cleaning.htm#prandomteds21) included the identification of random responders, or “clickers”. Clickers were identified from a combination of the following: incorrect responses on a quality control item (for example, we asked participants to select option ‘b’), too rapid responding (based on the mean item response time; web and smartphone app respondents only), or too uniform responding (i.e., selecting the same response over a series of consecutive items). Typically, around 1% of twins were excluded per questionnaire section but this varied from .3% to 5.3% (and of course 0% in themes without any measure containing a QC item). Twin data were excluded for an entire questionnaire section if the twin was identified as a clicker for any measure within that theme. Furthermore, twin data were excluded for an entire questionnaire (phase 1 or phase 2) if the twin was identified as a clicker for two or more themes within that questionnaire. For further details, including percentages of quality control exclusions, see Appendix S1.

### Statistical analyses

Our statistical analysis plan was registered in the Open Science Framework (OSF), prior to creation of the dataset and prior to analysis (osf.io/m6vwj). Scripts have been made available on the OSF site. All analyses were completed using SPSS Statistics software (v26, IBM Corp., Armonk, NY) and R version 4.0 (R Core Team, 2020).

### Data transformations

Several variables were highly skewed. We therefore used the rank-based van der Waerden’s transformation to achieve normality (Lehmann, 1975; Van Der Waerden, 1975). Sensitivity analyses were performed to examine the influence of the van der Waerden transformation on results. The results remained highly similar to untransformed data analyses. Results are reported both on untransformed data and transformed data (see Supplementary Materials).

### Phenotypic analyses

Because the present sample is a twin sample, we maintained independence of data by randomly selecting one twin per pair for all phenotypic analyses. We repeated analyses of descriptive statistics for the other half of the sample. These results are presented in Supplementary Material and are highly similar.

The 57 psychological trait measures were described in terms of means and variance, and compared between males and females, and identical and non-identical twins. Univariate analysis of variance (ANOVA) was used to test for mean differences for sex and zygosity and their interaction. Because significant, though small, sex differences emerged (Supplementary Table S2), we corrected all scores for mean sex differences using the regression method. Correcting for sex and age is important in the analysis of twin data because members of a twin pair are identical in age and identical twins are identical for sex, which, if uncorrected, would inflate twin estimates of shared environment (McGue & Bouchard, 1984). These age- and-sex adjusted standardised residuals were used in all subsequent analyses. For analyses using transformed data, we conducted the van der Waerden transformation prior to residualizing for age and sex as has been recommended by Pain et al. (Pain, Dudbridge, & Ronald, 2018).

We created standardised composite scores representing adverse mental health, adverse physical health and wellbeing using principal components analysis (PCA). We obtained the first principal component (1st PC) of behaviour problem/mental health phenotypes (see Figure 1b and Supplementary Table S1 for information about scales used). We also obtained the first principal component of physical health and wellbeing measures (see Table S1 for scales used). The 1st PCs of wellbeing, adverse mental health and adverse physical health were saved for subsequent analyses. Two separate items were used to assess educational outcomes: level of current education and level of planned education. We did not compute an educational attainment composite.

We calculated the phenotypic correlations between the 57 traits and the functional outcome factors (adverse mental health factor, adverse physical health factor, wellbeing factor, and two educational attainment variables) for the whole sample. We then repeated the analyses separately for males and females.

We also conducted multiple regression to assess how much variance in the functional outcomes could be explained in total by the 57 traits, reporting the adjusted R^2^ values.

### Genetic analyses

#### Univariate twin analyses

We applied the twin method, specifically the univariate ACE model, to investigate the aetiology of individual differences in psychological traits and functional outcomes (adverse physical health, adverse mental health, wellbeing, and educational attainment). This method capitalises on the fact that identical (monozygotic, MZ) twin pairs are genetically identical and share 100% of their genes, while non-identical (dizygotic, DZ) twin pairs share on average 50% of their segregating genes Shared environment is defined as aspects of the environment that make members of twin pairs similar to one another. Non-shared environmental influences are unique to individuals and do not contribute to similarities between twins. Using these family relatedness coefficients, it is possible to estimate the relative influence of additive genetic (A), shared environmental (C), and non-shared environmental (E) effects on the variance and covariance of phenotypes, by comparing intraclass correlations for MZ and DZ twins (Knopik, Neiderhiser, DeFries, & Plomin, 2017). In the model, non-shared environmental variance also includes any measurement error. Heritability can be roughly calculated by doubling the difference between MZ and DZ correlations, C can be calculated by subtracting heritability from the MZ correlation and E can be estimated by deducting the MZ correlation from unity (Rijsdijk & Sham, 2002). These parameters can be estimated more accurately using structural equation modelling (SEM), which also provides 95% confidence intervals and estimates of model fit. The SEM structural equation modelling program OpenMx was used for all model-fitting analyses (Boker et al., 2011). We report MZ and DZ same-sex twin intraclass correlations, model fit statistics and ACE estimates.

#### Sex-limitation analyses

The univariate model can be extended to the full sex limitation model to explore both qualitative and quantitative sex differences in the aetiology of individual differences in psychological traits, wellbeing, adverse physical health, adverse mental health and educational attainment. Qualitative sex differences indicate that different genetic or environmental factors influence a phenotype in males and females. Quantitative sex differences are observed when the same genetic and environmental factors influence variance in a given phenotype for males and females, but the magnitude of their effects differs across sexes. (For a detailed explanation of the full sex limitation model, see(Medland, 2004).) ACE estimates are also reported separately for males and females.

#### Bivariate Correlated Factors solution

The univariate model can be extended to a bivariate model to investigate the aetiology of the covariance between two traits. We used the bivariate twin design to calculate genetic correlations and estimate the A, C and E components of covariance between psychological traits versus wellbeing, adverse physical health, adverse mental health and educational attainment. The bivariate genetic method decomposes the covariance between phenotypes into A, C and E components by comparing the cross-trait cross-twin correlations between MZ and DZ twin pairs (Neale, Boker, Bergeman, & Maes, 2005). This method also enables estimation of the genetic correlation (rG), indicating the extent to which the same genetic variants influence two phenotypes. The shared environmental correlation (rC) and non-shared environmental correlation (rE) can also be estimated (Knopik et al., 2017; Rijsdijk & Sham, 2002). The bivariate genetic model also allows estimation of the genetic and environmental mediation of the phenotypic correlation between variables (see Supplementary Figure S1 for Cholesky composition). Genetic mediation of the phenotypic correlation is calculated by the genetic correlation between two variables weighted by their heritabilities. Shared environmental and non-shared environmental mediation of the phenotypic correlation can also be estimated in a similar manner.

#### Genome-based restricted maximum likelihood methods using genome-wide complex trait analyses (GREML-GCTA)

We used univariate GREML GCTA software to calculate SNP heritability for the 57 traits and adverse physical health, adverse mental health, wellbeing and educational attainment. GREML uses individual-level genotypic data to estimate the narrow-sense SNP-h^2^, that is, the proportion of phenotypic variation explained by the additive effects of SNPs assessed on a genotyping array and imputed SNPs (Yang, Lee, Goddard, & Visscher, 2013). Population stratification was controlled by using the 10 first principal components of the population structure as covariates in the model (based on the PCA), along with a categorical variable indexing genotyping batch. Univariate GREML can also be extended to bivariate GREML to decompose covariance between traits. Bivariate GREML-GCTA was used to calculate genetic correlations between psychological traits and adverse physical health, adverse mental health, wellbeing and educational attainment (Lee, Yang, Goddard, Visscher, & Wray, 2012).

## Results

### Descriptive statistics and comparisons of males and females

Means and standard deviations were calculated for the 57 emerging adulthood (EA) traits for the whole sample, for males and females separately (Figure 2), and for five sex and zygosity groups: monozygotic (MZ) males, dizygotic (DZ) males, MZ females, DZ females and DZ opposite-sex twin pairs (Supplementary Table S3). ANOVA results confirm the visual near-identity in the graph, by indicating that sex and zygosity together explain only 2% of the variance on average for psychological traits (Supplementary Table S3). These results are based on one twin randomly selected from each pair so that the data points are independent. We repeated these analyses with the other twin from each pair selected and results were highly similar (see Supplementary Tables S4 and S5).

**Figure 2.**
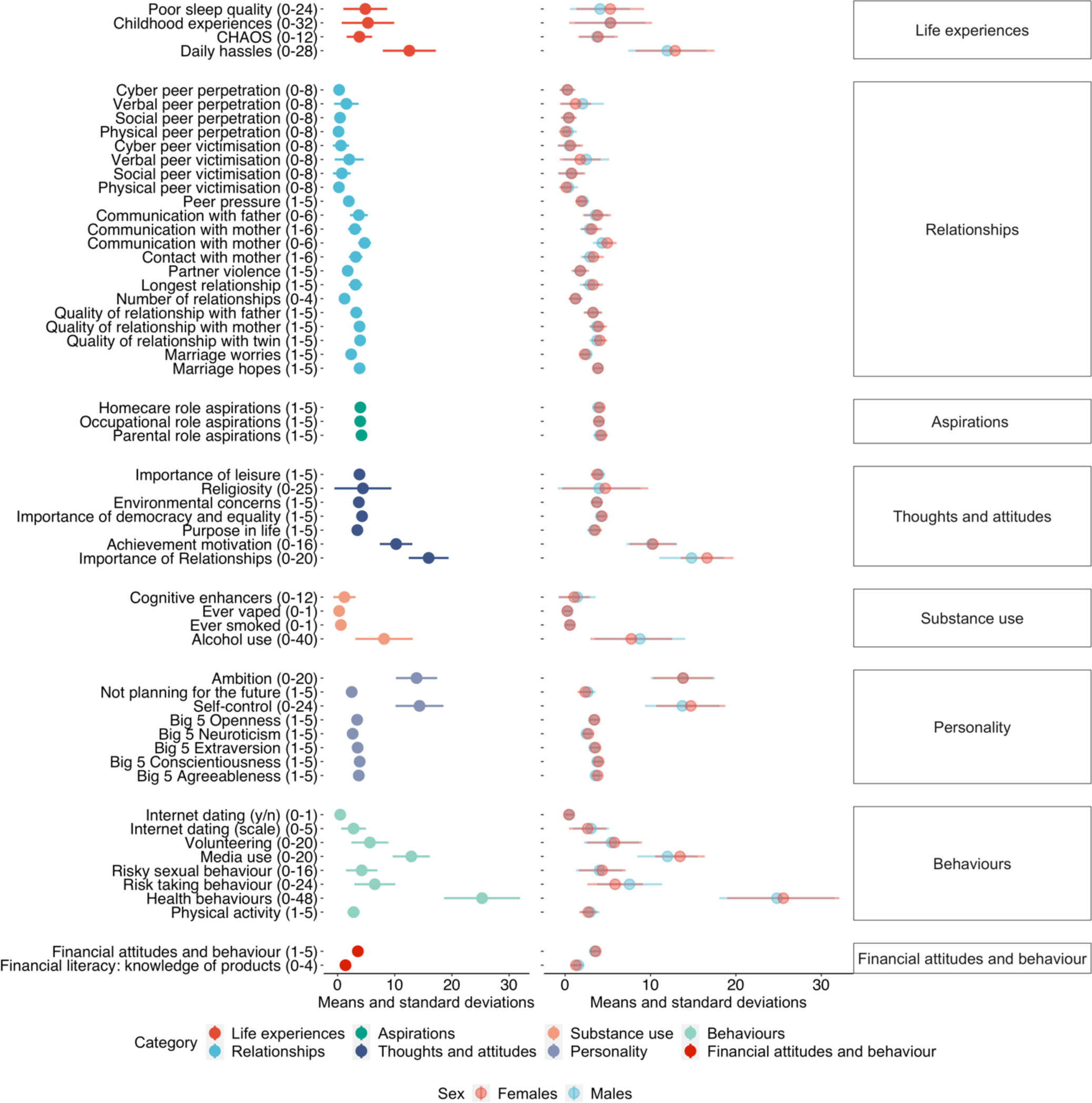
Descriptive statistics (means and standard deviations) for the whole sample and separately for males and females (minimum and maximum score for each scale in parentheses).

For subsequent analyses, the data were corrected for mean age and sex differences, as described in Methods.

### Wellbeing, adverse mental and physical health composite scores

Figure 1b lists the loadings on, and variance explained by, the first principal components of the functional outcomes (wellbeing, adverse mental health, adverse physical health). The variance explained by each 1st PC ranged from 37% to 44%, and all measures loaded on the respective 1st PC >.43. The 1st PC scores for wellbeing, adverse mental health and adverse physical health were saved for subsequent analyses. We did not include the Conner’s impulsivity and inattention subscales in this final mental health composite because these loaded <.4 on the 1st PC and removing the Conner’s subscales increased the variance explained by the 1st PC from 34% to 41%. Additionally, we included only the lifetime self-harm item, not the item about self-harm in the last year because the two self-harm items correlated highly (*r* = .55, *p* < .001, *N* = 3,887). We selected the lifetime item to maximise power (lifetime *N* = 4,727 *vs*. last year *N* = 3,887).

### Phenotypic analyses

Pearson correlations between the EA traits and the composite scores of adverse physical health, adverse mental health, wellbeing and educational attainment are shown by the total length of the bars in Figure 3 (see Supplementary Table S6 for correlation coefficients with 95% CI). These phenotypic correlations were mostly significant and moderate (the average correlation was .16 for wellbeing, .20 for adverse mental health, .12 for adverse physical health, and education .06). For wellbeing and adverse mental and adverse physical health, the strongest correlations emerged for life experience traits such as poor sleep quality and daily hassles, as well as for purpose in life, importance of leisure and neuroticism. Correlations were similar when calculated separately for males and females (see Supplementary Figure S2a and b). We also calculated these correlations after excluding opposite-sex DZ pairs from the analyses and the results are highly similar (see Supplementary Figure S2c).

**Figure 3.**
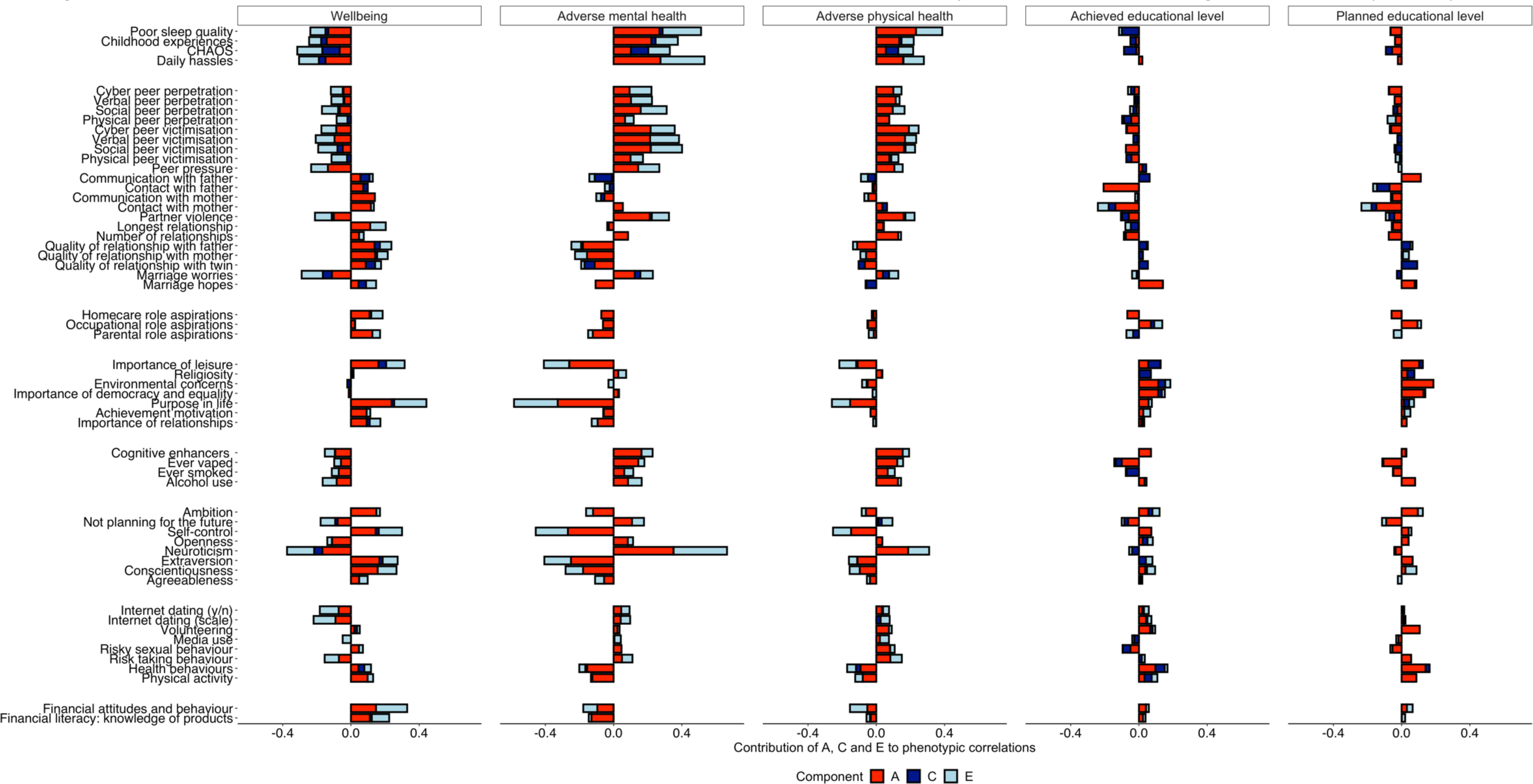
Phenotypic correlations between EA traits and key outcomes (indicated by the total length of the bar). Proportion of each correlation explained by additive genetic (A), shared environmental (C) and non-shared environmental factors are indicated by the red, dark blue and light blue bars, respectively.

Figure 4 summarises the results in Figure 3 using multiple regression to predict adverse physical and adverse mental health, wellbeing and achieved and planned educational level. Together, the emerging adulthood traits on average account for 31% (range = 8 to 68%) of the variance in the outcomes. The emerging adulthood traits are most strongly associated with adverse mental health, accounting for 68% of the variance. They were least strongly associated achieved (13%) and planned (8%) educational level. Full results of the multiple regression analyses, including results separately for males and females, are presented in Supplementary Table S7.

**Figure 4.**
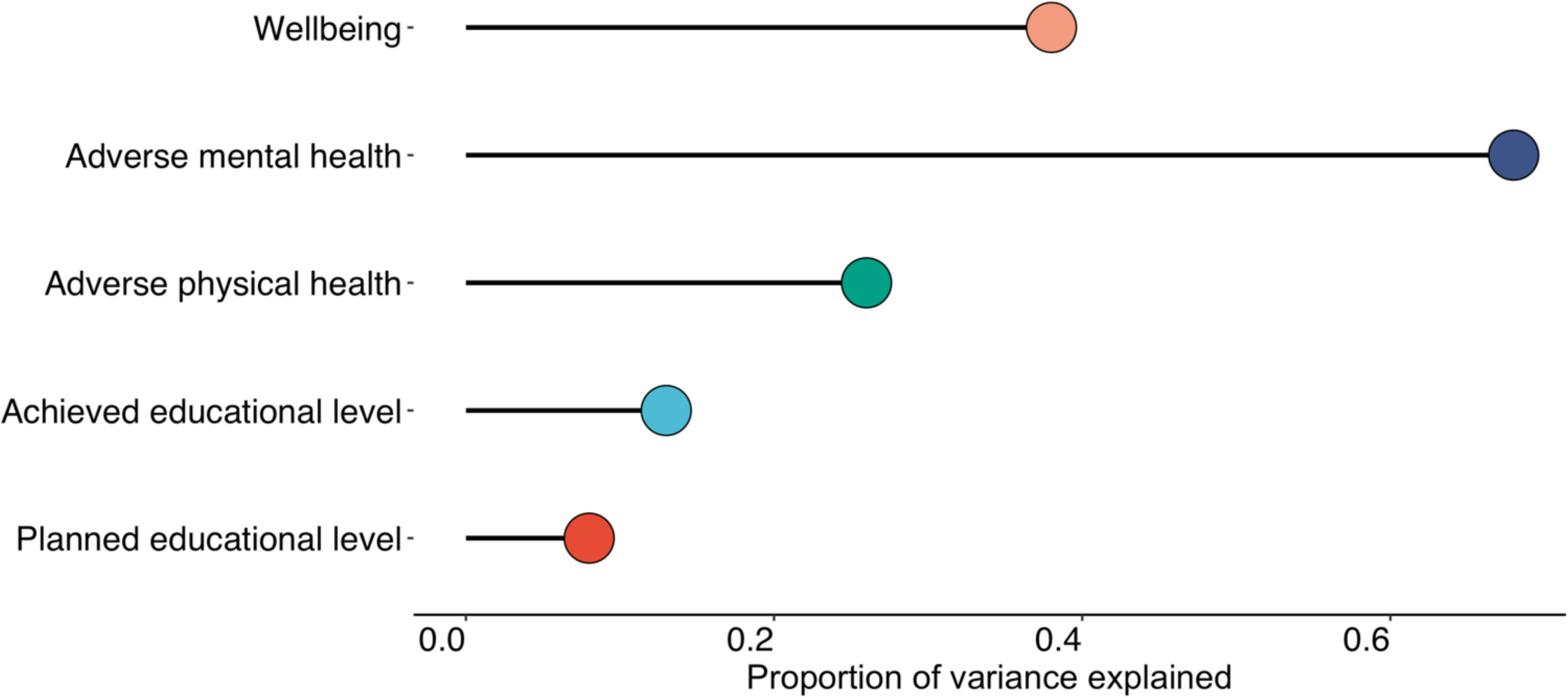
Summary of multiple regression analyses: variance explained (adjusted R^2^) in composite scores of adverse physical health, adverse mental health, wellbeing and educational attainment psychological traits

### Genetic analyses

#### Univariate twin analyses

All EA traits as well as functional outcomes showed significant heritability as presented in Figure 5. All twin correlations are presented in Supplementary Figure S3. Full ACE model-fitting results, including the 95% confidence intervals, are shown in Supplementary Table S9 and model-fit statistics in Supplementary Table S10. While for some variables the AE model showed better fit than the full ACE model, ACE model results are shown for completeness and to facilitate comparisons between variables.

**Figure 5.**
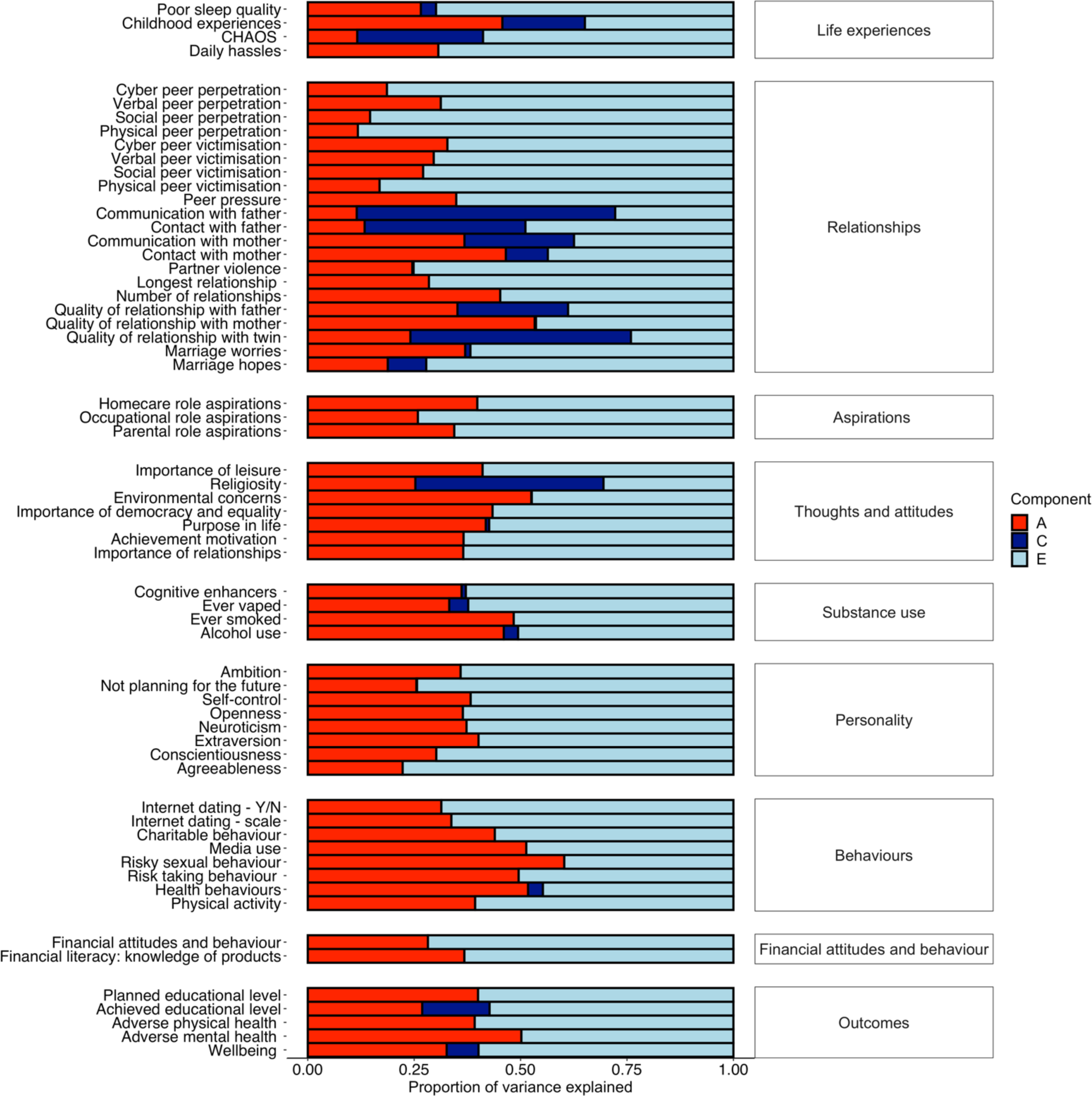
Univariate twin analyses of additive genetic (A), shared environmental (C), and non-shared environmental (E) components of variance for EA traits.

Across EA traits and functional outcomes, heritability (A) was modest on average (mean 34%, range 12% - 60%). The most heritable trait in emerging adulthood is risky sexual behaviour (.60; 95% CI: .52-.63). Other traits with substantial heritability include general risk-taking behaviour (.50; 95% CI: .42-.53), media use (.51; 95% CI: .48-.55), health behaviours (.52; 95% CI: .41-.58), environmental concerns (.53; 95% CI: .43-.56) and quality of relationship with mother (.53; 95% CI: .43-.57). These heritabilities are higher than those for the Big Five personality traits, which range from .22 (95% CI:.18-.27) to .40 (95% CI:.36-.44). For the outcome variables, the adverse mental health composite is more heritable (.50, 95% CI:.42-.54) than the adverse physical health composite (.39, 95% CI:.27-.43).

Shared environmental effects (C) were modest on average (mean 5%, range 0%–61%), but for a few variables substantial effects were observed. The greatest shared environmental effect was found for communication with father (.61; 95% CI: .53-.68). This might reflect estrangement due to divorce or father absence as it would result in greatly reduced communication with both members of the twin pair. Substantial shared environmental influence was also found for contact with father (.38; 95% CI: .24-.50) and quality of relationship with father (.26; 95% CI: .19-.33). Two other traits showing substantial shared environmental effects were relationship with twin (.52; 95% CI: .47-.56) and religiosity (.44; 95% CI: .39-.49). Non-shared environment (E) accounted for 59% of the variance on average (E range 24%–86%).

These univariate results were mostly similar when computed separately for males and females (Supplementary Figure S4a) and when DZ opposite-sex twin pairs were excluded (Supplementary Figure S4b).

#### Sex-limitation analyses

Some qualitative and quantitative sex differences emerged, for example, in childhood experiences, quality of relationship with mother and physical victimisation. Full model-fit statistics with nested models are presented in Supplementary Table S11; ACE estimates for males and females are presented in Supplementary Figure 4a and estimates with 95% confidence intervals in Supplementary Table S8. Two of the largest sex differences in heritability were found for chaos and marriage hopes. For chaos, heritability was .29 (95% CI:.10-.47) for males and .03 (95% CI:.00-.15) for females; for marriage hopes, heritability was .00 (95% CI:.00-.21) for males and .25 (95% CI:.11-.39) for females. However, as indicated by the overlapping confidence intervals for these estimates, limited power warrants caution in interpreting these results; they are presented here for completeness. As an additional sensitivity analysis, because we observed significant quantitative sex differences in the sex-limitation model-fitting analyses, we also calculated all estimates when excluding opposite-sex DZ twin pairs from analyses (presented in Supplementary Material) and the results remained highly similar.

#### Bivariate twin analyses

As mentioned earlier, Figure 3 showed the phenotypic correlations between EA traits and adverse physical health, adverse mental health, wellbeing and education. Figure 3 also included results for bivariate genetic analyses, which estimate the extent to which the phenotypic correlations between emerging adulthood traits and functional outcomes are mediated by genetic and environmental factors. The full results with 95% confidence intervals are presented in Supplementary Table S12.

On average, more than 50% of these phenotypic correlations were mediated by genetic factors, 60% for mental health, 56% for physical health, 50% for wellbeing and 52% for the two education outcomes. For example, the first row of Figure 3, shows that poor sleep quality is negatively correlated with wellbeing (r = -.23) and positively correlated with adverse mental health (.50) and adverse physical health (.38). These three correlations are largely mediated genetically – 56%, 54% and 60%, respectively. The rest of these correlations are mediated by nonshared environmental factors. For all of the substantial phenotypic correlations between EA traits and outcomes, genetic factors account for the majority of the correlations. Shared environment does not significantly contribute to any of these correlations.

Genetic contributions to the phenotypic correlations are based on the genetic correlation weighted by the heritability of the two traits. The genetic correlation estimates the extent to which the same genetic variants influence the two traits, regardless of their heritabilities. Figure 6 presents the genetic correlations between the EA traits and the outcome variables. Supplementary Table S13 shows the estimates with 95% confidence intervals. The average genetic correlation was .23 ignoring the sign of the correlation. Because all EA traits are moderately heritable, the pattern of results for genetic correlations in Figure 6 correspond to the results in Figure 3 showing the genetic contribution to the phenotypic correlations. For example, the genetic correlations between poor sleep quality and the outcome variables are -.45 for wellbeing, .74 for adverse mental health, and .74 for adverse physical health, all of which are statistically significant. Most of the other highest genetic correlations between EA traits and outcomes were observed for adverse mental health: daily hassles (.71), partner violence (.62), social peer victimisation (.60), Big 5 neuroticism (.83), and self-control (-.61). It should be noted that when the phenotypic correlation is negligible, the genetic correlation can be substantial but signifies little. For example, marriage hopes yields a genetic correlation of .61 with educational attainment but the phenotypic correlation is only .10 and it yields modest genetic correlations with other outcomes. These results are highly similar for males and females (Supplementary Figure S5a and b) and also similar when excluding opposite sex twin pairs (Supplementary Figure S5c).

**Figure 6.**
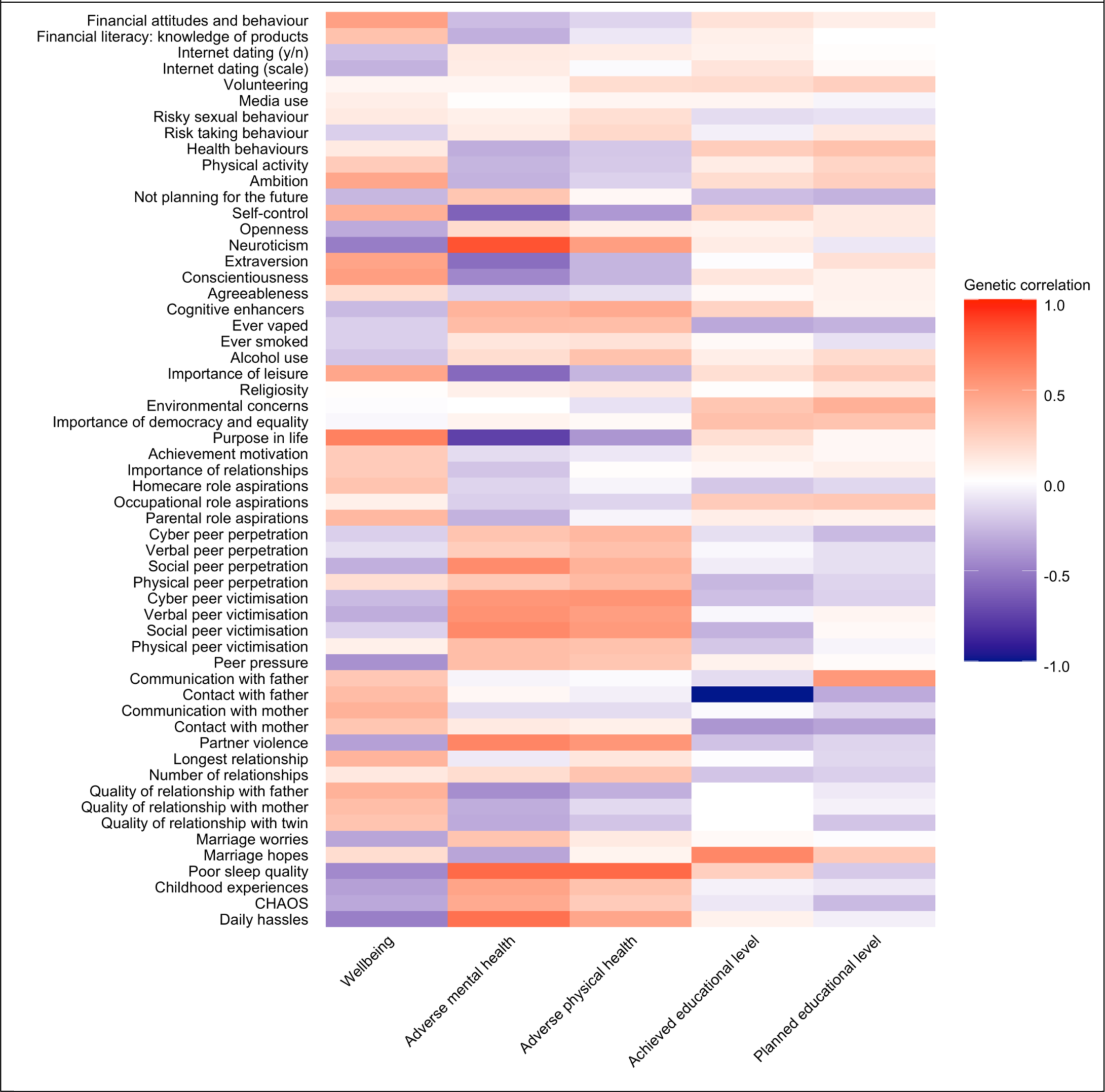
Genetic correlations between EA traits and outcomes derived from bivariate twin analyses

The shared environmental correlation (rC) and non-shared environmental correlation (rE) can also be estimated. The shared environmental correlations are illustrated in Supplementary Figure S6 and non-shared environmental correlations in Supplementary Figure S7. Supplementary Table S13 presents these estimates with 95% confidence intervals. None of the shared environmental correlations are statistically significant. Because shared environment contributes so little to the phenotypic correlations between EA traits and outcomes, the shared environmental correlations can be large, but the 95% confidence intervals indicate that these estimates are not significant.

### Univariate GREML-GCTA

Although the sample size with genotyped DNA limited power to detect SNP heritability, for completeness we present these estimates in Supplementary Table S14, which indicates the sample sizes for each measure. The mean SNP heritability was 10%, ranging up to 26%, as illustrated in Supplementary Figure S8. In addition, we conducted bivariate GREML analysis, again for completeness (Supplementary Table S15). However, little confidence can be placed in these results due to limited power.

### Sensitivity analyses

Univariate twin and GREML analyses reported above were repeated with van der Waerden transformed variables. The results of these analyses are presented in the Supplementary Materials (Figure S9 for univariate twin analyses; Figure S10 for univariate GREML analyses). Results do not differ with and without transformation.

## Discussion

Societal changes have stretched the transition from adolescence to adulthood, which has created a new developmental period called emerging adulthood. The prolonged uncertainties about identity, relationships and occupations can create stress but they also offer flexibility and unprecedented opportunities for young people to find their adult selves. These opportunities are amplified by the internet and social media for this first generation of digital natives.

As mentioned in the Introduction, this ever-expanding menu of idiosyncratic experiences for self-exploration during emerging adulthood suggests that environmental influence is likely to be critical for emerging adulthood traits. The present results from this first systematic analysis of emerging adulthood traits in a genetically sensitive design are consistent with this hypothesis in that the environment is the major source of individual differences in EA traits. Moreover, more than 90% of the environmental influence is due to non-shared environment, which includes measurement error. Non-shared environment could reflect idiosyncratic experiences in the chaotic world of emerging adulthood, experiences not likely to be shared by siblings. This non-shared environment is especially relevant when examining the differences between MZ twin pairs, who share their genes and home environment, are the same age and sex. Shared environmental influences account for only 5% of the variance of the 57 EA traits on average. In other words, most environmental effects on EA traits are not shared by siblings, even twins, who grow up in the same family. As siblings leave the family home in early adulthood, it seems likely that they each make their own independent way in the world, having trait-relevant experiences not shared with their sibling. Analyses investigating the environmental experiences that explain individual differences in emerging adulthood are part of our future research program using the longitudinal data collected at TEDS.

Another possibility is that the heterogeneity of experience in early adulthood provides an opportunity for genetic potential to be realised as individuals select, modify and create environments correlated with their genetic propensities. This is gene-environment correlation, which contributes to heritability. In support of this hypothesis, we found significant genetic influence for all 57 EA traits, many of which have not been investigated previously in emerging adulthood. On average, genetic influences accounted for 34% of the variance. Moreover, the most highly heritable traits in our study are traits that are especially relevant to emerging adulthood: risky sexual behaviour (60% heritability), general risk-taking behaviour (50%), media use (51%) and concerns about the environment (53%). An additional possibility is gene-environment interactions, individual react differentially to environmental influences and these particular resilience or vulnerability to environmental stressors or opportunities can be partly explained by genetic factors.

Although our results can be viewed as providing support for both hypotheses, it is also possible to interpret the results as support for neither hypothesis because both moderate genetic influence and substantial non-shared environmental influence are found for nearly all traits throughout development. Two of the ten most replicated findings from behavioural genetics are the findings that all psychological traits show significant and substantial genetic influence and most environmental effects are not shared by children growing up in the same family (Plomin, DeFries, Knopik, & Neiderhiser, 2016). Nonetheless, at the least it is useful to confirm these findings in emerging adulthood and especially for the many traits that have not been investigated previously at this age.

Moreover, it is possible that some EA traits show greater genetic or non-shared environmental influence in emerging adulthood as compared to earlier ages. Because many of these traits have not been studied previously in genetically sensitive designs, precise comparisons are not possible. One exception in TEDS is personality – the same measure of the Big Five personality traits was administered at age 16 and ages 21-25. The heritability of the Big Five personality traits was around 30% at age 16 (Rimfeld, Kovas, Dale, & Plomin, 2016) and 33% at age 21. This slight increase in heritability is not statistically significant and none of the five individual traits yielded significantly greater heritability at age 21 than at age 16. However, personality is highly stable across development (Turkheimer et al., 2014) and may be the least likely domain to show developmental changes in heritability.

TEDS also has some relevant data from a study of the psychological effects of COVID-19 (Rimfeld et al., 2020). In a subsample of TEDS, nine EA traits from the present study were assessed using the same measures two years later. For these EA traits, the average heritability was 34% at both ages. However, this study also found that change across the two-year period was significantly heritable (15% on average), suggesting that genetic influences change even during this two-year period in the early years of emerging adulthood. It remains to be seen whether the heritability of EA traits increases as the sample fully emerges into what has been called *established adulthood* (Mehta, Arnett, Palmer, & Nelson, 2020). The importance of non-shared environment in emerging adulthood is clear: it accounted for 82% of the variance on average for change across the two-year period.

Our finding of high heritability (50 to 60%) for traits most relevant to emerging adulthood and the evidence for genetic change during emerging adulthood leads us to predict that the heritability of EA traits will increase as we follow the twins into established adulthood. In any case, it seems that the turmoil of emerging adulthood does not shuffle the deck of individual differences. One firm conclusion is that heritability is not swamped by the environmental heterogeneity of emerging adulthood.

Our results also indicate that EA traits are associated with wellbeing, adverse mental and adverse physical health and educational outcomes. Although the associations were modest to moderate for individual EA traits and outcomes, the EA traits acted additively and together accounted for 31% of the variance in the outcomes on average. We show that adverse mental and physical health, wellbeing and educational attainment reflect a package of heritable EA traits. These findings also indicate that when studying individual differences in health, wellbeing or socioeconomic phenotypes in emerging adulthood, diverse traits should be considered such as life goals and aspirations and relationships. Results from bivariate genetic analyses show that these correlations are mediated mostly by genetic factors. These findings underscore the adage that correlation does not imply causation. For example, perceived daily hassles correlates negatively with wellbeing and positively with adverse mental and adverse physical health, but it cannot be assumed that daily hassles cause wellbeing or health.

Compared to health and wellbeing outcomes, we explained much less variance in educational outcomes (or hopes/aspirations for educational outcomes), compared to our previous reports in earlier childhood or even late adolescence (Krapohl et al., 2014). This can be explained in terms of the precision of the phenotype. Here we use achieved educational outcome, which is basically years of education, while previously we have used much finer-grained measures of educational achievement, such as teacher-graded performance or standardised examination results. However, similar to our previous report, we show that educational achievement and attainment can be partially explained by a range of behavioural and psychological traits, such as personality, poor sleep quality or risk-taking, all of which are also substantially heritable.

The limitations of this study include the usual limitations of twin design, which are described in detail elsewhere (Knopik et al., 2017; Rijsdijk & Sham, 2002). In addition, attrition and selection bias might have influenced the results. It is well documented that participants in most studies tend to be slightly healthier and more educated than the general population, and therefore, the associations observed between study variables might be biased by collider effects (Munafò, Tilling, Taylor, Evans, & Smith, 2018). However, we have shown that the TEDS twins in their early twenties remain reasonably representative of their birth cohort in terms of ethnicity and socioeconomic status (Rimfeld et al., 2019). An additional limitation is missing data and here we only used twins having all available datapoints. Missingness was minimal per data collection phase (e.g., it was less than 4% for the wellbeing composite) but was larger when both phase 1 and phase 2 data were used (e.g., around 18% for physical health). The representativeness of the sample, however, was very similar for phase 1 and phase 2 data collection. We repeated the analyses using mean scores of wellbeing, mental and physical health as a sensitivity analyses, allowing for missing data, and the patterns of associations and the aetiology of these composite measures remained highly similar. A conceptual limitation is that our EA traits were assessed at the same time as adverse mental and adverse physical health, wellbeing and education. Although we view the latter variables as outcomes, we acknowledge that longitudinal assessment of these outcomes is needed to investigate the extent to which EA traits predict them, which is part of our research plan for TEDS.

We conclude that the environmental uncertainties of emerging adulthood in the 21^st^ century do not diminish the importance of genetics. As adolescents travel down long and winding roads to adulthood, their trip is substantially influenced by genetic proclivities that nudge them down different paths that lead to different destinations.

## Acknowledgements

We gratefully acknowledge the ongoing contribution of the participants in the Twins Early Development Study (TEDS) and their families. TEDS is supported by a program grant to RP from the UK Medical Research Council (MR/M021475/1 and previously G0901245), with additional support from the US National Institutes of Health (AG046938). RP is supported by a Medical Research Council Professorship award (G19/2). KR is supported by a Sir Henry Wellcome Postdoctoral Fellowship. This study presents independent research [part-] funded by the National Institute for Health Research (NIHR) Biomedical Research Centre at South London and Maudsley NHS Foundation Trust and King’s College London. The views expressed are those of the author(s) and not necessarily those of the NHS, the NIHR or the Department of Health.

## Supporting information

### Appendix S1

#### Quality control

Two methods of identifying likely “clickers” are used in the TEDS-21 twin questionnaires:

1. *Quality control item error with uniform responding*. For a measure containing a quality control
2. (QC) item, if a twin’s response in the QC item was incorrect *and* the twin gave the same response in at least 3 of the adjacent 4 items within the measure, then the twin was labelled as a clicker and the theme was excluded. This definition could be applied in paper questionnaires as well as in web/app questionnaires. The rule had to be modified in some measures depending on the nature of the questions, etc.
3. *Quality control item error with rapid responding*. For a measure containing a QC item, if a twin’s response in the QC item was incorrect *and* the mean time taken per item across the entire theme was in the lowest 20%-ile of the distribution, then the twin was labelled as a clicker and the theme was excluded. This definition could only be applied for twins using web/app, where times could be measured, not for twins using paper. This rule was relaxed in only one theme where different conditions seemed to apply.

Hence, twins responding on paper could be excluded by the first criterion but not the second.

The rules above could not be applied uniformly across all themes in the two questionnaires. Some themes had no measures with QC items (hence no rule could be applied), some themes had one measure with a QC item, and some themes had more than one measure with a QC item.

The % excluded varies according to exclusion method, measure and theme. Below is an approximation of the % of twins excluded:

- Uniform responding (with QC item error): typically around 1% of twins were excluded per measure, but this varied from .01% to 6% in different measures.
- Rapid responding (with QC item error): typically around 1% of twins were excluded per measure, but this varied from .5% to 3%.
- Overall per theme: typically around 1% of twins were excluded per theme, but this varied from .3% to 5.3% (and of course 0% in themes without any measure containing a QC item).
- Overall exclusion for the entire questionnaire (questionnaire excluded if two or more themes excluded): 1.7% of twins in phase 1,.5% of twins in phase 2.
- Overall % of clickers (twin deemed to be a clicker and hence excluded from at least one theme): 5.5% of twins in phase 1, 4.4% of twins in phase 2.

As well as the varying number of measures with QC items, other factors caused variations in the percentages. For example, QC item errors were more likely in measures that were long, repetitive and perhaps “boring” to twins. The use of QC items was less well planned in phase 1, with the result that they were unevenly spread through the questionnaire and, in certain measures, the % of QC errors was higher than perhaps it should have been.

#### Supplementary Tables

**Table S1.** Summary of measured variables used

| Theme | Variable name | Scale/item<sup>a</sup> | No. of items used | Reference<sup>b</sup> |
| --- | --- | --- | --- | --- |
| **Emerging adulthood trait variables** |  |  |  |  |
| **Life experiences** | Daily hassles | Hassles | 7 | Delongis, A., Folkman, S. and Lazarus, R. S. (1988) |
|  | CHAOS | CHAOS<br>(Confusion, Hubbub and Order Scale) at home | 6 | Matheny, Wachs, Ludwig and Phillips (1995) |
|  | Childhood experiences | Childhood experiences | 8 | Taken from the ALSPAC "Life at 22+" questionnaire (section H) with some minor changes in wording:<br>http://www.bristol.ac.uk/media-library/sites/alspac/documents/questionnaires/YPB-life-at-22-plus.pdf |
|  | Sleep quality | The Pittsburgh Sleep Quality Index (PSQI) | 8 | Buyse D.J., Reynolds C.F., Monk T.H., Berman S.R. & Kupfer D.J. (1989) |
| **Relationships** | Marriage hopes | Marriage hopes | 4 | Kahn, S. (2007) |
|  | Marriage worries | Marriage worries | 4 | Kahn, S. (2007) |
|  | Quality of relationship with twin | Relationship with twin | 5 | Riggio, H. R. (2000) |
|  | Quality of relationship with mother | Relationship with mother | 5 | Fraley, R. C., Heffernan, M. E., Vicary, A. M., & Brumbaugh, C. C. (2011) |
|  | Quality of relationship with father | Relationship with father | 5 | Fraley, R. C., Heffernan, M. E., Vicary, A. M., & Brumbaugh, C. C. (2011) |
|  | Number of relationships | Number of relationships | 1 | Item created by the TEDS team. |
|  | Longest relationship | Longest relationship | 1 | Item created by the TEDS team. |
|  | Partner violence | Intimate Partner Violence Questions | 6 | Adapted from the 'Intimate Partner Violence Questions' section of the CDC<br>(Centers for Disease Control and Prevention) Violence Prevention questionnaire: |
|  |  |  |  | https://www.cdc.gov/violenceprevention/pdf/ipv/ipvandsvscreening.pdf |
|  | Contact with mother | Parent contact | 3 | Items created by TEDS researchers. |
|  | Communication with mother | Parent contact | 2 | Items created by TEDS researchers. |
|  | Contact with father | Parent contact | 3 | Items created by TEDS researchers. |
|  | Communication with father | Parent contact | 2 | Items created by TEDS researchers. |
|  | Peer pressure | Peer pressure | 7 | Shortened from Santor, D. A., Messervey, D., & Kusumakar, V. (2000) |
|  | Physical peer victimisation | Multidimensional Peer Victimization Scale | 4 | Mynard, H., and Joseph (2000) |
|  | Social peer victimisation | Multidimensional Peer Victimization Scale | 4 | Mynard, H., and Joseph (2000) |
|  | Verbal peer victimisation | Multidimensional Peer Victimization Scale | 4 | Mynard, H., and Joseph (2000) |
|  | Cyber peer victimisation | Multidimensional Peer Victimization Scale | 4 | Mynard, H., and Joseph (2000) |
|  | Physical peer Perpetration | Multidimensional Peer Victimization Scale | 4 | Mynard, H., and Joseph (2000) |
|  | Social peer Perpetration | Multidimensional Peer Victimization Scale | 4 | Mynard, H., and Joseph (2000) |
|  | Verbal peer Perpetration | Multidimensional Peer Victimization Scale | 4 | Mynard, H., and Joseph (2000) |
|  | Cyber peer Perpetration | Multidimensional Peer Victimization Scale | 4 | Mynard, H., and Joseph (2000) |
| **Aspirations** | Parental role aspirations | Life Role Salience (LRSS), Parental role | 4 | Amatea, E.S., Cross, E.G., Clark, J.E. and Bobby, C.L. (1986) |
|  | Occupational role aspirations | Life Role Salience (LRSS), Occupational role | 4 | Amatea, E.S., Cross, E.G., Clark, J.E. and Bobby, C.L. (1986) |
|  | Homecare role aspirations | Life Role Salience (LRSS), Homecare role | 4 | Amatea, E.S., Cross, E.G., Clark, J.E. and Bobby, C.L. (1986) |
| **Thoughts and attitudes** | Importance of Relationships | GOALS – relationships | 5 | Pöhlmann and Brunstein (1997) |
|  | Achievement Motivation | GOALS – self-fulfilment | 4 | Pöhlmann and Brunstein (1997) |
|  | Purpose in Life | Purpose in Life | 5 | Crumbaugh and Maholick (1964) |
|  | Importance of democracy and equality | BSA Democracy and Government | 5 | Taken from the NatCen Social Research's British Social Attitudes (BSA) survey (http://natcen.ac.uk/our-research/research/british-social-attitudes/). |
|  | Environmental concerns | BSA Environmental concerns | 6 | Taken from the NatCen Social Research's British Social Attitudes (BSA) Survey (http://natcen.ac.uk/our-research/research/british-social-attitudes/). |
|  | Religiosity | Religiosity: the Centrality of Religiosity Scale (CRS) | 5 | Huber, S., & Huber, O. W. (2012) |
|  | Importance of leisure | Leisure and Hobbies | 5 | Lavallee, L. F., Hatch, P. M., Michalos, A. C., & McKinley, T. (2007). |
| **Substance use** | Alcohol use | Alcohol Use Disorders Identification Test (AUDIT) scale | 10 | Adapted from items 1-10 of the AUDIT scale: https://www.drugabuse.gov/sites/default/files/files/AUDIT.pdf |
|  | Ever smoked | Smoking (inc. vaping) | 1 | Adapted from Heatherton, Kozlowski, Frecker, and Fagerstrom (1991) |
|  | Ever vaped | Smoking (inc. vaping) | 1 | Adapted from Heatherton, Kozlowski, Frecker, and Fagerstrom (1991) |
|  | Cognitive enhancers | Cognitive Enhancers | 3 | Items created by the TEDS team. |
| **Personality** | Big 5 personality | Big 5 Personality scale (Agreeableness, conscientiousness, extraversion, neuroticism, and openness subscales). | 30 | Mullins-Sweatt, S. N., Jamerson, J. E., Samuel, D. B., Olson, D. R., Widiger, T. A. (2006). |
|  | Self-control | Self-Control. | 6 | Tangney, J. P., Boone, A. L., & Baumeister, R. F. (2018) |
|  | Not planning for the future | Consideration of Future Consequences Scale. | 4 | Strathman, A., Gleicher, F., Boninger, D. S., & Edwards, C. S. (1994). |
|  | Ambition | Grit: Perseverance and passion for long-term goals. | 5 | Duckworth, A.L., Peterson, C., Matthews, M.D., & Kelly, D.R. (2007) |
| **Behaviours** | Physical activity | Activity questionnaire | 3 | Questions devised by TEDS researchers. |
|  | Health behaviours | Rapid Eating and Activity Assessment for Patients<br>(REAP) Health Behaviours | 12 | Adapted from Gans, K. M., Risica, P. M., Wylie-Rosett, J., Ross, E. M., Strolla, L. O., McMurray, J., & Eaton, C. B. (2006) |
|  | Risk taking behaviour | Risk taking index | 6 | Nicholson, N., Soane, E., Fenton-O'Creevy, M., & Willman, P. (2005) |
|  | Risky sexual behaviour | Sexual Behaviours and Experience | 5 | Adapted from the ALSPAC measure:<br>http://www.bristol.ac.uk/media-library/sites/alspac/documents/questionnaires/YPC-me-at-23.pdf |
|  | Media use | Media and Technology Usage and Attitudes Scale | 4 | Rosen, Whaling, Carrier, Cheever, & Rökkum (2013) |
|  | Volunteering | Volunteering | 5 | Adapted from page 42 of the ALSPAC questionnaire "It's all about you" at age 20: http://www.bristol.ac.uk/media-library/sites/alspac/migrated/documents/ques-yp20-all-about-you.pdf |
|  | Internet dating | Internet dating | 2 | Items created by TEDS researchers, in order to understand the proportion of emerging adults that take part in internet dating. |
| **Financial attitudes and behaviour** | Financial literacy: Knowledge of Products | OECD Financial literacy/products | 13 | Adapted from sections of the OECD instrument for measuring financial literacy:<br>https://www.oecd.org/finance/financial-education/49319977.pdf |
|  | Financial attitudes and behaviour | OECD Attitudes towards money | 6 | Adapted from sections of the OECD instrument for measuring financial literacy:<br>https://www.oecd.org/finance/financial-education/49319977.pdf |
| **Wellbeing variables** |  |  |  |  |
| - | Love and Relationships | CLAS, Love and Relationships | 3 | Lavallee, Hatch, Michalos, and McKinley (2007) |
| - | Community satisfaction | CLAS Life Satisfaction Scale, Community. | 5 | Lavallee, Hatch, Michalos, and McKinley (2007) |
| - | Financial wellbeing | CLAS Financial wellbeing | 5 | Lavallee, Hatch, Michalos, and McKinley (2007) |
| **Mental health variables** |  |  |  |  |
| - | Conners | Conners, 3rd edition. Impulsivity and inattention subscales. | 20 | Conners, C. K. 2008) |
| - | Aggression | Brief Aggression Questionnaire (BAQ). | 8 | Webster, G. D., DeWall, C. N., Pond Jr, R. S., Deckman, T., Jonason, P. K., Le, B. M., ... & Smith, C. V. (2014) |
| - | Conduct problems | SDQ, Conduct problems | 5 | Goodman (1997) |
| - | Emotional problems | SDQ, Emotional problems | 5 | Goodman (1997) |
| - | Hyperactivity | SDQ, Hyperactivity | 5 | Goodman (1997) |
| - | Peer problems | SDQ, Peer problems | 5 | Goodman (1997) |
| - | General anxiety | General Anxiety: the Severity Measure for Generalized Anxiety Disorder | 10 | Craske et al. (2013) |
| - | Depression | Short Mood and Feeling Questionnaire (SMFQ) | 8 | Adapted from Angold et al. (1995) |
| - | Self-harm | CASE, Self-harm | 2 | Adapted from Madge et al. (2008) |
| - | Psychotic experiences, paranoia | SPEQ: Specific Psychotic Experiences Questionnaire: Paranoia subscale | 12 | Ronald, A., Sieradzka, D., Cardno, A. G., Haworth, C. M. A, McGuire, P., Freeman, D. (2014) |
| - | Psychotic experiences, hallucinations | SPEQ: Specific Psychotic Experiences Questionnaire: Hallucinations subscale | 12 | Ronald, A., Sieradzka, D., Cardno, A. G., Haworth, C. M. A, McGuire, P., Freeman, D. (2014) |
| **Physical health variables** |  |  |  |  |
| - | Use of painkillers | Use of over-the-counter painkillers | 1 | Devised by TEDS researchers. |
| - | Use of antibiotics | Use of antibiotics | 1 | Devised by TEDS researchers. |
| - | General self-reported health | RAND Short-Form Health Survey | 5 | Adapted from Ware, J., Jr., & Sherbourne, C.D. The MOS 36-Item Short-Form Health Survey |
|  |  |  |  | (SF-36): I. Conceptual Framework and Item Selection, in Medical Care, June 1992, Vol. 30, No. 6, pp 473-483 |
| - | Hospitalisation | RAND Short-Form Health Survey. | 2 | Adapted from Ware, J., Jr., & Sherbourne, C.D. The MOS 36-Item Short-Form Health Survey (SF-36): I. Conceptual Framework and Item Selection, in Medical Care, June 1992, Vol. 30, No. 6, pp 473-483 |
| - | Short Health Economics | Short Health Economics.<br>Questions from the EQ-5D-5L measure. | 4 | Herdman M., Gudex, C., Lloyd, A. (2011). |
| **Education variables** |  |  |  |  |
| - | Achieved educational level | Educational level (current) | 1 | Questions devised by TEDS researchers. |
| - | Planned education level | Educational level (aspirational) | 1 | Questions devised by TEDS researchers. |

<sup>a</sup> More information about the variables and the references can be found in the TEDS data dictionary
(http://www.teds.ac.uk/datadictionary/studies/measures/21yr_measures.htm).
<sup>b</sup> Note that measures were shortened and adapted from the referenced measures.

**Table S2.** Descriptive statistics for psychological traits, for the whole and males and females separately.

| Trait (possible range) | All |  |  |  |  | Males |  |  |  |  | Females |  |  |  |  | ANOVA<sup>a</sup> |  |  |
| --- | --- | --- | --- | --- | --- | --- | --- | --- | --- | --- | --- | --- | --- | --- | --- | --- | --- | --- |
|  | *N* | Min | Max | *M* | *SD* | *n* | Min | Max | *M* | *SD* | *n* | Min | Max | *M* | *SD* | Sex | *p* | *R*<sup>2</sup> |
| Daily hassles (0-28) | 4324 | .00 | 28.00 | 12.54 | 4.63 | 1609 | .00 | 28.00 | 11.96 | 4.58 | 2715 | .00 | 28.00 | 12.88 | 4.62 | 4.40 | <.001 | .01 |
| CHAOS (0-12) | 4875 | .00 | 12.00 | 3.81 | 2.22 | 1860 | .00 | 12.00 | 3.73 | 2.05 | 3015 | .00 | 12.00 | 3.86 | 2.32 | 4.05 | .044 | .00 |
| Childhood experiences (0-32) | 4321 | .00 | 32.00 | 5.31 | 4.59 | 1607 | .00 | 32.00 | 5.27 | 4.11 | 2714 | .00 | 32.00 | 5.33 | 4.85 | .19 | .665 | .00 |
| Poor sleep quality (0-24) | 4250 | .00 | 24.00 | 4.85 | 3.84 | 1578 | .00 | 22.00 | 4.09 | 3.52 | 2672 | .00 | 24.00 | 5.29 | 3.96 | 104.74 | <.001 | .02 |
| Marriage hopes (1-5) | 4800 | 1.00 | 5.00 | 3.85 | .60 | 1829 | 1.00 | 5.00 | 3.85 | .59 | 2971 | 1.00 | 5.00 | 3.84 | .60 | .39 | .532 | .00 |
| Marriage worries (1-5) | 4801 | 1.00 | 5.00 | 2.37 | .68 | 1829 | 1.00 | 5.00 | 2.51 | .69 | 2972 | 1.00 | 5.00 | 2.29 | .66 | 126.53 | <.001 | .03 |
| Quality of relationship with twin (1-5) | 4710 | 1.00 | 5.00 | 3.94 | .88 | 1785 | 1.00 | 5.00 | 3.72 | .87 | 2925 | 1.00 | 5.00 | 4.07 | .86 | 179.85 | <.001 | .04 |
| Quality of relationship with mother (1-5) | 4658 | 1.00 | 5.00 | 3.83 | .96 | 1764 | 1.00 | 5.00 | 3.71 | .89 | 2894 | 1.00 | 5.00 | 3.91 | 1.00 | 52.03 | <.001 | .01 |
| Quality of relationship with father (1-5) | 4541 | 1.00 | 5.00 | 3.27 | 1.06 | 1716 | 1.00 | 5.00 | 3.26 | 1.01 | 2825 | 1.00 | 5.00 | 3.27 | 1.09 | .20 | .654 | .00 |
| Number of relationships (0-4) | 4798 | .00 | 4.00 | 1.22 | .80 | 1827 | .00 | 4.00 | 1.24 | .85 | 2971 | .00 | 4.00 | 1.20 | .76 | 2.47 | .116 | .00 |
| Longest relationship (1-5) | 4022 | 1.00 | 5.00 | 3.13 | 1.20 | 1510 | 1.00 | 5.00 | 2.91 | 1.20 | 2512 | 1.00 | 5.00 | 3.26 | 1.19 | 81.70 | <.001 | .02 |
| Partner violence (1-5) | 4642 | 1.00 | 5.00 | 1.77 | .96 | 1749 | 1.00 | 5.00 | 1.75 | .82 | 2893 | 1.00 | 5.00 | 1.78 | 1.03 | 1.39 | .239 | .00 |
| Contact with mother (1-6) | 1960 | 1.00 | 6.00 | 3.17 | 1.19 | 708 | 1.00 | 6.00 | 2.91 | 1.10 | 1252 | 1.00 | 6.00 | 3.31 | 1.22 | 54.86 | <.001 | .03 |
| Communication with mother (0-6) | 1999 | .00 | 6.00 | 4.72 | 1.13 | 722 | .00 | 6.00 | 4.33 | 1.10 | 1277 | .00 | 6.00 | 4.94 | 1.10 | 143.39 | <.001 | .07 |
| Communication with mother (1-6) | 2140 | 1.00 | 6.00 | 3.02 | 1.19 | 772 | 1.00 | 6.00 | 2.88 | 1.13 | 1368 | 1.00 | 6.00 | 3.11 | 1.21 | 19.06 | <.001 | .01 |
| Communication with father (0-6) | 2335 | .00 | 6.00 | 3.72 | 1.56 | 840 | .00 | 6.00 | 3.57 | 1.51 | 1495 | .00 | 6.00 | 3.80 | 1.59 | 12.40 | <.001 | .01 |
| Peer pressure (1-5) | 4745 | 1.00 | 5.00 | 1.96 | .72 | 1812 | 1.00 | 4.57 | 2.08 | .74 | 2933 | 1.00 | 5.00 | 1.90 | .70 | 7.32 | <.001 | .01 |
| Physical peer victimisation (0-8) | 4196 | .00 | 8.00 | .23 | .88 | 1566 | .00 | 8.00 | .39 | 1.12 | 2630 | .00 | 8.00 | .14 | .68 | 63.59 | <.001 | .02 |
| Social peer victimisation (0-8) | 4195 | .00 | 8.00 | .74 | 1.55 | 1566 | .00 | 8.00 | .70 | 1.47 | 2629 | .00 | 8.00 | .77 | 1.60 | 1.69 | .194 | .00 |
| Verbal peer victimisation (0-8) | 4195 | .00 | 8.00 | 2.03 | 2.53 | 1565 | .00 | 8.00 | 2.47 | 2.69 | 2630 | .00 | 8.00 | 1.77 | 2.39 | 71.07 | <.001 | .02 |
| Cyber peer victimisation (0-8) | 4195 | .00 | 8.00 | .59 | 1.43 | 1565 | .00 | 8.00 | .50 | 1.36 | 2630 | .00 | 8.00 | .63 | 1.47 | 8.82 | .003 | .00 |
| Physical peer Perpetration (0-8) | 4195 | .00 | 8.00 | .16 | .74 | 1564 | .00 | 8.00 | .32 | 1.06 | 2631 | .00 | 7.00 | .07 | .44 | 79.75 | <.001 | .03 |
| Social peer Perpetration (0-8) | 4195 | .00 | 8.00 | .42 | .90 | 1564 | .00 | 8.00 | .41 | .88 | 2631 | .00 | 8.00 | .43 | .92 | .88 | .349 | .00 |
| Verbal peer Perpetration (0-8) | 4195 | .00 | 8.00 | 1.53 | 2.14 | 1564 | .00 | 8.00 | 2.05 | 2.49 | 2631 | .00 | 8.00 | 1.22 | 1.84 | 129.84 | <.001 | .03 |
| Cyber peer Perpetration (0-8) | 4195 | .00 | 8.00 | .27 | .89 | 1564 | .00 | 8.00 | .29 | .95 | 2631 | .00 | 8.00 | .26 | .85 | 1.17 | .280 | .00 |
| Parental role aspirations (1-5) | 4334 | 1.00 | 5.00 | 4.18 | .76 | 1612 | 1.00 | 5.00 | 4.06 | .77 | 2722 | 1.00 | 5.00 | 4.25 | .74 | 62.14 | <.001 | .01 |
| Occupational role aspirations (1-5) | 4336 | 1.00 | 5.00 | 3.96 | .57 | 1614 | 1.00 | 5.00 | 3.94 | .58 | 2722 | 1.25 | 5.00 | 3.97 | .56 | 2.21 | .137 | .00 |
| Homecare role aspirations (1-5) | 4336 | 1.00 | 5.00 | 3.98 | .64 | 1614 | 1.00 | 5.00 | 3.86 | .68 | 2722 | 1.75 | 5.00 | 4.05 | .60 | 89.96 | <.001 | .02 |
| Importance of Relationships (0-20) | 4859 | .00 | 2.00 | 15.93 | 3.49 | 1860 | .00 | 2.00 | 14.81 | 3.79 | 2999 | .00 | 2.00 | 16.62 | 3.09 | 301.17 | <.001 | .06 |
| Achievement Motivation (0-16) | 4859 | .00 | 16.00 | 1.24 | 2.85 | 1860 | .00 | 16.00 | 1.16 | 3.00 | 2999 | .00 | 16.00 | 1.28 | 2.75 | 2.27 | .132 | .00 |
| Purpose in Life (1-5) | 4854 | 1.00 | 5.00 | 3.46 | .80 | 1858 | 1.00 | 5.00 | 3.38 | .81 | 2996 | 1.00 | 5.00 | 3.51 | .80 | 26.79 | <.001 | .01 |
| Importance of democracy and equality (1-5) | 4846 | 1.00 | 5.00 | 4.28 | .55 | 1854 | 1.00 | 5.00 | 4.20 | .58 | 2992 | 1.00 | 5.00 | 4.32 | .53 | 52.04 | <.001 | .01 |
| Environmental concerns (1-5) | 4851 | 1.00 | 5.00 | 3.72 | .69 | 1857 | 1.00 | 5.00 | 3.71 | .72 | 2994 | 1.00 | 5.00 | 3.73 | .67 | 1.06 | .303 | .00 |
| Religiosity (0-25) | 4751 | .00 | 25.00 | 4.43 | 4.98 | 1816 | .00 | 25.00 | 3.99 | 4.85 | 2935 | .00 | 25.00 | 4.70 | 5.04 | 23.80 | <.001 | .00 |
| Importance of leisure (1-5) | 4243 | 1.00 | 5.00 | 3.83 | .77 | 1569 | 1.00 | 5.00 | 3.94 | .75 | 2674 | 1.00 | 5.00 | 3.76 | .77 | 6.63 | <.001 | .01 |
| Alcohol use (0-40) | 3889 | .00 | 36.00 | 8.13 | 5.01 | 1461 | .00 | 36.00 | 8.76 | 5.29 | 2428 | .00 | 34.00 | 7.75 | 4.79 | 35.78 | <.001 | .01 |
| Ever smoked (0-1) | 4254 | .00 | 1.00 | .57 | .50 | 1579 | .00 | 1.00 | .57 | .50 | 2675 | .00 | 1.00 | .56 | .50 | .42 | .518 | .00 |
| Ever vaped (0-1) | 4254 | .00 | 1.00 | .27 | .44 | 1578 | .00 | 1.00 | .31 | .46 | 2676 | .00 | 1.00 | .24 | .43 | 24.64 | <.001 | .01 |
| Cognitive enhancers (0-12) | 4213 | .00 | 12.00 | 1.19 | 1.95 | 1560 | .00 | 12.00 | 1.42 | 2.14 | 2653 | .00 | 12.00 | 1.06 | 1.82 | 32.00 | <.001 | .01 |
| Big 5 Agreeableness (1-5) | 4708 | 1.00 | 5.00 | 3.71 | .57 | 1834 | 1.00 | 5.00 | 3.55 | .58 | 2874 | 1.33 | 5.00 | 3.81 | .54 | 246.80 | <.001 | .05 |
| Big 5 Conscientiousness (1-5) | 4702 | 1.00 | 5.00 | 3.87 | .62 | 1832 | 1.00 | 5.00 | 3.76 | .65 | 2870 | 1.17 | 5.00 | 3.93 | .59 | 78.99 | <.001 | .02 |
| Big 5 Extraversion (1-5) | 4726 | 1.00 | 5.00 | 3.50 | .66 | 1843 | 1.00 | 5.00 | 3.42 | .68 | 2883 | 1.00 | 5.00 | 3.55 | .63 | 4.33 | <.001 | .01 |
| Big 5 Neuroticism (1-5) | 4733 | 1.00 | 4.83 | 2.64 | .66 | 1847 | 1.00 | 4.67 | 2.51 | .63 | 2886 | 1.00 | 4.83 | 2.72 | .67 | 116.17 | <.001 | .02 |
| Big 5 Openness (1-5) | 4715 | 1.00 | 5.00 | 3.42 | .61 | 1838 | 1.00 | 5.00 | 3.39 | .63 | 2877 | 1.25 | 5.00 | 3.44 | .60 | 7.29 | .007 | .00 |
| Self-control (0-24) | 4699 | .00 | 24.00 | 14.33 | 4.20 | 1831 | .00 | 24.00 | 13.73 | 4.36 | 2868 | .00 | 24.00 | 14.72 | 4.05 | 6.13 | <.001 | .01 |
| Not planning for the future (1-5) | 4697 | 1.00 | 5.00 | 2.47 | .94 | 1831 | 1.00 | 5.00 | 2.64 | .95 | 2866 | 1.00 | 5.00 | 2.35 | .92 | 105.81 | <.001 | .02 |
| Ambition (0-20) | 4326 | .00 | 2.00 | 13.81 | 3.61 | 1610 | 1.00 | 2.00 | 13.80 | 3.74 | 2716 | .00 | 2.00 | 13.82 | 3.52 | .03 | .875 | .00 |
| Physical activity (1-5) | 4758 | 1.00 | 5.00 | 2.80 | 1.08 | 1804 | 1.00 | 5.00 | 2.93 | 1.10 | 2954 | 1.00 | 5.00 | 2.71 | 1.06 | 44.39 | <.001 | .01 |
| Health behaviours (0-48) | 4769 | 1.00 | 48.00 | 25.27 | 6.65 | 1811 | 1.00 | 48.00 | 24.81 | 6.75 | 2958 | 1.00 | 46.00 | 25.55 | 6.57 | 13.75 | <.001 | .00 |
| Risk taking behaviour (0-24) | 4697 | .00 | 23.00 | 6.51 | 3.58 | 1830 | .00 | 21.00 | 7.54 | 3.82 | 2867 | .00 | 23.00 | 5.86 | 3.25 | 244.01 | <.001 | .05 |
| Risky sexual behaviour (0-16) | 4554 | .00 | 12.67 | 4.22 | 2.75 | 1736 | .00 | 12.67 | 4.02 | 2.75 | 2818 | .00 | 12.00 | 4.34 | 2.75 | 14.52 | <.001 | .00 |
| Media use (0-20) | 4719 | .00 | 2.00 | 12.90 | 3.24 | 1794 | .00 | 2.00 | 11.99 | 3.53 | 2925 | .00 | 2.00 | 13.45 | 2.91 | 217.54 | <.001 | .05 |
| Volunteering (0-20) | 4754 | .00 | 19.00 | 5.65 | 3.24 | 1817 | .00 | 18.00 | 5.44 | 3.22 | 2937 | .00 | 19.00 | 5.77 | 3.24 | 11.97 | .001 | .00 |
| Internet dating (scale) (0-5) | 4706 | .00 | 5.00 | 2.79 | 2.17 | 1788 | .00 | 5.00 | 3.04 | 2.11 | 2918 | .00 | 5.00 | 2.64 | 2.19 | 39.25 | <.001 | .01 |
| Internet dating (y/n) (0-1) | 4717 | .00 | 1.00 | .46 | .50 | 1793 | .00 | 1.00 | .51 | .50 | 2924 | .00 | 1.00 | .44 | .50 | 21.50 | <.001 | .00 |
| Financial literacy: Knowledge of Products (0-4) | 4637 | .00 | 4.00 | 1.38 | .69 | 1778 | .00 | 4.00 | 1.55 | .73 | 2859 | .00 | 4.00 | 1.27 | .64 | 182.00 | <.001 | .04 |
| Financial attitudes and behaviour (1-5) | 4636 | 1.17 | 5.00 | 3.54 | .61 | 1778 | 1.17 | 5.00 | 3.48 | .64 | 2858 | 1.17 | 5.00 | 3.59 | .59 | 34.52 | <.001 | .01 |

*Note.* Raw uncorrected scores were used. These results are based on one twin randomly selected from each pair so that the data points are independent.
<sup>a</sup> *F* and *p* values are reported with Welch's correction (Welch, 1951) as the assumption of homogeneity of variance was violated for several variables.

**Table S3.** Descriptive statistics for psychological traits, for five sex and zygosity groups.

| Trait (possible range) | MZ males |  |  | DZ males |  |  | MZ females |  |  | DZ females |  |  | DZ opposite sex |  |  | ANOVA |  |  |
| --- | --- | --- | --- | --- | --- | --- | --- | --- | --- | --- | --- | --- | --- | --- | --- | --- | --- | --- |
|  | *n* | *M* | *SD* | *n* | *M* | *SD* | *n* | *M* | *SD* | *n* | *M* | *SD* | *n* | *M* | *SD* | Zygosity | Sex*<br>Zygosity | *R*<sup>2</sup><br>(adjusted) |
| Daily hassles (0-28) | 543 | 11.76 | 4.63 | 478 | 11.94 | 4.59 | 1055 | 12.88 | 4.69 | 893 | 12.97 | 4.64 | 1313 | 12.51 | 4.53 | .80 | .21 | .013 (.010) |
| CHAOS (0-12) | 606 | 3.69 | 2.09 | 555 | 3.80 | 2.10 | 1152 | 3.76 | 2.27 | 993 | 3.95 | 2.35 | 1514 | 3.76 | 2.17 | .29 | 3.65 | .004 (.001) |
| Childhood experiences (0-32) | 544 | 5.36 | 3.80 | 475 | 5.06 | 4.19 | 1052 | 5.50 | 5.22 | 892 | 5.23 | 4.52 | 1316 | 5.22 | 4.48 | .26 | .34 | .001 (-.002) |
| Poor sleep quality (0-24) | 534 | 4.04 | 3.70 | 470 | 4.03 | 3.29 | 1031 | 5.45 | 4.08 | 882 | 5.17 | 3.82 | 1291 | 4.75 | 3.78 | .00 | 1.24 | .025 (.022) |
| Marriage hopes (1-5) | 593 | 3.85 | .58 | 548 | 3.89 | .60 | 1134 | 3.85 | .61 | 978 | 3.82 | .58 | 1494 | 3.85 | .60 | 1.55 | .05 | .002 (-.001) |
| Marriage worries (1-5) | 593 | 2.49 | .68 | 548 | 2.50 | .66 | 1134 | 2.27 | .67 | 978 | 2.29 | .66 | 1495 | 2.40 | .70 | .03 | .04 | .033 (.030) |
| Quality of relationship with twin (1-5) | 580 | 4.07 | .79 | 537 | 3.57 | .84 | 1114 | 4.35 | .78 | 962 | 4.04 | .83 | 1465 | 3.64 | .89 | 8.22** | .00 | .106 (.103) |
| Quality of relationship with mother (1-5) | 569 | 3.68 | .93 | 534 | 3.73 | .87 | 1105 | 3.91 | 1.01 | 948 | 3.91 | .99 | 1452 | 3.84 | .95 | .19 | .14 | .009 (.006) |
| Quality of relationship with father (1-5) | 559 | 3.25 | 1.07 | 514 | 3.32 | .96 | 1075 | 3.25 | 1.11 | 923 | 3.24 | 1.09 | 1421 | 3.28 | 1.05 | .09 | .92 | .001 (-.002) |
| Number of relationships (0-4) | 593 | 1.18 | .86 | 546 | 1.28 | .84 | 1132 | 1.18 | .78 | 979 | 1.20 | .73 | 1495 | 1.26 | .81 | .00 | 2.41 | .005 (.003) |
| Longest relationship (1-5) | 473 | 2.94 | 1.18 | 467 | 2.97 | 1.22 | 939 | 3.26 | 1.18 | 839 | 3.26 | 1.22 | 1260 | 3.08 | 1.19 | .36 | .10 | .019 (.016) |
| Partner violence (1-5) | 569 | 1.73 | .82 | 523 | 1.74 | .84 | 1099 | 1.76 | 1.02 | 956 | 1.78 | 1.03 | 1442 | 1.79 | .94 | .04 | .03 | .001 (-.002) |
| Contact with mother (1-6) | 254 | 3.00 | 1.15 | 195 | 2.87 | 1.02 | 482 | 3.31 | 1.22 | 419 | 3.38 | 1.27 | 598 | 3.07 | 1.15 | .99 | .34 | .023 (.020) |
| Communication with mother (0-6) | 258 | 4.40 | 1.10 | 199 | 4.34 | 1.06 | 494 | 4.91 | 1.18 | 425 | 5.02 | 1.07 | 610 | 4.63 | 1.10 | .00 | .08 | .103 (.100) |
| Communication with mother (1-6) | 270 | 2.97 | 1.21 | 218 | 2.90 | 1.10 | 516 | 3.05 | 1.18 | 460 | 3.22 | 1.29 | 658 | 2.94 | 1.12 | .08 | 1.08 | .009 (.006) |
| Communication with father (0-6) | 297 | 3.62 | 1.57 | 233 | 3.64 | 1.46 | 571 | 3.73 | 1.61 | 498 | 3.90 | 1.60 | 718 | 3.65 | 1.53 | .26 | .13 | .019 (.016) |
| Peer pressure (1-5) | 585 | 1.98 | .71 | 546 | 2.06 | .74 | 1118 | 1.87 | .68 | 968 | 1.89 | .70 | 1479 | 2.04 | .74 | 3.50 | .53 | .015 (.012) |
| Physical peer victimisation (0-8) | 528 | .39 | 1.16 | 469 | .44 | 1.12 | 1020 | .14 | .70 | 869 | .11 | .55 | 1269 | .25 | .94 | .41 | .53 | .035 (.032) |
| Social peer victimisation (0-8) | 528 | .63 | 1.33 | 469 | .69 | 1.50 | 1020 | .74 | 1.58 | 869 | .78 | 1.60 | 1268 | .79 | 1.59 | .56 | .90 | .001 (-.001) |
| Verbal peer victimisation (0-8) | 528 | 2.28 | 2.59 | 468 | 2.57 | 2.71 | 1020 | 1.64 | 2.34 | 869 | 1.77 | 2.35 | 1269 | 2.23 | 2.64 | 2.56 | 1.18 | .036 (.034) |
| Cyber peer victimisation (0-8) | 528 | .49 | 1.35 | 468 | .54 | 1.37 | 1020 | .63 | 1.47 | 869 | .58 | 1.41 | 1269 | .60 | 1.47 | .03 | .40 | .001 (-.001) |
| Physical peer Perpetration (0-8) | 526 | .34 | 1.14 | 469 | .36 | 1.07 | 1020 | .07 | .44 | 870 | .07 | .42 | 1269 | .17 | .73 | 1.15 | 2.85 | .044 (.041) |
| Social peer Perpetration (0-8) | 526 | .40 | .83 | 469 | .40 | .94 | 1020 | .42 | .92 | 870 | .43 | .93 | 1269 | .43 | .88 | .01 | .02 | .000 (-.003) |
| Verbal peer Perpetration (0-8) | 526 | 1.92 | 2.45 | 469 | 2.01 | 2.48 | 1020 | 1.08 | 1.73 | 870 | 1.28 | 1.87 | 1269 | 1.74 | 2.26 | 2.75 | .00 | .040 (.038) |
| Cyber peer Perpetration (0-8) | 526 | .24 | .82 | 469 | .33 | 1.07 | 1020 | .25 | .85 | 870 | .23 | .76 | 1269 | .30 | .96 | 2.12 | .21 | .006 (.003) |
| Parental role aspirations (1-5) | 548 | 4.07 | .76 | 476 | 4.03 | .74 | 1057 | 4.27 | .71 | 895 | 4.25 | .73 | 1316 | 4.16 | .80 | .03 | .00 | .007 (.004) |
| Occupational role aspirations (1-5) | 548 | 3.94 | .58 | 476 | 3.91 | .60 | 1057 | 3.98 | .55 | 895 | 3.96 | .57 | 1318 | 3.97 | .57 | .08 | .10 | .001 (-.002) |
| Homecare role aspirations (1-5) | 548 | 3.90 | .68 | 476 | 3.81 | .69 | 1057 | 4.08 | .60 | 895 | 4.04 | .58 | 1318 | 3.96 | .65 | .45 | .20 | .012 (.010) |
| Importance of Relationships (0-20) | 603 | 14.74 | 3.90 | 556 | 14.85 | 3.73 | 1141 | 16.65 | 3.11 | 993 | 16.53 | 3.06 | 1512 | 15.87 | 3.51 | .96 | 1.31 | .060 (.057) |
| Achievement Motivation (0-16) | 603 | 1.26 | 3.01 | 556 | 1.06 | 3.06 | 1141 | 1.30 | 2.77 | 993 | 1.31 | 2.69 | 1512 | 1.20 | 2.87 | 1.87 | 3.96* | .004 (.002) |
| Purpose in Life (1-5) | 602 | 3.40 | .83 | 556 | 3.39 | .80 | 1139 | 3.52 | .80 | 993 | 3.48 | .78 | 1510 | 3.45 | .80 | .06 | .80 | .005 (.002) |
| Importance of democracy and equality (1-5) | 602 | 4.22 | .57 | 554 | 4.22 | .54 | 1135 | 4.31 | .51 | 992 | 4.31 | .54 | 1508 | 4.27 | .59 | 1.48 | .01 | .021 (.018) |
| Environmental concerns (1-5) | 602 | 3.70 | .73 | 556 | 3.74 | .70 | 1137 | 3.72 | .65 | 992 | 3.75 | .69 | 1509 | 3.70 | .70 | 1.36 | .10 | .002 (-.001) |
| Religiosity (0-25) | 586 | 4.28 | 5.30 | 548 | 4.07 | 4.83 | 1119 | 4.87 | 5.25 | 968 | 4.45 | 4.73 | 1480 | 4.22 | 4.79 | 1.00 | .27 | .004 (.001) |
| Importance of leisure (1-5) | 528 | 3.93 | .76 | 471 | 3.97 | .73 | 1031 | 3.72 | .77 | 885 | 3.75 | .78 | 1286 | 3.88 | .76 | 1.60 | 1.37 | .018 (.016) |
| Alcohol use (0-40) | 484 | 8.56 | 5.29 | 436 | 8.76 | 5.24 | 932 | 7.49 | 4.68 | 807 | 7.88 | 4.94 | 1195 | 8.41 | 5.05 | .11 | 1.63 | .008 (.005) |
| Ever smoked (0-1) | 530 | .52 | .50 | 471 | .58 | .49 | 1034 | .54 | .50 | 883 | .57 | .50 | 1295 | .61 | .49 | 2.33 | 1.01 | .006 (.003) |
| Ever vaped (0-1) | 530 | .30 | .46 | 472 | .29 | .45 | 1035 | .22 | .42 | 884 | .24 | .43 | 1292 | .30 | .46 | 1.76 | 2.09 | .005 (.003) |
| Cognitive enhancers (0-12) | 521 | 1.30 | 2.07 | 467 | 1.36 | 2.11 | 1029 | 1.07 | 1.88 | 876 | 1.05 | 1.76 | 1280 | 1.29 | 2.03 | .33 | .05 | .003 (.001) |
| Big 5 Agreeableness (1-5) | 598 | 3.56 | .55 | 551 | 3.55 | .58 | 1095 | 3.83 | .53 | 956 | 3.80 | .54 | 1459 | 3.68 | .59 | .57 | .18 | .046 (.044) |
| Big 5 Conscientiousness (1-5) | 598 | 3.83 | .62 | 550 | 3.74 | .67 | 1094 | 3.96 | .57 | 954 | 3.91 | .59 | 1457 | 3.83 | .63 | 2.05 | .35 | .012 (.009) |
| Big 5 Extraversion (1-5) | 601 | 3.39 | .70 | 552 | 3.44 | .67 | 1101 | 3.52 | .63 | 956 | 3.54 | .64 | 1466 | 3.53 | .65 | .00 | .13 | .010 (.008) |
| Big 5 Neuroticism (1-5) | 602 | 2.50 | .63 | 553 | 2.53 | .64 | 1104 | 2.73 | .66 | 956 | 2.73 | .67 | 1468 | 2.60 | .66 | .43 | .13 | .034 (.032) |
| Big 5 Openness (1-5) | 600 | 3.39 | .61 | 552 | 3.39 | .63 | 1098 | 3.42 | .60 | 956 | 3.45 | .60 | 1460 | 3.41 | .62 | 3.32 | .44 | .007 (.004) |
| Self-control (0-24) | 598 | 14.02 | 4.21 | 550 | 13.73 | 4.43 | 1092 | 14.92 | 4.00 | 954 | 14.47 | 4.12 | 1456 | 14.18 | 4.25 | 1.89 | 4.02* | .019 (.016) |
| Not planning for the future (1-5) | 598 | 2.60 | .95 | 550 | 2.64 | .94 | 1092 | 2.34 | .93 | 952 | 2.37 | .92 | 1456 | 2.51 | .95 | .21 | .12 | .021 (.018) |
| Ambition (0-20) | 544 | 13.93 | 3.69 | 478 | 13.65 | 3.84 | 1055 | 13.86 | 3.55 | 893 | 13.79 | 3.52 | 1314 | 13.81 | 3.61 | 2.32 | 1.36 | .003 (.001) |
| Physical activity (1-5) | 585 | 2.95 | 1.11 | 543 | 2.92 | 1.09 | 1129 | 2.67 | 1.07 | 970 | 2.68 | 1.03 | 1478 | 2.87 | 1.09 | .83 | .22 | .011 (.008) |
| Health behaviours (0-48) | 588 | 24.85 | 6.56 | 543 | 24.89 | 6.85 | 1129 | 25.41 | 6.62 | 973 | 25.49 | 6.48 | 1483 | 25.30 | 6.73 | .62 | .89 | .003 (.000) |
| Risk taking behaviour (0-24) | 598 | 7.26 | 3.86 | 548 | 7.44 | 3.73 | 1093 | 5.59 | 3.10 | 953 | 5.92 | 3.43 | 1457 | 6.96 | 3.61 | 5.71* | 2.78 | .046 (.044) |
| Risky sexual behaviour (0-16) | 561 | 3.91 | 2.85 | 522 | 4.02 | 2.71 | 1071 | 4.16 | 2.80 | 923 | 4.38 | 2.71 | 1427 | 4.37 | 2.71 | .93 | 9.93* | .018 (.015) |
| Media use (0-20) | 581 | 11.91 | 3.67 | 540 | 11.88 | 3.61 | 1122 | 13.48 | 2.89 | 958 | 13.45 | 2.82 | 1466 | 12.85 | 3.22 | 3.17 | 2.13 | .042 (.039) |
| Volunteering (0-20) | 586 | 5.57 | 3.24 | 548 | 5.51 | 3.19 | 1120 | 5.66 | 3.29 | 969 | 5.82 | 3.21 | 1481 | 5.61 | 3.21 | 1.86 | 6.48* | .010 (.008) |
| Internet dating (scale) (0-5) | 579 | 3.00 | 2.14 | 541 | 2.92 | 2.09 | 1116 | 2.57 | 2.19 | 959 | 2.59 | 2.21 | 1459 | 2.98 | 2.13 | .29 | .98 | .014 (.012) |
| Internet dating (y/n) (0-1) | 581 | .50 | .50 | 542 | .47 | .50 | 1120 | .42 | .49 | 960 | .43 | .50 | 1462 | .50 | .50 | .86 | 1.79 | .011 (.008) |
| Financial literacy: Knowledge of Products (0-4) | 582 | 1.60 | .75 | 539 | 1.56 | .73 | 1088 | 1.26 | .63 | 936 | 1.30 | .65 | 1440 | 1.38 | .67 | .79 | .79 | .046 (.043) |
| Financial attitudes and behaviour (1-5) | 582 | 3.53 | .60 | 539 | 3.45 | .65 | 1088 | 3.59 | .59 | 936 | 3.59 | .59 | 1439 | 3.52 | .63 | .20 | .80 | .018 (.015) |

*Note.* MZ=monozygotic; DZ=dizygotic. Raw uncorrected scores were used. These results are based on one twin randomly selected from each pair so that the data points are independent.
Group difference = *F* statistics; \*\**p* < .05. \*\*\**p* < .001.

**Table S4.** Descriptive statistics for psychological traits, for the whole and males and females separately (repeated with the twin that was not randomly selected from each pair in the main analyses).

| Trait (possible range) | All |  |  |  |  | Males |  |  |  |  | Females |  |  |  |  | ANOVA<sup>a</sup> |  |  |
| --- | --- | --- | --- | --- | --- | --- | --- | --- | --- | --- | --- | --- | --- | --- | --- | --- | --- | --- |
|  | *N* | Min | Max | *M* | *SD* | *n* | Min | Max | *M* | *SD* | *n* | Min | Max | *M* | *SD* | Sex | *p* | *R*<sup>2</sup> |
| Daily hassles (0-28) | 4327 | .00 | 28.00 | 12.58 | 4.70 | 1553 | .00 | 28.00 | 11.96 | 4.67 | 2774 | .00 | 28.00 | 12.92 | 4.68 | 4.40 | .000 | .01 |
| CHAOS (0-12) | 4856 | .00 | 12.00 | 3.85 | 2.22 | 1795 | .00 | 12.00 | 3.82 | 2.13 | 3061 | .00 | 12.00 | 3.87 | 2.27 | 4.05 | .044 | .00 |
| Childhood experiences (0-32) | 4323 | .00 | 32.00 | 5.34 | 4.56 | 1548 | .00 | 31.00 | 5.19 | 4.06 | 2775 | .00 | 32.00 | 5.43 | 4.81 | .19 | .665 | .00 |
| Poor sleep quality (0-24) | 4239 | .00 | 24.00 | 4.95 | 3.90 | 1515 | .00 | 21.00 | 4.10 | 3.49 | 2724 | .00 | 24.00 | 5.43 | 4.04 | 104.74 | .000 | .02 |
| Marriage hopes (1-5) | 4776 | 1.00 | 5.00 | 3.85 | .59 | 1772 | 1.00 | 5.00 | 3.84 | .58 | 3004 | 1.00 | 5.00 | 3.86 | .60 | .39 | .532 | .00 |
| Marriage worries (1-5) | 4775 | 1.00 | 5.00 | 2.38 | .69 | 1771 | 1.00 | 5.00 | 2.55 | .69 | 3004 | 1.00 | 5.00 | 2.29 | .67 | 126.53 | .000 | .03 |
| Quality of relationship with twin (1-5) | 4680 | 1.00 | 5.00 | 3.93 | .90 | 1722 | 1.00 | 5.00 | 3.73 | .87 | 2958 | 1.00 | 5.00 | 4.04 | .90 | 179.85 | .000 | .04 |
| Quality of relationship with mother (1-5) | 4646 | 1.00 | 5.00 | 3.83 | .96 | 1706 | 1.00 | 5.00 | 3.65 | .90 | 2940 | 1.00 | 5.00 | 3.93 | .97 | 52.03 | .000 | .01 |
| Quality of relationship with father (1-5) | 4539 | 1.00 | 5.00 | 3.21 | 1.08 | 1658 | 1.00 | 5.00 | 3.24 | .99 | 2881 | 1.00 | 5.00 | 3.20 | 1.13 | .20 | .654 | .00 |
| Number of relationships (0-4) | 4776 | .00 | 4.00 | 1.21 | .79 | 1772 | .00 | 4.00 | 1.25 | .85 | 3004 | .00 | 4.00 | 1.18 | .75 | 2.47 | .116 | .00 |
| Longest relationship (1-5) | 4024 | 1.00 | 5.00 | 3.09 | 1.22 | 1479 | 1.00 | 5.00 | 2.86 | 1.21 | 2545 | 1.00 | 5.00 | 3.22 | 1.21 | 81.70 | .000 | .02 |
| Partner violence (1-5) | 4618 | 1.00 | 5.00 | 1.77 | .97 | 1693 | 1.00 | 5.00 | 1.73 | .84 | 2925 | 1.00 | 5.00 | 1.79 | 1.03 | 1.39 | .239 | .00 |
| Contact with mother (1-6) | 1960 | 1.00 | 6.00 | 3.19 | 1.20 | 676 | 1.00 | 6.00 | 2.94 | 1.09 | 1284 | 1.00 | 6.00 | 3.32 | 1.23 | 54.86 | .000 | .03 |
| Communication with mother (0-6) | 2007 | .00 | 6.00 | 4.67 | 1.19 | 690 | .00 | 6.00 | 4.30 | 1.12 | 1317 | .00 | 6.00 | 4.86 | 1.18 | 143.39 | .000 | .07 |
| Communication with mother (1-6) | 2121 | 1.00 | 6.00 | 3.02 | 1.18 | 728 | 1.00 | 6.00 | 2.92 | 1.14 | 1393 | 1.00 | 6.00 | 3.07 | 1.20 | 19.06 | .000 | .01 |
| Communication with father (0-6) | 2328 | .00 | 6.00 | 3.70 | 1.55 | 788 | .00 | 6.00 | 3.62 | 1.43 | 1540 | .00 | 6.00 | 3.75 | 1.61 | 12.40 | .000 | .01 |
| Peer pressure (1-5) | 4725 | 1.00 | 4.86 | 1.97 | .71 | 1758 | 1.00 | 4.86 | 2.07 | .73 | 2967 | 1.00 | 4.86 | 1.91 | .70 | 7.32 | .000 | .01 |
| Physical peer victimisation (0-8) | 4187 | .00 | 8.00 | .22 | .86 | 1503 | .00 | 8.00 | .39 | 1.14 | 2684 | .00 | 8.00 | .12 | .63 | 63.59 | .000 | .02 |
| Social peer victimisation (0-8) | 4187 | .00 | 8.00 | .75 | 1.57 | 1503 | .00 | 8.00 | .72 | 1.52 | 2684 | .00 | 8.00 | .77 | 1.60 | 1.69 | .194 | .00 |
| Verbal peer victimisation (0-8) | 4186 | .00 | 8.00 | 2.07 | 2.53 | 1502 | .00 | 8.00 | 2.49 | 2.73 | 2684 | .00 | 8.00 | 1.84 | 2.39 | 71.07 | .000 | .02 |
| Cyber peer victimisation (0-8) | 4185 | .00 | 8.00 | .59 | 1.44 | 1503 | .00 | 8.00 | .57 | 1.44 | 2682 | .00 | 8.00 | .60 | 1.44 | 8.82 | .003 | .00 |
| Physical peer Perpetration (0-8) | 4186 | .00 | 8.00 | .16 | .76 | 1503 | .00 | 8.00 | .31 | 1.07 | 2683 | .00 | 8.00 | .07 | .49 | 79.75 | .000 | .03 |
| Social peer Perpetration (0-8) | 4184 | .00 | 8.00 | .42 | .88 | 1502 | .00 | 8.00 | .43 | .95 | 2682 | .00 | 8.00 | .42 | .83 | .88 | .349 | .00 |
| Verbal peer Perpetration (0-8) | 4185 | .00 | 8.00 | 1.54 | 2.12 | 1503 | .00 | 8.00 | 2.08 | 2.49 | 2682 | .00 | 8.00 | 1.24 | 1.81 | 129.84 | .000 | .03 |
| Cyber peer Perpetration (0-8) | 4185 | .00 | 8.00 | .27 | .92 | 1503 | .00 | 8.00 | .31 | 1.02 | 2682 | .00 | 8.00 | .25 | .85 | 1.17 | .280 | .00 |
| Parental role aspirations (1-5) | 4328 | 1.00 | 5.00 | 4.16 | .77 | 1551 | 1.00 | 5.00 | 4.04 | .76 | 2777 | 1.00 | 5.00 | 4.23 | .76 | 62.14 | .000 | .01 |
| Occupational role aspirations (1-5) | 4331 | 1.00 | 5.00 | 3.95 | .61 | 1551 | 1.00 | 5.00 | 3.96 | .60 | 2780 | 1.00 | 5.00 | 3.94 | .61 | 2.21 | .137 | .00 |
| Homecare role aspirations (1-5) | 4331 | 1.00 | 5.00 | 4.00 | .63 | 1551 | 1.00 | 5.00 | 3.91 | .65 | 2780 | 1.25 | 5.00 | 4.04 | .61 | 89.96 | .000 | .02 |
| Importance of Relationships (0-20) | 4822 | .00 | 2.00 | 15.92 | 3.56 | 1790 | .00 | 2.00 | 14.88 | 3.87 | 3032 | .00 | 2.00 | 16.53 | 3.21 | 301.17 | .000 | .06 |
| Achievement Motivation (0-16) | 4821 | .00 | 16.00 | 1.19 | 2.91 | 1789 | .00 | 16.00 | 1.17 | 3.09 | 3032 | .00 | 16.00 | 1.21 | 2.80 | 2.27 | .132 | .00 |
| Purpose in Life (1-5) | 4819 | 1.00 | 5.00 | 3.47 | .82 | 1788 | 1.00 | 5.00 | 3.42 | .82 | 3031 | 1.00 | 5.00 | 3.49 | .81 | 26.79 | .000 | .01 |
| Importance of democracy and equality (1-5) | 4808 | 1.00 | 5.00 | 4.28 | .54 | 1785 | 1.00 | 5.00 | 4.19 | .56 | 3023 | 1.00 | 5.00 | 4.33 | .51 | 52.04 | .000 | .01 |
| Environmental concerns (1-5) | 4815 | 1.00 | 5.00 | 3.70 | .70 | 1787 | 1.00 | 5.00 | 3.64 | .74 | 3028 | 1.17 | 5.00 | 3.73 | .67 | 1.06 | .303 | .00 |
| Religiosity (0-25) | 4730 | .00 | 25.00 | 4.36 | 4.96 | 1762 | .00 | 25.00 | 3.91 | 4.91 | 2968 | .00 | 25.00 | 4.64 | 4.97 | 23.80 | .000 | .00 |
| Importance of leisure (1-5) | 4243 | 1.00 | 5.00 | 3.82 | .76 | 1518 | 1.00 | 5.00 | 3.93 | .71 | 2725 | 1.00 | 5.00 | 3.76 | .77 | 6.63 | .000 | .01 |
| Alcohol use (0-40) | 3899 | .00 | 37.00 | 8.05 | 5.06 | 1425 | .00 | 32.00 | 8.79 | 5.29 | 2474 | .00 | 37.00 | 7.61 | 4.88 | 35.78 | .000 | .01 |
| Ever smoked (0-1) | 4262 | .00 | 1.00 | .56 | .50 | 1528 | .00 | 1.00 | .57 | .50 | 2734 | .00 | 1.00 | .56 | .50 | .42 | .518 | .00 |
| Ever vaped (0-1) | 4259 | .00 | 1.00 | .25 | .44 | 1524 | .00 | 1.00 | .29 | .45 | 2735 | .00 | 1.00 | .24 | .43 | 24.64 | .000 | .01 |
| Cognitive enhancers (0-12) | 4217 | .00 | 12.00 | 1.19 | 1.96 | 1506 | .00 | 12.00 | 1.38 | 2.16 | 2711 | .00 | 12.00 | 1.08 | 1.83 | 32.00 | .000 | .01 |
| Big 5 Agreeableness (1-5) | 4674 | 1.00 | 5.00 | 3.72 | .57 | 1760 | 1.00 | 5.00 | 3.55 | .57 | 2914 | 1.00 | 5.00 | 3.82 | .55 | 246.80 | .000 | .05 |
| Big 5 Conscientiousness (1-5) | 4666 | 1.00 | 5.00 | 3.88 | .62 | 1757 | 1.00 | 5.00 | 3.78 | .64 | 2909 | 1.00 | 5.00 | 3.94 | .60 | 78.99 | .000 | .02 |
| Big 5 Extraversion (1-5) | 4694 | 1.00 | 5.00 | 3.52 | .65 | 1772 | 1.00 | 5.00 | 3.45 | .68 | 2922 | 1.00 | 5.00 | 3.56 | .63 | 4.33 | .000 | .01 |
| Big 5 Neuroticism (1-5) | 4700 | 1.00 | 5.00 | 2.62 | .67 | 1774 | 1.00 | 5.00 | 2.48 | .66 | 2926 | 1.00 | 5.00 | 2.70 | .66 | 116.17 | .000 | .02 |
| Big 5 Openness (1-5) | 4688 | 1.00 | 5.00 | 3.40 | .62 | 1769 | 1.00 | 5.00 | 3.37 | .64 | 2919 | 1.00 | 5.00 | 3.42 | .61 | 7.29 | .007 | .00 |
| Self-control (0-24) | 4661 | .00 | 24.00 | 14.46 | 4.21 | 1753 | .00 | 24.00 | 13.86 | 4.30 | 2908 | .00 | 24.00 | 14.83 | 4.11 | 6.13 | .000 | .01 |
| Not planning for the future (1-5) | 4655 | 1.00 | 5.00 | 2.47 | .92 | 1749 | 1.00 | 5.00 | 2.58 | .92 | 2906 | 1.00 | 5.00 | 2.40 | .91 | 105.81 | .000 | .02 |
| Ambition (0-20) | 4328 | 1.00 | 2.00 | 13.85 | 3.61 | 1553 | 1.00 | 2.00 | 14.00 | 3.67 | 2775 | 1.00 | 2.00 | 13.77 | 3.58 | .03 | .875 | .00 |
| Physical activity (1-5) | 4699 | 1.00 | 5.00 | 2.83 | 1.09 | 1733 | 1.00 | 5.00 | 2.96 | 1.10 | 2966 | 1.00 | 5.00 | 2.75 | 1.08 | 44.39 | .000 | .01 |
| Health behaviours (0-48) | 4721 | 2.00 | 48.00 | 25.24 | 6.67 | 1742 | 4.00 | 48.00 | 24.61 | 6.63 | 2979 | 2.00 | 47.00 | 25.61 | 6.66 | 13.75 | .000 | .00 |
| Risk taking behaviour (0-24) | 4651 | .00 | 24.00 | 6.49 | 3.61 | 1748 | .00 | 21.00 | 7.46 | 3.74 | 2903 | .00 | 24.00 | 5.91 | 3.40 | 244.01 | .000 | .05 |
| Risky sexual behaviour (0-16) | 4568 | .00 | 14.00 | 4.17 | 2.74 | 1706 | .00 | 12.33 | 3.92 | 2.71 | 2862 | .00 | 14.00 | 4.32 | 2.76 | 14.52 | .000 | .00 |
| Media use (0-20) | 4686 | .00 | 2.00 | 12.98 | 3.23 | 1732 | .00 | 2.00 | 12.04 | 3.66 | 2954 | .00 | 2.00 | 13.54 | 2.79 | 217.54 | .000 | .05 |
| Volunteering (0-20) | 4735 | .00 | 2.00 | 5.66 | 3.23 | 1763 | .00 | 18.00 | 5.52 | 3.32 | 2972 | .00 | 2.00 | 5.74 | 3.17 | 11.97 | .001 | .00 |
| Internet dating (scale) (0-5) | 4669 | .00 | 5.00 | 2.81 | 2.17 | 1726 | .00 | 5.00 | 3.09 | 2.09 | 2943 | .00 | 5.00 | 2.64 | 2.20 | 39.25 | .000 | .01 |
| Internet dating (y/n) (0-1) | 4689 | .00 | 1.00 | .47 | .50 | 1734 | .00 | 1.00 | .52 | .50 | 2955 | .00 | 1.00 | .44 | .50 | 21.50 | .000 | .00 |
| Financial literacy: Knowledge of Products (0-4) | 4612 | .00 | 4.00 | 1.39 | .70 | 1717 | .00 | 4.00 | 1.58 | .73 | 2895 | .00 | 4.00 | 1.28 | .65 | 182.00 | .000 | .04 |
| Financial attitudes and behaviour (1-5) | 4610 | 1.00 | 5.00 | 3.55 | .62 | 1717 | 1.00 | 5.00 | 3.48 | .65 | 2893 | 1.00 | 5.00 | 3.59 | .61 | 34.52 | .000 | .01 |

*Note.* Raw uncorrected scores were used. These results are based on one twin randomly selected from each pair so that the data points are independent.
<sup>a</sup> *F* and *p* values are reported with Welch's correction (Welch, 1951) as the assumption of homogeneity of variance was violated for several variables.

**Table S5.**
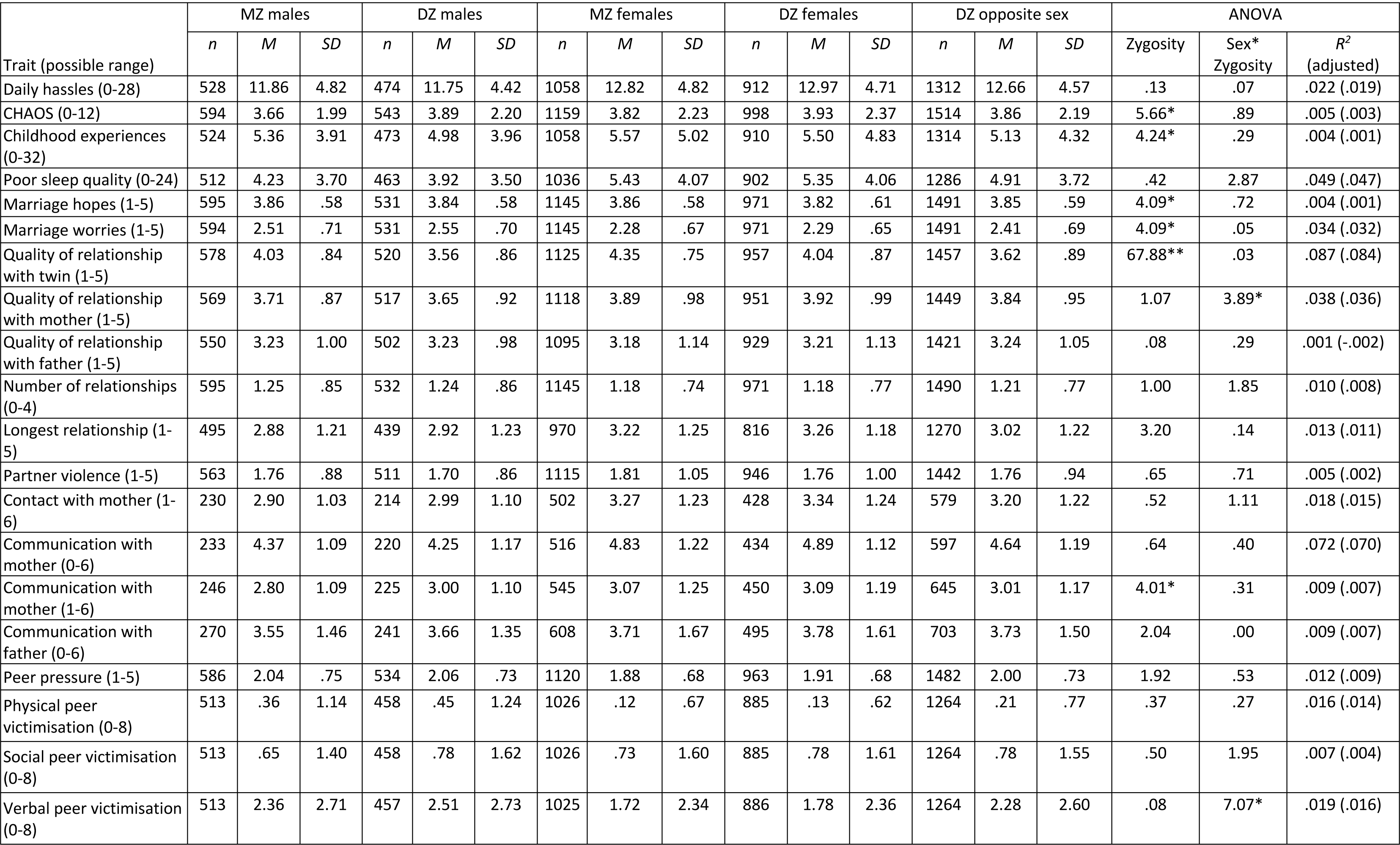
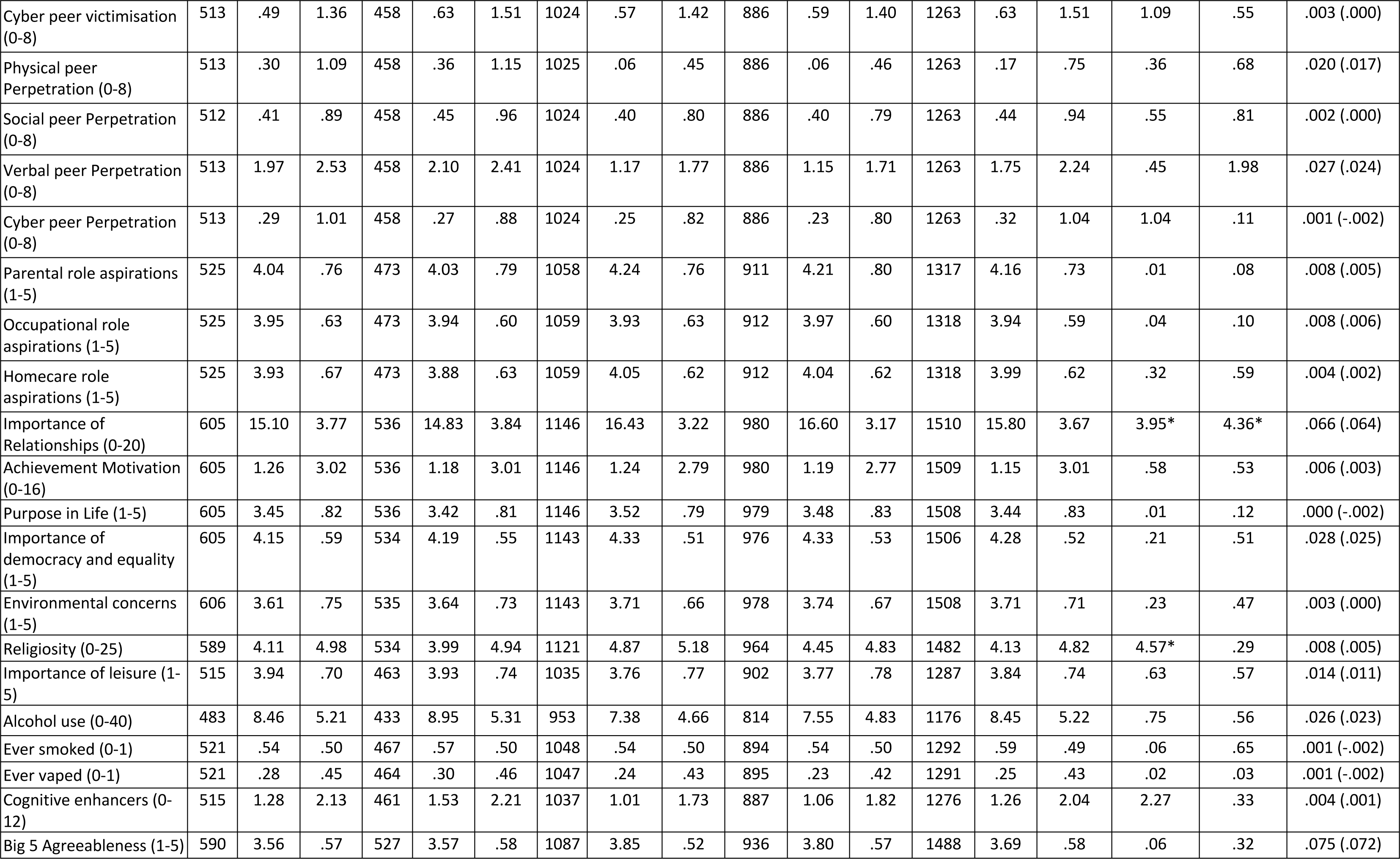
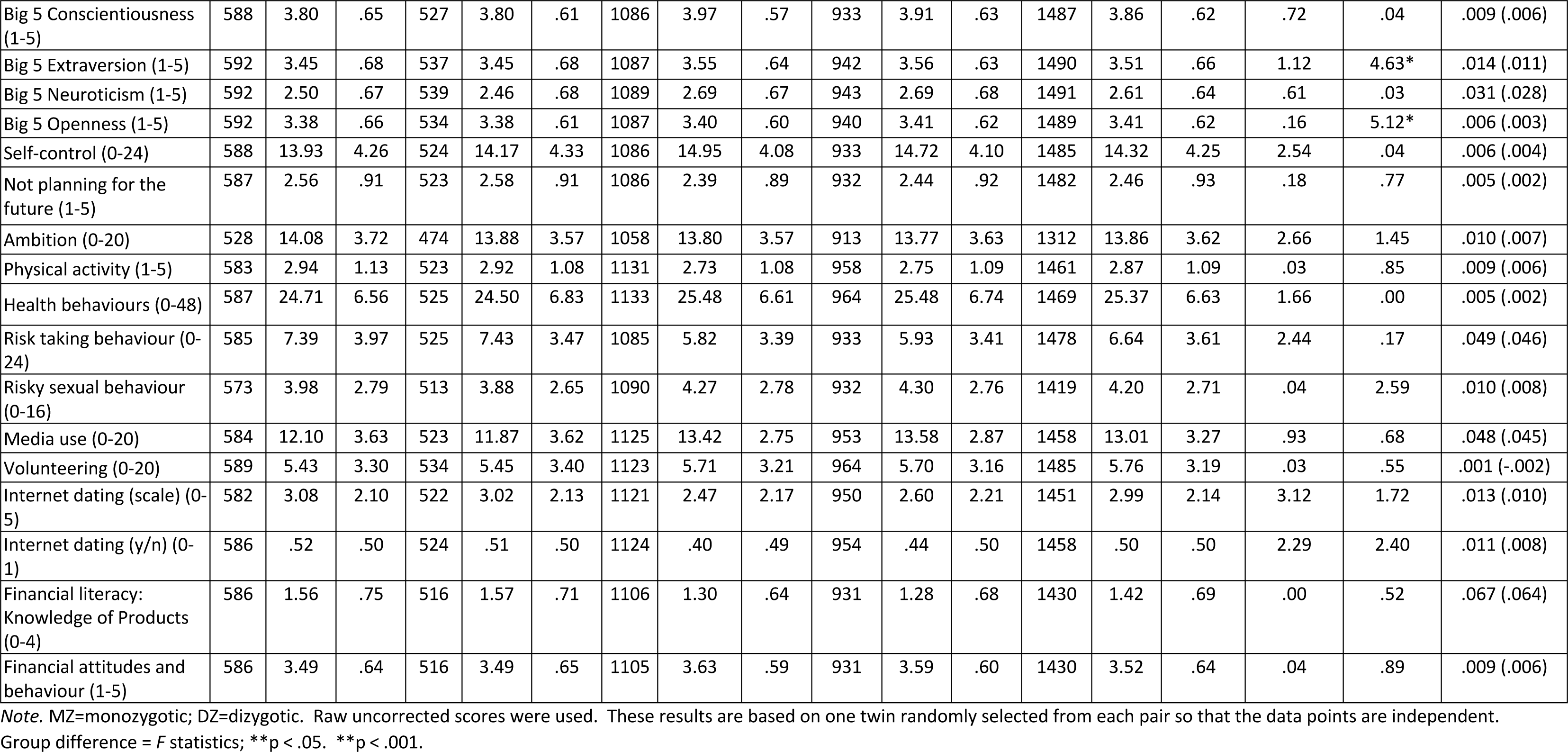
Descriptive statistics for psychological traits, for five sex and zygosity groups (repeated with the twin that was not randomly selected from each pair in the main analyses).

**Table S6.** Phenotypic correlations between psychological traits and composite scores of adverse physical and adverse mental health, wellbeing and educational attainment, for the whole sample (a) and then separately for males (b) and females (c) (95% confidence intervals are in parentheses).

| (a) |  |  |  |  |  |  |  |  |  |  |  |  |  |  |  |
| --- | --- | --- | --- | --- | --- | --- | --- | --- | --- | --- | --- | --- | --- | --- | --- |
| Trait | All |  |  |  |  |  |  |  |  |  |  |  |  |  |  |
|  | Wellbeing |  |  | Adverse mental health |  |  | Adverse physical health |  |  | Achieved educational level |  |  | Planned educational level |  |  |
|  | *r* | *p* | *n* | *r* | *p* | *n* | *r* | *p* | *n* | *r* | *p* | *n* | *r* | *p* | *n* |
| Daily hassles | -.29<br>(-.31,-.26) | <.001 | 3890 | .52<br>(.49,.54) | <.001 | 3607 | .28<br>(.25,.31) | <.001 | 3877 | .01<br>(-.02,.04) | .573 | 3965 | -.01<br>(-.05,.03) | .510 | 2656 |
| CHAOS | -.29<br>(-.33,-.27) | <.001 | 4625 | .32<br>(.29,.35) | <.001 | 3609 | .22<br>(.18,.24) | <.001 | 3879 | -.09<br>(-.12,-.06) | <.001 | 4728 | -.08<br>(-.11,-.04) | <.001 | 3121 |
| Childhood experiences | -.24<br>(-.27,-.21) | <.001 | 3885 | .35<br>(.32,.37) | <.001 | 3591 | .22<br>(.19,.25) | <.001 | 3854 | -.04<br>(-.07,-.01) | .007 | 3959 | -.06<br>(-.10,-.02) | .002 | 2654 |
| Poor sleep quality | -.23<br>(-.26,-.20) | <.001 | 3832 | .50<br>(.47,.52) | <.001 | 3590 | .38<br>(.35,.41) | <.001 | 3875 | -.04<br>(-.07,-.01) | .019 | 3908 | -.03<br>(-.07,.01) | .160 | 2615 |
| Marriage hopes | .14<br>(.11,.17) | <.001 | 4625 | -.08<br>(-.11,-.05) | <.001 | 3610 | -.05<br>(-.08,-.02) | .002 | 3879 | .10<br>(.07,.13) | <.001 | 4727 | .05<br>(.02,.09) | .005 | 3121 |
| Marriage worries | -.28<br>(-.31,-.25) | <.001 | 4626 | .22<br>(.18,.25) | <.001 | 3611 | .12<br>(.09,.15) | <.001 | 3880 | -.03<br>(-.06,.00) | .044 | 4728 | -.05<br>(-.08,-.01) | .013 | 3121 |
| Quality of relationship with twin | .18<br>(.15,.21) | <.001 | 4555 | -.18<br>(-.21,-.14) | <.001 | 3571 | -.09<br>(-.12,-.06) | <.001 | 3832 | .03<br>(.00,.06) | .035 | 4658 | .00<br>(-.03,.04) | .904 | 3080 |
| Quality of relationship with mother | .21<br>(.18,.24) | <.001 | 4510 | -.22<br>(-.25,-.18) | <.001 | 3538 | -.08<br>(-.11,-.05) | <.001 | 3794 | .00<br>(-.03,.03) | .830 | 4612 | .00<br>(-.04,.03) | .872 | 3048 |
| Quality of relationship with father | .25<br>(.22,.27) | <.001 | 4401 | -.26<br>(-.29,-.23) | <.001 | 3465 | -.14<br>(-.17,-.11) | <.001 | 3719 | .04<br>(.01,.07) | .005 | 4497 | .01<br>(-.03,.04) | .726 | 2969 |
| Number of relationships | .06<br>(.04,.09) | <.001 | 4623 | .11<br>(.07,.14) | <.001 | 3608 | .16<br>(.13,.19) | <.001 | 3877 | -.07<br>(-.09,-.04) | <.001 | 4726 | -.06<br>(-.10,-.03) | .001 | 3119 |
| Longest relationship | .17<br>(.14,.20) | <.001 | 3881 | -.02<br>(-.06,.01) | .205 | 3005 | .00<br>(-.03,.04) | .859 | 3232 | -.07<br>(-.10,-.03) | <.001 | 3963 | -.04<br>(-.08,.00) | .061 | 2602 |
| Partner violence | -.21<br>(-.24,-.18) | <.001 | 4476 | .32<br>(.29,.35) | <.001 | 3496 | .22<br>(.19,.25) | <.001 | 3758 | -.10<br>(-.12,-.07) | <.001 | 4573 | -.07<br>(-.10,-.03) | <.001 | 3034 |
| Contact with mother | .10<br>(.05,.14) | <.001 | 1798 | -.02<br>(-.07,.03) | .372 | 1669 | .03<br>(-.02,.07) | .271 | 1772 | -.25<br>(-.29,-.20) | <.001 | 1817 | -.19<br>(-.24,-.14) | <.001 | 1263 |
| Communication with mother | .10<br>(.05,.14) | <.001 | 1831 | -.09<br>(-.14,-.05) | <.001 | 1698 | -.04<br>(-.08,.01) | .138 | 1803 | -.04<br>(-.09,.00) | .066 | 1849 | -.03<br>(-.08,.03) | .302 | 1283 |
| Contact with father | .10<br>(.06,.14) | <.001 | 1957 | -.08<br>(-.13,-.04) | <.001 | 1819 | -.01<br>(-.06,.03) | .614 | 1934 | -.14<br>(-.18,-.09) | <.001 | 1980 | -.15<br>(-.20,-.10) | <.001 | 1340 |
| Communication with father | .13<br>(.09,.18) | <.001 | 2138 | -.16<br>(-.20,-.11) | <.001 | 1977 | -.09<br>(-.13,-.04) | <.001 | 2107 | .05<br>(.01,.09) | .024 | 2164 | .02<br>(-.03,.07) | .416 | 1471 |
| Peer pressure | -.20<br>(-.23,-.17) | <.001 | 4537 | .27<br>(.23,.30) | <.001 | 3608 | .15<br>(.11,.18) | <.001 | 3820 | .02<br>(.00,.05) | .102 | 4642 | -.02<br>(-.05,.02) | .338 | 3067 |
| Physical peer victimisation | -.09<br>(-.13,-.06) | <.001 | 3781 | .16<br>(.12,.19) | <.001 | 3535 | .11<br>(.08,.15) | <.001 | 3792 | -.04<br>(-.07,.00) | .028 | 3852 | -.03<br>(-.07,.01) | .114 | 2584 |
| Social peer victimisation | -.19<br>(-.22,-.16) | <.001 | 3780 | .39<br>(.37,.42) | <.001 | 3534 | .21<br>(.18,.24) | <.001 | 3791 | -.03<br>(-.06,.00) | .084 | 3851 | -.05<br>(-.09,-.01) | .013 | 2584 |
| Verbal peer victimisation | -.18<br>(-.21,-.15) | <.001 | 3780 | .38<br>(.35,.41) | <.001 | 3534 | .22<br>(.18,.25) | <.001 | 3791 | .00<br>(-.03,.03) | .870 | 3851 | .00<br>(-.04,.04) | .876 | 2583 |
| Cyber peer victimisation | -.17<br>(-.20,-.13) | <.001 | 3780 | .35<br>(.32,.38) | <.001 | 3534 | .23<br>(.20,.26) | <.001 | 3791 | -.05<br>(-.08,-.01) | .005 | 3851 | -.07<br>(-.11,-.03) | .001 | 2583 |
| Physical peer Perpetration | -.06<br>(-.09,-.02) | .001 | 3781 | .12<br>(.08,.15) | <.001 | 3537 | .08<br>(.05,.11) | <.001 | 3793 | -.07<br>(-.10,-.04) | <.001 | 3852 | -.10<br>(-.14,-.06) | <.001 | 2584 |
| Social peer Perpetration | -.16<br>(-.19,-.13) | <.001 | 3781 | .30<br>(.26,.32) | <.001 | 3537 | .15<br>(.12,.18) | <.001 | 3793 | -.06<br>(-.09,-.03) | <.001 | 3852 | -.06<br>(-.10,-.03) | .001 | 2584 |
| Verbal peer Perpetration | -.10<br>(-.14,-.07) | <.001 | 3781 | .22<br>(.19,.25) | <.001 | 3537 | .13<br>(.10,.16) | <.001 | 3793 | -.02<br>(-.05,.01) | .285 | 3852 | -.02<br>(-.06,.01) | .226 | 2584 |
| Cyber peer Perpetration | -.09<br>(-.12,-.06) | <.001 | 3781 | .19<br>(.16,.22) | <.001 | 3537 | .14<br>(.11,.17) | <.001 | 3793 | -.05<br>(-.08,-.02) | .003 | 3852 | -.08<br>(-.11,-.04) | <.001 | 2584 |
| Parental role aspirations | .15<br>(.12,.18) | <.001 | 3897 | -.14<br>(-.17,-.10) | <.001 | 3598 | -.04<br>(-.07,-.01) | .009 | 3861 | -.05<br>(-.08,-.01) | .004 | 3970 | -.02<br>(-.06,.02) | .294 | 2661 |
| Occupational role aspirations | .02<br>(-.01,.05) | .223 | 3897 | -.05<br>(-.08,-.02) | .004 | 3598 | -.03<br>(-.06,.00) | .046 | 3861 | .12<br>(.09,.15) | <.001 | 3970 | .07<br>(.04,.11) | <.001 | 2661 |
| Homecare role aspirations | .19<br>(.16,.22) | <.001 | 3897 | -.09<br>(-.12,-.06) | <.001 | 3598 | -.04<br>(-.07,-.01) | .011 | 3861 | -.05<br>(-.08,-.02) | .002 | 3970 | -.05<br>(-.09,-.01) | .011 | 2661 |
| Importance of Relationships | .18<br>(.15,.20) | <.001 | 4577 | -.10<br>(-.13,-.07) | <.001 | 3590 | -.01<br>(-.04,.02) | .549 | 3841 | .04<br>(.01,.07) | .004 | 4681 | .02<br>(-.02,.05) | .417 | 3094 |
| Achievement Motivation | .09<br>(.06,.12) | <.001 | 4577 | -.04<br>(-.08,-.01) | .010 | 3590 | -.01<br>(-.04,.02) | .470 | 3841 | .05<br>(.02,.07) | .002 | 4681 | .04<br>(.00,.07) | .033 | 3094 |
| Purpose in Life | .44<br>(.41,.46) | <.001 | 4576 | -.56<br>(-.58,-.54) | <.001 | 3590 | -.23<br>(-.26,-.20) | <.001 | 3841 | .06<br>(.03,.09) | <.001 | 4680 | .08<br>(.05,.12) | <.001 | 3094 |
| Importance of democracy and equality | -.02<br>(-.05,.01) | .205 | 4580 | .05<br>(.01,.08) | .007 | 3587 | .00<br>(-.03,.03) | .988 | 3841 | .16<br>(.13,.18) | <.001 | 4684 | .11<br>(.08,.15) | <.001 | 3096 |
| Environmental concerns | -.01<br>(-.04,.02) | .514 | 4581 | -.02<br>(-.05,.02) | .322 | 3588 | -.07<br>(-.10,-.04) | <.001 | 3842 | .18<br>(.16,.21) | <.001 | 4685 | .19<br>(.15,.22) | <.001 | 3096 |
| Religiosity | .02<br>(-.01,.05) | .212 | 4543 | .03<br>(.00,.06) | .095 | 3610 | .02<br>(-.01,.05) | .229 | 3823 | .06<br>(.03,.09) | <.001 | 4647 | .07<br>(.03,.10) | <.001 | 3068 |
| Importance of leisure | .32<br>(.29,.35) | <.001 | 3830 | -.40<br>(-.43,-.37) | <.001 | 3610 | -.22<br>(-.25,-.19) | <.001 | 3857 | .11<br>(.08,.14) | <.001 | 3905 | .10<br>(.06,.14) | <.001 | 2614 |
| Alcohol use | -.11<br>(-.14,-.07) | <.001 | 3516 | .14<br>(.11,.18) | <.001 | 3283 | .10<br>(.07,.14) | <.001 | 3517 | .03<br>(.00,.06) | .059 | 3580 | .03<br>(-.01,.06) | .211 | 2430 |
| Ever smoked | -.09<br>(-.12,-.06) | <.001 | 3834 | .12<br>(.08,.15) | <.001 | 3574 | .10<br>(.07,.13) | <.001 | 3837 | -.07<br>(-.10,-.04) | <.001 | 3906 | -.06<br>(-.10,-.02) | .002 | 2622 |
| Ever vaped | -.10<br>(-.13,-.06) | <.001 | 3835 | .16<br>(.12,.19) | <.001 | 3577 | .17<br>(.13,.20) | <.001 | 3837 | -.14<br>(-.17,-.10) | <.001 | 3907 | -.11<br>(-.15,-.08) | <.001 | 2624 |
| Cognitive enhancers | -.15<br>(-.18,-.12) | <.001 | 3809 | .23<br>(.20,.26) | <.001 | 3550 | .14<br>(.11,.17) | <.001 | 3816 | .00<br>(-.03,.03) | .899 | 3877 | .02<br>(-.02,.06) | .280 | 2609 |
| Big 5 Agreeableness | .10<br>(.07,.13) | <.001 | 4360 | -.10<br>(-.14,-.07) | <.001 | 3608 | -.07<br>(-.10,-.03) | <.001 | 3665 | .02<br>(-.01,.05) | .188 | 4449 | .01<br>(-.03,.05) | .547 | 2961 |
| Big 5 Conscientiousness | .26<br>(.23,.29) | <.001 | 4361 | -.29<br>(-.32,-.26) | <.001 | 3608 | -.16<br>(-.19,-.13) | <.001 | 3665 | .09<br>(.06,.12) | <.001 | 4450 | .08<br>(.04,.11) | <.001 | 2961 |
| Big 5 Extraversion | .27<br>(.25,.30) | <.001 | 4360 | -.41<br>(-.44,-.39) | <.001 | 3608 | -.16<br>(-.19,-.13) | <.001 | 3665 | .07<br>(.04,.10) | <.001 | 4449 | .03<br>(-.01,.07) | .101 | 2961 |
| Big 5 Neuroticism | -.36<br>(-.38,-.33) | <.001 | 4361 | .65<br>(.63,.67) | <.001 | 3608 | .30<br>(.27,.33) | <.001 | 3665 | -.04<br>(-.06,-.01) | .020 | 4450 | -.03<br>(-.07,.00) | .067 | 2961 |
| Big 5 Openness | -.14<br>(-.17,-.11) | <.001 | 4360 | .13<br>(.09,.16) | <.001 | 3608 | .04<br>(.01,.07) | .012 | 3665 | .07<br>(.04,.10) | <.001 | 4449 | .02<br>(-.02,.05) | .358 | 2961 |
| Self-control | .28<br>(.25,.31) | <.001 | 4361 | -.45<br>(-.48,-.43) | <.001 | 3608 | -.25<br>(-.28,-.22) | <.001 | 3665 | .07<br>(.04,.10) | <.001 | 4450 | .07<br>(.03,.10) | <.001 | 2961 |
| Not planning for the future | -.16<br>(-.19,-.13) | <.001 | 4361 | .17<br>(.14,.20) | <.001 | 3608 | .08<br>(.05,.12) | <.001 | 3665 | -.10<br>(-.13,-.07) | <.001 | 4450 | -.11<br>(-.15,-.08) | <.001 | 2961 |
| Ambition | .14<br>(.11,.17) | <.001 | 3890 | -.15<br>(-.18,-.12) | <.001 | 3607 | -.07<br>(-.10,-.04) | <.001 | 3877 | .11<br>(.08,.14) | <.001 | 3965 | .09<br>(.06,.13) | <.001 | 2656 |
| Physical activity | .15<br>(.12,.18) | <.001 | 4616 | -.15<br>(-.18,-.11) | <.001 | 3606 | -.14<br>(-.17,-.11) | <.001 | 3874 | .10<br>(.07,.13) | <.001 | 4720 | .07<br>(.04,.11) | <.001 | 3118 |
| Health behaviours | .11<br>(.08,.14) | <.001 | 4623 | -.19<br>(-.22,-.16) | <.001 | 3610 | -.15<br>(-.18,-.11) | <.001 | 3881 | .17<br>(.14,.19) | <.001 | 4728 | .15<br>(.12,.19) | <.001 | 3122 |
| Risk taking behaviour | -.12<br>(-.15,-.09) | <.001 | 4363 | .12<br>(.09,.15) | <.001 | 3611 | .14<br>(.11,.17) | <.001 | 3665 | .01<br>(-.02,.04) | .429 | 4452 | -.01<br>(-.05,.02) | .460 | 2958 |
| Risky sexual behaviour | .06<br>(.03,.09) | <.001 | 4386 | .04<br>(.01,.08) | .010 | 3445 | .10<br>(.06,.13) | <.001 | 3690 | -.08<br>(-.11,-.05) | <.001 | 4487 | -.05<br>(-.09,-.02) | .004 | 2970 |
| Media use | .00<br>(-.03,.03) | .925 | 4605 | .04<br>(.00,.07) | .025 | 3587 | .08<br>(.05,.11) | <.001 | 3851 | .00<br>(-.03,.03) | .982 | 4696 | -.01<br>(-.04,.03) | .684 | 3105 |
| Volunteering | .06<br>(.04,.09) | <.001 | 4543 | .03<br>(.00,.07) | .048 | 3611 | .09<br>(.06,.12) | <.001 | 3825 | .11<br>(.08,.13) | <.001 | 4647 | .09<br>(.05,.12) | <.001 | 3070 |
| Internet dating<br>(scale) | -.18<br>(-.21,-.15) | <.001 | 4587 | .11<br>(.07,.14) | <.001 | 3578 | .09<br>(.06,.13) | <.001 | 3844 | .08<br>(.06,.11) | <.001 | 4683 | .03<br>(-.01,.06) | .121 | 3102 |
| Internet dating<br>(y/n) | -.15<br>(-.18,-.12) | <.001 | 4597 | .10<br>(.07,.14) | <.001 | 3586 | .10<br>(.07,.13) | <.001 | 3852 | .07<br>(.04,.10) | <.001 | 4694 | .02<br>(-.02,.05) | .301 | 3105 |
| Financial literacy:<br>Knowledge of<br>Products | .22<br>(.20,.25) | <.001 | 4626 | -.14<br>(-.17,-.11) | <.001 | 3535 | -.04<br>(-.07,-.01) | .017 | 3789 | .02<br>(-.01,.05) | .239 | 4616 | .01<br>(-.02,.05) | .548 | 3059 |
| Financial attitudes<br>and behaviour | .32<br>(.29,.34) | <.001 | 4625 | -.19<br>(-.22,-.16) | <.001 | 3534 | -.14<br>(-.18,-.11) | <.001 | 3788 | .07<br>(.04,.09) | <.001 | 4615 | .06<br>(.02,.09) | .002 | 3058 |

(b) Males
| Trait | Wellbeing |  |  | Adverse mental health |  |  | Adverse physical health |  |  | Achieved educational level |  |  | Planned educational level |  |  |
| --- | --- | --- | --- | --- | --- | --- | --- | --- | --- | --- | --- | --- | --- | --- | --- |
|  | *r* | *p* | *n* | *r* | *p* | *n* | *r* | *p* | *n* | *r* | *p* | *n* | *r* | *p* | *n* |
| Daily hassles | -.29<br>(-.34,-.25) | <.001 | 1424 | .51<br>(.47,.55) | <.001 | 1312 | .28<br>(.23,.32) | <.001 | 1395 | -.02<br>(-.07,.03) | .563 | 1439 | -.06<br>(-.12,.00) | .050 | 939 |
| CHAOS | -.26<br>(-.30,-.21) | <.001 | 1774 | .28<br>(.23,.33) | <.001 | 1312 | .17<br>(.12,.22) | <.001 | 1395 | -.11<br>(-.16,-.06) | <.001 | 1798 | -.08<br>(-.14,-.02) | .006 | 1149 |
| Childhood<br>experiences | -.22<br>(-.27,-.17) | <.001 | 1418 | .32<br>(.27,.37) | <.001 | 1305 | .17<br>(.12,.22) | <.001 | 1383 | -.07<br>(-.12,-.02) | .013 | 1434 | -.10<br>(-.16,-.04) | .003 | 936 |
| Poor sleep quality | -.22<br>(-.27,-.17) | <.001 | 1401 | .45<br>(.40,.49) | <.001 | 1303 | .32<br>(.28,.37) | <.001 | 1394 | -.05<br>(-.10,.00) | .083 | 1416 | -.06<br>(-.12,.00) | .075 | 924 |
| Marriage hopes | .12<br>(.08,.17) | <.001 | 1774 | -.08<br>(-.14,-.03) | .003 | 1312 | -.02<br>(-.07,.04) | .576 | 1395 | .11<br>(.06,.16) | <.001 | 1798 | .02<br>(-.04,.08) | .442 | 1149 |
| Marriage worries | -.20<br>(-.24,-.15) | <.001 | 1774 | .16<br>(.10,.21) | <.001 | 1312 | .05<br>(.00,.11) | .047 | 1395 | -.02<br>(-.07,.03) | .375 | 1798 | -.06<br>(-.12,.00) | .044 | 1149 |
| Quality of<br>relationship with<br>twin | .16<br>(.11,.20) | <.001 | 1740 | -.09<br>(-.14,-.03) | .001 | 1297 | -.08<br>(-.13,-.03) | .003 | 1373 | .03<br>(-.02,.08) | .173 | 1764 | -.03<br>(-.09,.03) | .364 | 1129 |
| Quality of<br>relationship with<br>mother | .17<br>(.12,.22) | <.001 | 1722 | -.13<br>(-.19,-.08) | <.001 | 1282 | -.09<br>(-.14,-.03) | .002 | 1357 | .03<br>(-.02,.08) | .236 | 1746 | -.03<br>(-.09,.03) | .369 | 1116 |
| Quality of<br>relationship with<br>father | .27<br>(.23,.31) | <.001 | 1676 | -.25<br>(-.30,-.20) | <.001 | 1257 | -.14<br>(-.19,-.09) | <.001 | 1330 | .05<br>(.00,.10) | .050 | 1698 | .03<br>(-.03,.09) | .409 | 1086 |
| Number of relationships | .10<br>(.06,.15) | <.001 | 1772 | .04<br>(-.02,.09) | .173 | 1310 | .11<br>(.06,.16) | <.001 | 1393 | -.06<br>(-.11,-.01) | .013 | 1796 | -.09<br>(-.15,-.03) | .002 | 1148 |
| Longest relationship | .16<br>(.11,.21) | <.001 | 1468 | -.06<br>(-.12,.00) | .049 | 1070 | -.02<br>(-.08,.04) | .568 | 1137 | -.03<br>(-.08,.02) | .192 | 1485 | .03<br>(-.03,.09) | .439 | 955 |
| Partner violence | -.19<br>(-.23,-.14) | <.001 | 1700 | .28<br>(.23,.33) | <.001 | 1255 | .14<br>(.09,.19) | <.001 | 1334 | -.10<br>(-.15,-.05) | <.001 | 1720 | -.07<br>(-.13,-.01) | .015 | 1108 |
| Contact with mother | .11<br>(.03,.19) | .005 | 634 | .00<br>(-.08,.08) | .971 | 591 | .00<br>(-.08,.08) | .997 | 614 | -.21<br>(-.28,-.13) | <.001 | 637 | -.12<br>(-.21,-.03) | .015 | 435 |
| Communication with mother | .07<br>(-.01,.15) | .072 | 645 | -.02<br>(-.10,.06) | .633 | 601 | -.05<br>(-.13,.03) | .236 | 625 | -.04<br>(-.12,.04) | .350 | 648 | .00<br>(-.09,.09) | .942 | 443 |
| Contact with father | .11<br>(.04,.19) | .003 | 689 | -.10<br>(-.17,-.02) | .015 | 642 | -.03<br>(-.10,.05) | .496 | 669 | -.10<br>(-.17,-.03) | .010 | 693 | -.11<br>(-.20,-.02) | .025 | 458 |
| Communication with father | .16<br>(.09,.23) | <.001 | 750 | -.18<br>(-.25,-.11) | <.001 | 694 | -.04<br>(-.11,.03) | .256 | 727 | .07<br>(.00,.14) | .073 | 754 | -.02<br>(-.11,.07) | .657 | 500 |
| Peer pressure | -.17<br>(-.21,-.12) | <.001 | 1740 | .23<br>(.18,.28) | <.001 | 1311 | .15<br>(.10,.20) | <.001 | 1375 | -.01<br>(-.06,.04) | .609 | 1766 | -.04<br>(-.10,.02) | .141 | 1128 |
| Physical peer victimisation | -.10<br>(-.15,-.05) | <.001 | 1388 | .17<br>(.12,.22) | <.001 | 1291 | .11<br>(.05,.16) | <.001 | 1369 | -.02<br>(-.07,.03) | .449 | 1402 | -.05<br>(-.11,.01) | .165 | 917 |
| Social peer victimisation | -.17<br>(-.22,-.12) | <.001 | 1388 | .35<br>(.30,.40) | <.001 | 1291 | .16<br>(.11,.21) | <.001 | 1369 | .01<br>(-.04,.06) | .840 | 1402 | -.05<br>(-.11,.01) | .128 | 917 |
| Verbal peer victimisation | -.13<br>(-.18,-.07) | <.001 | 1387 | .28<br>(.23,.33) | <.001 | 1290 | .13<br>(.08,.18) | <.001 | 1368 | .04<br>(-.01,.09) | .145 | 1401 | .01<br>(-.05,.07) | .766 | 916 |
| Cyber peer victimisation | -.14<br>(-.19,-.09) | <.001 | 1387 | .27<br>(.22,.32) | <.001 | 1290 | .12<br>(.07,.17) | <.001 | 1368 | .01<br>(-.04,.06) | .681 | 1401 | -.05<br>(-.11,.01) | .146 | 916 |
| Physical peer Perpetration | -.07<br>(-.12,-.01) | .012 | 1386 | .13<br>(.07,.18) | <.001 | 1291 | .08<br>(.03,.13) | .002 | 1368 | -.07<br>(-.12,-.02) | .006 | 1400 | -.14<br>(-.20,-.08) | <.001 | 916 |
| Social peer Perpetration | -.16<br>(-.21,-.11) | <.001 | 1386 | .31<br>(.26,.36) | <.001 | 1291 | .13<br>(.07,.18) | <.001 | 1368 | -.04<br>(-.09,.01) | .120 | 1400 | -.07<br>(-.13,-.01) | .051 | 916 |
| Verbal peer Perpetration | -.08<br>(-.13,-.02) | .004 | 1386 | .17<br>(.12,.22) | <.001 | 1291 | .10<br>(.05,.15) | <.001 | 1368 | .02<br>(-.03,.07) | .572 | 1400 | .00<br>(-.06,.06) | .955 | 916 |
| Cyber peer Perpetration | -.08<br>(-.14,-.03) | .002 | 1386 | .17<br>(.11,.22) | <.001 | 1291 | .10<br>(.04,.15) | <.001 | 1368 | -.03<br>(-.08,.02) | .277 | 1400 | -.06<br>(-.12,.00) | .097 | 916 |
| Parental role aspirations | .14<br>(.09,.19) | <.001 | 1423 | -.13<br>(-.18,-.08) | <.001 | 1309 | -.05<br>(-.10,.01) | .088 | 1387 | .01<br>(-.04,.06) | .781 | 1438 | -.01<br>(-.07,.05) | .676 | 938 |
| Occupational role aspirations | .02<br>(-.03,.07) | .464 | 1423 | -.09<br>(-.15,-.04) | .001 | 1309 | -.05<br>(-.10,.01) | .082 | 1387 | .09<br>(.04,.14) | <.001 | 1438 | .08<br>(.02,.14) | .017 | 938 |
| Homecare role aspirations | .20<br>(.15,.25) | <.001 | 1423 | -.08<br>(-.13,-.02) | .005 | 1309 | -.07<br>(-.12,-.02) | .010 | 1387 | -.02<br>(-.07,.03) | .398 | 1438 | -.02<br>(-.08,.04) | .630 | 938 |
| Importance of Relationships | .15<br>(.10,.20) | <.001 | 1755 | -.04<br>(-.10,.01) | .143 | 1305 | .03<br>(-.02,.09) | .215 | 1380 | .06<br>(.01,.11) | .012 | 1780 | .03<br>(-.03,.09) | .253 | 1139 |
| Achievement Motivation | .10<br>(.05,.14) | <.001 | 1755 | -.03<br>(-.09,.02) | .259 | 1305 | .01<br>(-.04,.06) | .705 | 1380 | .07<br>(.02,.12) | .005 | 1780 | .04<br>(-.02,.10) | .182 | 1139 |
| Purpose in Life | .43<br>(.39,.46) | <.001 | 1755 | -.50<br>(-.54,-.46) | <.001 | 1305 | -.20<br>(-.25,-.15) | <.001 | 1380 | .07<br>(.02,.12) | .002 | 1780 | .12<br>(.06,.18) | <.001 | 1139 |
| Importance of democracy and equality | -.01<br>(-.05,.04) | .748 | 1755 | .04<br>(-.02,.09) | .189 | 1303 | .02<br>(-.03,.07) | .465 | 1379 | .12<br>(.07,.17) | <.001 | 1780 | .07<br>(.01,.13) | .013 | 1140 |
| Environmental concerns | .00<br>(-.04,.05) | .913 | 1756 | -.03<br>(-.09,.02) | .256 | 1304 | -.06<br>(-.11,.00) | .035 | 1380 | .19<br>(.14,.23) | <.001 | 1781 | .22<br>(.16,.27) | <.001 | 1140 |
| Religiosity | .04<br>(-.01,.08) | .110 | 1744 | -.01<br>(-.06,.05) | .756 | 1312 | .01<br>(-.04,.06) | .728 | 1377 | .04<br>(-.01,.09) | .133 | 1769 | .04<br>(-.02,.10) | .203 | 1128 |
| Importance of leisure | .32<br>(.27,.36) | <.001 | 1394 | -.39<br>(-.43,-.34) | <.001 | 1312 | -.19<br>(-.24,-.13) | <.001 | 1380 | .10<br>(.05,.15) | <.001 | 1408 | .11<br>(.05,.17) | .001 | 919 |
| Alcohol use | -.08<br>(-.13,-.02) | .004 | 1297 | .11<br>(.05,.16) | <.001 | 1209 | .10<br>(.04,.15) | <.001 | 1279 | .06<br>(.01,.11) | .037 | 1313 | .04<br>(-.03,.11) | .197 | 863 |
| Ever smoked | -.06<br>(-.11,.00) | .041 | 1400 | .09<br>(.03,.14) | .002 | 1298 | .10<br>(.05,.16) | <.001 | 1380 | -.06<br>(-.11,-.01) | .016 | 1415 | -.06<br>(-.12,.00) | .056 | 929 |
| Ever vaped | -.06<br>(-.11,-.01) | .019 | 1400 | .13<br>(.07,.18) | <.001 | 1300 | .15<br>(.09,.20) | <.001 | 1378 | -.13<br>(-.18,-.08) | <.001 | 1415 | -.13<br>(-.19,-.07) | <.001 | 927 |
| Cognitive enhancers | -.10<br>(-.16,-.05) | <.001 | 1387 | .15<br>(.09,.20) | <.001 | 1285 | .12<br>(.07,.17) | <.001 | 1369 | .00<br>(-.05,.05) | .929 | 1400 | .04<br>(-.02,.10) | .274 | 921 |
| Big 5 Agreeableness | .06<br>(.01,.11) | .013 | 1677 | -.03<br>(-.08,.03) | .368 | 1312 | -.07<br>(-.12,-.01) | .015 | 1325 | .05<br>(.00,.10) | .065 | 1702 | .00<br>(-.06,.06) | .996 | 1095 |
| Big 5 Conscientiousness | .26<br>(.21,.30) | <.001 | 1677 | -.29<br>(-.34,-.24) | <.001 | 1312 | -.15<br>(-.20,-.10) | <.001 | 1325 | .09<br>(.04,.14) | <.001 | 1702 | .05<br>(-.01,.11) | .091 | 1095 |
| Big 5 Extraversion | .25<br>(.21,.30) | <.001 | 1677 | -.35<br>(-.40,-.31) | <.001 | 1312 | -.13<br>(-.18,-.07) | <.001 | 1325 | .04<br>(-.01,.09) | .093 | 1702 | -.01<br>(-.07,.05) | .695 | 1095 |
| Big 5 Neuroticism | -.34<br>(-.39,-.30) | <.001 | 1677 | .61<br>(.57,.64) | <.001 | 1312 | .31<br>(.26,.35) | <.001 | 1325 | -.08<br>(-.13,-.03) | .002 | 1702 | -.05<br>(-.11,.01) | .086 | 1095 |
| Big 5 Openness | -.15<br>(-.20,-.10) | <.001 | 1677 | .17<br>(.12,.22) | <.001 | 1312 | .06<br>(.01,.12) | .024 | 1325 | .07<br>(.02,.12) | .006 | 1702 | .02<br>(-.04,.08) | .538 | 1095 |
| Self-control | .28<br>(.23,.32) | <.001 | 1677 | -.46<br>(-.50,-.41) | <.001 | 1312 | -.22<br>(-.27,-.17) | <.001 | 1325 | .08<br>(.03,.13) | .001 | 1702 | .09<br>(.03,.15) | .004 | 1095 |
| Not planning for the future | -.16<br>(-.21,-.12) | <.001 | 1677 | .20<br>(.15,.26) | <.001 | 1312 | .10<br>(.05,.16) | <.001 | 1325 | -.10<br>(-.15,-.05) | <.001 | 1702 | -.10<br>(-.16,-.04) | .001 | 1095 |
| Ambition | .16<br>(.11,.21) | <.001 | 1424 | -.21<br>(-.26,-.15) | <.001 | 1312 | -.09<br>(-.14,-.04) | .001 | 1395 | .09<br>(.04,.14) | .001 | 1439 | .09<br>(.03,.15) | .009 | 939 |
| Physical activity | .20<br>(.15,.24) | <.001 | 1767 | -.20<br>(-.25,-.15) | <.001 | 1309 | -.16<br>(-.21,-.11) | <.001 | 1390 | .08<br>(.03,.13) | .002 | 1791 | .04<br>(-.02,.10) | .142 | 1147 |
| Health behaviours | .09<br>(.05,.14) | <.001 | 1771 | -.16<br>(-.21,-.10) | <.001 | 1311 | -.14<br>(-.19,-.08) | <.001 | 1395 | .15<br>(.10,.19) | <.001 | 1796 | .14<br>(.08,.20) | <.001 | 1149 |
| Risk taking behaviour | -.08<br>(-.13,-.04) | .001 | 1677 | .08<br>(.02,.13) | .005 | 1312 | .17<br>(.12,.23) | <.001 | 1324 | -.01<br>(-.06,.04) | .722 | 1702 | -.06<br>(-.12,.00) | .057 | 1093 |
| Risky sexual behaviour | .09<br>(.04,.13) | <.001 | 1682 | .03<br>(-.02,.09) | .261 | 1252 | .11<br>(.05,.16) | <.001 | 1329 | -.08<br>(-.13,-.03) | .001 | 1708 | -.05<br>(-.11,.01) | .088 | 1094 |
| Media use | .02<br>(-.02,.07) | .305 | 1763 | -.01<br>(-.06,.05) | .852 | 1301 | .06<br>(.00,.11) | .037 | 1382 | .04<br>(-.01,.09) | .125 | 1785 | .02<br>(-.04,.08) | .456 | 1142 |
| Volunteering | .08<br>(.03,.12) | .001 | 1744 | -.03<br>(-.08,.03) | .336 | 1312 | .06<br>(.00,.11) | .037 | 1377 | .09<br>(.04,.14) | <.001 | 1769 | .03<br>(-.03,.09) | .252 | 1128 |
| Internet dating (scale) | -.15<br>(-.20,-.10) | <.001 | 1756 | .10<br>(.04,.15) | .001 | 1298 | .03<br>(-.02,.08) | .257 | 1379 | .10<br>(.05,.15) | <.001 | 1779 | -.01<br>(-.07,.05) | .840 | 1141 |
| Internet dating (y/n) | -.11<br>(-.16,-.06) | <.001 | 1760 | .08<br>(.03,.14) | .003 | 1301 | .03<br>(-.02,.08) | .276 | 1382 | .08<br>(.03,.13) | .001 | 1784 | -.02<br>(-.08,.04) | .554 | 1142 |
| Financial literacy: Knowledge of Products | .23<br>(.19,.28) | <.001 | 1774 | -.15<br>(-.20,-.10) | <.001 | 1293 | -.05<br>(-.11,.00) | .046 | 1374 | .05<br>(.00,.10) | .044 | 1769 | .04<br>(-.02,.10) | .181 | 1137 |
| Financial attitudes and behaviour | .31<br>(.27,.36) | <.001 | 1774 | -.22<br>(-.27,-.17) | <.001 | 1293 | -.18<br>(-.23,-.13) | <.001 | 1374 | .06<br>(.01,.11) | .010 | 1769 | .08<br>(.02,.14) | .006 | 1137 |

(c) Females
| Trait | Wellbeing |  |  | Adverse mental health |  |  | Adverse physical health |  |  | Achieved educational level |  |  | Planned educational level |  |  |
| --- | --- | --- | --- | --- | --- | --- | --- | --- | --- | --- | --- | --- | --- | --- | --- |
|  | *r* | *p* | *n* | *r* | *p* | *n* | *r* | *p* | *n* | *r* | *p* | *n* | *r* | *p* | *n* |
| Daily hassles | -.28<br>(-.32,-.24) | <.001 | 2466 | .52<br>(.49,.55) | <.001 | 2295 | .28<br>(.24,.32) | <.001 | 2482 | .02<br>(-.02,.06) | .239 | 2526 | .02<br>(-.03,.07) | .535 | 1717 |
| CHAOS | -.33<br>(-.36,-.29) | <.001 | 2851 | .34<br>(.31,.38) | <.001 | 2297 | .23<br>(.19,.27) | <.001 | 2484 | -.08<br>(-.12,-.04) | <.001 | 2930 | -.08<br>(-.12,-.04) | .001 | 1972 |
| Childhood experiences | -.25<br>(-.28,-.21) | <.001 | 2467 | .36<br>(.32,.39) | <.001 | 2286 | .23<br>(.20,.27) | <.001 | 2471 | -.03<br>(-.07,.01) | .113 | 2525 | -.05<br>(-.10,.00) | .058 | 1718 |
| Poor sleep quality | -.24<br>(-.27,-.20) | <.001 | 2431 | .52<br>(.49,.55) | <.001 | 2287 | .40<br>(.37,.44) | <.001 | 2481 | -.03<br>(-.07,.01) | .098 | 2492 | -.01<br>(-.06,.04) | .614 | 1691 |
| Marriage hopes | .15<br>(.11,.18) | <.001 | 2851 | -.08<br>(-.12,-.04) | <.001 | 2298 | -.07<br>(-.10,-.03) | .001 | 2484 | .10<br>(.06,.14) | <.001 | 2929 | .07<br>(.03,.11) | .003 | 1972 |
| Marriage worries | -.33<br>(-.37,-.30) | <.001 | 2852 | .25<br>(.21,.28) | <.001 | 2299 | .16<br>(.12,.19) | <.001 | 2485 | -.04<br>(-.08,.00) | .059 | 2930 | -.04<br>(-.08,.00) | .114 | 1972 |
| Quality of relationship with twin | .19<br>(.16,.23) | <.001 | 2815 | -.22<br>(-.26,-.18) | <.001 | 2274 | -.09<br>(-.13,-.05) | <.001 | 2459 | .03<br>(-.01,.07) | .107 | 2894 | .02<br>(-.02,.06) | .374 | 1951 |
| Quality of relationship with mother | .24<br>(.20,.27) | <.001 | 2788 | -.25<br>(-.29,-.21) | <.001 | 2256 | -.08<br>(-.12,-.04) | <.001 | 2437 | -.01<br>(-.05,.03) | .551 | 2866 | .01<br>(-.03,.05) | .668 | 1932 |
| Quality of relationship with father | .23<br>(.19,.27) | <.001 | 2725 | -.26<br>(-.30,-.22) | <.001 | 2208 | -.14<br>(-.18,-.10) | <.001 | 2389 | .04<br>(.00,.08) | .039 | 2799 | .00<br>(-.05,.05) | .873 | 1883 |
| Number of relationships | .04<br>(.00,.07) | .047 | 2851 | .14<br>(.10,.18) | <.001 | 2298 | .19<br>(.15,.22) | <.001 | 2484 | -.07<br>(-.11,-.03) | <.001 | 2930 | -.04<br>(-.08,.00) | .076 | 1971 |
| Longest relationship | .17<br>(.14,.21) | <.001 | 2413 | -.01<br>(-.05,.04) | .820 | 1935 | .01<br>(-.03,.05) | .572 | 2095 | -.09<br>(-.13,-.05) | <.001 | 2478 | -.08<br>(-.13,-.03) | .002 | 1647 |
| Partner violence | -.22<br>(-.26,-.19) | <.001 | 2776 | .33<br>(.29,.37) | <.001 | 2241 | .25<br>(.21,.29) | <.001 | 2424 | -.10<br>(-.14,-.06) | <.001 | 2853 | -.07<br>(-.11,-.03) | .004 | 1926 |
| Contact with mother | .09<br>(.03,.14) | .003 | 1164 | -.03<br>(-.09,.03) | .323 | 1078 | .04<br>(-.02,.09) | .216 | 1158 | -.27<br>(-.32,-.22) | <.001 | 1180 | -.23<br>(-.29,-.16) | <.001 | 828 |
| Communication with mother | .11<br>(.05,.17) | <.001 | 1186 | -.13<br>(-.18,-.07) | <.001 | 1097 | -.03<br>(-.09,.03) | .311 | 1178 | -.05<br>(-.11,.01) | .122 | 1201 | -.04<br>(-.11,.03) | .237 | 840 |
| Contact with father | .10<br>(.04,.15) | .001 | 1268 | -.08<br>(-.13,-.02) | .009 | 1177 | -.01<br>(-.06,.05) | .859 | 1265 | -.16<br>(-.21,-.11) | <.001 | 1287 | -.17<br>(-.23,-.11) | <.001 | 882 |
| Communication with father | .12<br>(.07,.17) | <.001 | 1388 | -.15<br>(-.20,-.09) | <.001 | 1283 | -.10<br>(-.15,-.05) | <.001 | 1380 | .04<br>(-.01,.09) | .124 | 1410 | .04<br>(-.02,.10) | .190 | 971 |
| Peer pressure | -.22<br>(-.26,-.19) | <.001 | 2797 | .29<br>(.25,.32) | <.001 | 2297 | .14<br>(.11,.18) | <.001 | 2445 | .05<br>(.01,.09) | .009 | 2876 | .00<br>(-.04,.04) | 1<.001 | 1939 |
| Physical peer victimisation | -.10<br>(-.13,-.06) | <.001 | 2393 | .16<br>(.12,.20) | <.001 | 2244 | .14<br>(.10,.17) | <.001 | 2423 | -.05<br>(-.09,-.01) | .009 | 2450 | -.02<br>(-.07,.03) | .428 | 1667 |
| Social peer victimisation | -.20<br>(-.24,-.16) | <.001 | 2392 | .41<br>(.38,.45) | <.001 | 2243 | .23<br>(.19,.26) | <.001 | 2422 | -.05<br>(-.09,-.01) | .022 | 2449 | -.05<br>(-.10,.00) | .047 | 1667 |
| Verbal peer victimisation | -.22<br>(-.25,-.18) | <.001 | 2393 | .44<br>(.40,.47) | <.001 | 2244 | .26<br>(.23,.30) | <.001 | 2423 | -.03<br>(-.07,.01) | .127 | 2450 | -.01<br>(-.06,.04) | .632 | 1667 |
| Cyber peer victimisation | -.18<br>(-.22,-.14) | <.001 | 2393 | .39<br>(.35,.42) | <.001 | 2244 | .27<br>(.23,.31) | <.001 | 2423 | -.08<br>(-.12,-.04) | <.001 | 2450 | -.08<br>(-.13,-.03) | .002 | 1667 |
| Physical peer Perpetration | -.05<br>(-.09,-.01) | .013 | 2395 | .14<br>(.10,.18) | <.001 | 2246 | .10<br>(.06,.14) | <.001 | 2425 | -.08<br>(-.12,-.04) | <.001 | 2452 | -.06<br>(-.11,-.01) | .010 | 1668 |
| Social peer Perpetration | -.16<br>(-.20,-.12) | <.001 | 2395 | .29<br>(.25,.33) | <.001 | 2246 | .17<br>(.13,.21) | <.001 | 2425 | -.07<br>(-.11,-.03) | .001 | 2452 | -.06<br>(-.11,-.01) | .010 | 1668 |
| Verbal peer Perpetration | -.13<br>(-.17,-.09) | <.001 | 2395 | .26<br>(.22,.30) | <.001 | 2246 | .16<br>(.12,.19) | <.001 | 2425 | -.04<br>(-.08,.00) | .030 | 2452 | -.04<br>(-.09,.01) | .088 | 1668 |
| Cyber peer Perpetration | -.10<br>(-.13,-.06) | <.001 | 2395 | .21<br>(.17,.25) | <.001 | 2246 | .17<br>(.13,.21) | <.001 | 2425 | -.06<br>(-.10,-.02) | .003 | 2452 | -.09<br>(-.14,-.04) | <.001 | 1668 |
| Parental role aspirations | .15<br>(.11,.19) | <.001 | 2474 | -.14<br>(-.18,-.10) | <.001 | 2289 | -.04<br>(-.08,.00) | .045 | 2474 | -.08<br>(-.12,-.04) | <.001 | 2532 | -.03<br>(-.08,.02) | .309 | 1723 |
| Occupational role aspirations | .02<br>(-.02,.06) | .337 | 2474 | -.03<br>(-.07,.02) | .210 | 2289 | -.03<br>(-.06,.01) | .208 | 2474 | .14<br>(.10,.18) | <.001 | 2532 | .07<br>(.02,.12) | .003 | 1723 |
| Homecare role aspirations | .18<br>(.14,.22) | <.001 | 2474 | -.09<br>(-.13,-.05) | <.001 | 2289 | -.03<br>(-.07,.01) | .202 | 2474 | -.07<br>(-.11,-.03) | .001 | 2532 | -.07<br>(-.12,-.02) | .004 | 1723 |
| Importance of Relationships | .20<br>(.16,.23) | <.001 | 2822 | -.14<br>(-.18,-.09) | <.001 | 2285 | -.03<br>(-.07,.01) | .106 | 2461 | .03<br>(-.01,.07) | .135 | 2901 | .00<br>(-.04,.04) | .990 | 1955 |
| Achievement Motivation | .09<br>(.05,.12) | <.001 | 2822 | -.05<br>(-.09,-.01) | .018 | 2285 | -.02<br>(-.06,.02) | .296 | 2461 | .03<br>(-.01,.07) | .095 | 2901 | .04<br>(.00,.08) | .096 | 1955 |
| Purpose in Life | .44<br>(.41,.47) | <.001 | 2821 | -.59<br>(-.61,-.56) | <.001 | 2285 | -.25<br>(-.28,-.21) | <.001 | 2461 | .05<br>(.01,.09) | .006 | 2900 | .06<br>(.02,.10) | .007 | 1955 |
| Importance of democracy and equality | -.03<br>(-.06,.01) | .161 | 2825 | .05<br>(.01,.09) | .018 | 2284 | -.01<br>(-.05,.03) | .567 | 2462 | .18<br>(.14,.22) | <.001 | 2904 | .14<br>(.10,.18) | <.001 | 1956 |
| Environmental concerns | -.02<br>(-.05,.02) | .352 | 2825 | -.01<br>(-.05,.03) | .650 | 2284 | -.08<br>(-.12,-.04) | <.001 | 2462 | .18<br>(.14,.22) | <.001 | 2904 | .17<br>(.13,.21) | <.001 | 1956 |
| Religiosity | .01<br>(-.03,.04) | .721 | 2799 | .05<br>(.00,.09) | .030 | 2298 | .02<br>(-.02,.06) | .242 | 2446 | .07<br>(.03,.11) | <.001 | 2878 | .08<br>(.04,.12) | <.001 | 1940 |
| Importance of leisure | .32<br>(.28,.35) | <.001 | 2436 | -.41<br>(-.44,-.37) | <.001 | 2298 | -.24<br>(-.27,-.20) | <.001 | 2477 | .12<br>(.08,.16) | <.001 | 2497 | .10<br>(.05,.15) | <.001 | 1695 |
| Alcohol use | -.13<br>(-.17,-.08) | <.001 | 2219 | .16<br>(.12,.21) | <.001 | 2074 | .11<br>(.07,.15) | <.001 | 2238 | .01<br>(-.03,.05) | .514 | 2267 | .02<br>(-.03,.07) | .557 | 1567 |
| Ever smoked | -.11<br>(-.15,-.07) | <.001 | 2434 | .13<br>(.09,.17) | <.001 | 2276 | .10<br>(.06,.14) | <.001 | 2457 | -.07<br>(-.11,-.03) | .001 | 2491 | -.06<br>(-.11,-.01) | .014 | 1693 |
| Ever vaped | -.12<br>(-.16,-.08) | <.001 | 2435 | .18<br>(.13,.21) | <.001 | 2277 | .18<br>(.14,.22) | <.001 | 2459 | -.14<br>(-.18,-.10) | <.001 | 2492 | -.10<br>(-.15,-.05) | <.001 | 1697 |
| Cognitive enhancers | -.18<br>(-.22,-.14) | <.001 | 2422 | .28<br>(.24,.32) | <.001 | 2265 | .15<br>(.11,.19) | <.001 | 2447 | -.01<br>(-.05,.03) | .799 | 2477 | .01<br>(-.04,.06) | .624 | 1688 |
| Big 5 Agreeableness | .12<br>(.08,.16) | <.001 | 2683 | -.15<br>(-.18,-.10) | <.001 | 2296 | -.07<br>(-.11,-.02) | .002 | 2340 | .00<br>(-.04,.04) | .928 | 2747 | .02<br>(-.03,.07) | .421 | 1866 |
| Big 5 Conscientiousness | .27<br>(.23,.30) | <.001 | 2684 | -.29<br>(-.33,-.25) | <.001 | 2296 | -.17<br>(-.21,-.13) | <.001 | 2340 | .09<br>(.05,.13) | <.001 | 2748 | .09<br>(.04,.13) | <.001 | 1866 |
| Big 5 Extraversion | .29<br>(.25,.32) | <.001 | 2683 | -.45<br>(-.48,-.42) | <.001 | 2296 | -.17<br>(-.21,-.13) | <.001 | 2340 | .09<br>(.05,.13) | <.001 | 2747 | .06<br>(.01,.11) | .012 | 1866 |
| Big 5 Neuroticism | -.36<br>(-.39,-.33) | <.001 | 2684 | .67<br>(.64,.69) | <.001 | 2296 | .30<br>(.27,.34) | <.001 | 2340 | -.01<br>(-.05,.03) | .626 | 2748 | -.02<br>(-.07,.03) | .311 | 1866 |
| Big 5 Openness | -.13<br>(-.17,-.10) | <.001 | 2683 | .10<br>(.06,.14) | <.001 | 2296 | .03<br>(-.01,.07) | .120 | 2340 | .07<br>(.03,.11) | <.001 | 2747 | .02<br>(-.03,.07) | .493 | 1866 |
| Self-control | .29<br>(.25,.32) | <.001 | 2684 | -.45<br>(-.48,-.42) | <.001 | 2296 | -.26<br>(-.30,-.23) | <.001 | 2340 | .07<br>(.03,.11) | <.001 | 2748 | .05<br>(.00,.10) | .031 | 1866 |
| Not planning for the future | -.16<br>(-.20,-.13) | <.001 | 2684 | .16<br>(.11,.19) | <.001 | 2296 | .07<br>(.03,.11) | <.001 | 2340 | -.10<br>(-.14,-.06) | <.001 | 2748 | -.12<br>(-.16,-.08) | <.001 | 1866 |
| Ambition | .13<br>(.10,.17) | <.001 | 2466 | -.12<br>(-.16,-.08) | <.001 | 2295 | -.06<br>(-.10,-.02) | .001 | 2482 | .12<br>(.08,.16) | <.001 | 2526 | .10<br>(.05,.15) | <.001 | 1717 |
| Physical activity | .12<br>(.08,.16) | <.001 | 2849 | -.12<br>(-.16,-.08) | <.001 | 2297 | -.13<br>(-.17,-.10) | <.001 | 2484 | .12<br>(.08,.16) | <.001 | 2929 | .09<br>(.05,.13) | <.001 | 1971 |
| Health behaviours | .12<br>(.08,.15) | <.001 | 2852 | -.21<br>(-.24,-.17) | <.001 | 2299 | -.15<br>(-.19,-.11) | <.001 | 2486 | .17<br>(.13,.20) | <.001 | 2932 | .16<br>(.12,.20) | <.001 | 1973 |
| Risk taking behaviour | -.14<br>(-.18,-.11) | <.001 | 2686 | .15<br>(.11,.19) | <.001 | 2299 | .13<br>(.09,.17) | <.001 | 2341 | .03<br>(-.01,.07) | .150 | 2750 | .02<br>(-.03,.07) | .439 | 1865 |
| Risky sexual behaviour | .04<br>(.01,.08) | .025 | 2704 | .05<br>(.01,.09) | .019 | 2193 | .10<br>(.06,.14) | <.001 | 2361 | -.08<br>(-.12,-.04) | <.001 | 2779 | -.05<br>(-.10,.00) | .021 | 1876 |
| Media use | -.02<br>(-.06,.02) | .267 | 2842 | .06<br>(.02,.10) | .003 | 2286 | .10<br>(.06,.14) | <.001 | 2469 | -.03<br>(-.07,.01) | .118 | 2911 | -.03<br>(-.07,.01) | .196 | 1963 |
| Volunteering | .06<br>(.02,.09) | .003 | 2799 | .06<br>(.02,.10) | .003 | 2299 | .10<br>(.06,.14) | <.001 | 2448 | .12<br>(.08,.16) | <.001 | 2878 | .12<br>(.08,.16) | <.001 | 1942 |
| Internet dating (scale) | -.20<br>(-.23,-.16) | <.001 | 2831 | .11<br>(.07,.15) | <.001 | 2280 | .12<br>(.08,.16) | <.001 | 2465 | .08<br>(.04,.12) | <.001 | 2904 | .05<br>(.01,.09) | .035 | 1961 |
| Internet dating (y/n) | -.18<br>(-.21,-.14) | <.001 | 2837 | .12<br>(.08,.16) | <.001 | 2285 | .13<br>(.09,.17) | <.001 | 2470 | .06<br>(.02,.10) | .001 | 2910 | .04<br>(.00,.08) | .071 | 1963 |
| Financial literacy: Knowledge of Products | .22<br>(.18,.25) | <.001 | 2852 | -.13<br>(-.17,-.09) | <.001 | 2242 | -.03<br>(-.07,.01) | .126 | 2415 | -.01<br>(-.05,.03) | .773 | 2847 | -.01<br>(-.05,.03) | .679 | 1922 |
| Financial attitudes and behaviour | .32<br>(.29,.35) | <.001 | 2851 | -.17<br>(-.21,-.13) | <.001 | 2241 | -.13<br>(-.17,-.09) | <.001 | 2414 | .07<br>(.03,.11) | <.001 | 2846 | .04<br>(.00,.08) | .078 | 1921 |

*Note.* Scores were corrected for mean age and sex differences (see Methods). These results are based on one twin randomly selected from each pair so that the data points are independent.

**Table S7.**
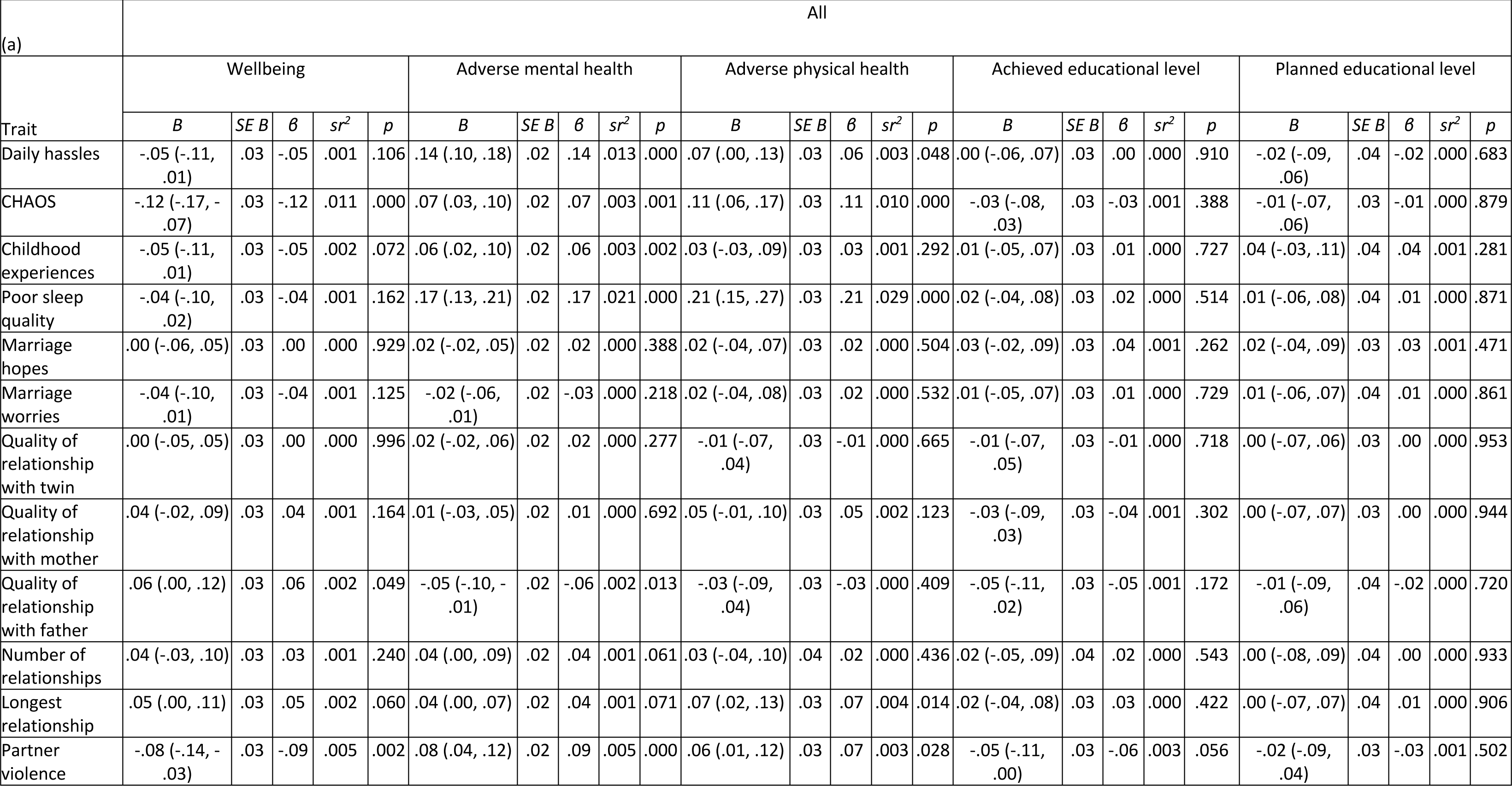
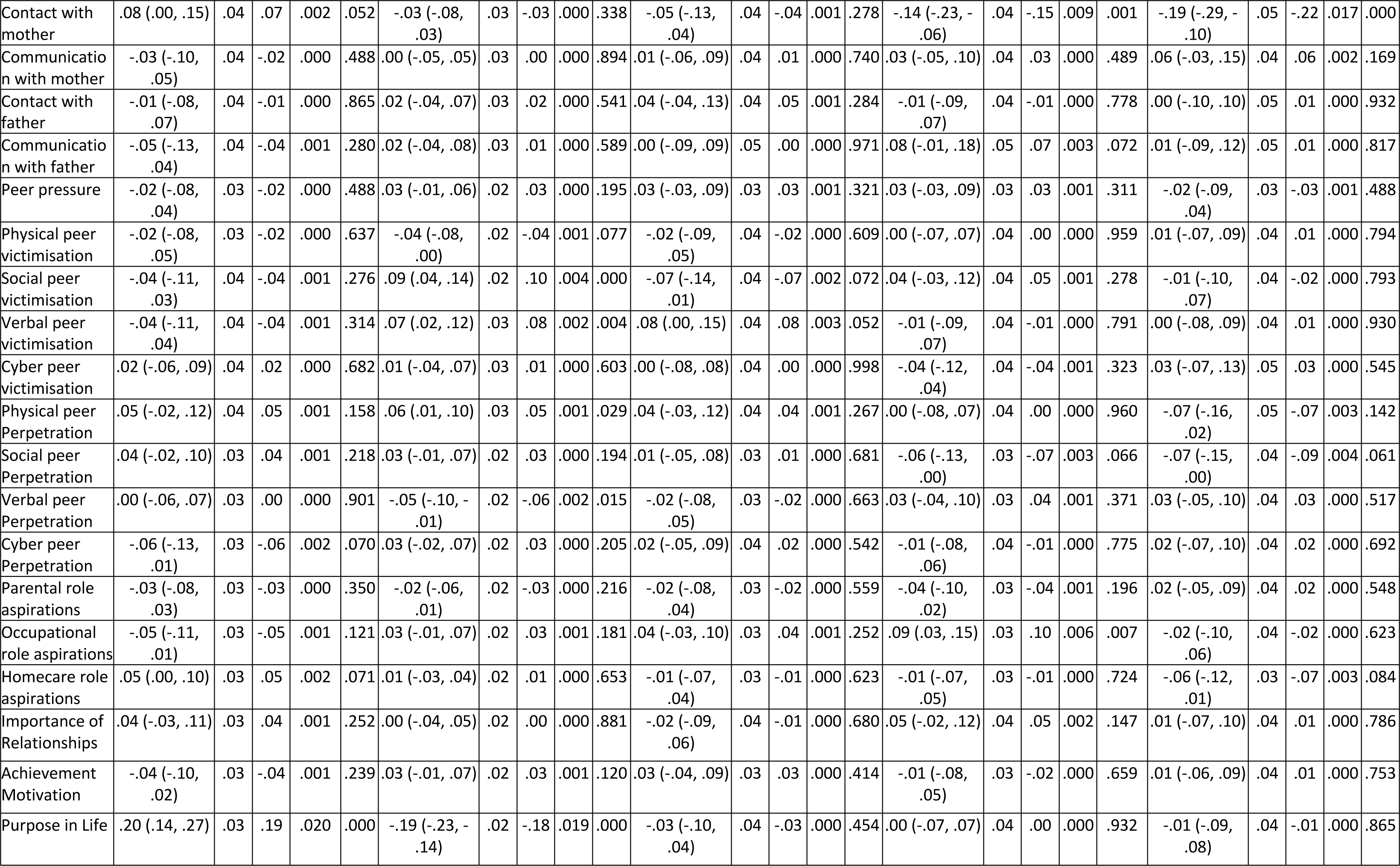
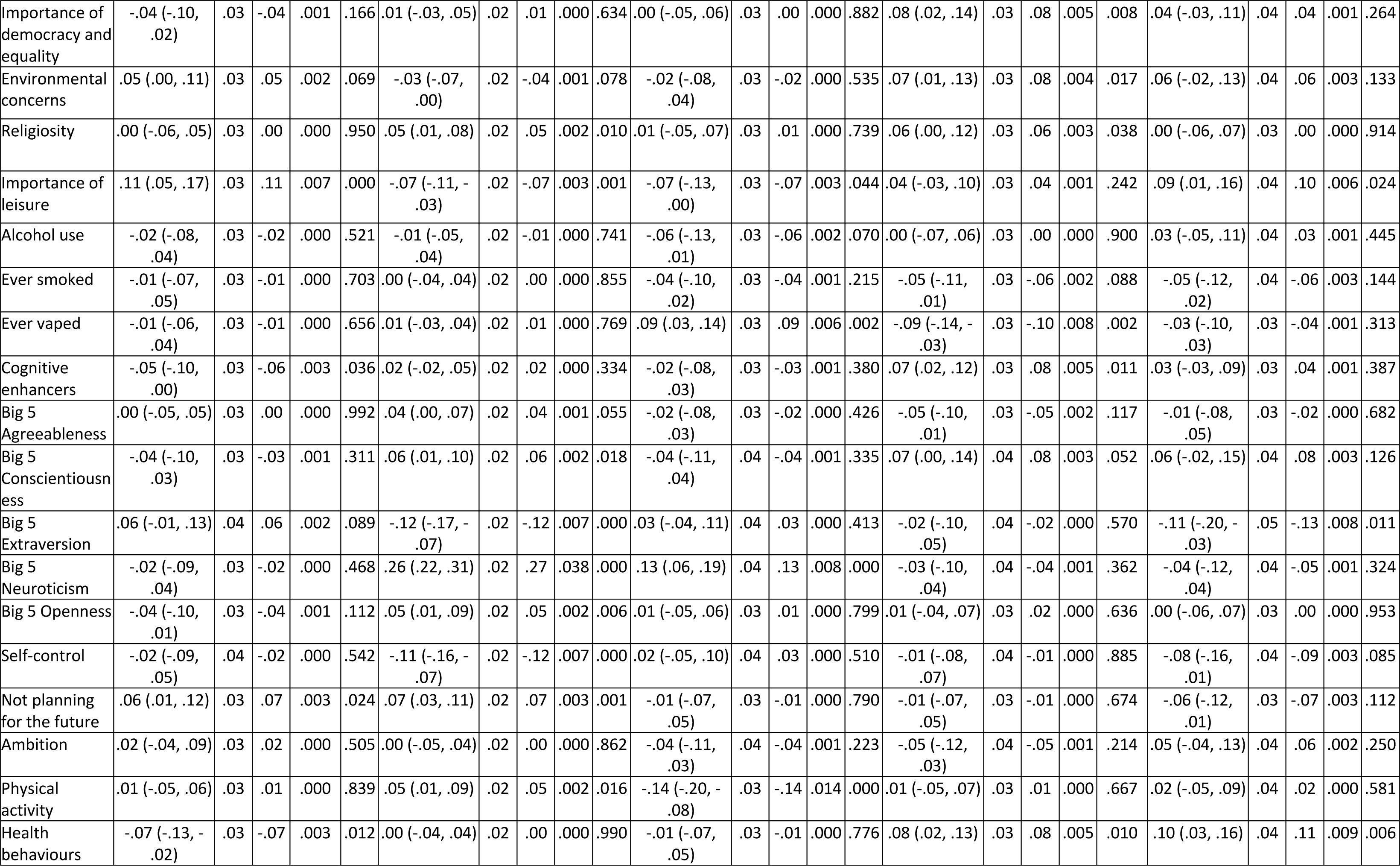
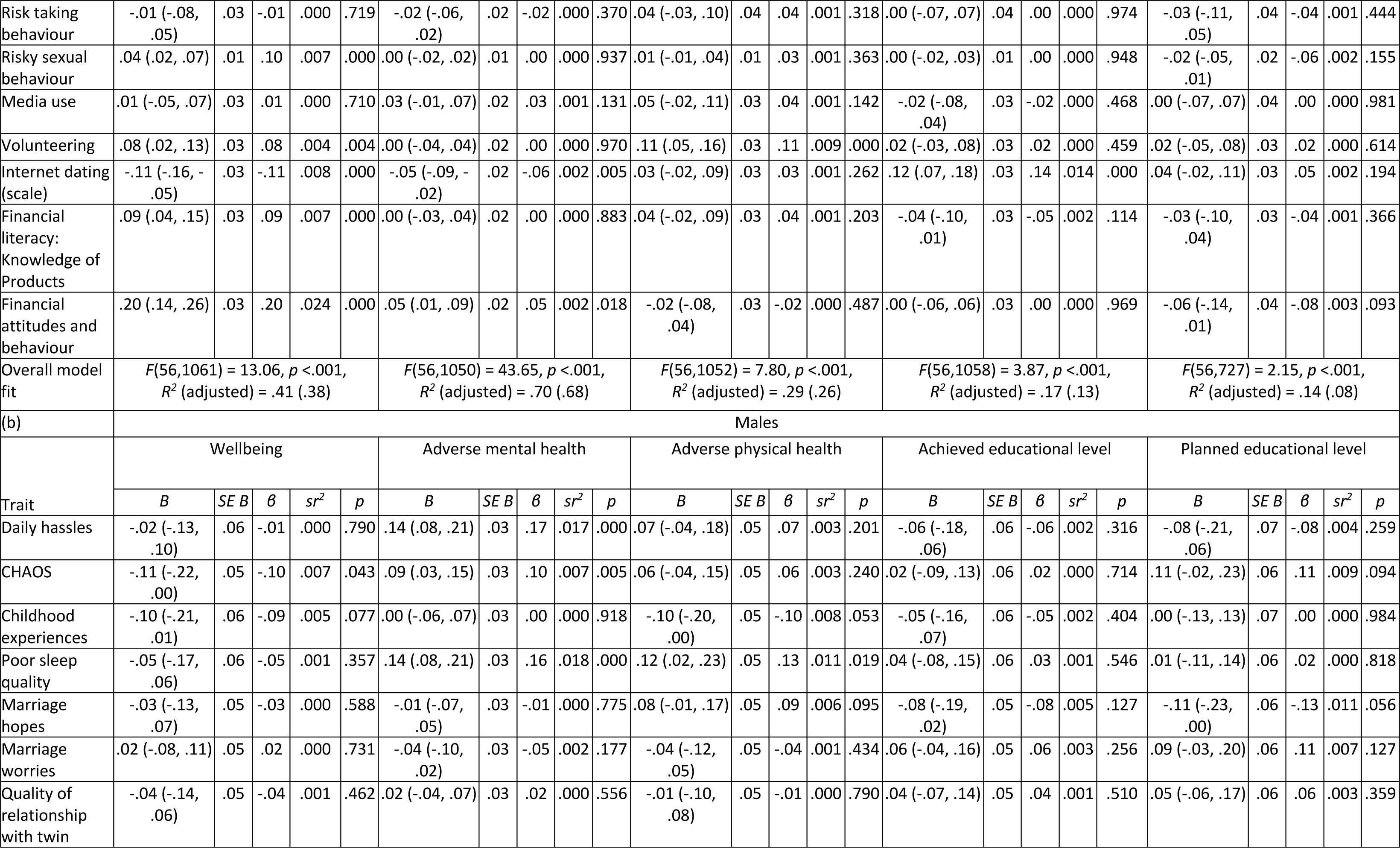
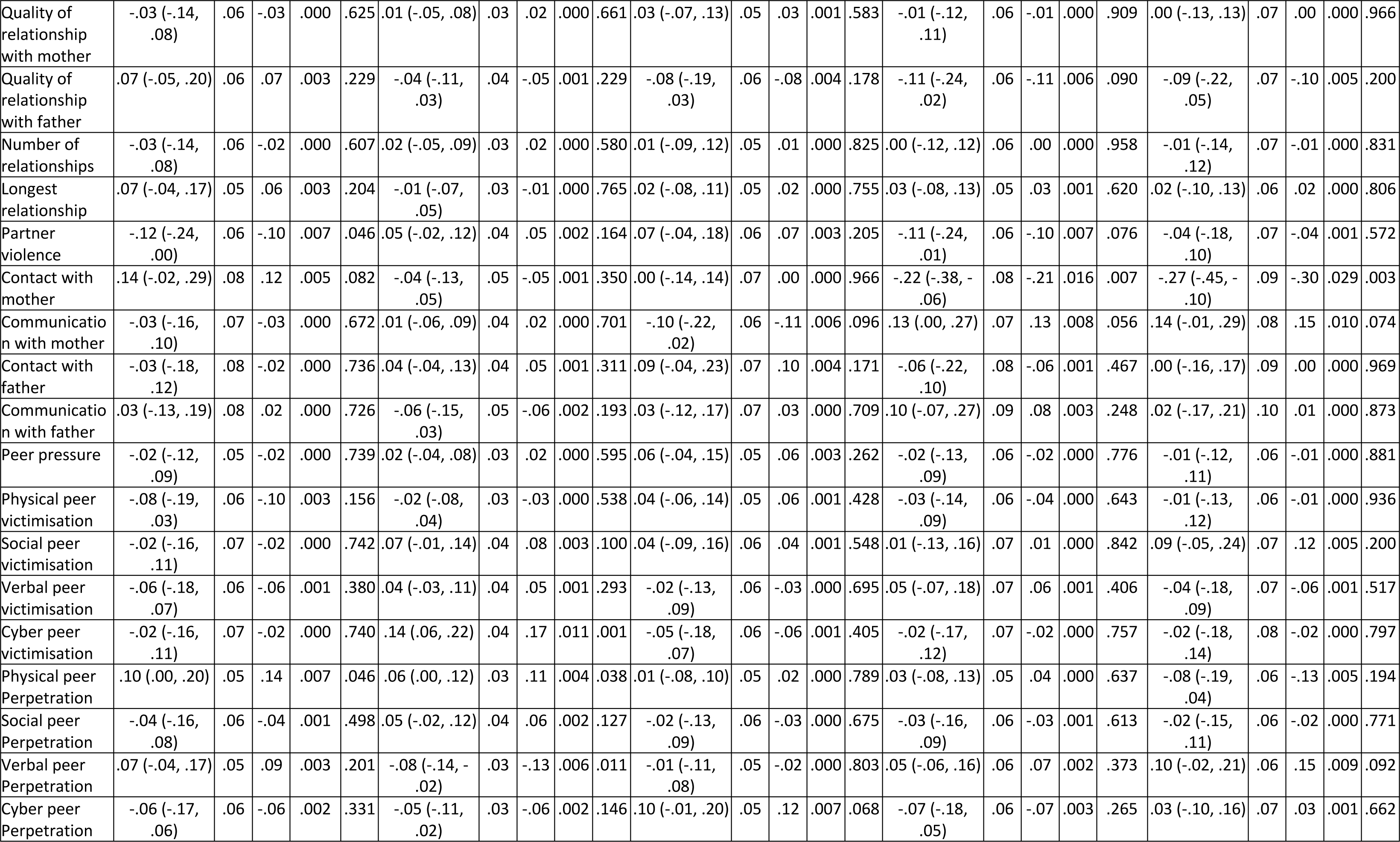
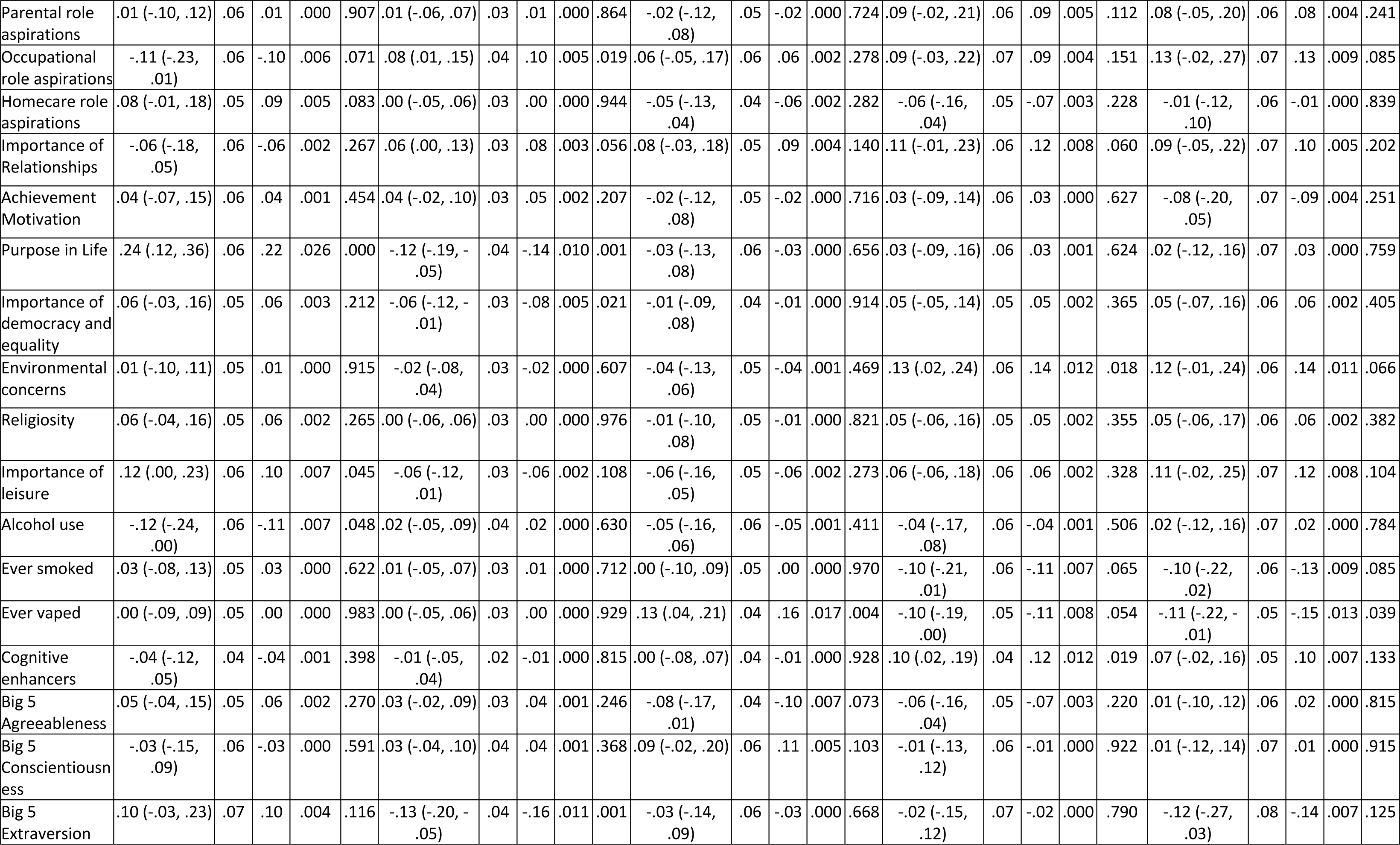
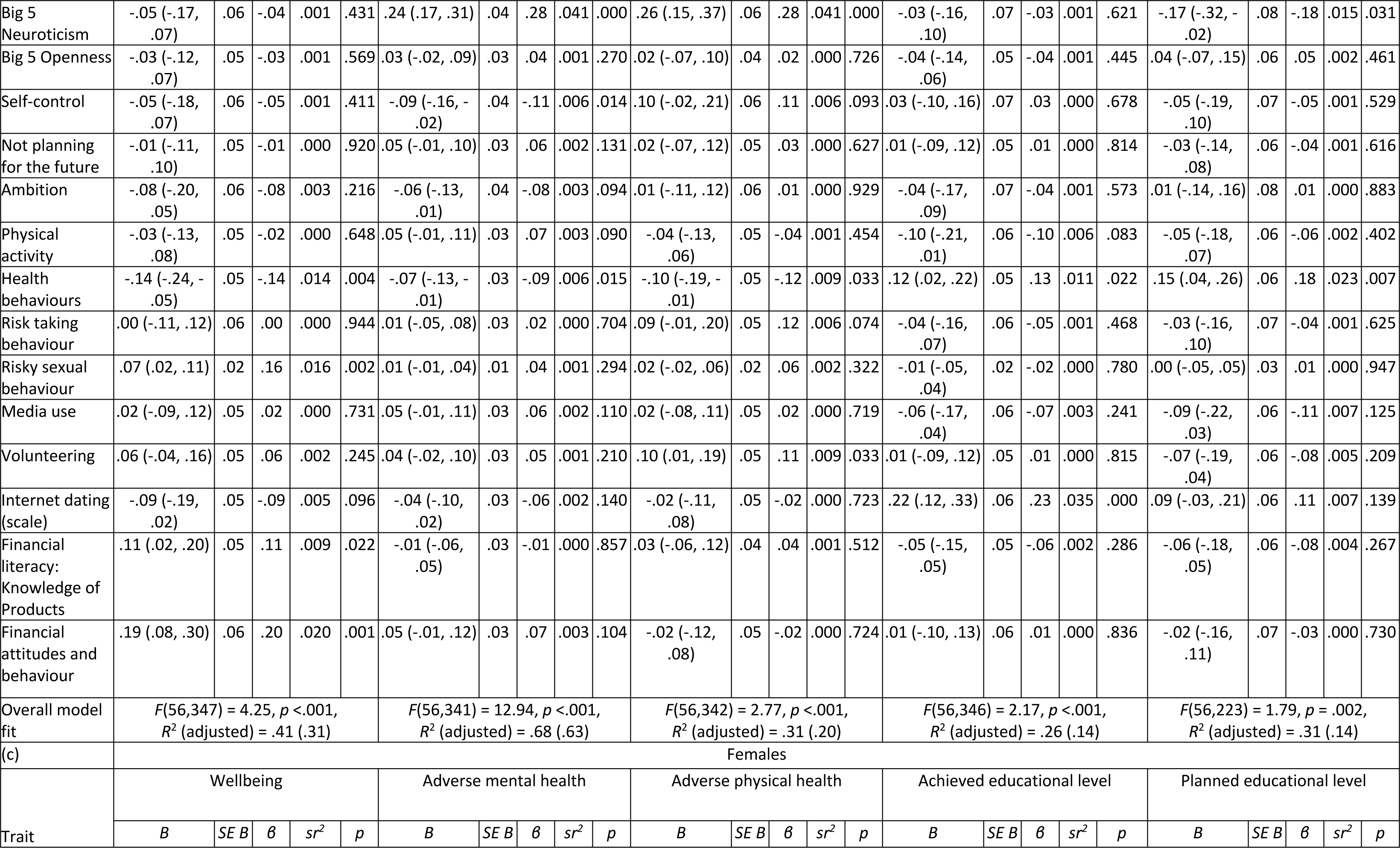
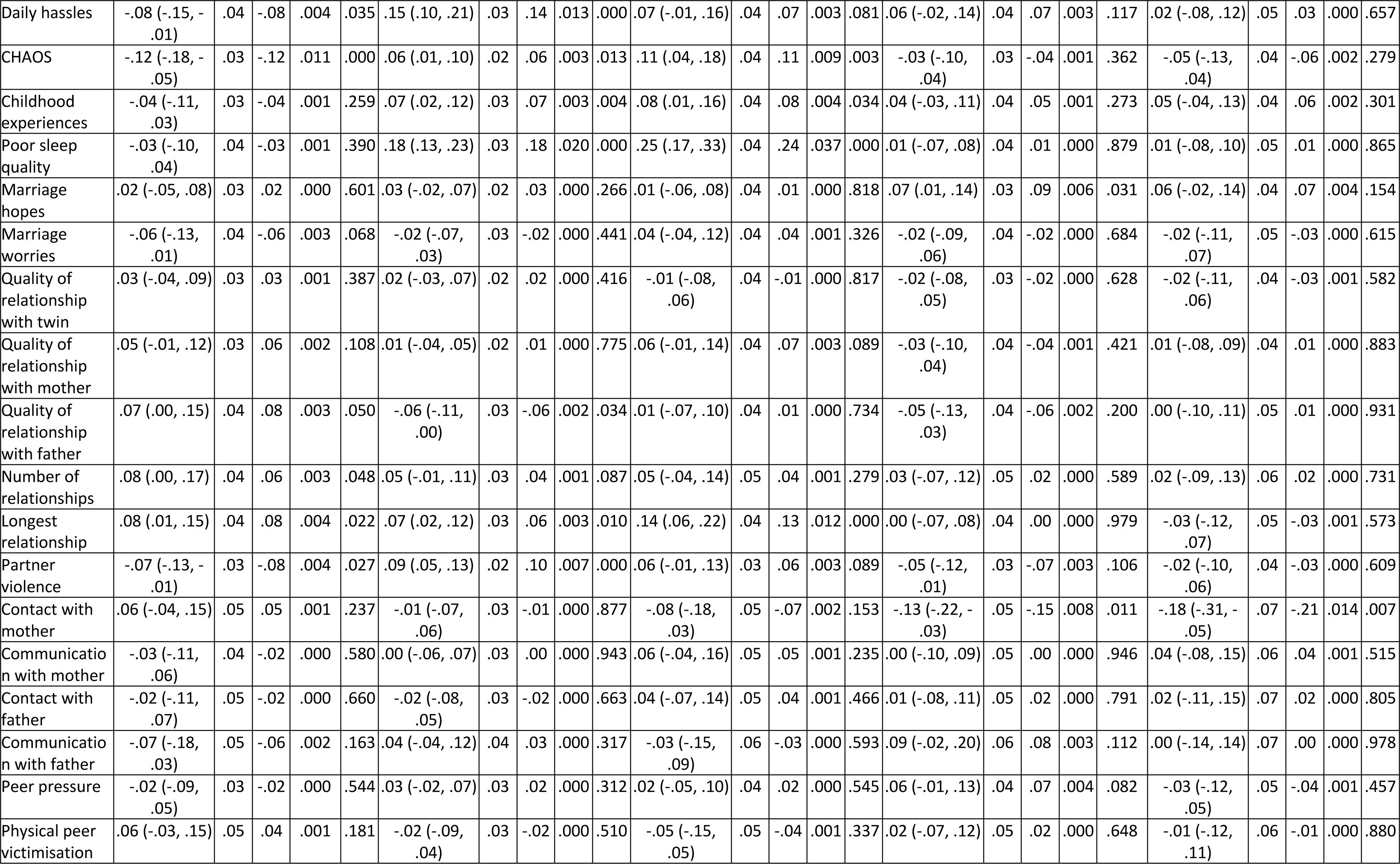
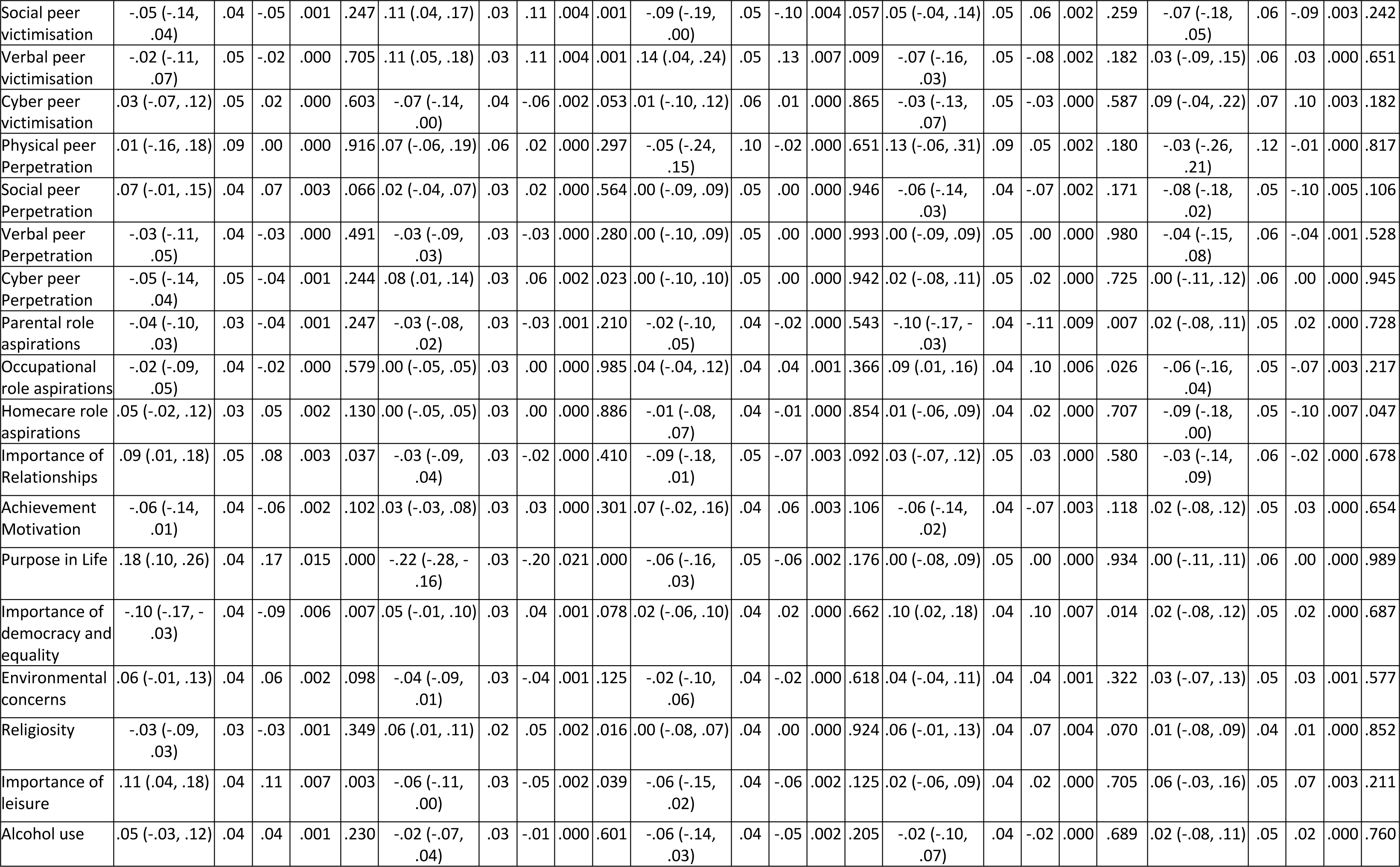
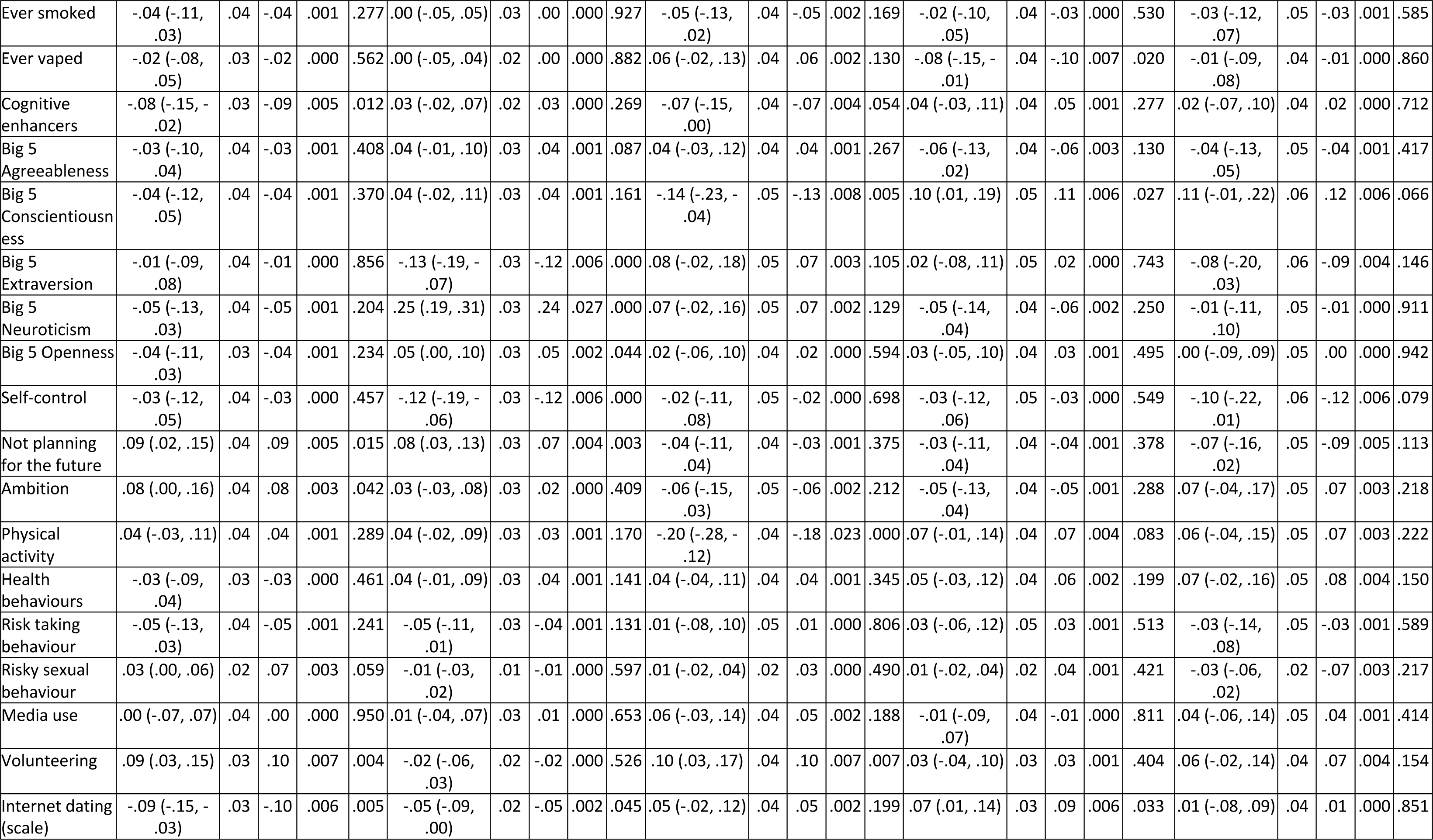
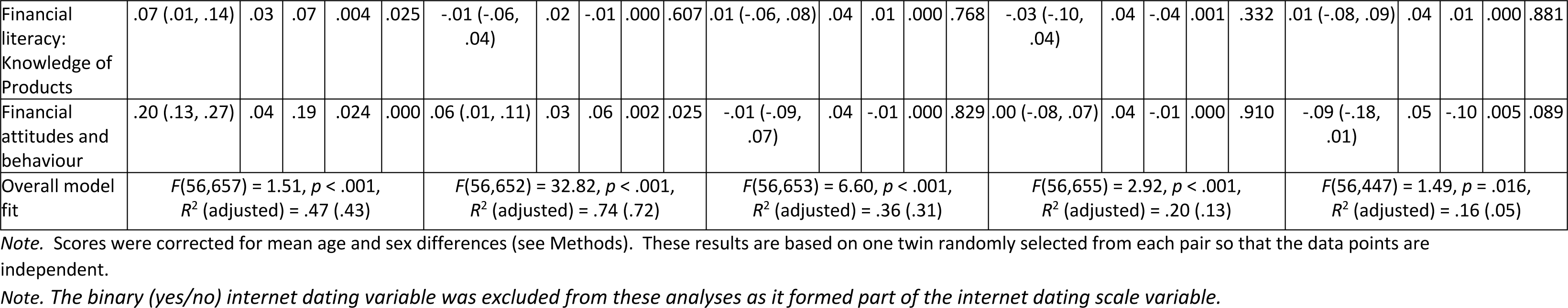
Summary of multiple regression analyses: variance explained in composite scores of adverse physical and adverse mental health, wellbeing and educational attainment psychological traits, for the whole sample (a) and then separately for males (b) and females (c) (95% confidence intervals are in parentheses).

**Table S8.** Twin intraclass correlations and Falconer ACE estimates, and model fitting results for univariate analyses of additive genetic (A), shared environmental (C), and non-shared environmental (E) components of variance for variables for psychological traits and functional outcomes (95% confidence intervals are in parentheses). (a) for the whole sample; (b) males only; (c) females only; (d) MZ and DZ same sex twin pairs (opposite sex DZ twins excluded).

| Trait | MZ |  |  | DZ<sup>a</sup> |  |  | Falconer ACE estimates |  |  |
| --- | --- | --- | --- | --- | --- | --- | --- | --- | --- |
|  | *r* | *p* | *n* | *r* | *p* | *n* | A | C | E |
| Daily hassles | .31<br>(.26,.36) | <.001 | 1376 | .15<br>(.10, .19) | <.001 | 2050 | .31<sup>b</sup> | .00 | .69 |
| CHAOS | .40<br>(.36,.44) | <.001 | 1588 | .36<br>(.32, .39) | <.001 | 2516 | .07 | .32 | .61 |
| Childhood experiences | .66<br>(.63,.69) | <.001 | 1367 | .40<br>(.36, .43) | <.001 | 2039 | .52 | .14 | .34 |
| Poor sleep quality | .31<br>(.26,.36) | <.001 | 1334 | .15<br>(.11, .20) | <.001 | 2001 | .31<sup>b</sup> | .00 | .69 |
| Marriage hopes | .28<br>(.23,.33) | <.001 | 1550 | .18<br>(.14, .22) | <.001 | 2417 | .19 | .09 | .73 |
| Marriage worries | .38<br>(.34,.42) | <.001 | 1549 | .19<br>(.15, .23) | <.001 | 2418 | .37 | .01 | .62 |
| Quality of relationship with twin | .69<br>(.66,.72) | <.001 | 1497 | .62<br>(.59, .64) | <.001 | 2339 | .14 | .55 | .31 |
| Quality of relationship with mother | .53<br>(.49,.57) | <.001 | 1478 | .26<br>(.22, .29) | <.001 | 2308 | .53<sup>b</sup> | .00 | .47 |
| Quality of relationship with father | .61<br>(.58,.64) | <.001 | 1422 | .41<br>(.38, .44) | <.001 | 2229 | .39 | .22 | .39 |
| Number of relationships | .46<br>(.42,.50) | <.001 | 1548 | .19<br>(.15, .23) | <.001 | 2416 | .46<sup>b</sup> | .00 | .54 |
| Longest relationship | .32<br>(.27,.37) | <.001 | 1152 | .08<br>(.03, .12) | .002 | 1736 | .32<sup>b</sup> | .00 | .68 |
| Partner violence | .25<br>(.20,.30) | <.001 | 1457 | .12<br>(.07, .16) | <.001 | 2264 | .25<sup>b</sup> | .00 | .75 |
| Contact with mother | .55<br>(.48,.61) | <.001 | 426 | .30<br>(.22, .37) | <.001 | 559 | .50 | .05 | .45 |
| Communication with mother | .64<br>(.58,.69) | <.001 | 440 | .38<br>(.30, .45) | <.001 | 574 | .54 | .11 | .36 |
| Contact with father | .50<br>(.43,.56) | <.001 | 511 | .40<br>(.33, .46) | <.001 | 694 | .21 | .29 | .50 |
| Communication with father | .73<br>(.69,.77) | <.001 | 575 | .64<br>(.59, .68) | <.001 | 764 | .18 | .54 | .27 |
| Peer pressure | .35<br>(.30,.39) | <.001 | 1494 | .16<br>(.12, .19) | <.001 | 2371 | .35<sup>b</sup> | .00 | .65 |
| Physical peer victimisation | .20<br>(.15,.25) | <.001 | 1308 | .02<br>(-.03, .06) | .417 | 1935 | .20<sup>b</sup> | .00 | .80 |
| Social peer victimisation | .24<br>(.19,.29) | <.001 | 1308 | .14<br>(.10, .18) | <.001 | 1934 | .21 | .03 | .76 |
| Verbal peer victimisation | .29<br>(.24,.34) | <.001 | 1308 | .13<br>(.09, .18) | <.001 | 1934 | .29<sup>b</sup> | .00 | .71 |
| Cyber peer victimisation | .33<br>(.28,.38) | <.001 | 1307 | .12<br>(.07, .16) | <.001 | 1934 | .33<sup>b</sup> | .00 | .67 |
| Physical peer Perpetration | .14<br>(.09,.19) | <.001 | 1308 | .01<br>(-.03, .06) | .569 | 1936 | .14<sup>b</sup> | .00 | .86 |
| Social peer Perpetration | .14<br>(.09,.19) | <.001 | 1307 | .05<br>(.00, .09) | .030 | 1936 | .14<sup>b</sup> | .00 | .86 |
| Verbal peer Perpetration | .32<br>(.27,.37) | <.001 | 1307 | .12<br>(.07, .16) | <.001 | 1936 | .32<sup>b</sup> | .00 | .68 |
| Cyber peer Perpetration | .20<br>(.15,.25) | <.001 | 1307 | .03<br>(-.02, .07) | .247 | 1936 | .20<sup>b</sup> | .00 | .80 |
| Parental role aspirations | .35<br>(.30,.40) | <.001 | 1373 | .16<br>(.11, .20) | <.001 | 2042 | .35<sup>b</sup> | .00 | .65 |
| Occupational role aspirations | .27<br>(.22,.32) | <.001 | 1374 | .11<br>(.07, .15) | <.001 | 2045 | .27<sup>b</sup> | .00 | .73 |
| Homecare role aspirations | .42<br>(.38,.46) | <.001 | 1374 | .17<br>(.13, .21) | <.001 | 2045 | .42<sup>b</sup> | .00 | .58 |
| Importance of Relationships | .41<br>(.37,.45) | <.001 | 1543 | .10<br>(.06, .13) | <.001 | 2434 | .41<sup>b</sup> | .00 | .59 |
| Achievement Motivation | .37<br>(.33,.41) | <.001 | 1543 | .18<br>(.14, .21) | <.001 | 2434 | .37<sup>b</sup> | .00 | .63 |
| Purpose in Life | .42<br>(.38,.46) | <.001 | 1542 | .21<br>(.18, .25) | <.001 | 2432 | .41 | .01 | .58 |
| Importance of democracy and equality | .43<br>(.39,.47) | <.001 | 1541 | .18<br>(.14, .22) | <.001 | 2424 | .43<sup>b</sup> | .00 | .57 |
| Environmental concerns | .51<br>(.47,.55) | <.001 | 1542 | .27<br>(.23, .30) | <.001 | 2428 | .49 | .02 | .49 |
| Religiosity | .72<br>(.69,.74) | <.001 | 1498 | .56<br>(.54, .59) | <.001 | 2374 | .32 | .41 | .28 |
| Importance of leisure | .41<br>(.36,.45) | <.001 | 1338 | .19<br>(.15, .23) | <.001 | 1997 | .41<sup>b</sup> | .00 | .59 |
| Alcohol use | .45<br>(.40,.50) | <.001 | 1126 | .26<br>(.22, .31) | <.001 | 1717 | .38 | .07 | .55 |
| Ever smoked | .49<br>(.45,.53) | <.001 | 1342 | .23<br>(.19, .27) | <.001 | 1995 | .49<sup>b</sup> | .00 | .51 |
| Ever vaped | .36<br>(.31,.41) | <.001 | 1342 | .20<br>(.16, .24) | <.001 | 1990 | .31 | .04 | .64 |
| Cognitive enhancers | .33<br>(.28,.38) | <.001 | 1322 | .18<br>(.14, .23) | <.001 | 1955 | .29 | .04 | .67 |
| Big 5 Agreeableness | .27<br>(.22,.32) | <.001 | 1419 | .03<br>(-.01, .07) | .117 | 2266 | .27<sup>b</sup> | .00 | .73 |
| Big 5 Conscientiousness | .34<br>(.29,.39) | <.001 | 1418 | .07<br>(.03, .11) | .001 | 2262 | .34<sup>b</sup> | .00 | .66 |
| Big 5 Extraversion | .44<br>(.40,.48) | <.001 | 1424 | .13<br>(.09, .17) | <.001 | 2276 | .44<sup>b</sup> | .00 | .56 |
| Big 5 Neuroticism | .37<br>(.32,.41) | <.001 | 1426 | .18<br>(.14, .22) | <.001 | 2280 | .37<sup>b</sup> | .00 | .63 |
| Big 5 Openness | .40<br>(.36,.44) | <.001 | 1421 | .11<br>(.07, .15) | <.001 | 2271 | .40<sup>b</sup> | .00 | .60 |
| Self-control | .40<br>(.36,.44) | <.001 | 1417 | .14<br>(.10, .18) | <.001 | 2259 | .40<sup>b</sup> | .00 | .60 |
| Not planning for the future | .25<br>(.20,.30) | <.001 | 1416 | .13<br>(.09, .17) | <.001 | 2258 | .24 | .01 | .75 |
| Ambition | .37<br>(.32,.41) | <.001 | 1376 | .16<br>(.11, .20) | <.001 | 2051 | .37<sup>b</sup> | .00 | .63 |
| Physical activity | .40<br>(.36,.44) | <.001 | 1528 | .17<br>(.13, .21) | <.001 | 2371 | .40<sup>b</sup> | .00 | .60 |
| Health behaviours | .54<br>(.50,.57) | <.001 | 1533 | .29<br>(.25, .33) | <.001 | 2391 | .50 | .04 | .46 |
| Risk taking behaviour | .48<br>(.44,.52) | <.001 | 1413 | .23<br>(.19, .27) | <.001 | 2258 | .48<sup>b</sup> | .00 | .52 |
| Risky sexual behaviour | .61<br>(.58,.64) | <.001 | 1413 | .29<br>(.25, .33) | <.001 | 2206 | .61<sup>b</sup> | .00 | .39 |
| Media use | .53<br>(.49,.57) | <.001 | 1520 | .19<br>(.15, .22) | <.001 | 2350 | .53<sup>b</sup> | .00 | .47 |
| Volunteering | .45<br>(.41,.49) | <.001 | 1500 | .20<br>(.16, .24) | <.001 | 2378 | .45<sup>b</sup> | .00 | .55 |
| Internet dating<br>(scale) | .34<br>(.29,.38) | <.001 | 1512 | .17<br>(.13, .21) | <.001 | 2337 | .34<sup>b</sup> | .00 | .66 |
| Internet dating<br>(y/n) | .32<br>(.27,.36) | <.001 | 1519 | .15<br>(.11, .18) | <.001 | 2351 | .32<sup>b</sup> | .00 | .68 |
| Financial literacy: Knowledge of<br>Products | .38<br>(.34,.42) | <.001 | 1481 | .15<br>(.11, .19) | <.001 | 2266 | .38<sup>b</sup> | .00 | .62 |
| Financial attitudes and behaviour | .30<br>(.25,.35) | <.001 | 1480 | .09<br>(.05, .13) | <.001 | 2266 | .30<sup>b</sup> | .00 | .70 |
| Functional outcome | MZ |  |  | DZ<sup>a</sup> |  |  | Falconer ACE estimates |  |  |
|  | *r* | *p* | *n* | *r* | *p* | *n* | A | C | E |
| Wellbeing factor (1st PC) | .39<br>(.35, .43) | <.001 | 1477 | .23<br>(.19, .27) | <.001 | 2249 | .32 | .07 | .61 |
| Adverse mental health factor (1st<br>PC) | .50<br>(.45, .54) | <.001 | 1042 | .21<br>(.17, .26) | <.001 | 1578 | .50<sup>b</sup> | .00 | .50 |
| Adverse physical health factor (1st<br>PC) | .40<br>(.35, .45) | <.001 | 1234 | .17<br>(.13, .22) | <.001 | 1786 | .40<sup>b</sup> | .00 | .60 |
| Achieved educational level | .40<br>(.36, .44) | <.001 | 1520 | .28<br>(.24, .32) | <.001 | 2365 | .24 | .16 | .60 |
| Planned educational level | .37<br>(.31, .43) | <.001 | 735 | .11<br>(.05, .17) | <.001 | 1068 | .37<sup>b</sup> | .00 | .63 |

*Note.* MZ=monozygotic; DZ=dizygotic; and genetic (A), shared environmental (C) and non-shared environmental (E) estimates. Scores were corrected for mean age and sex differences (see Methods). Falconer's ACE estimates were derived from the intraclass twin correlations using Falconer's formula (Rijsdijk & Sham, 2002).
<sup>a</sup> Same-sex and opposite-sex DZ twins combined.
<sup>b</sup> rMZ reported as A would otherwise exceed rMZ.

(b) Males only
| Trait | MZ |  |  | DZ<sup>a</sup> |  |  | Falconer ACE estimates |  |  |
| --- | --- | --- | --- | --- | --- | --- | --- | --- | --- |
|  | *r* | *p* | *n* | *r* | *p* | *n* | A | C | E |
| Daily hassles | .31<br>(.22, .39) | <.001 | 445 | .07<br>(-.04, .17) | .230 | 344 | .31<sup>b</sup> | .00 | .69 |
| CHAOS | .38<br>(.30, .45) | <.001 | 523 | .24<br>(.15, .33) | <.001 | 438 | .28 | .11 | .62 |
| Childhood experiences | .52<br>(.45, .59) | <.001 | 439 | .35<br>(.25, .44) | <.001 | 336 | .35 | .17 | .48 |
| Poor sleep quality | .26<br>(.17, .35) | <.001 | 432 | .19<br>(.08, .29) | <.001 | 334 | .13 | .12 | .74 |
| Marriage hopes | .14<br>(.05, .22) | .001 | 512 | .14<br>(.05, .23) | .005 | 424 | .01 | .13 | .86 |
| Marriage worries | .37<br>(.29, .44) | <.001 | 511 | .15<br>(.06, .24) | .002 | 424 | .37<sup>b</sup> | .00 | .63 |
| Quality of relationship with twin | .70<br>(.65, .74) | <.001 | 488 | .62<br>(.56, .68) | <.001 | 409 | .16 | .54 | .30 |
| Quality of relationship with mother | .48<br>(.41, .55) | <.001 | 479 | .15<br>(.05, .24) | .003 | 407 | .48<sup>b</sup> | .00 | .53 |
| Quality of relationship with father | .56<br>(.49, .62) | <.001 | 461 | .34<br>(.25, .43) | <.001 | 386 | .44 | .12 | .44 |
| Number of relationships | .50<br>(.43, .56) | <.001 | 512 | .21<br>(.12, .30) | <.001 | 423 | .50<sup>b</sup> | .00 | .50 |
| Longest relationship | .26<br>(.16, .35) | <.001 | 368 | .13<br>(.02, .24) | .030 | 298 | .26<sup>b</sup> | .00 | .74 |
| Partner violence | .24<br>(.15, .32) | <.001 | 471 | .10<br>(.00, .20) | .060 | 389 | .24<sup>b</sup> | .00 | .76 |
| Contact with mother | .48<br>(.34, .60) | <.001 | 132 | .40<br>(.21, .56) | <.001 | 88 | .15 | .33 | .52 |
| Communication with mother | .58<br>(.46, .68) | <.001 | 136 | .44<br>(.26, .59) | <.001 | 91 | .29 | .29 | .42 |
| Contact with father | .46<br>(.33, .57) | <.001 | 159 | .43<br>(.26, .57) | <.001 | 106 | .07 | .39 | .54 |
| Communication with father | .69<br>(.60, .76) | <.001 | 179 | .49<br>(.34, .62) | <.001 | 116 | .41 | .28 | .31 |
| Peer pressure | .40<br>(.32, .47) | <.001 | 495 | .18<br>(.09, .27) | <.001 | 422 | .40<sup>b</sup> | .00 | .60 |
| Physical peer victimisation | .25<br>(.16, .34) | <.001 | 427 | .04<br>(-.07, .15) | .486 | 331 | .25<sup>b</sup> | .00 | .75 |
| Social peer victimisation | .13<br>(.04, .22) | .005 | 427 | .26<br>(.16, .36) | <.001 | 331 | .00 | .13 | .87 |
| Verbal peer victimisation | .24<br>(.15, .33) | <.001 | 427 | .13<br>(.02, .23) | .017 | 329 | .22 | .02 | .76 |
| Cyber peer victimisation | .23<br>(.14, .32) | <.001 | 427 | .13<br>(.02, .23) | .017 | 330 | .19 | .04 | .77 |
| Physical peer Perpetration | .17<br>(.08, .26) | .001 | 426 | .09<br>(-.02, .20) | .115 | 331 | .16 | .01 | .84 |
| Social peer Perpetration | .16<br>(.07, .25) | .001 | 426 | .00<br>(-.11, .11) | .993 | 331 | .16<sup>b</sup> | .00 | .85 |
| Verbal peer Perpetration | .36<br>(.27, .44) | <.001 | 426 | .11<br>(.00, .22) | .053 | 331 | .36<sup>b</sup> | .00 | .64 |
| Cyber peer Perpetration | .18<br>(.09, .27) | <.001 | 426 | -.04<br>(-.15, .07) | .482 | 331 | .18<sup>b</sup> | .00 | .82 |
| Parental role aspirations | .29<br>(.20, .37) | <.001 | 442 | .02<br>(-.09, .13) | .660 | 337 | .29<sup>b</sup> | .00 | .71 |
| Occupational role aspirations | .23<br>(.14, .32) | <.001 | 442 | .10<br>(-.01, .20) | .067 | 337 | .23<sup>b</sup> | .00 | .77 |
| Homecare role aspirations | .46<br>(.38, .53) | <.001 | 442 | .20<br>(.10, .30) | <.001 | 337 | .46<sup>b</sup> | .00 | .54 |
| Importance of Relationships | .47<br>(.40, .54) | <.001 | 510 | .06<br>(-.04, .15) | .199 | 425 | .47<sup>b</sup> | .00 | .53 |
| Achievement Motivation | .34<br>(.26, .41) | <.001 | 510 | .24<br>(.15, .33) | <.001 | 425 | .20 | .14 | .66 |
| Purpose in Life | .43<br>(.36, .50) | <.001 | 510 | .30<br>(.21, .38) | <.001 | 425 | .26 | .17 | .57 |
| Importance of democracy and equality | .44<br>(.37, .51) | <.001 | 511 | .24<br>(.15, .33) | <.001 | 423 | .40 | .04 | .56 |
| Environmental concerns | .55<br>(.49, .61) | <.001 | 511 | .29<br>(.20, .37) | <.001 | 425 | .54 | .02 | .45 |
| Religiosity | .74<br>(.70, .78) | <.001 | 497 | .62<br>(.56, .68) | <.001 | 424 | .24 | .50 | .26 |
| Importance of leisure | .38<br>(.30, .46) | <.001 | 432 | .18<br>(.07, .28) | .001 | 331 | .38<sup>b</sup> | .00 | .63 |
| Alcohol use | .40<br>(.31, .48) | <.001 | 369 | .22<br>(.11, .33) | <.001 | 292 | .36 | .04 | .60 |
| Ever smoked | .48<br>(.40, .55) | <.001 | 433 | .24<br>(.14, .34) | <.001 | 333 | .47 | .01 | .53 |
| Ever vaped | .36<br>(.28, .44) | <.001 | 432 | .26<br>(.16, .36) | <.001 | 332 | .20 | .16 | .64 |
| Cognitive enhancers | .31<br>(.22, .39) | <.001 | 420 | .17<br>(.06, .27) | .003 | 325 | .28 | .02 | .69 |
| Big 5 Agreeableness | .32<br>(.24, .40) | <.001 | 483 | .10<br>(.00, .20) | .049 | 404 | .32<sup>b</sup> | .00 | .68 |
| Big 5 Conscientiousness | .38<br>(.30, .45) | <.001 | 482 | .13<br>(.03, .22) | .009 | 404 | .38<sup>b</sup> | .00 | .62 |
| Big 5 Extraversion | .47<br>(.40, .54) | <.001 | 485 | .15<br>(.05, .24) | .003 | 408 | .47<sup>b</sup> | .00 | .53 |
| Big 5 Neuroticism | .31<br>(.23, .39) | <.001 | 485 | .20<br>(.11, .29) | <.001 | 410 | .21 | .10 | .69 |
| Big 5 Openness | .41<br>(.33, .48) | <.001 | 484 | .17<br>(.07, .26) | <.001 | 407 | .41<sup>b</sup> | .00 | .59 |
| Self-control | .37<br>(.29, .44) | <.001 | 482 | .14<br>(.04, .23) | .006 | 401 | .37<sup>b</sup> | .00 | .63 |
| Not planning for the future | .31<br>(.23, .39) | <.001 | 481 | .13<br>(.03, .23) | .011 | 401 | .31<sup>b</sup> | .00 | .69 |
| Ambition | .37<br>(.29, .45) | <.001 | 445 | .15<br>(.05, .25) | .005 | 344 | .37<sup>b</sup> | .00 | .63 |
| Physical activity | .41<br>(.33, .48) | <.001 | 501 | .17<br>(.08, .26) | .001 | 416 | .41<sup>b</sup> | .00 | .59 |
| Health behaviours | .53<br>(.46, .59) | <.001 | 504 | .25<br>(.16, .34) | <.001 | 417 | .53<sup>b</sup> | .00 | .47 |
| Risk taking behaviour | .53<br>(.46, .59) | <.001 | 479 | .24<br>(.15, .33) | <.001 | 400 | .53<sup>b</sup> | .00 | .47 |
| Risky sexual behaviour | .61<br>(.55, .66) | <.001 | 468 | .31<br>(.22, .40) | <.001 | 389 | .59 | .01 | .39 |
| Media use | .57<br>(.51, .63) | <.001 | 498 | .12<br>(.02, .21) | .014 | 413 | .57<sup>b</sup> | .00 | .44 |
| Volunteering | .49<br>(.42, .55) | <.001 | 497 | .27<br>(.18, .36) | <.001 | 424 | .44 | .05 | .51 |
| Internet dating (scale) | .36<br>(.28, .43) | <.001 | 498 | .19<br>(.10, .28) | <.001 | 414 | .34 | .02 | .64 |
| Internet dating (y/n) | .33<br>(.25, .41) | <.001 | 499 | .21<br>(.12, .30) | <.001 | 416 | .25 | .08 | .67 |
| Financial literacy: Knowledge of Products | .48<br>(.41, .54) | <.001 | 500 | .27<br>(.18, .36) | <.001 | 407 | .42 | .06 | .52 |
| Financial attitudes and behaviour | .30<br>(.22, .38) | <.001 | 500 | .14<br>(.04, .23) | .004 | 407 | .30<sup>b</sup> | .00 | .70 |
| Functional outcome | MZ |  |  | DZ<sup>a</sup> |  |  | Falconer ACE estimates |  |  |
|  | *r* | *p* | *n* | *r* | *p* | *n* | A | C | E |
| Wellbeing factor (1st PC) | .34<br>(.26, .42) | <.001 | 497 | .25<br>(.16, .34) | <.001 | 404 | .18 | .16 | .66 |
| Adverse mental health factor (1st PC) | .37<br>(.27, .46) | <.001 | 338 | .22<br>(.10, .33) | <.001 | 271 | .29 | .08 | .64 |
| Adverse physical health factor (1st PC) | .39<br>(.30, .47) | <.001 | 383 | .13<br>(.02, .24) | .020 | 302 | .39<sup>b</sup> | .00 | .61 |
| Achieved educational level | .40<br>(.32, .47) | <.001 | 502 | .19<br>(.10, .28) | <.001 | 412 | .40<sup>b</sup> | .00 | .60 |
| Planned educational level | .43<br>(.32, .53) | <.001 | 247 | .16<br>(.01, .30) | .033 | 172 | .43<sup>b</sup> | .00 | .57 |

*Note.* MZ=monozygotic; DZ=dizygotic; and genetic (A), shared environmental (C) and non-shared environmental (E) estimates. Scores were corrected for mean age and sex differences (see Methods). Falconer's ACE estimates were derived from the intraclass twin correlations using Falconer's formula (Rijsdijk & Sham, 2002).
<sup>a</sup> Same-sex male DZ twins.
<sup>b</sup> rMZ reported as A would otherwise exceed rMZ.

(c) females only
|  |  |  |  |  |  |  |  |  |  |
| --- | --- | --- | --- | --- | --- | --- | --- | --- | --- |
| Trait | MZ |  |  | DZ<sup>a</sup> |  |  | Falconer ACE estimates |  |  |
|  | *r* | *p* | *n* | *r* | *p* | *n* | A | C | E |
| Daily hassles | .32<br>(.26, .38) | <.001 | 931 | .16<br>(.09, .23) | <.001 | 757 | .32<sup>b</sup> | .00 | .68 |
| CHAOS | .40<br>(.35, .45) | <.001 | 1065 | .43<br>(.37, .48) | <.001 | 873 | .00 | .40 | .60 |
| Childhood experiences | .70<br>(.67, .73) | <.001 | 928 | .47<br>(.41, .52) | <.001 | 752 | .46 | .24 | .30 |
| Poor sleep quality | .33<br>(.27, .39) | <.001 | 902 | .21<br>(.14, .28) | <.001 | 740 | .26 | .08 | .67 |
| Marriage hopes | .34<br>(.29, .39) | <.001 | 1038 | .23<br>(.16, .29) | <.001 | 833 | .21 | .13 | .66 |
| Marriage worries | .38<br>(.33, .43) | <.001 | 1038 | .24<br>(.17, .30) | <.001 | 833 | .29 | .10 | .62 |
| Quality of relationship with twin | .69<br>(.66, .72) | <.001 | 1009 | .64<br>(.60, .68) | <.001 | 813 | .09 | .60 | .32 |
| Quality of relationship with mother | .56<br>(.52, .60) | <.001 | 999 | .35<br>(.29, .41) | <.001 | 799 | .41 | .15 | .45 |
| Quality of relationship with father | .63<br>(.59, .67) | <.001 | 961 | .47<br>(.41, .52) | <.001 | 770 | .32 | .31 | .37 |
| Number of relationships | .44<br>(.39, .49) | <.001 | 1036 | .26<br>(.20, .32) | <.001 | 833 | .34 | .09 | .57 |
| Longest relationship | .34<br>(.28, .40) | <.001 | 784 | .12<br>(.04, .20) | .005 | 600 | .34<sup>b</sup> | .00 | .66 |
| Partner violence | .25<br>(.19, .31) | <.001 | 986 | .11<br>(.04, .18) | .003 | 794 | .25<sup>b</sup> | .00 | .75 |
| Contact with mother | .57<br>(.49, .64) | <.001 | 294 | .34<br>(.22, .45) | <.001 | 224 | .46 | .11 | .43 |
| Communication with mother | .67<br>(.60, .73) | <.001 | 304 | .33<br>(.21, .44) | <.001 | 228 | .67<sup>b</sup> | .00 | .33 |
| Contact with father | .52<br>(.44, .59) | <.001 | 352 | .45<br>(.35, .54) | <.001 | 271 | .15 | .37 | .48 |
| Communication with father | .74<br>(.69, .78) | <.001 | 396 | .70<br>(.64, .75) | <.001 | 298 | .08 | .66 | .26 |
| Peer pressure | .31<br>(.25, .37) | <.001 | 999 | .15<br>(.08, .22) | <.001 | 812 | .31<sup>b</sup> | .00 | .69 |
| Physical peer victimisation | .09<br>(.02, .16) | .006 | 881 | .01<br>(-.06, .08) | .776 | 712 | .09<sup>b</sup> | .00 | .91 |
| Social peer victimisation | .29<br>(.23, .35) | <.001 | 881 | .14<br>(.07, .21) | <.001 | 712 | .29<sup>b</sup> | .00 | .71 |
| Verbal peer victimisation | .32<br>(.26, .38) | <.001 | 881 | .16<br>(.09, .23) | <.001 | 713 | .32 | .00 | .68 |
| Cyber peer victimisation | .37<br>(.31, .43) | <.001 | 880 | .06<br>(-.01, .13) | .129 | 713 | .37<sup>b</sup> | .00 | .63 |
| Physical peer Perpetration | .08<br>(.01, .15) | .016 | 882 | -.02<br>(-.09, .05) | .551 | 714 | .08<sup>b</sup> | .00 | .92 |
| Social peer Perpetration | .13<br>(.06, .19) | <.001 | 881 | .15<br>(.08, .22) | <.001 | 714 | .00 | .13 | .87 |
| Verbal peer Perpetration | .27<br>(.21, .33) | <.001 | 881 | .16<br>(.09, .23) | <.001 | 714 | .22 | .05 | .73 |
| Cyber peer Perpetration | .22<br>(.16, .28) | <.001 | 881 | .14<br>(.07, .21) | <.001 | 714 | .17 | .05 | .78 |
| Parental role aspirations | .38<br>(.32, .43) | <.001 | 931 | .21<br>(.14, .28) | <.001 | 754 | .33 | .04 | .63 |
| Occupational role aspirations | .30<br>(.24, .36) | <.001 | 932 | .15<br>(.08, .22) | <.001 | 755 | .29 | .00 | .70 |
| Homecare role aspirations | .39<br>(.33, .44) | <.001 | 932 | .24<br>(.17, .31) | <.001 | 755 | .29 | .10 | .61 |
| Importance of Relationships | .38<br>(.33, .43) | <.001 | 1033 | .13<br>(.06, .20) | <.001 | 840 | .38<sup>b</sup> | .00 | .63 |
| Achievement Motivation | .38 (.33, .43) | <.001 | 1033 | .15<br>(.08, .22) | <.001 | 840 | .38<sup>b</sup> | .00 | .62 |
| Purpose in Life | .41<br>(.36, .46) | <.001 | 1032 | .20<br>(.13, .26) | <.001 | 839 | .41<sup>b</sup> | .00 | .59 |
| Importance of democracy and equality | .42<br>(.37, .47) | <.001 | 1030 | .14<br>(.07, .21) | <.001 | 836 | .42<sup>b</sup> | .00 | .58 |
| Environmental concerns | .49<br>(.44, .54) | <.001 | 1031 | .30<br>(.24, .36) | <.001 | 838 | .38 | .11 | .51 |
| Religiosity | .71<br>(.68, .74) | <.001 | 1001 | .56<br>(.51, .61) | <.001 | 813 | .32 | .40 | .29 |
| Importance of leisure | .43<br>(.38, .48) | <.001 | 906 | .23<br>(.16, .30) | <.001 | 743 | .39 | .04 | .57 |
| Alcohol use | .49<br>(.43, .54) | <.001 | 757 | .31<br>(.24, .38) | <.001 | 624 | .34 | .14 | .52 |
| Ever smoked | .50<br>(.45, .55) | <.001 | 909 | .23<br>(.16, .30) | <.001 | 733 | .50<sup>b</sup> | .00 | .50 |
| Ever vaped | .36<br>(.30, .42) | <.001 | 910 | .20<br>(.13, .27) | <.001 | 734 | .32 | .04 | .65 |
| Cognitive enhancers | .34<br>(.28, .40) | <.001 | 902 | .24<br>(.17, .31) | <.001 | 725 | .20 | .14 | .66 |
| Big 5 Agreeableness | .24<br>(.18, .30) | <.001 | 936 | -.02<br>(-.09, .05) | .643 | 768 | .24<sup>b</sup> | .00 | .76 |
| Big 5 Conscientiousness | .32<br>(.26, .38) | <.001 | 936 | .03<br>(-.04, .10) | .368 | 766 | .32<sup>b</sup> | .00 | .68 |
| Big 5 Extraversion | .43<br>(.38, .48) | <.001 | 939 | .09<br>(.02, .16) | .009 | 771 | .43<sup>b</sup> | .00 | .58 |
| Big 5 Neuroticism | .40<br>(.34, .45) | <.001 | 941 | .21<br>(.14, .28) | <.001 | 772 | .37 | .02 | .61 |
| Big 5 Openness | .40<br>(.34, .45) | <.001 | 937 | .10<br>(.03, .17) | .007 | 770 | .40<sup>b</sup> | .00 | .60 |
| Self-control | .41<br>(.36, .46) | <.001 | 935 | .15<br>(.08, .22) | <.001 | 766 | .41<sup>b</sup> | .00 | .59 |
| Not planning for the future | .22<br>(.16, .28) | <.001 | 935 | .11 (.04, .18) | .002 | 766 | .22 | .00 | .78 |
| Ambition | .37<br>(.31, .42) | <.001 | 931 | .16<br>(.09, .23) | <.001 | 757 | .37<sup>b</sup> | .00 | .63 |
| Physical activity | .40<br>(.35, .45) | <.001 | 1027 | .21<br>(.14, .27) | <.001 | 823 | .40 | .01 | .60 |
| Health behaviours | .55<br>(.51, .59) | <.001 | 1029 | .34<br>(.28, .40) | <.001 | 831 | .41 | .14 | .45 |
| Risk taking behaviour | .45<br>(.40, .50) | <.001 | 934 | .23<br>(.16, .30) | <.001 | 768 | .45<sup>b</sup> | .00 | .55 |
| Risky sexual behaviour | .61 (.57, .65) | <.001 | 945 | .33<br>(.26, .39) | <.001 | 755 | .57 | .04 | .39 |
| Media use | .51 (.46, .55) | <.001 | 1022 | .21<br>(.14, .27) | <.001 | 813 | .51<sup>b</sup> | .00 | .49 |
| Volunteering | .43 (.38, .48) | <.001 | 1003 | .19<br>(.12, .26) | <.001 | 814 | .43<sup>b</sup> | .00 | .57 |
| Internet dating<br>(scale) | .32 (.26, .37) | <.001 | 1014 | .19<br>(.12, .26) | <.001 | 811 | .27 | .05 | .68 |
| Internet dating<br>(y/n) | .31 (.25, .36) | <.001 | 1020 | .18<br>(.11, .25) | <.001 | 815 | .25 | .06 | .69 |
| Financial literacy: Knowledge of<br>Products | .31 (.25, .37) | <.001 | 981 | .15<br>(.08, .22) | <.001 | 777 | .31<sup>b</sup> | .00 | .69 |
| Financial attitudes and behaviour | .30 (.24, .36) | <.001 | 980 | .08<br>(.01, .15) | .025 | 777 | .30<sup>b</sup> | .00 | .70 |

| Functional outcome | MZ |  |  | DZ<sup>a</sup> |  |  | Falconer ACE estimates |  |  |
| --- | --- | --- | --- | --- | --- | --- | --- | --- | --- |
|  | *r* | *p* | *n* | *r* | *p* | *n* | A | C | E |
| Wellbeing factor (1st PC) | .41<br>(.36, .46) | <.001 | 980 | .31<br>(.24, .37) | <.001 | 770 | .21 | .21 | .59 |
| Adverse mental health factor (1st PC) | .54<br>(.49, .59) | <.001 | 704 | .26<br>(.18, .33) | <.001 | 584 | .54<sup>b</sup> | .00 | .46 |
| Adverse physical health factor (1st PC) | .40<br>(.34, .46) | <.001 | 851 | .25<br>(.18, .32) | <.001 | 673 | .31 | .09 | .60 |
| Achieved educational level | .40<br>(.35, .45) | <.001 | 1018 | .31<br>(.25, .37) | <.001 | 816 | .17 | .23 | .60 |
| Planned educational level | .33<br>(.25, .41) | <.001 | 488 | .14<br>(.04, .23) | .007 | 399 | .33<sup>b</sup> | .00 | .67 |

*Note.* MZ=monozygotic; DZ=dizygotic; and genetic (A), shared environmental (C) and non-shared environmental (E) estimates. Scores were corrected for mean age and sex differences (see Methods). Falconer's ACE estimates were derived from the intraclass twin correlations using Falconer's formula (Rijsdijk & Sham, 2002).
<sup>a</sup> Same-sex female DZ twins.
<sup>b</sup> rMZ reported as A would otherwise exceed rMZ.

**Table S9.** Univariate twin analyses presenting additive genetic (A), shared environmental (C), and non-shared environmental (E) components of variance EA traits and functional outcomes (95% confidence intervals are in parentheses).

| Trait | Model fitting results |  |  |  |  |  |  |  |  |  |  |  |
| --- | --- | --- | --- | --- | --- | --- | --- | --- | --- | --- | --- | --- |
|  | Whole sample |  |  | Males only |  |  | Females only |  |  | Same-sex MZ and DZ twins |  |  |
|  | A | C | E | A | C | E | A | C | E | A | C | E |
| Daily hassles | .31<br>(.19,.35) | .00<br>(.00,.09) | .69<br>(.65,.74) | .28<br>(.13,.35) | .00<br>(.00,.00) | .72<br>(.65,.79) | .28<br>(.13,.37) | .03<br>(.00,.15) | .69<br>(.63,.74) | .31<br>(.19,.35) | .00<br>(.00,.10) | .69<br>(.65,.74) |
| CHAOS | .12<br>(.01,.22) | .30<br>(.22,.37) | .59<br>(.55,.63) | .28<br>(.10,.45) | .13<br>(.00,.26) | .59<br>(.52,.66) | .03<br>(.00,.15) | .39<br>(.29,.44) | .58<br>(.54,.62) | .11<br>(.00,.22) | .31<br>(.21,.40) | .58<br>(.54,.62) |
| Childhood experiences | .46<br>(.37,.54) | .19<br>(.12,.27) | .35<br>(.32,.38) | .50<br>(.33,.63) | .08<br>(.00,.20) | .43<br>(.37,.49) | .40<br>(.30,.50) | .29<br>(.19,.37) | .32<br>(.29,.35) | .39<br>(.29,.49) | .27<br>(.17,.36) | .34<br>(.31,.37) |
| Poor sleep quality | .27<br>(.14,.35) | .04<br>(.00,.14) | .70<br>(.65,.75) | .23<br>(.01,.33) | .02<br>(.00,.18) | .75<br>(.67,.84) | .27<br>(.11,.38) | .05<br>(.00,.18) | .67<br>(.62,.73) | .16<br>(.01,.32) | .14<br>(.01,.27) | .69<br>(.65,.74) |
| Marriage hopes | .19<br>(.07,.31) | .09<br>(.00,.18) | .72<br>(.68,.77) | .00<br>(.00,.21) | .15<br>(.00,.20) | .85<br>(.76,.90) | .25<br>(.11,.38) | .08<br>(.00,.19) | .67<br>(.62,.72) | .15<br>(.01,.29) | .13<br>(.01,.24) | .72<br>(.68,.77) |
| Marriage worries | .37<br>(.26,.42) | .01<br>(.00,.10) | .62<br>(.58,.66) | .36<br>(.22,.42) | .00<br>(.00,.10) | .64<br>(.58,.71) | .31<br>(.17,.43) | .07<br>(.00,.18) | .61<br>(.57,.67) | .32<br>(.19,.42) | .06<br>(.00,.17) | .62<br>(.58,.66) |
| Quality of relationship with twin | .24<br>(.19,.30) | .52<br>(.47,.56) | .24<br>(.22,.26) | .20<br>(.11,.29) | .55<br>(.48,.62) | .24<br>(.21,.28) | .26<br>(.19,.34) | .49<br>(.43,.55) | .24<br>(.22,.27) | .18<br>(.12,.25) | .55<br>(.49,.61) | .27<br>(.25,.29) |
| Quality of relationship with mother | .53<br>(.43,.57) | .00<br>(.00,.08) | .46<br>(.43,.50) | .47<br>(.37,.52) | .00<br>(.00,.06) | .53<br>(.48,.60) | .48<br>(.36,.59) | .08<br>(.00,.18) | .44<br>(.40,.48) | .48<br>(.36,.57) | .06<br>(.00,.16) | .46<br>(.43,.49) |
| Quality of relationship with father | .35<br>(.27,.44) | .26<br>(.19,.33) | .39<br>(.36,.42) | .38<br>(.22,.53) | .19<br>(.07,.31) | .43<br>(.38,.49) | .33<br>(.23,.43) | .30<br>(.21,.39) | .37<br>(.33,.40) | .33<br>(.23,.44) | .28<br>(.19,.37) | .38<br>(.35,.41) |
| Number of relationships | .45<br>(.40,.49) | .00<br>(.00,.00) | .55<br>(.51,.58) | .46<br>(.38,.52) | .00<br>(.00,.05) | .54<br>(.48,.60) | .44<br>(.35,.48) | .00<br>(.00,.00) | .56<br>(.52,.60) | .44<br>(.32,.50) | .03<br>(.00,.13) | .53<br>(.50,.57) |
| Longest relationship | .28<br>(.23,.33) | .00<br>(.00,.00) | .72<br>(.67,.76) | .21<br>(.09,.29) | .00<br>(.00,.00) | .79<br>(.71,.87) | .32<br>(.25,.38) | .00<br>(.00,.00) | .68<br>(.62,.73) | .32<br>(.22,.36) | .00<br>(.00,.08) | .68<br>(.64,.73) |
| Partner violence | .25<br>(.12,.29) | 0<br>(.00,.10) | .75<br>(.71,.80) | .27<br>(.05,.35) | .00<br>(.00,.00) | .73<br>(.65,.82) | .24<br>(.09,.30) | .00<br>(.00,.12) | .76<br>(.70,.81) | .25<br>(.12,.29) | .00<br>(.00,.10) | .75<br>(.71,.80) |
| Contact with mother | .47<br>(.28,.62) | .10<br>(.00,.25) | .44<br>(.38,.50) | .44<br>(.09,.63) | .09<br>(.00,.35) | .47<br>(.37,.61) | .48<br>(.25,.64) | .10<br>(.00,.29) | .42<br>(.36,.49) | .38<br>(.17,.60) | .19<br>(.00,.37) | .43<br>(.38,.50) |
| Communication with mother | .37<br>(.21,.54) | .26<br>(.10,.39) | .37<br>(.33,.43) | .37<br>(.08,.66) | .24<br>(.00,.46) | .39<br>(.31,.50) | .35<br>(.15,.57) | .28<br>(.08,.45) | .37<br>(.31,.43) | .41<br>(.21,.63) | .22<br>(.01,.40) | .37<br>(.32,.42) |
| Contact with father | .13<br>(.00,.30) | .38<br>(.24,.50) | .49<br>(.43,.55) | .03<br>(.00,.33) | .41<br>(.16,.50) | .56<br>(.46,.65) | .19<br>(.00,.39) | .36<br>(.19,.52) | .45<br>(.39,.52) | .10<br>(.00,.29) | .43<br>(.25,.54) | .48<br>(.42,.54) |
| Communication with father | .12<br>(.02,.21) | .61<br>(.53,.68) | .28<br>(.25,.31) | .18<br>(.00,.38) | .48<br>(.32,.63) | .33<br>(.27,.41) | .07<br>(.00,.18) | .68<br>(.59,.75) | .25<br>(.22,.29) | .09<br>(.00,.20) | .64<br>(.54,.73) | .27<br>(.24,.31) |
| Peer pressure | .35<br>(.27,.39) | .00<br>(.00,.05) | .65<br>(.61,.69) | .40<br>(.27,.46) | .00<br>(.00,.09) | .60<br>(.54,.67) | .31<br>(.21,.36) | .00<br>(.00,.08) | .69<br>(.64,.74) | .35<br>(.24,.39) | .00<br>(.00,.09) | .65<br>(.61,.69) |
| Physical peer victimisation | .17<br>(.11,.22) | .00<br>(.00,.00) | .83<br>(.78,.88) | .19<br>(.06,.26) | .00<br>(.00,.00) | .81<br>(.74,.88) | .09<br>(.00,.17) | .00<br>(.00,.07) | .91<br>(.83,.99) | .19<br>(.10,.25) | .00<br>(.00,.00) | .81<br>(.75,.86) |
| Social peer victimisation | .27<br>(.14,.32) | .00<br>(.00,.09) | .73<br>(.68,.78) | .00<br>(.00,.23) | .17<br>(.02,.22) | .83<br>(.73,.88) | .29<br>(.20,.34) | .00<br>(.00,.06) | .71<br>(.66,.77) | .20<br>(.05,.32) | .07<br>(.00,.19) | .73<br>(.68,.79) |
| Verbal peer victimisation | .30<br>(.21,.34) | .00<br>(.00,.06) | .70<br>(.66,.75) | .18<br>(.00,.33) | .07<br>(.00,.22) | .76<br>(.67,.85) | .32<br>(.24,.37) | .00<br>(.00,.00) | .68<br>(.63,.74) | .30<br>(.14,.34) | .00<br>(.00,.13) | .70<br>(.66,.75) |
| Cyber peer victimisation | .33<br>(.26,.37) | .00<br>(.00,.04) | .67<br>(.63,.72) | .06<br>(.00,.29) | .18<br>(.02,.27) | .75<br>(.66,.83) | .34<br>(.29,.40) | .00<br>(.00,.03) | .66<br>(.60,.71) | .32<br>(.25,.37) | .00<br>(.00,.05) | .68<br>(.63,.73) |
| Physical peer Perpetration | .12<br>(.05,.17) | .00<br>(.00,.00) | .88<br>(.83,.93) | .13<br>(.00,.20) | .00<br>(.00,.13) | .87<br>(.80,.95) | .05<br>(.00,.12) | .00<br>(.00,.05) | .95<br>(.88,1.00) | .13<br>(.00,.21) | .02<br>(.00,.16) | .85<br>(.79,.91) |
| Social peer Perpetration | .15<br>(.06,.2) | .00<br>(.00,.06) | .85<br>(.80,.91) | .15<br>(.01,.25) | .00<br>(.00,.00) | .85<br>(.75,.94) | .14<br>(.00,.20) | .00<br>(.00,.11) | .86<br>(.80,.92) | .14<br>(.00,.22) | .02<br>(.00,.15) | .84<br>(.78,.90) |
| Verbal peer Perpetration | .31<br>(.26,.36) | .00<br>(.00,.03) | .69<br>(.64,.73) | .32<br>(.19,.38) | .00<br>(.00,.00) | .68<br>(.62,.75) | .29<br>(.21,.35) | .00<br>(.00,.05) | .71<br>(.65,.77) | .32<br>(.21,.36) | .00<br>(.00,.09) | .68<br>(.64,.73) |
| Cyber peer Perpetration | .19<br>(.13,.24) | 0<br>(.00,.03) | .81<br>(.76,.87) | .14<br>(.02,.23) | .00<br>(.00,.00) | .86<br>(.77,.95) | .21<br>(.13,.27) | .00<br>(.00,.06) | .79<br>(.73,.85) | .20<br>(.10,.25) | .00<br>(.00,.08) | .80<br>(.75,.85) |
| Parental role aspirations | .34<br>(.26,.38) | .00<br>(.00,.06) | .66<br>(.62,.70) | .25<br>(.14,.32) | .00<br>(.00,.00) | .75<br>(.68,.83) | .35<br>(.20,.44) | .03<br>(.00,.15) | .61<br>(.56,.67) | .35<br>(.26,.39) | .00<br>(.00,.00) | .65<br>(.61,.70) |
| Occupational role aspirations | .26<br>(.18,.3) | .00<br>(.00,.06) | .74<br>(.70,.78) | .20<br>(.06,.27) | .00<br>(.00,.10) | .80<br>(.73,.87) | .29<br>(.16,.34) | .00<br>(.00,.10) | .71<br>(.66,.76) | .27<br>(.13,.32) | .00<br>(.00,.12) | .73<br>(.68,.78) |
| Homecare role aspirations | .40<br>(.33,.44) | .00<br>(.00,.05) | .60<br>(.56,.64) | .41<br>(.31,.47) | .00<br>(.00,.07) | .59<br>(.53,.65) | .39<br>(.27,.43) | .00<br>(.00,.09) | .61<br>(.57,.66) | .36<br>(.22,.45) | .05<br>(.00,.17) | .59<br>(.55,.63) |
| Importance of Relationships | .37<br>(.33,.4) | .00<br>(.00,.02) | .63<br>(.60,.67) | .38<br>(.32,.44) | .00<br>(.00,.04) | .62<br>(.56,.68) | .34<br>(.29,.39) | .00<br>(.00,.03) | .66<br>(.61,.71) | .39<br>(.34,.43) | .00<br>(.00,.03) | .61<br>(.57,.65) |
| Achievement Motivation | .37<br>(.28,.4) | .00<br>(.00,.06) | .63<br>(.60,.67) | .31<br>(.12,.41) | .03<br>(.00,.17) | .66<br>(.59,.74) | .38<br>(.29,.42) | .00<br>(.00,.06) | .62<br>(.58,.67) | .35<br>(.22,.41) | .01<br>(.00,.13) | .63<br>(.59,.68) |
| Purpose in Life | .42<br>(.31,.46) | .01<br>(.00,.09) | .57<br>(.54,.61) | .39<br>(.21,.49) | .04<br>(.00,.17) | .57<br>(.51,.64) | .42<br>(.30,.47) | .00<br>(.00,.00) | .58<br>(.53,.63) | .37<br>(.25,.46) | .05<br>(.00,.16) | .57<br>(.54,.62) |
| Importance of democracy and equality | .43<br>(.38,.47) | .00<br>(.00,.03) | .57<br>(.53,.60) | .42<br>(.24,.49) | .02<br>(.00,.15) | .56<br>(.51,.63) | .43<br>(.37,.47) | .00<br>(.00,.00) | .57<br>(.53,.62) | .44<br>(.37,.47) | .00<br>(.00,.05) | .56<br>(.53,.60) |
| Environmental concerns | .53<br>(.43,.56) | .00<br>(.00,.08) | .47<br>(.44,.51) | .54<br>(.41,.59) | .00<br>(.00,.00) | .46<br>(.41,.52) | .48<br>(.35,.55) | .03<br>(.00,.13) | .49<br>(.45,.54) | .47<br>(.35,.55) | .06<br>(.00,.15) | .48<br>(.45,.51) |
| Religiosity | .25<br>(.19,.32) | .44<br>(.39,.49) | .31<br>(.28,.33) | .22<br>(.12,.32) | .48<br>(.39,.55) | .30<br>(.26,.34) | .28<br>(.20,.36) | .41<br>(.34,.48) | .31<br>(.28,.34) | .22<br>(.14,.30) | .48<br>(.41,.55) | .30<br>(.28,.32) |
| Importance of leisure | .41<br>(.31,.45) | .00<br>(.00,.08) | .59<br>(.55,.63) | .38<br>(.17,.45) | .01<br>(.00,.16) | .61<br>(.55,.70) | .42<br>(.30,.47) | .00<br>(.00,.00) | .58<br>(.53,.63) | .40<br>(.26,.46) | .02<br>(.00,.14) | .58<br>(.54,.63) |
| Alcohol use | .46<br>(.34,.54) | .03<br>(.00,.13) | .51<br>(.46,.55) | .29<br>(.07,.49) | .13<br>(.00,.29) | .58<br>(.50,.66) | .53<br>(.40,.57) | .00<br>(.00,.00) | .47<br>(.43,.52) | .40<br>(.26,.53) | .09<br>(.00,.21) | .51<br>(.47,.56) |
| Ever smoked | .48<br>(.41,.52) | .00<br>(.00,.06) | .52<br>(.48,.55) | .45<br>(.34,.51) | .00<br>(.00,.08) | .55<br>(.49,.61) | .50<br>(.36,.54) | .00<br>(.00,.11) | .50<br>(.46,.55) | .49<br>(.38,.52) | .00<br>(.00,.00) | .51<br>(.48,.55) |
| Ever vaped | .33<br>(.21,.42) | .04<br>(.00,.14) | .62<br>(.58,.67) | .35<br>(.14,.44) | .01<br>(.00,.17) | .63<br>(.56,.72) | .30<br>(.15,.43) | .08<br>(.00,.20) | .62<br>(.57,.68) | .28<br>(.13,.41) | .09<br>(.00,.21) | .63<br>(.58,.68) |
| Cognitive enhancers | .36<br>(.23,.42) | .01<br>(.00,.11) | .63<br>(.58,.68) | .34<br>(.14,.41) | .00<br>(.00,.14) | .66<br>(.59,.75) | .32<br>(.16,.43) | .06<br>(.00,.18) | .62<br>(.57,.68) | .26<br>(.10,.40) | .10<br>(.00,.23) | .64<br>(.59,.69) |
| Big 5 Agreeableness | .22<br>(.18,.27) | .00<br>(.00,.00) | .78<br>(.73,.82) | .30<br>(.19,.37) | .00<br>(.00,.00) | .70<br>(.63,.77) | .17<br>(.11,.23) | .00<br>(.00,.02) | .83<br>(.77,.89) | .24<br>(.19,.29) | .00<br>(.00,.03) | .76<br>(.71,.80) |
| Big 5 Conscientiousness | .30<br>(.26,.34) | .00<br>(.00,.02) | .70<br>(.66,.74) | .33<br>(.24,.39) | .00<br>(.00,.04) | .67<br>(.61,.74) | .28<br>(.22,.34) | .00<br>(.00,.00) | .72<br>(.66,.77) | .33<br>(.28,.37) | .00<br>(.00,.03) | .67<br>(.63,.72) |
| Big 5 Extraversion | .40<br>(.36,.44) | .00<br>(.00,.02) | .60<br>(.56,.64) | .41<br>(.33,.47) | .00<br>(.00,.05) | .59<br>(.53,.65) | .39<br>(.34,.44) | .00<br>(.00,.03) | .61<br>(.56,.66) | .42<br>(.37,.46) | .00<br>(.00,.00) | .58<br>(.54,.62) |
| Big 5 Neuroticism | .37<br>(.26,.41) | .00<br>(.00,.09) | .63<br>(.59,.67) | .25<br>(.04,.39) | .07<br>(.00,.21) | .68<br>(.61,.77) | .39<br>(.29,.44) | .00<br>(.00,.08) | .61<br>(.56,.66) | .35<br>(.22,.42) | .03<br>(.00,.14) | .62<br>(.58,.67) |
| Big 5 Openness | .36<br>(.32,.4) | 0<br>(.00,.02) | .64<br>(.60,.68) | .37<br>(.28,.43) | .00<br>(.00,.06) | .63<br>(.57,.70) | .36<br>(.30,.41) | .00<br>(.00,.00) | .64<br>(.59,.69) | .38<br>(.33,.42) | .00<br>(.00,.04) | .62<br>(.58,.66) |
| Self-control | .38<br>(.34,.42) | .00<br>(.00,.03) | .62<br>(.58,.66) | .35<br>(.24,.41) | .00<br>(.00,.07) | .65<br>(.59,.72) | .40<br>(.34,.45) | .00<br>(.00,.04) | .60<br>(.55,.65) | .39<br>(.34,.44) | .00<br>(.00,.04) | .61<br>(.56,.65) |
| Not planning for the future | .26<br>(.13,.3) | .00<br>(.00,.09) | .74<br>(.70,.79) | .29<br>(.16,.35) | .00<br>(.00,.08) | .71<br>(.65,.78) | .17<br>(.01,.29) | .06<br>(.00,.18) | .77<br>(.71,.84) | .25<br>(.13,.30) | .00<br>(.00,.00) | .75<br>(.70,.80) |
| Ambition | .36<br>(.29,.4) | .00<br>(.00,.05) | .64<br>(.60,.68) | .34<br>(.22,.41) | .00<br>(.00,.00) | .66<br>(.59,.73) | .37<br>(.26,.41) | .00<br>(.00,.08) | .63<br>(.59,.68) | .37<br>(.27,.41) | .00<br>(.00,.08) | .63<br>(.59,.68) |
| Physical activity | .39<br>(.33,.43) | .00<br>(.00,.05) | .61<br>(.57,.64) | .39<br>(.26,.45) | .00<br>(.00,.09) | .61<br>(.55,.68) | .40<br>(.30,.44) | .00<br>(.00,.08) | .60<br>(.56,.65) | .40<br>(.28,.44) | .00<br>(.00,.10) | .60<br>(.56,.64) |
| Health behaviours | .52<br>(.42,.58) | .03<br>(.00,.11) | .45<br>(.42,.48) | .55<br>(.42,.60) | .00<br>(.00,.10) | .45<br>(.40,.51) | .47<br>(.35,.58) | .08<br>(.00,.18) | .45<br>(.41,.49) | .48<br>(.37,.58) | .07<br>(.00,.17) | .45<br>(.42,.48) |
| Risk taking behaviour | .50<br>(.42,.53) | .00<br>(.00,.06) | .50<br>(.47,.54) | .47<br>(.30,.56) | .04<br>(.00,.17) | .49<br>(.44,.55) | .47<br>(.40,.52) | .00<br>(.00,.06) | .53<br>(.48,.57) | .50<br>(.39,.53) | .00<br>(.00,.09) | .50<br>(.47,.54) |
| Risky sexual behaviour | .60<br>(.52,.63) | .00<br>(.00,.07) | .40<br>(.37,.43) | .59<br>(.47,.63) | .00<br>(.00,.09) | .41<br>(.37,.47) | .61<br>(.49,.64) | .00<br>(.00,.10) | .39<br>(.36,.43) | .55<br>(.44,.63) | .06<br>(.00,.16) | .39<br>(.36,.43) |
| Media use | .51<br>(.48,.55) | .00<br>(.00,.02) | .49<br>(.45,.52) | .48<br>(.42,.53) | .00<br>(.00,.00) | .52<br>(.47,.58) | .53<br>(.47,.57) | .00<br>(.00,.04) | .47<br>(.43,.51) | .52<br>(.48,.56) | .00<br>(.00,.00) | .48<br>(.44,.51) |
| Volunteering | .44<br>(.36,.47) | .00<br>(.00,.06) | .56<br>(.53,.60) | .47<br>(.34,.53) | .00<br>(.00,.10) | .53<br>(.47,.59) | .42<br>(.32,.46) | .00<br>(.00,.00) | .58<br>(.54,.63) | .45<br>(.32,.48) | .00<br>(.00,.11) | .55<br>(.52,.59) |
| Internet dating (scale) | .34<br>(.22,.37) | .00<br>(.00,.09) | .66<br>(.63,.71) | .36<br>(.17,.42) | .00<br>(.00,.14) | .64<br>(.58,.71) | .32<br>(.19,.37) | .00<br>(.00,.11) | .68<br>(.63,.73) | .30<br>(.16,.38) | .04<br>(.00,.15) | .66<br>(.62,.71) |
| Internet dating (y/n) | .31<br>(.23,.35) | .00<br>(.00,.00) | .69<br>(.65,.73) | .33<br>(.15,.39) | .00<br>(.00,.13) | .67<br>(.61,.74) | .30<br>(.20,.35) | .00<br>(.00,.08) | .70<br>(.65,.75) | .26<br>(.12,.36) | .06<br>(.00,.18) | .68<br>(.64,.73) |
| Financial literacy: Knowledge of Products | .37<br>(.31,.41) | .00<br>(.00,.00) | .63<br>(.59,.67) | .44<br>(.30,.50) | .00<br>(.00,.11) | .56<br>(.50,.62) | .31<br>(.23,.36) | .00<br>(.00,.00) | .69<br>(.64,.75) | .38<br>(.25,.43) | .01<br>(.00,.12) | .61<br>(.57,.66) |
| Financial attitudes and behaviour | .28<br>(.23,.32) | .00<br>(.00,.00) | .72<br>(.68,.76) | .28<br>(.13,.35) | .00<br>(.00,.00) | .72<br>(.65,.79) | .28<br>(.21,.33) | .00<br>(.00,.04) | .72<br>(.67,.77) | .29<br>(.22,.34) | .00<br>(.00,.05) | .71<br>(.66,.75) |
| Functional outcome | Model fitting results |  |  | Males only |  |  | Females only |  |  | Same-sex MZ and DZ twins |  |  |
|  | A | C | E | A | C | E | A | C | E | A | C | E |
| Wellbeing factor (1st PC) | .33<br>(.21,.44) | .07<br>(.00,.16) | .60<br>(.56,.64) | .36<br>(.16,.42) | .00<br>(.00,.14) | .64<br>(.58,.72) | .28<br>(.15,.42) | .14<br>(.02,.24) | .58<br>(.53,.63) | .22<br>(.09,.35) | .18<br>(.07,.29) | .60<br>(.56,.64) |
| Adverse mental health factor (1st PC) | .50<br>(.42,.54) | .00<br>(.00,.07) | .50<br>(.46,.54) | .33<br>(.09,.49) | .07<br>(.00,.24) | .60<br>(.51,.70) | .54<br>(.46,.59) | .00<br>(.00,.07) | .46<br>(.41,.51) | .52<br>(.37,.56) | .00<br>(.00,.13) | .48<br>(.44,.53) |
| Adverse physical health factor (1st PC) | .39<br>(.27,.43) | .00<br>(.00,.10) | .61<br>(.57,.65) | .37<br>(.23,.45) | .00<br>(.00,.10) | .63<br>(.55,.70) | .36<br>(.20,.45) | .04<br>(.00,.17) | .60<br>(.55,.66) | .32<br>(.17,.44) | .08<br>(.00,.21) | .60<br>(.55,.64) |
| Achieved educational level | .27<br>(.16,.38) | .16<br>(.07,.24) | .57<br>(.53,.62) | .25<br>(.07,.43) | .15<br>(.01,.28) | .60<br>(.53,.67) | .27<br>(.14,.41) | .17<br>(.06,.27) | .56<br>(.51,.61) | .29<br>(.16,.42) | .14<br>(.03,.25) | .57<br>(.53,.61) |
| Planned educational level | .40<br>(.29,.46) | .00<br>(.00,.08) | .60<br>(.54,.66) | .41<br>(.21,.50) | .00<br>(.00,.16) | .59<br>(.50,.69) | .39<br>(.24,.47) | .00<br>(.00,.12) | .61<br>(.53,.69) | .43<br>(.25,.49) | .00<br>(.00,.16) | .57<br>(.51,.64) |

Note. MZ=monozygotic; DZ=dizygotic; and genetic (A), shared environmental (C) and non-shared environmental (E) estimates. Scores were corrected for mean age and sex differences (see Methods).

**Table S10.**
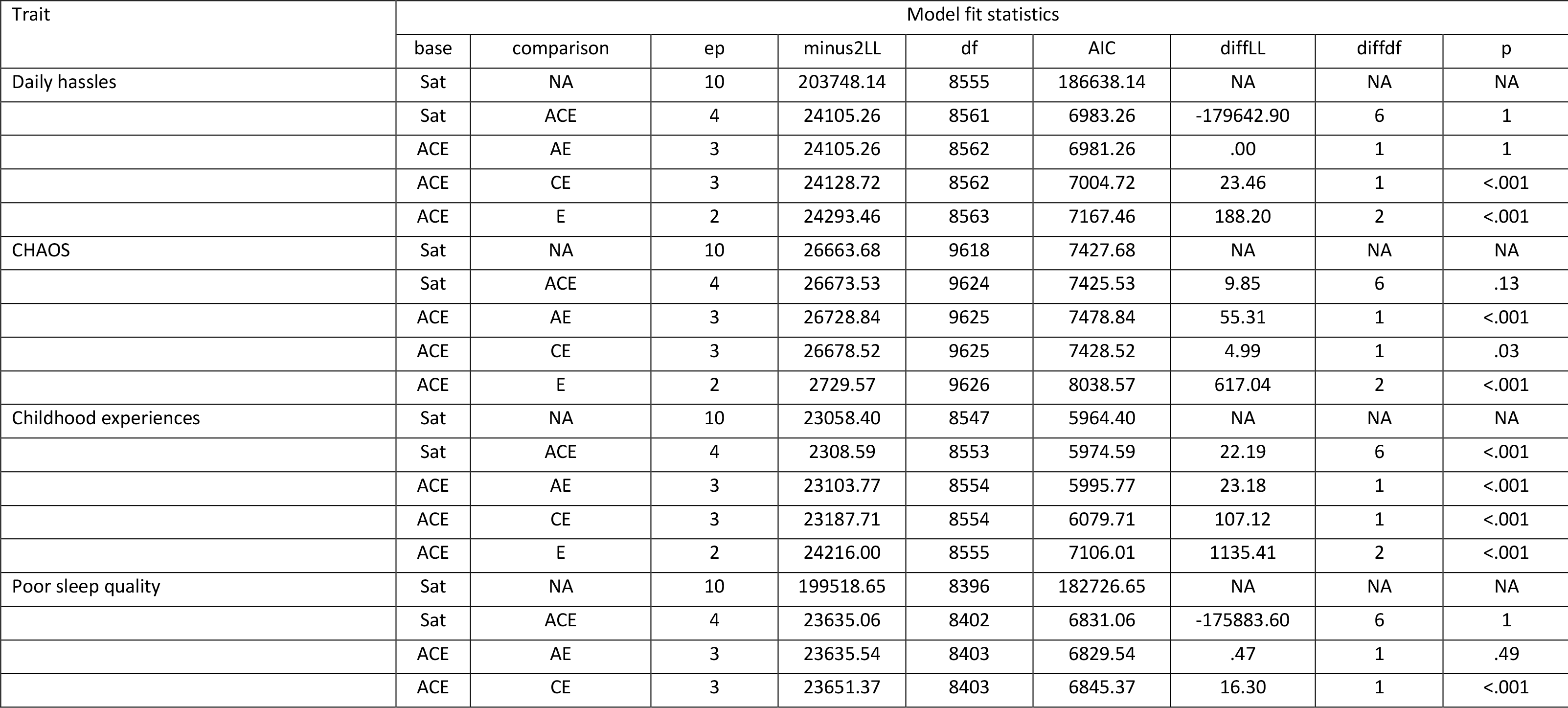
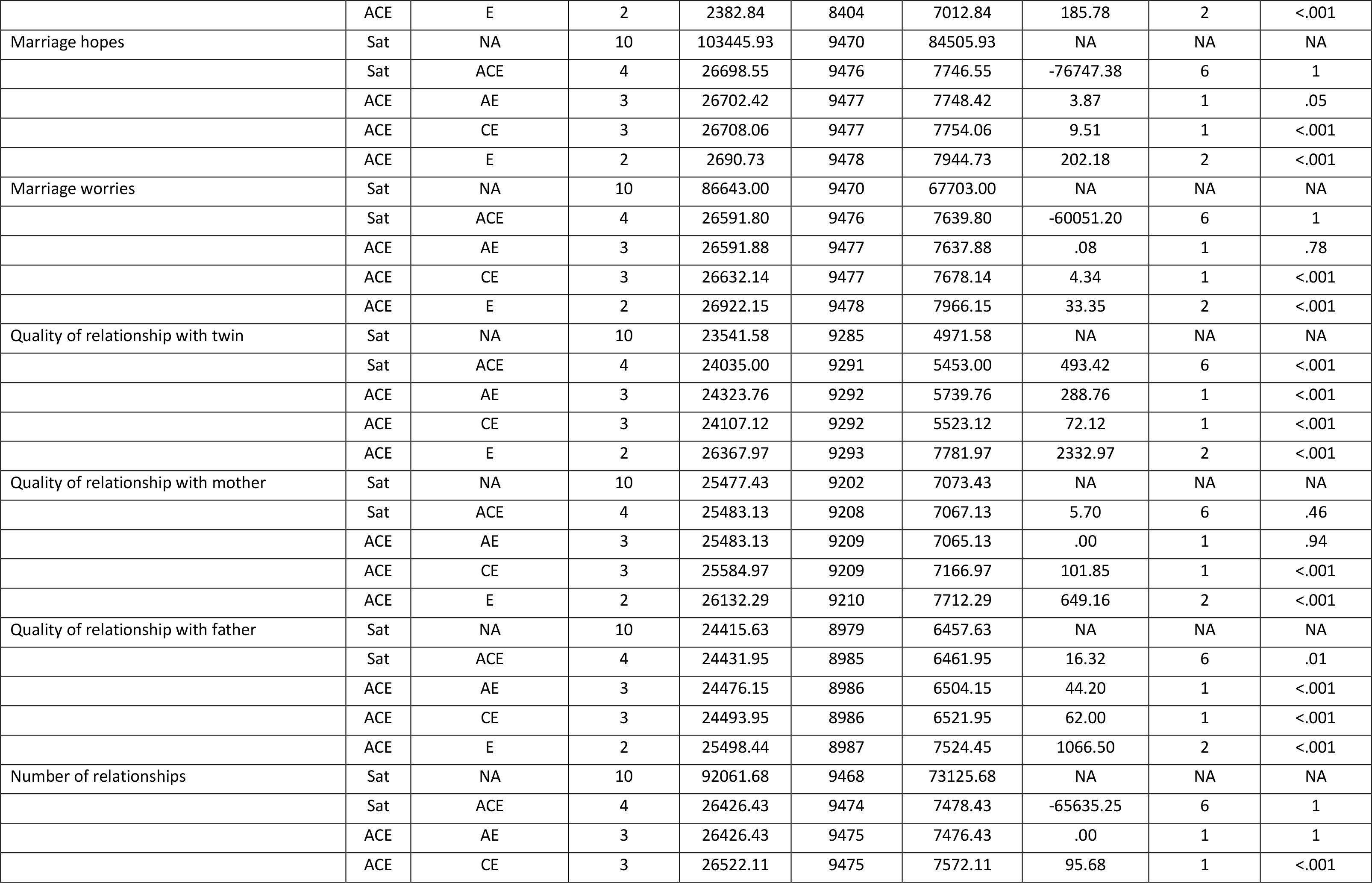
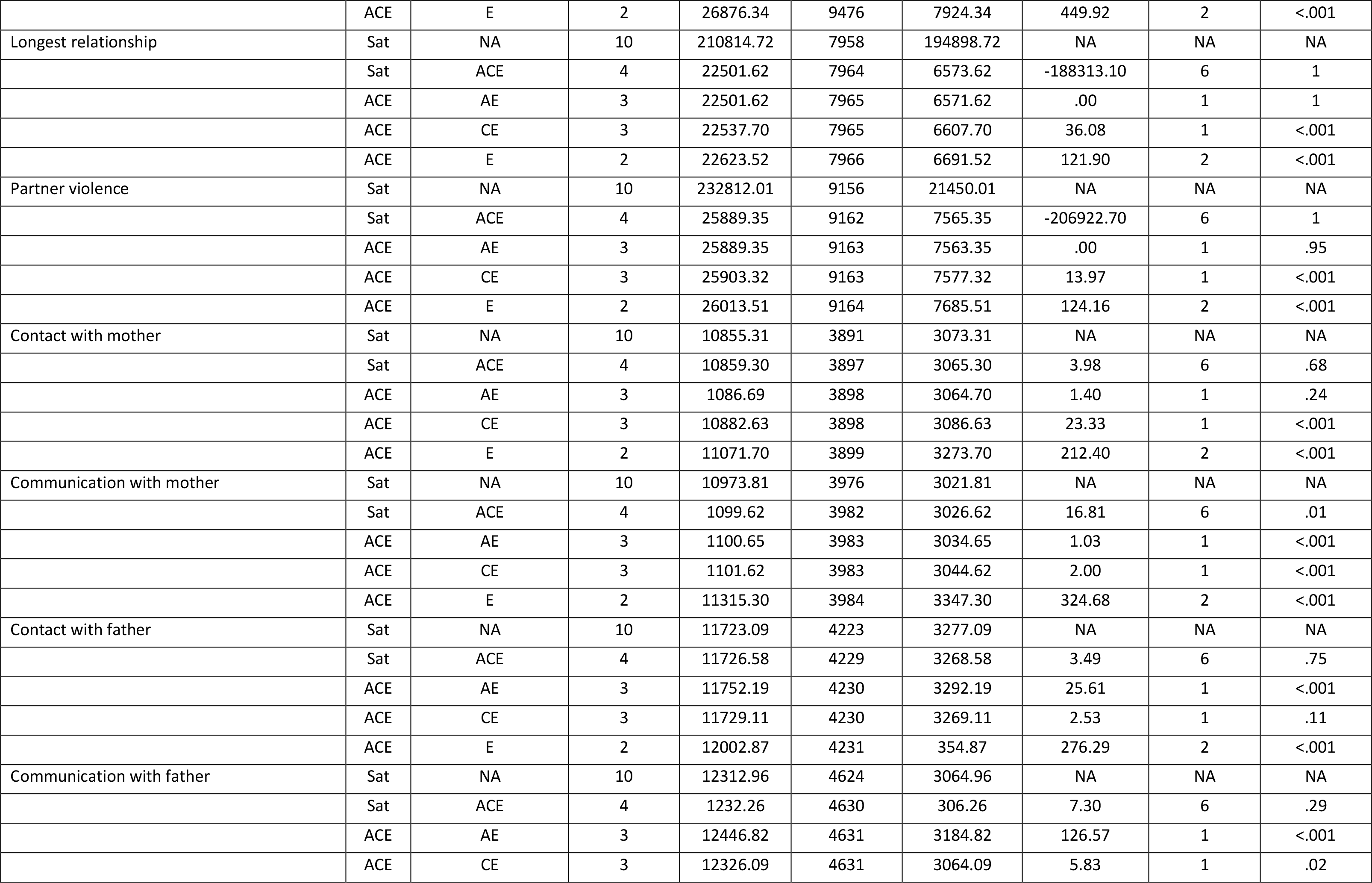
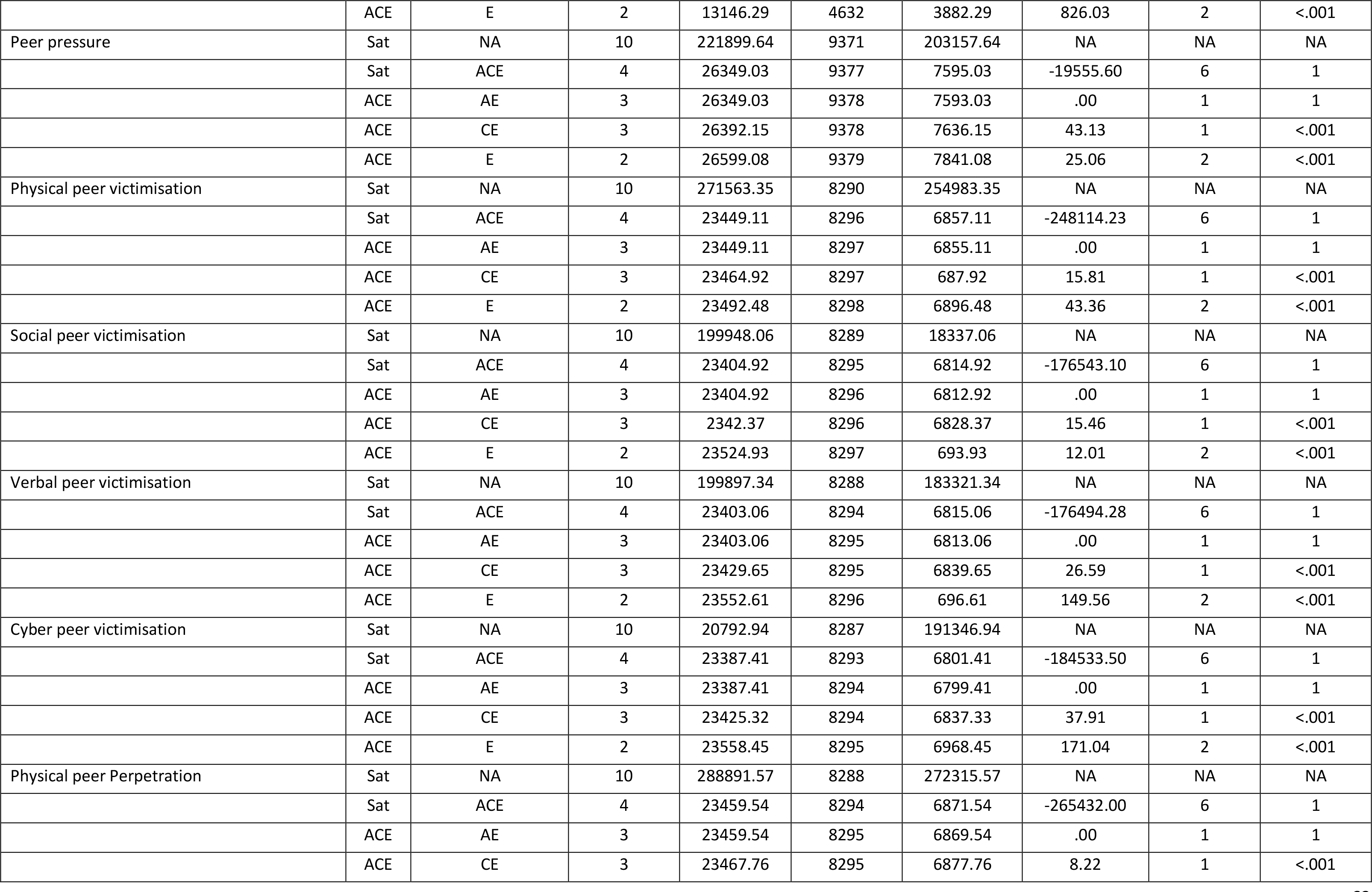
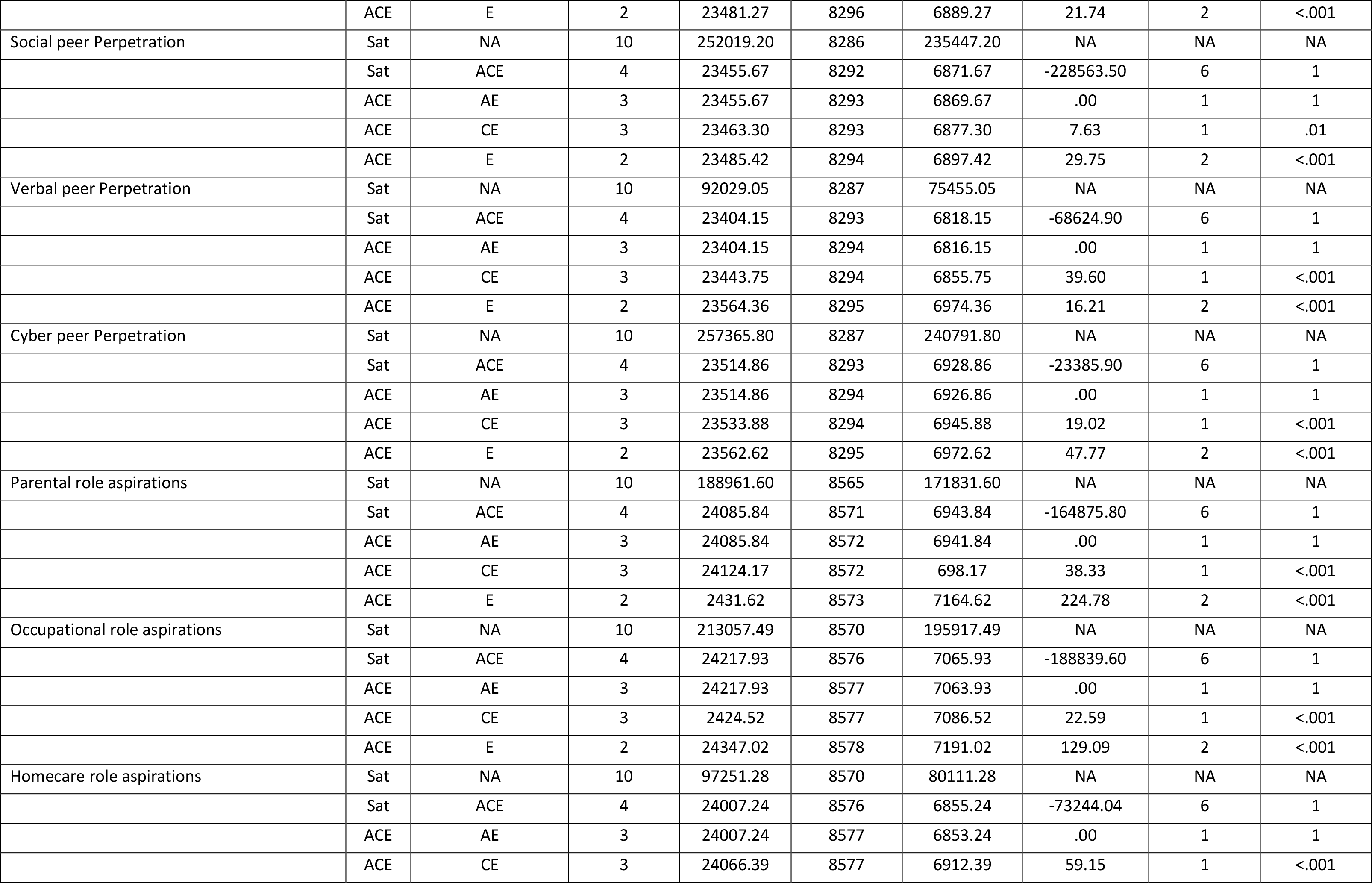
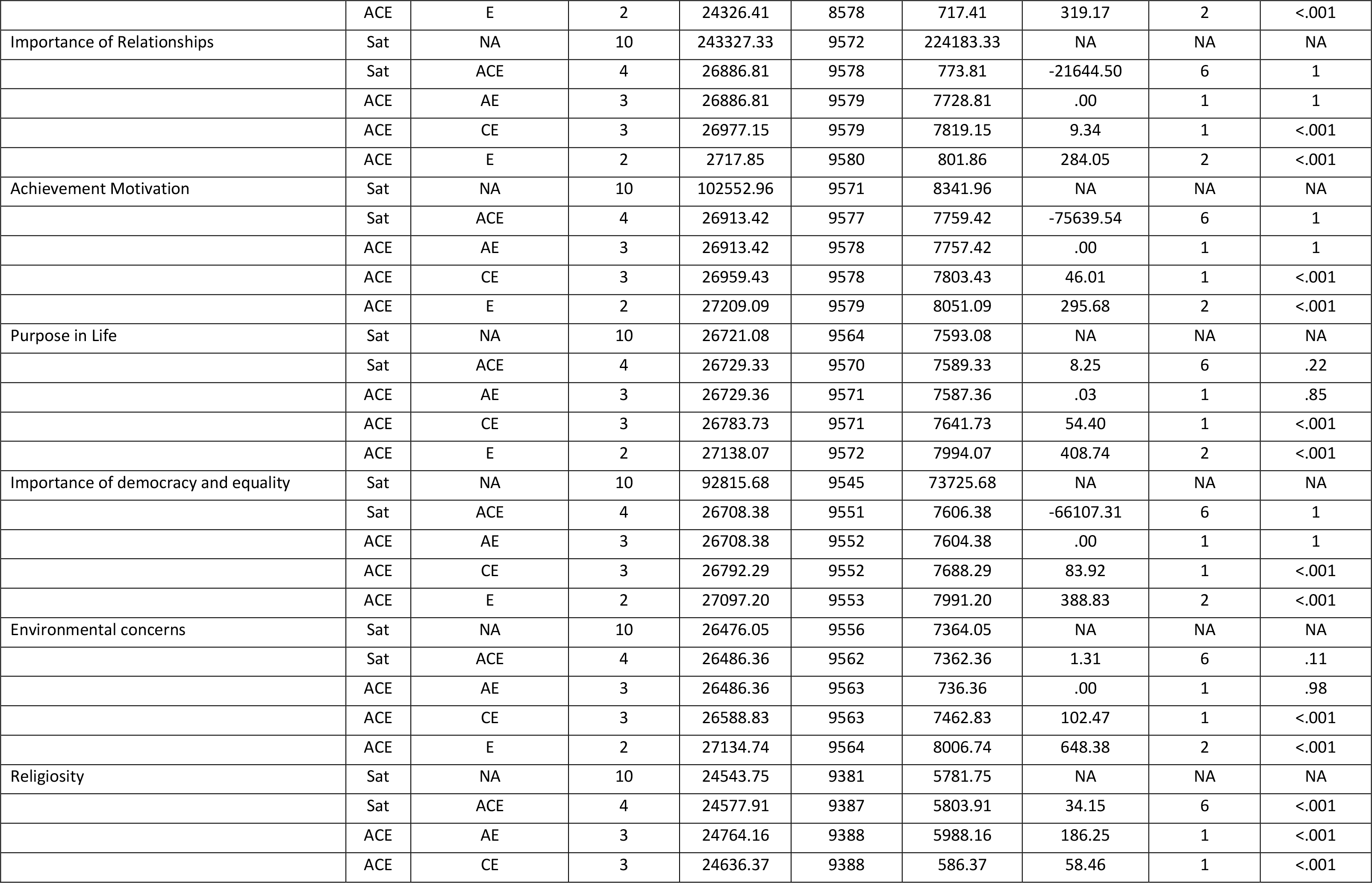
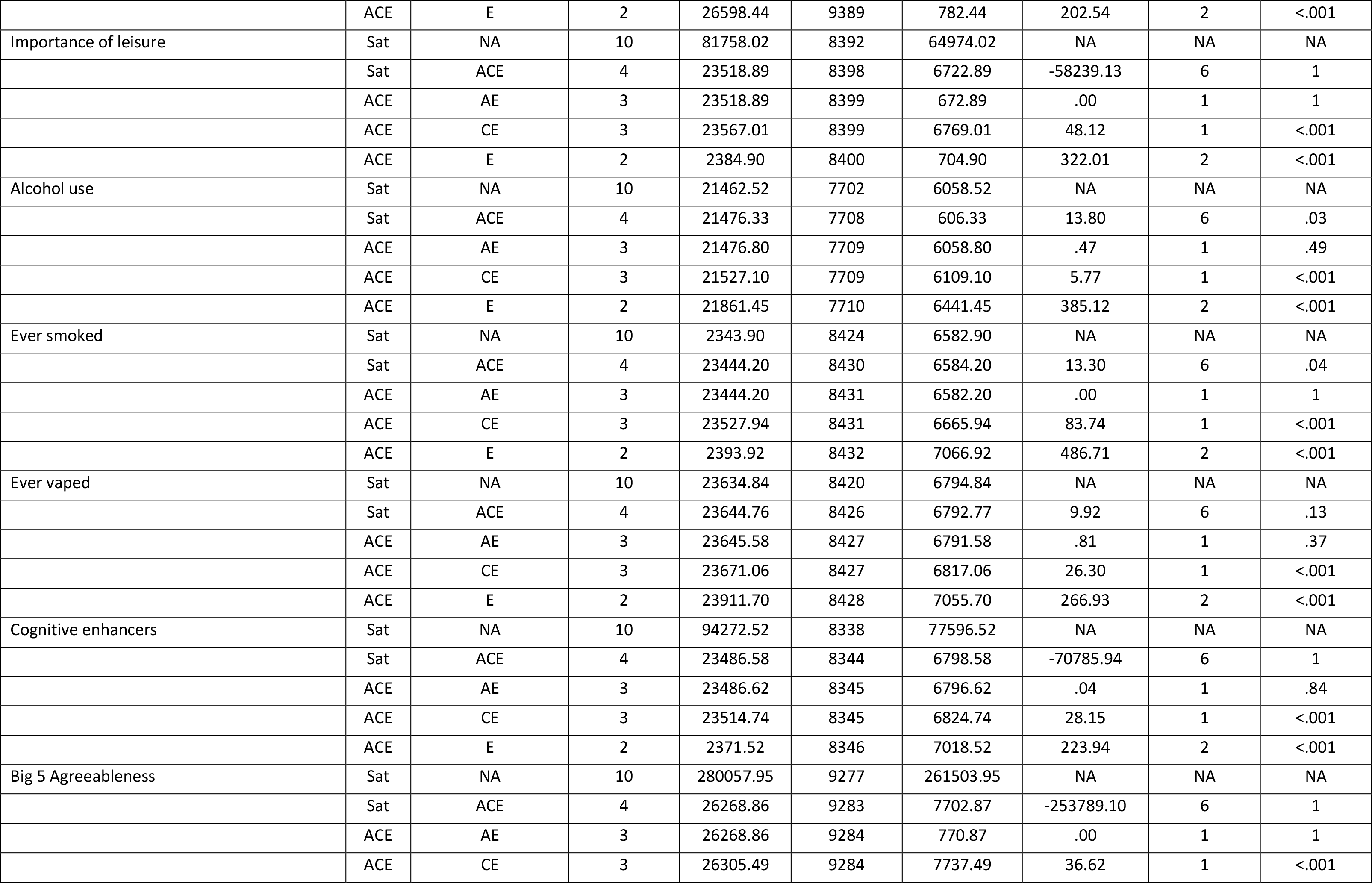
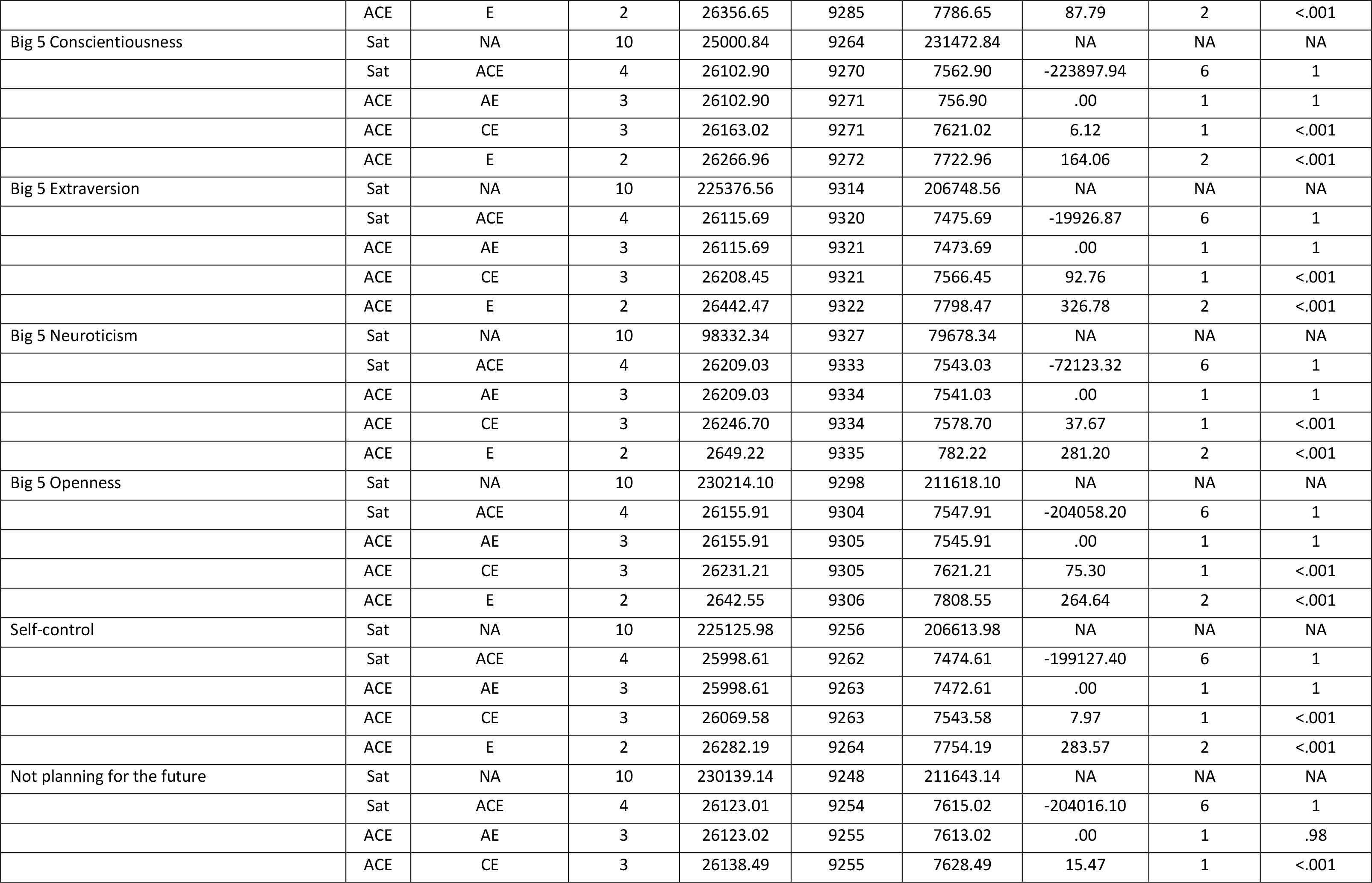
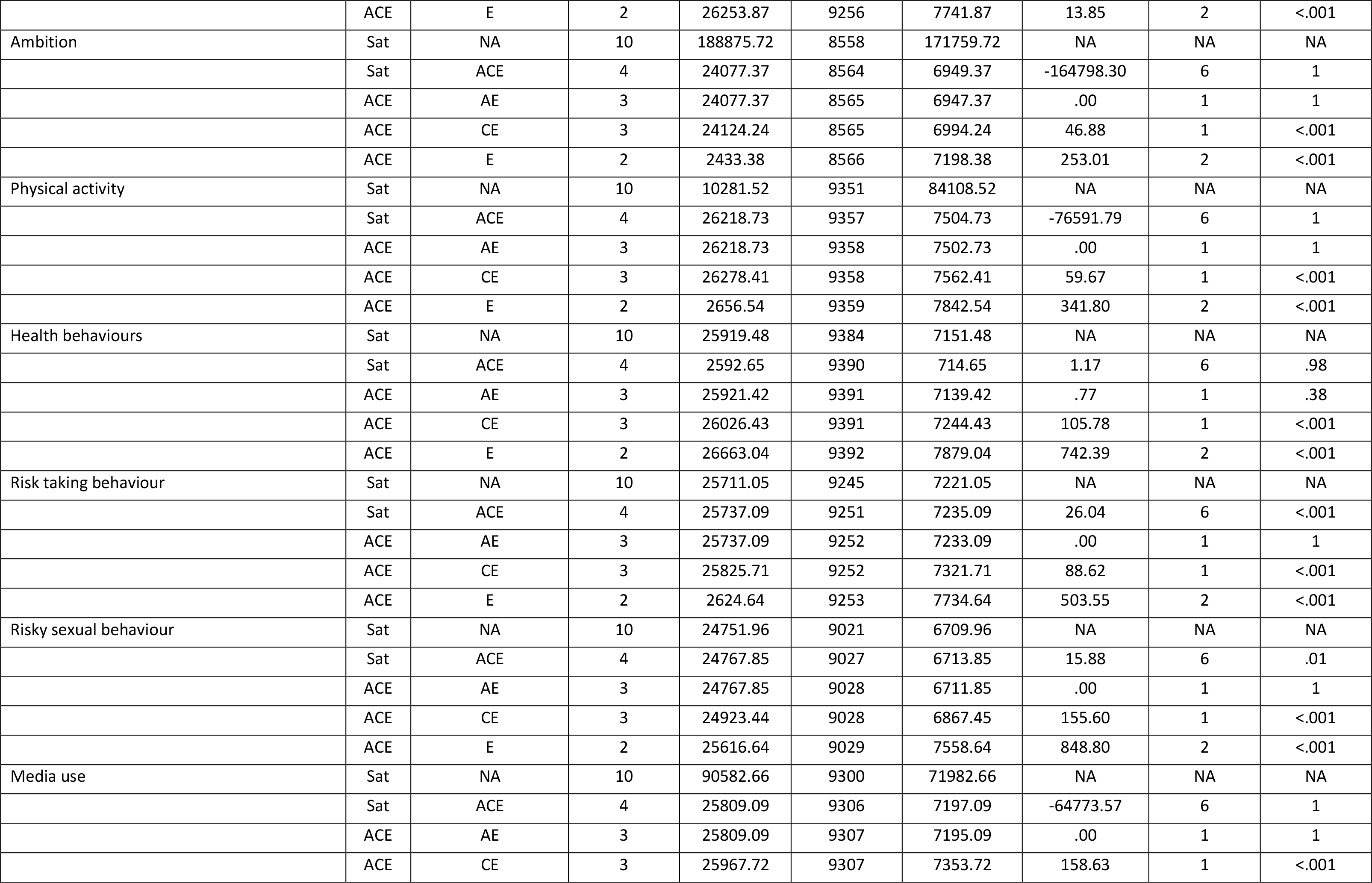
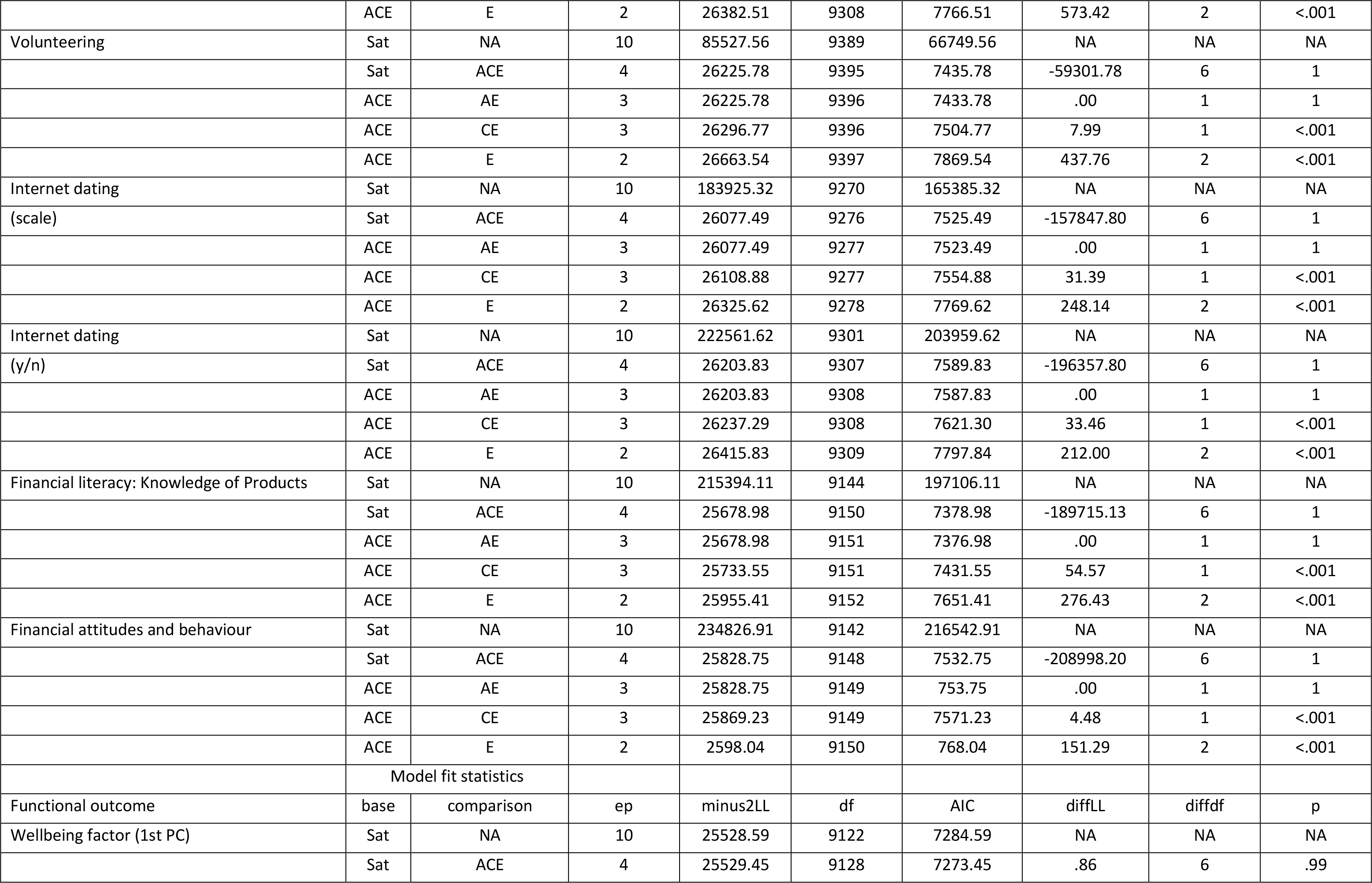
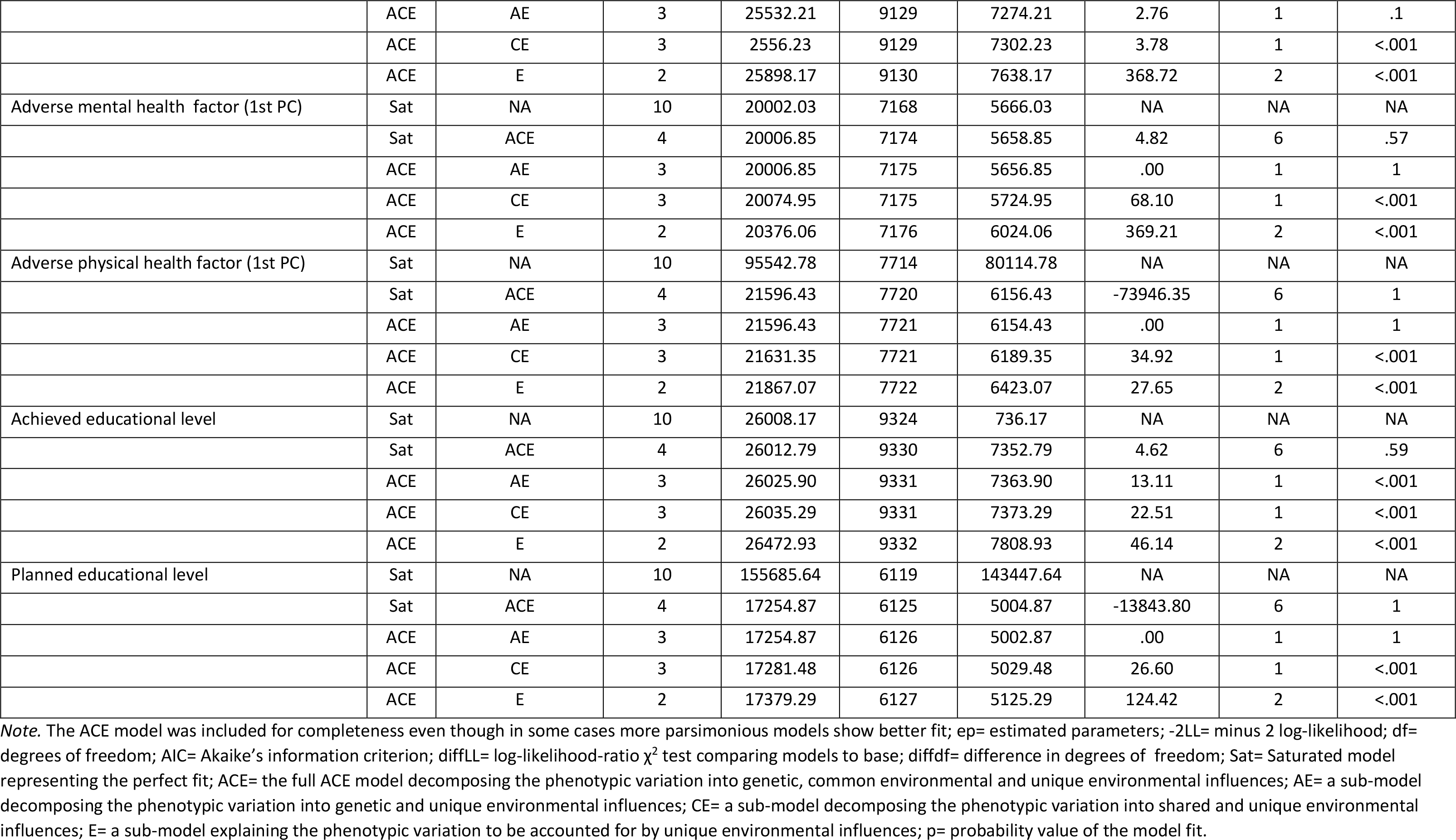
Model-fit statistics for the univariate twin analyses

**Table S11.**
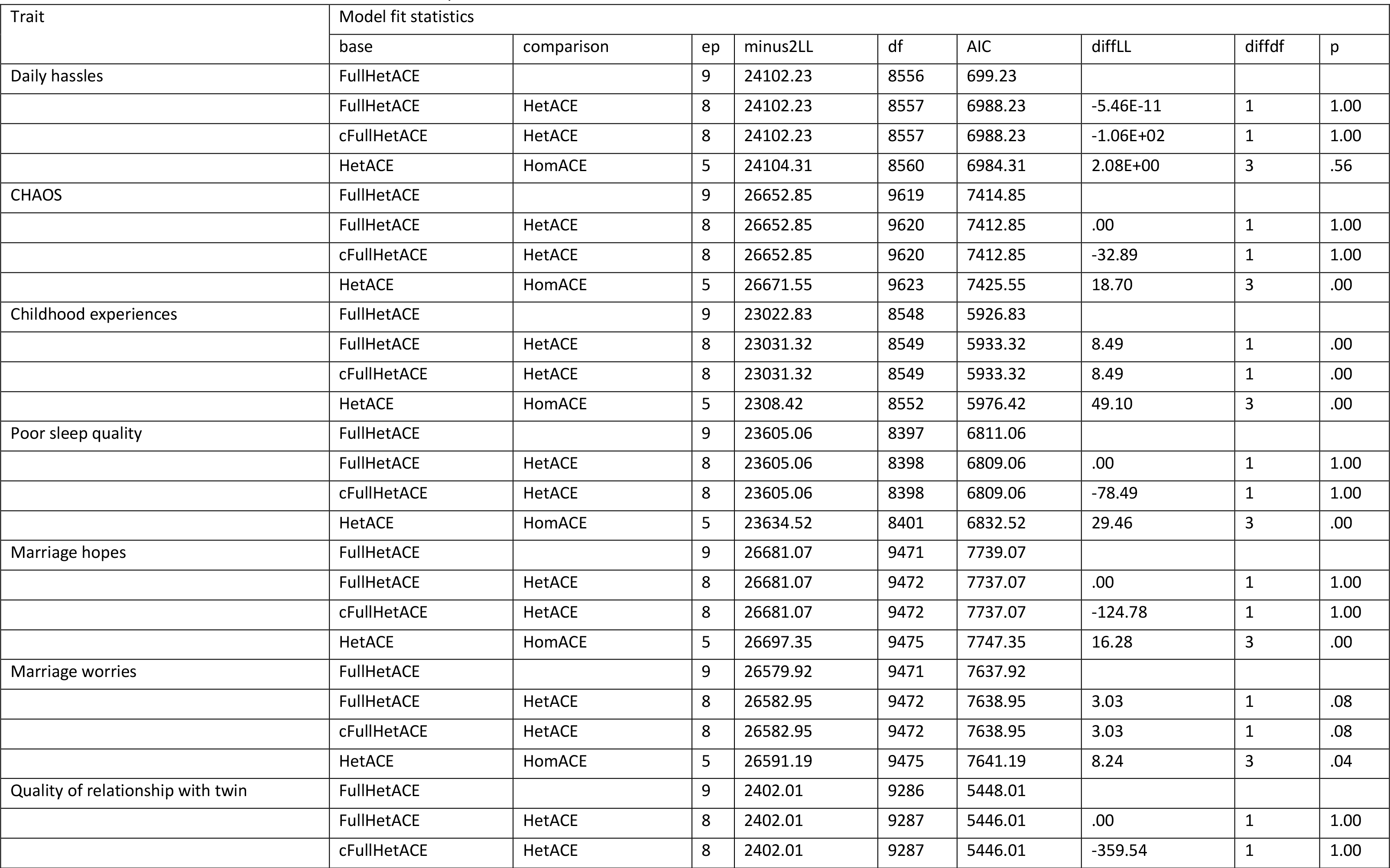
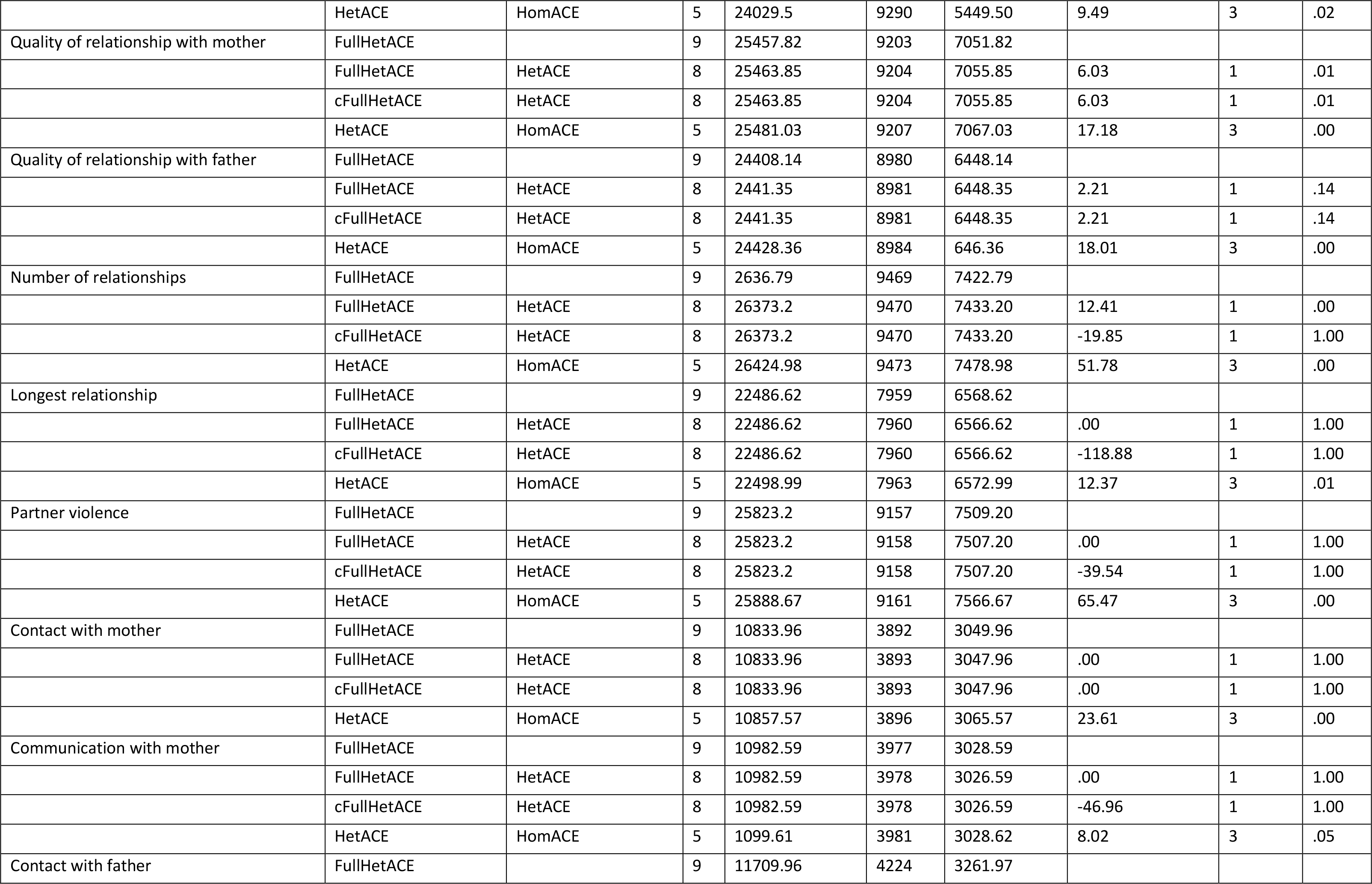
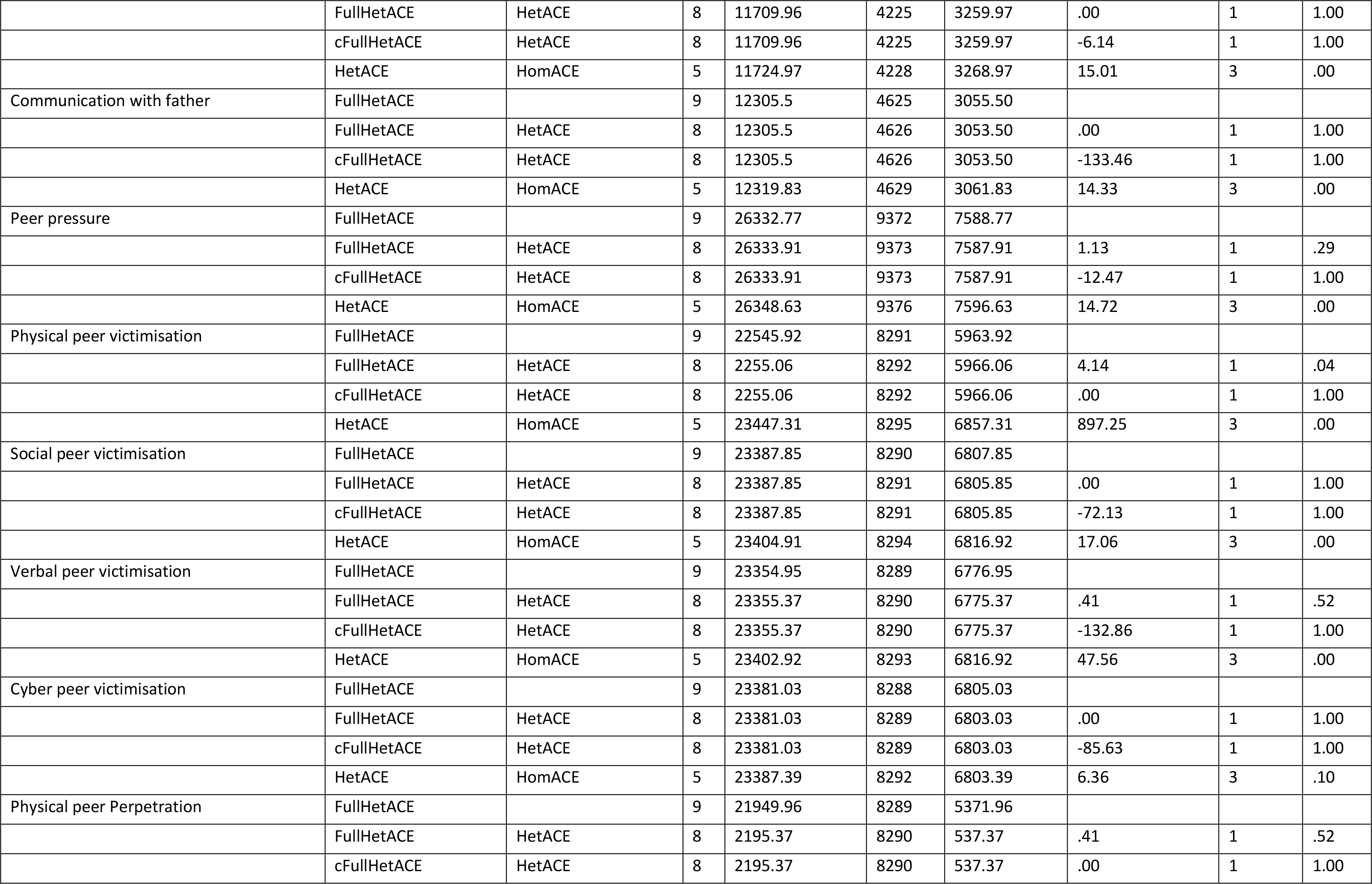
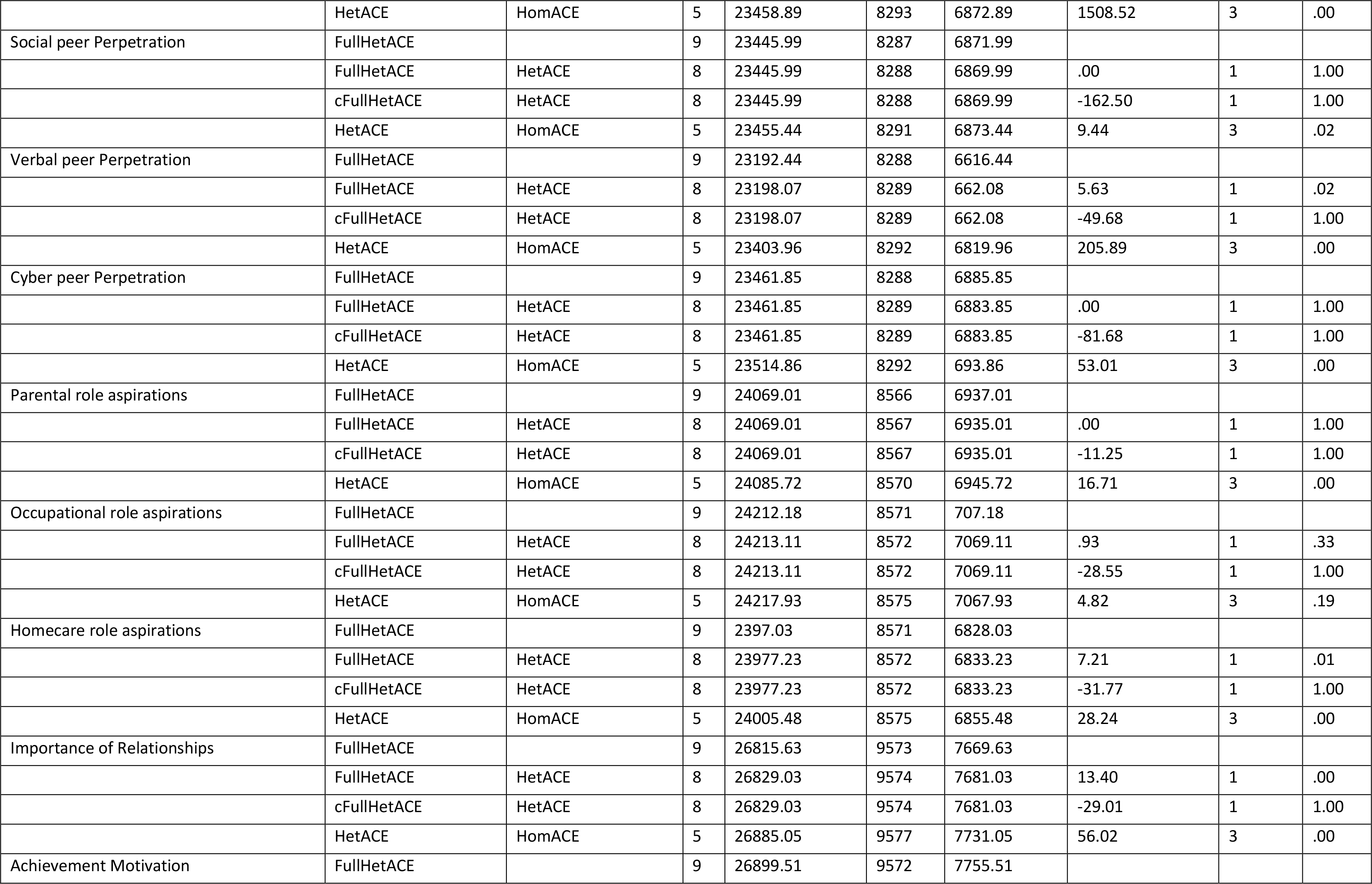
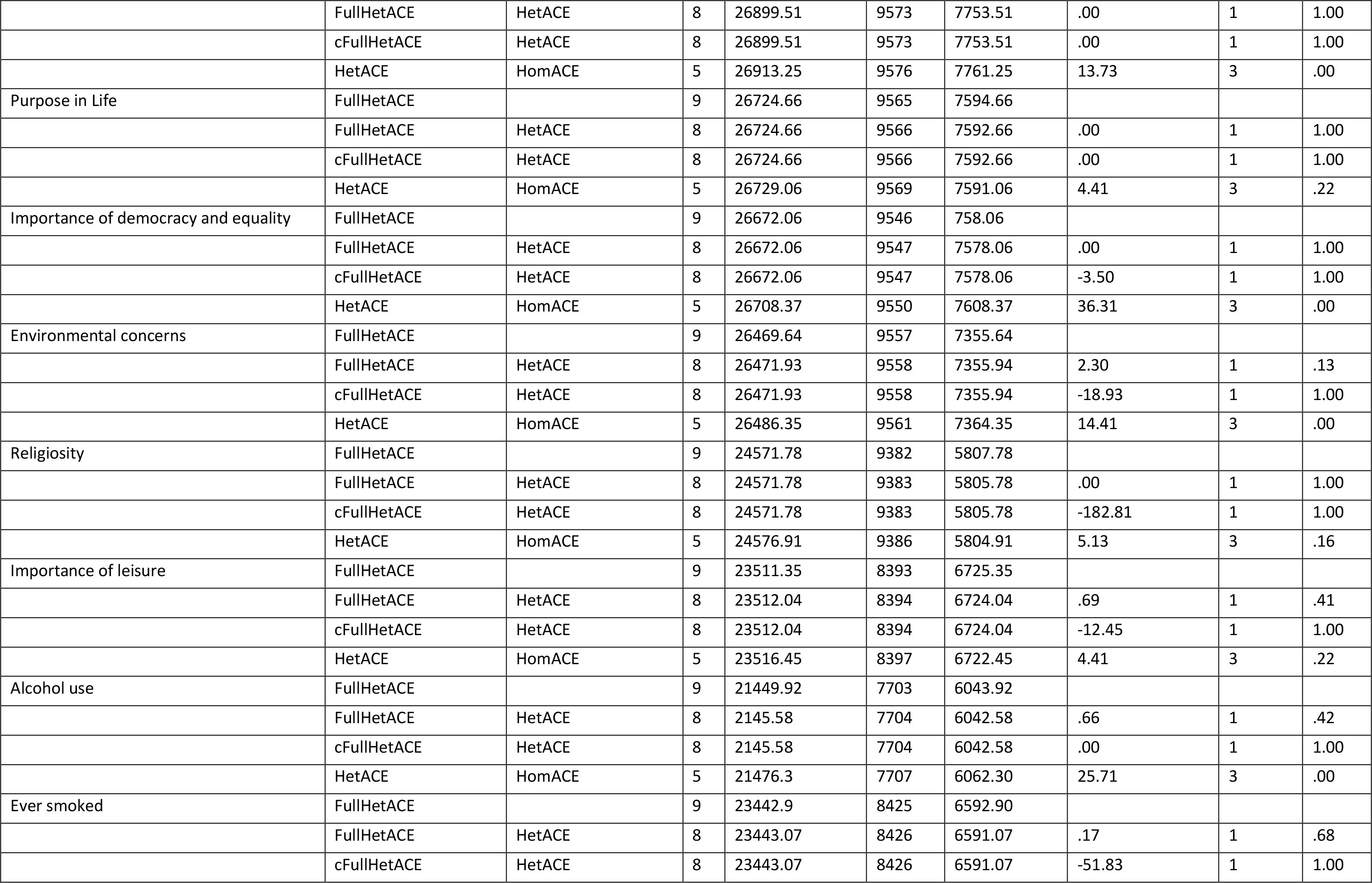
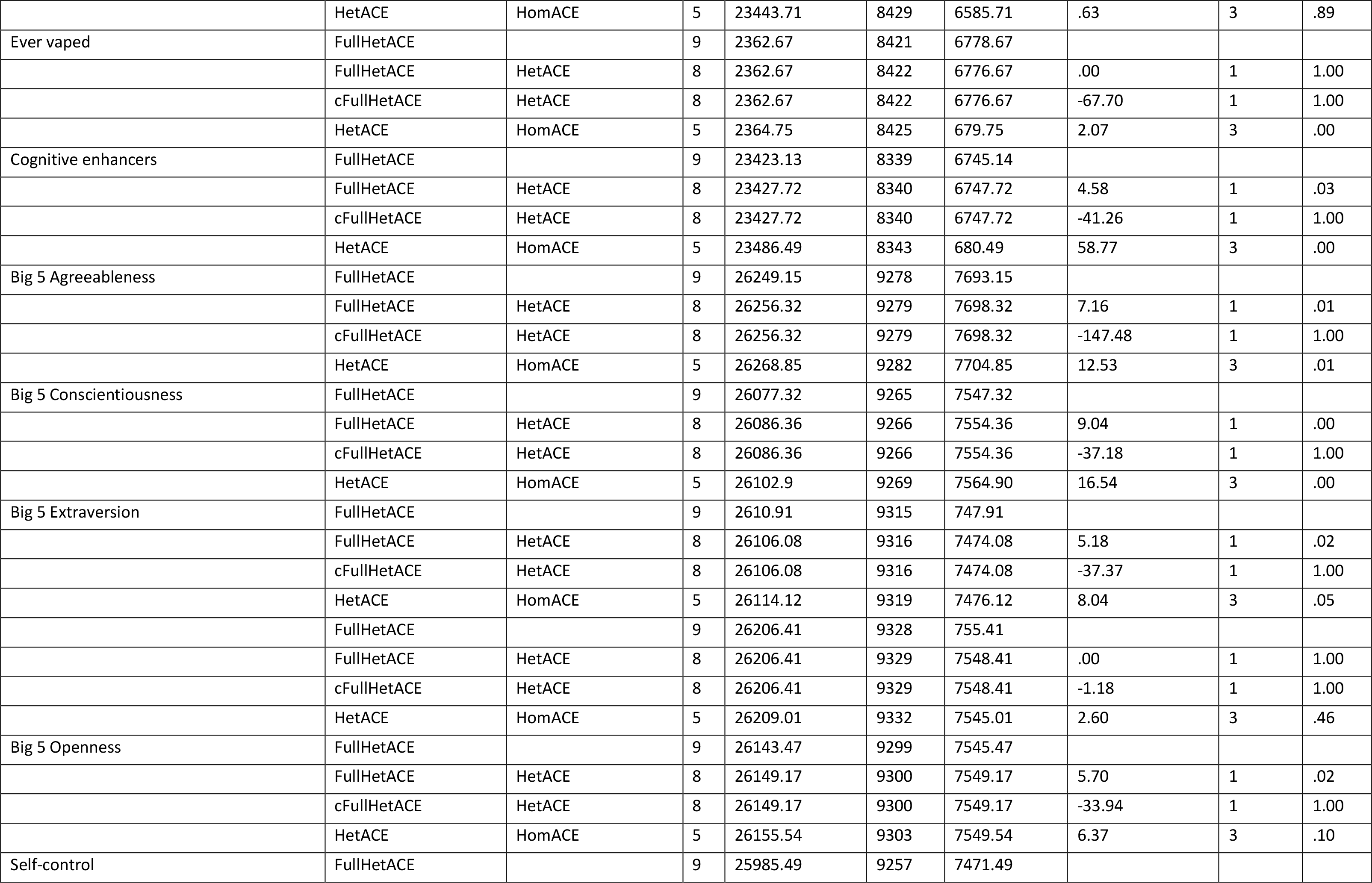
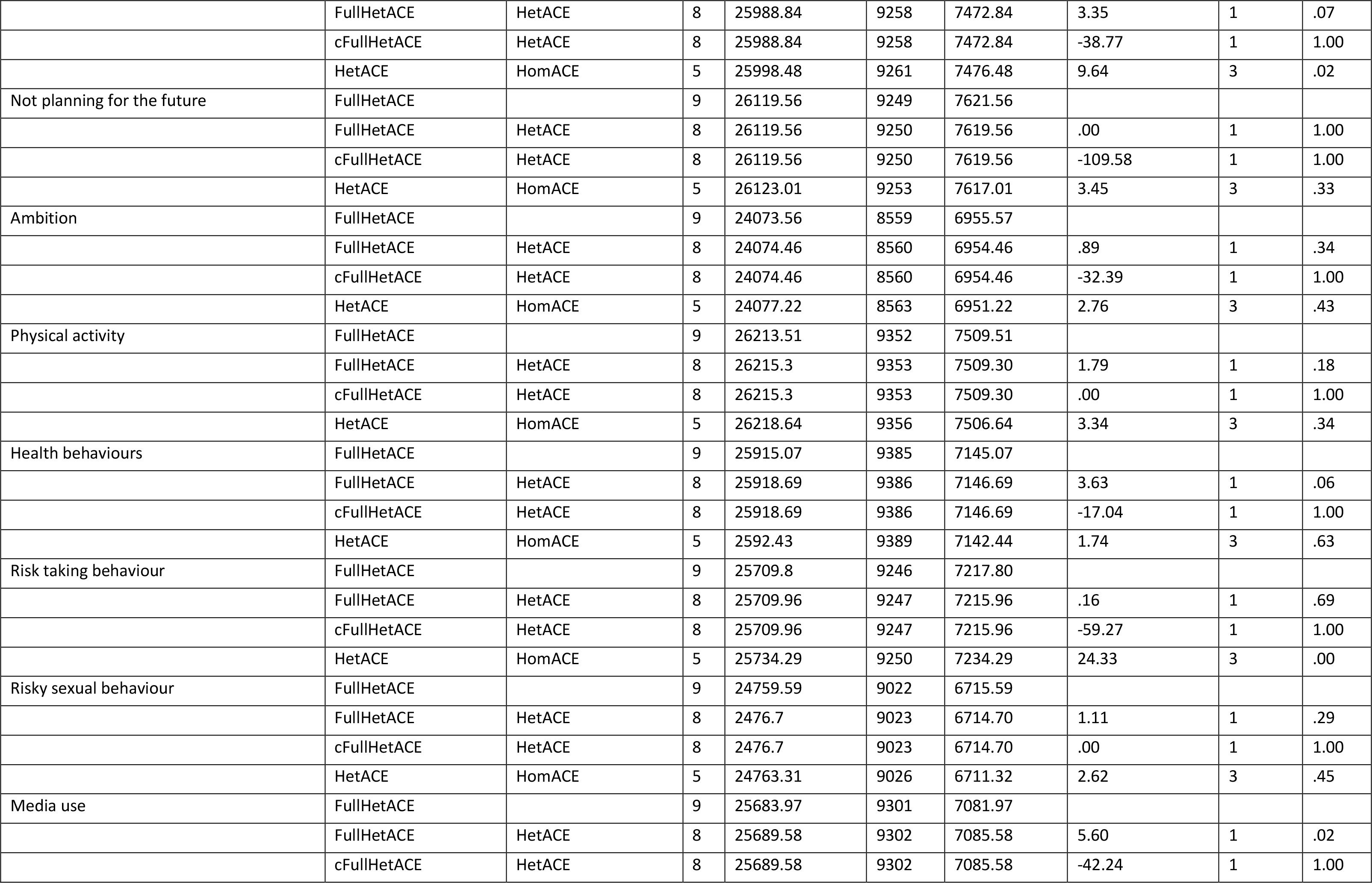
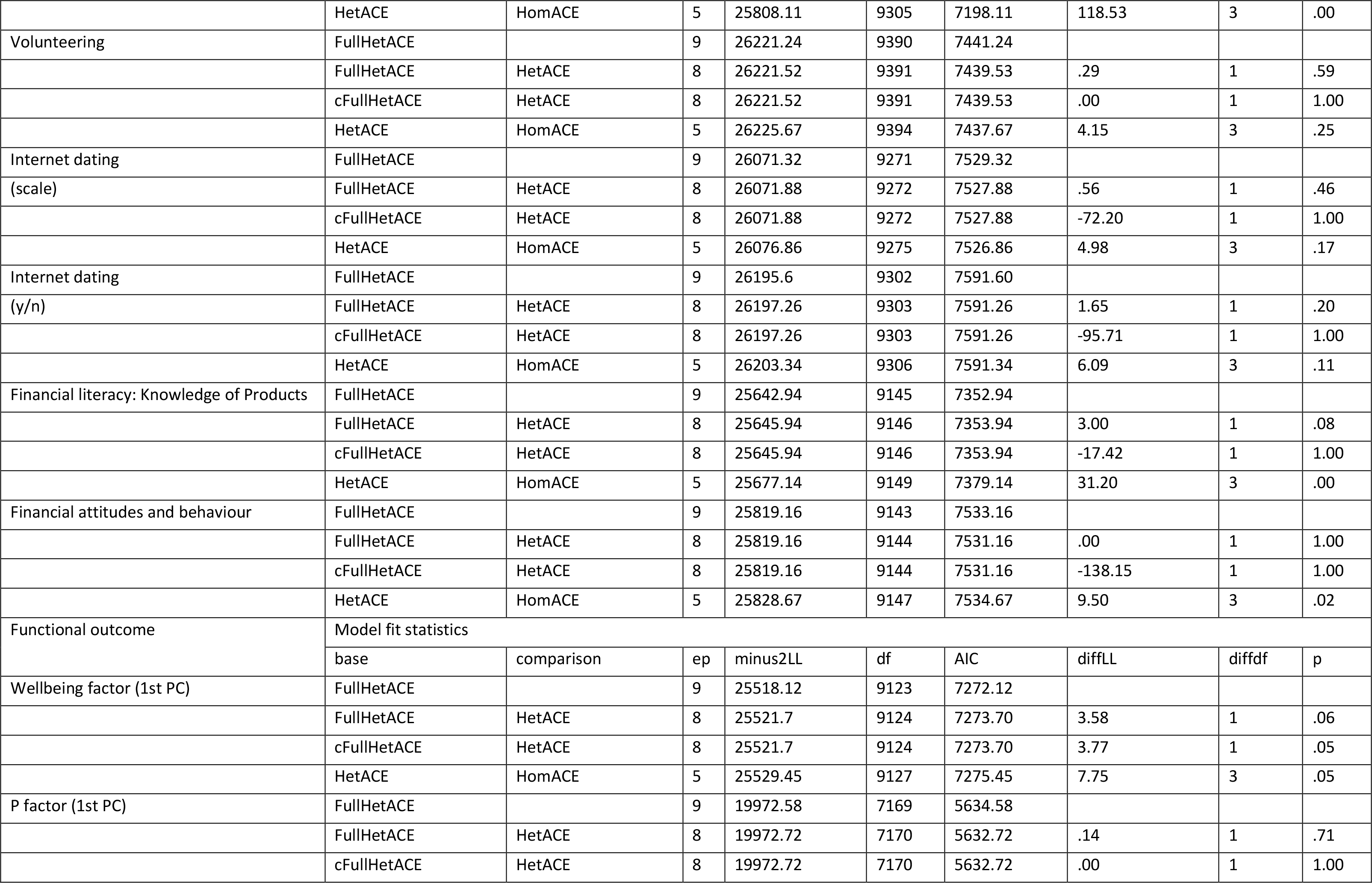
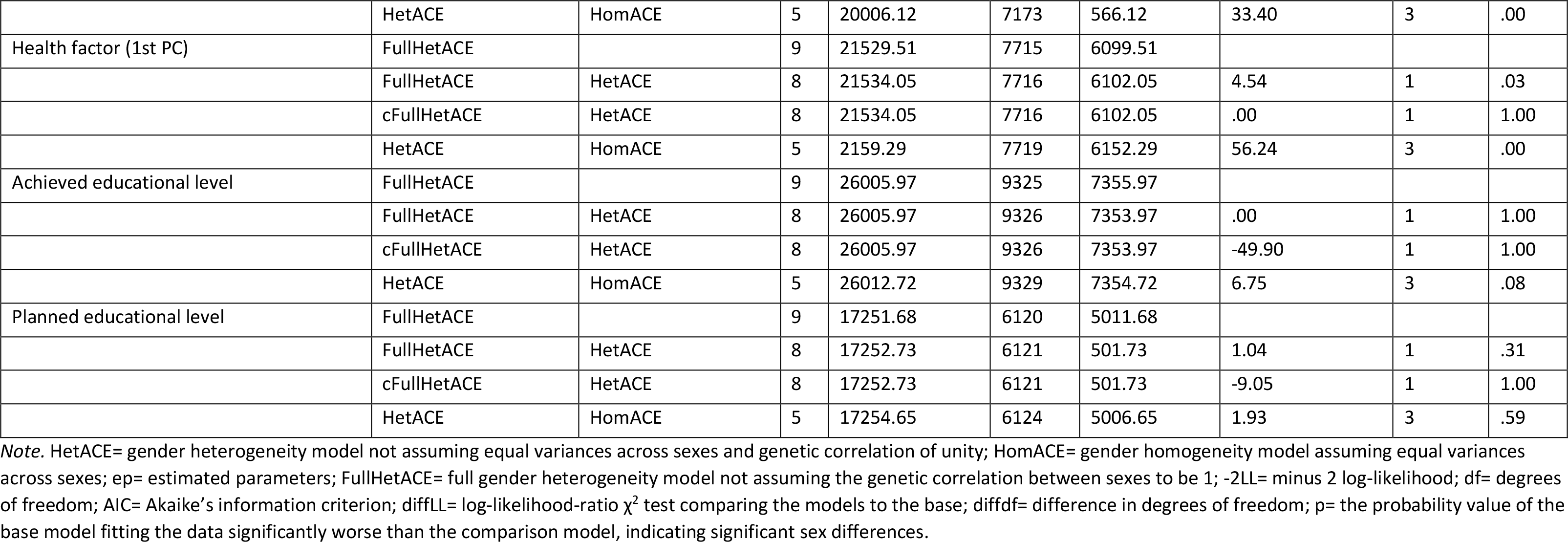
Model-fit statistics for the sex limitation twin analyses.

**Table S12.**
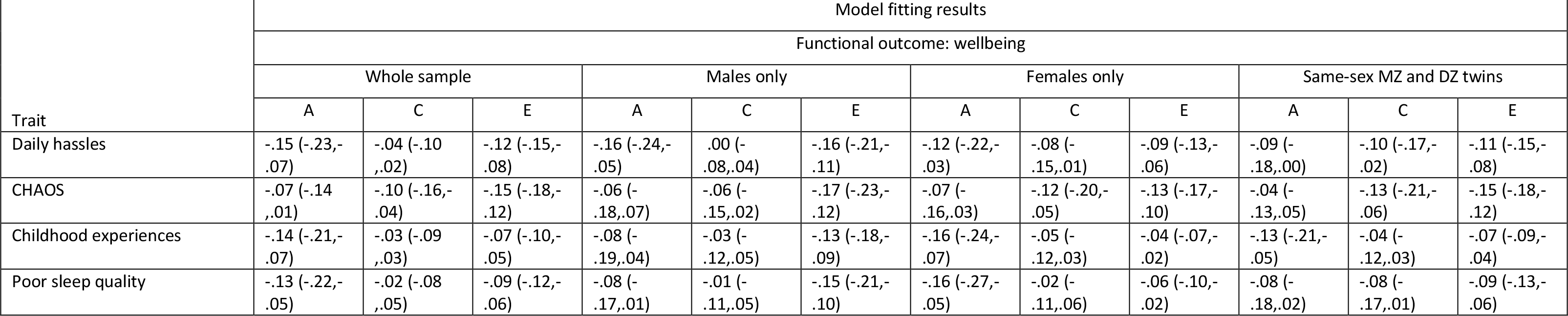
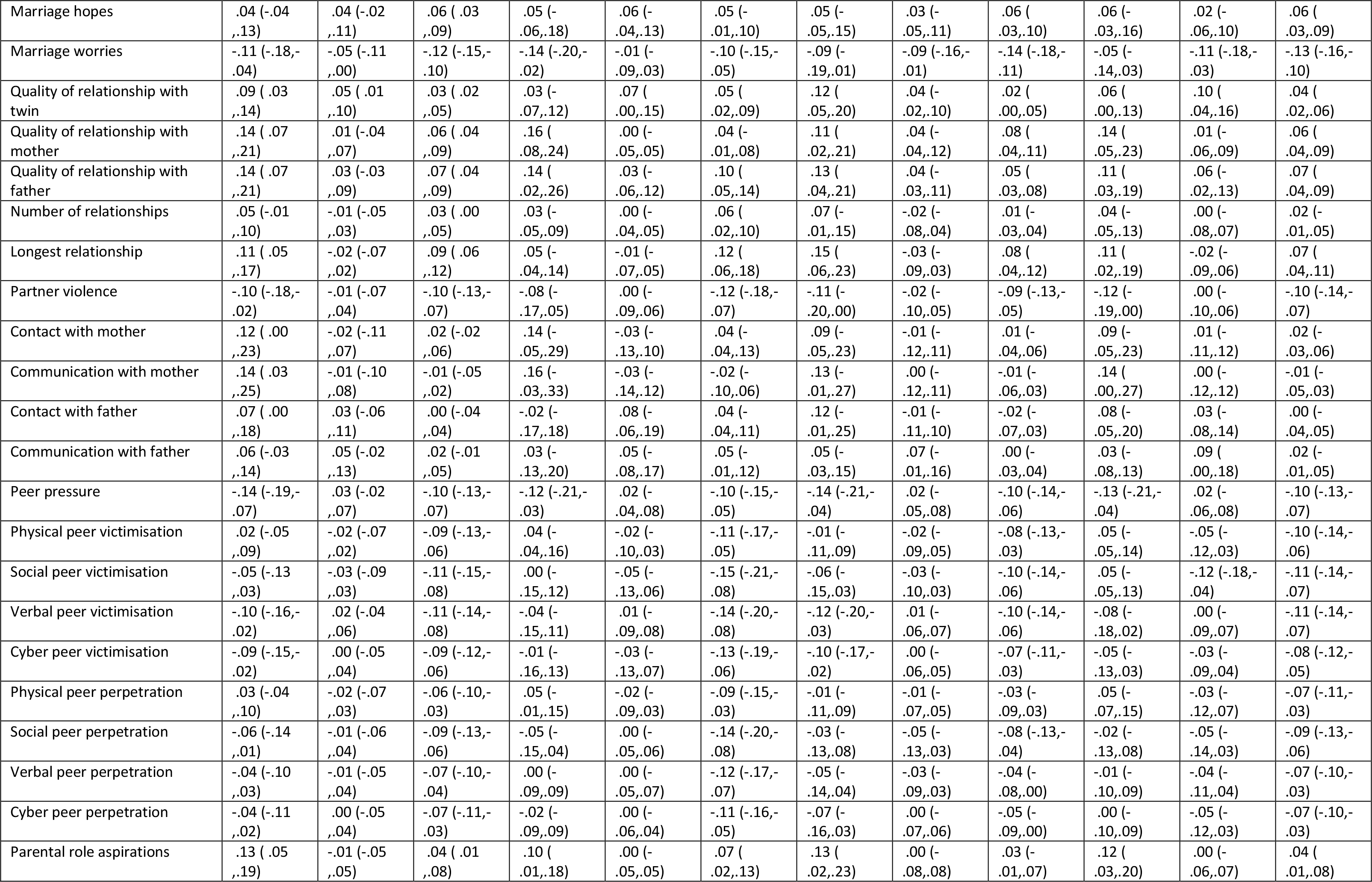
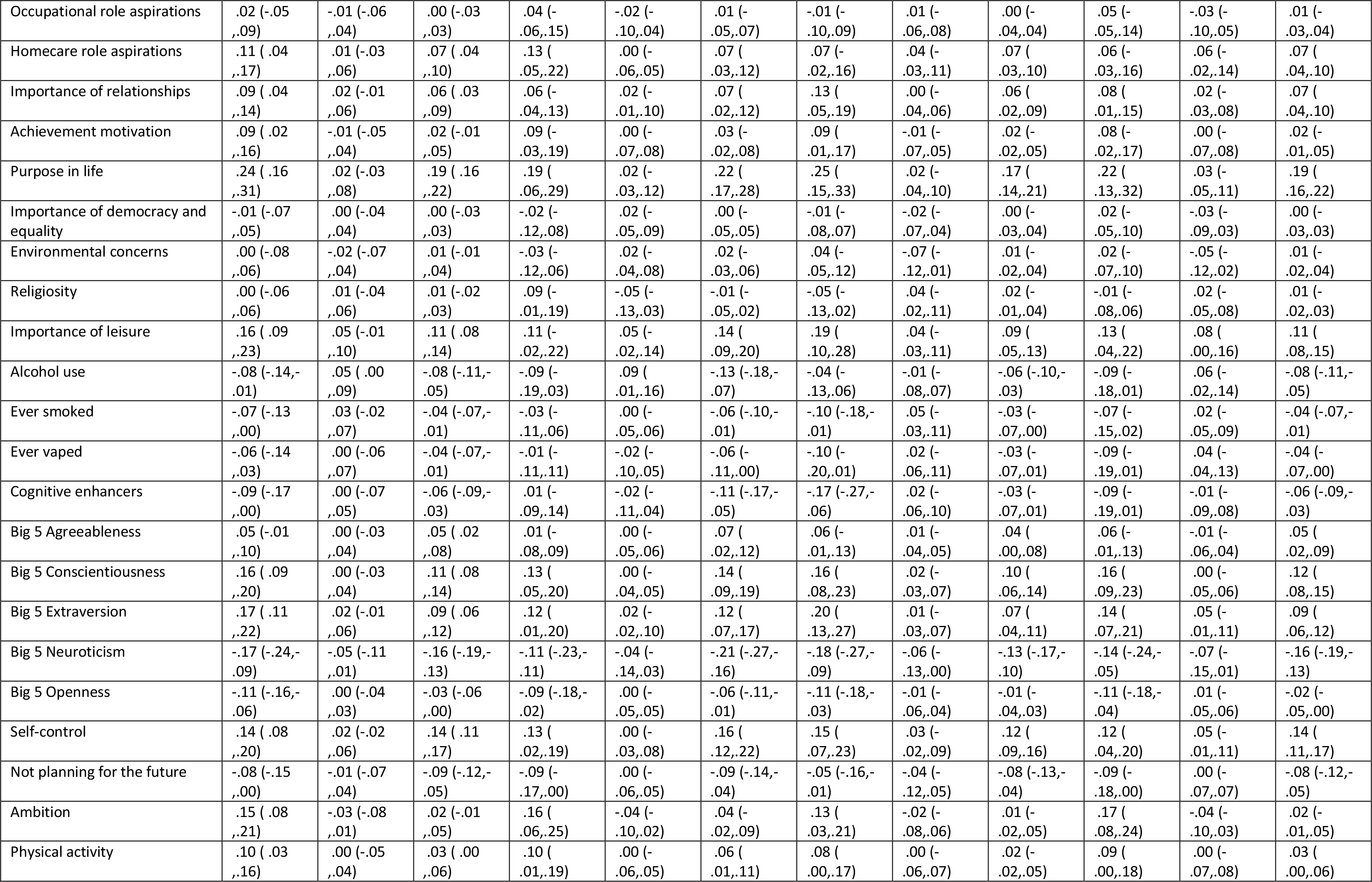
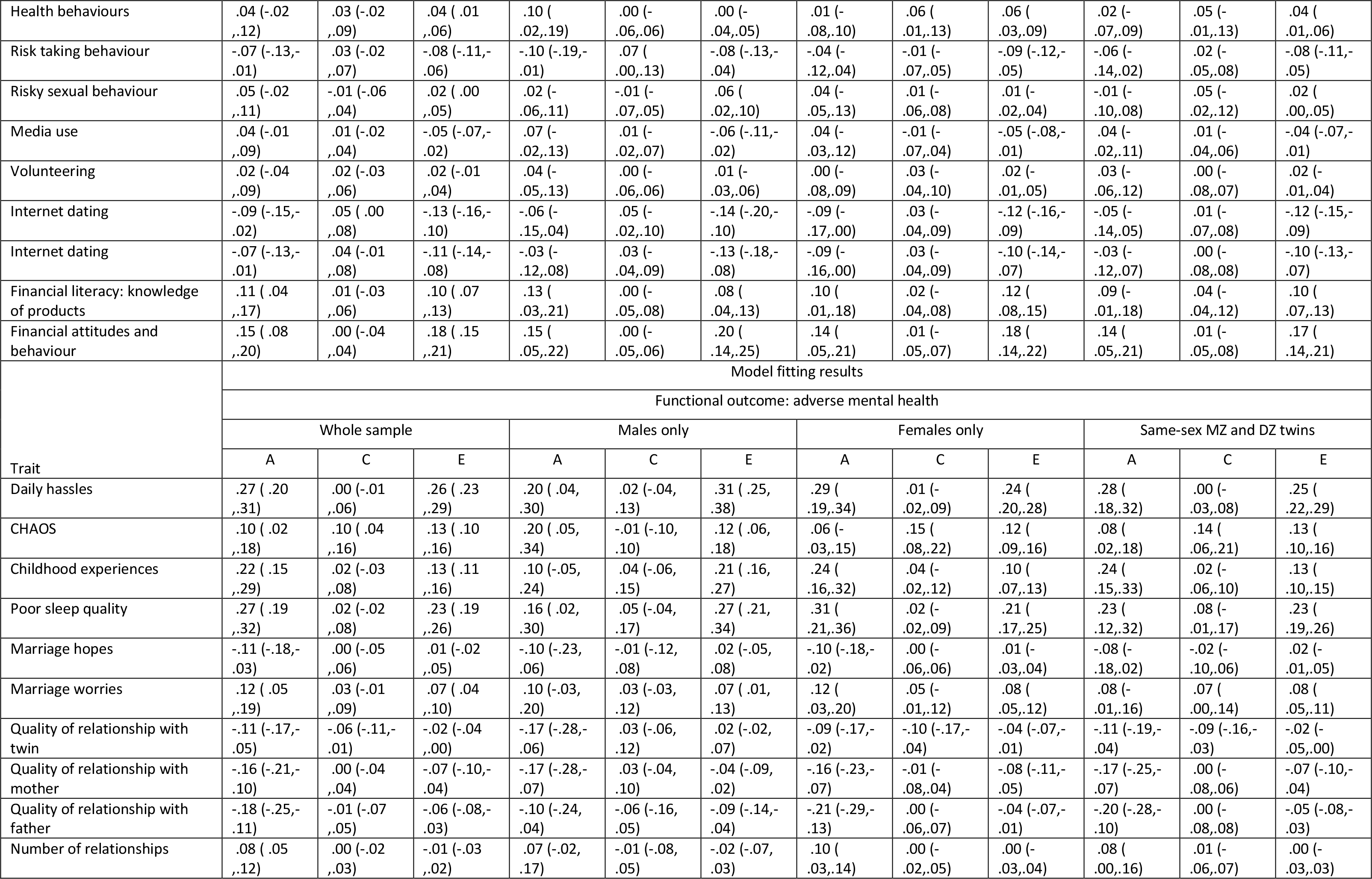
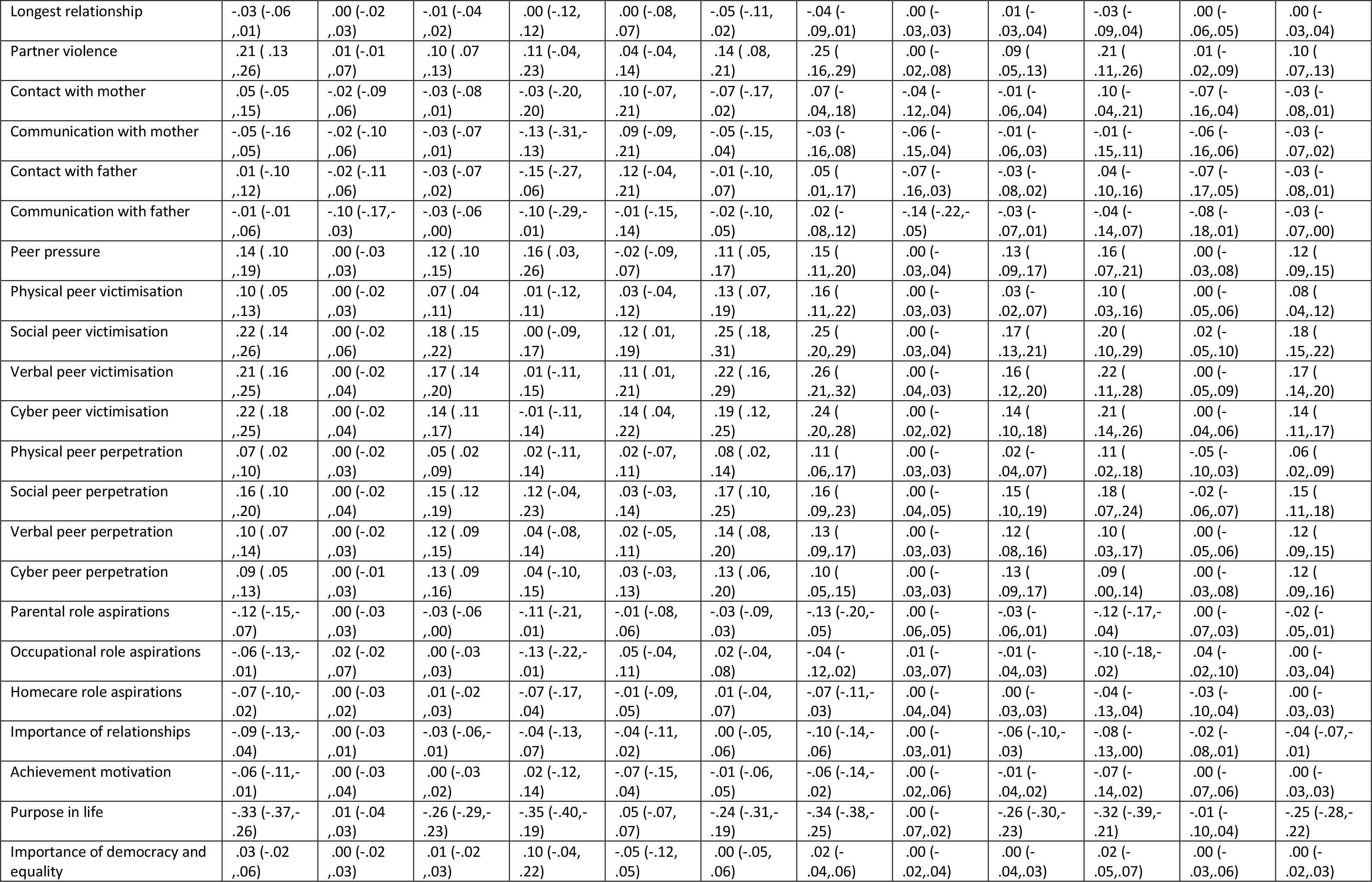
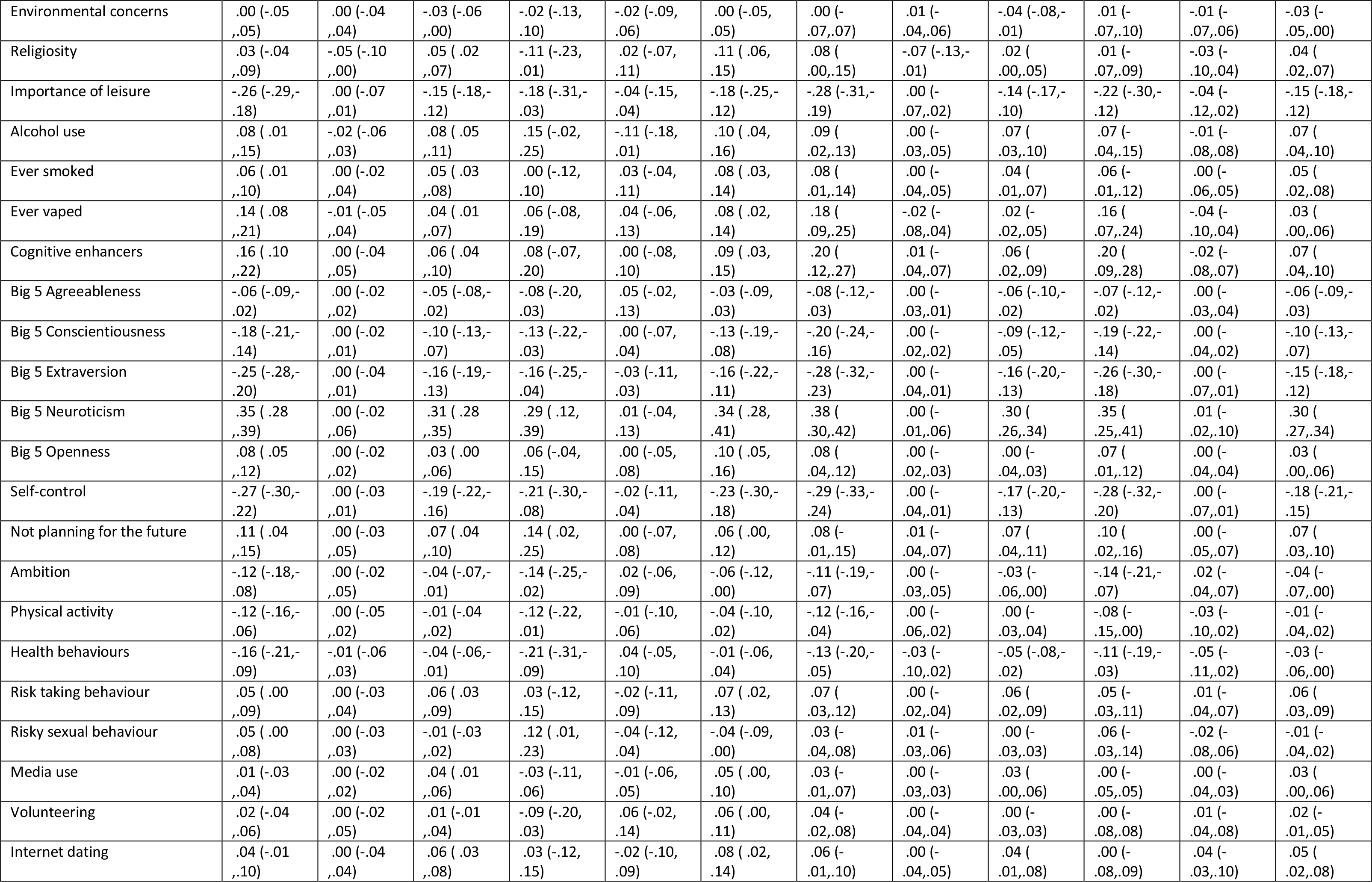
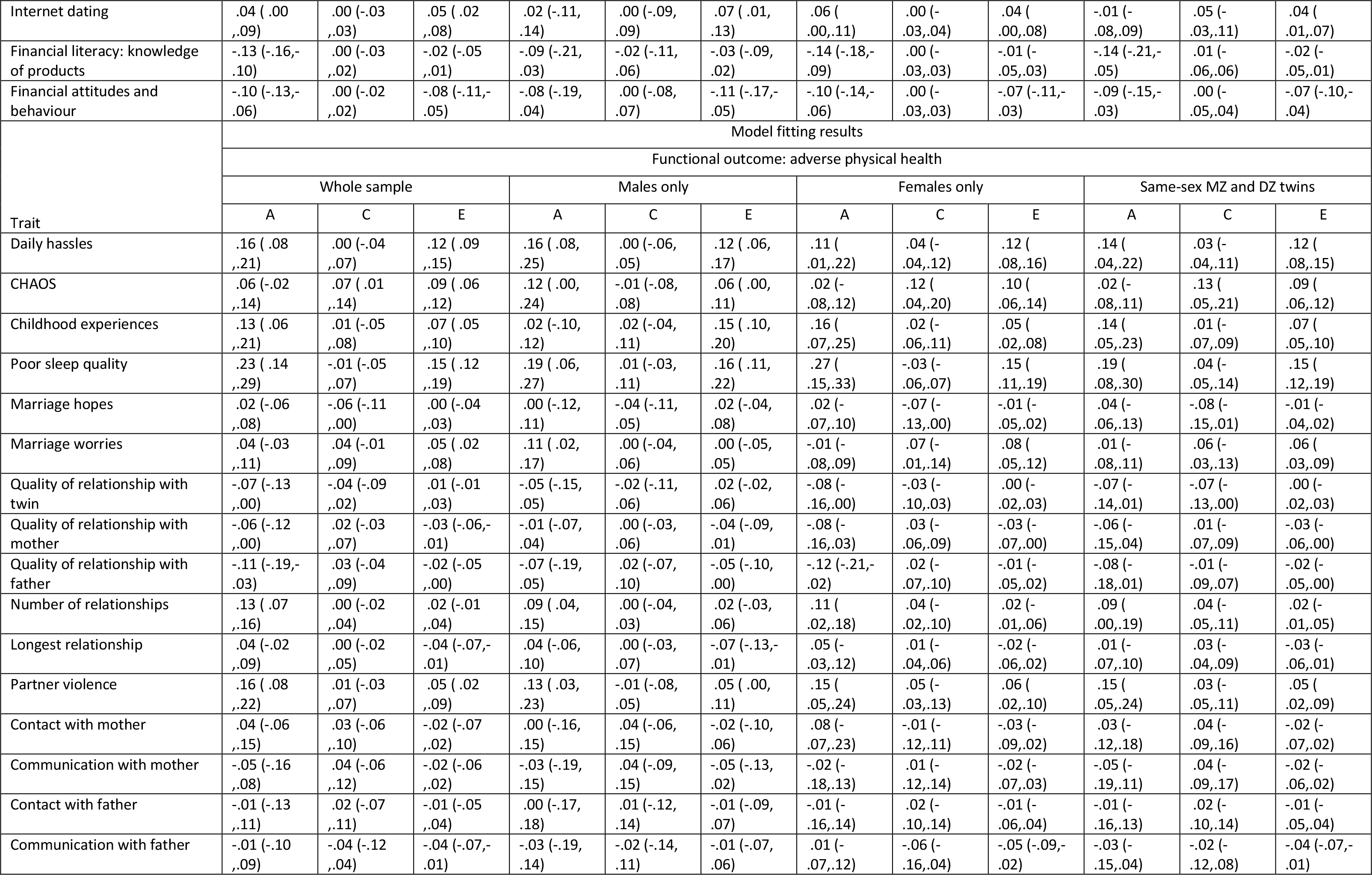
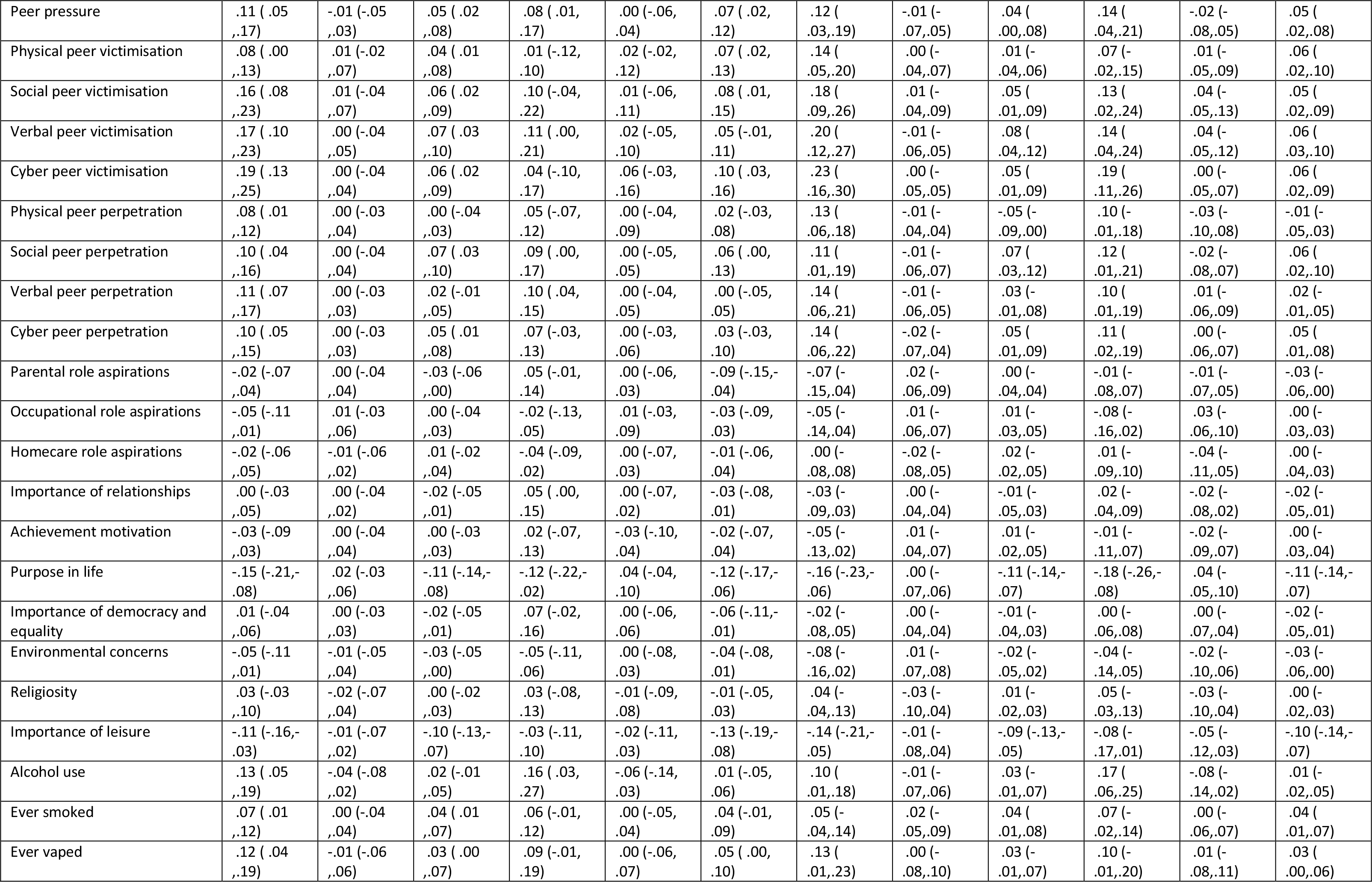
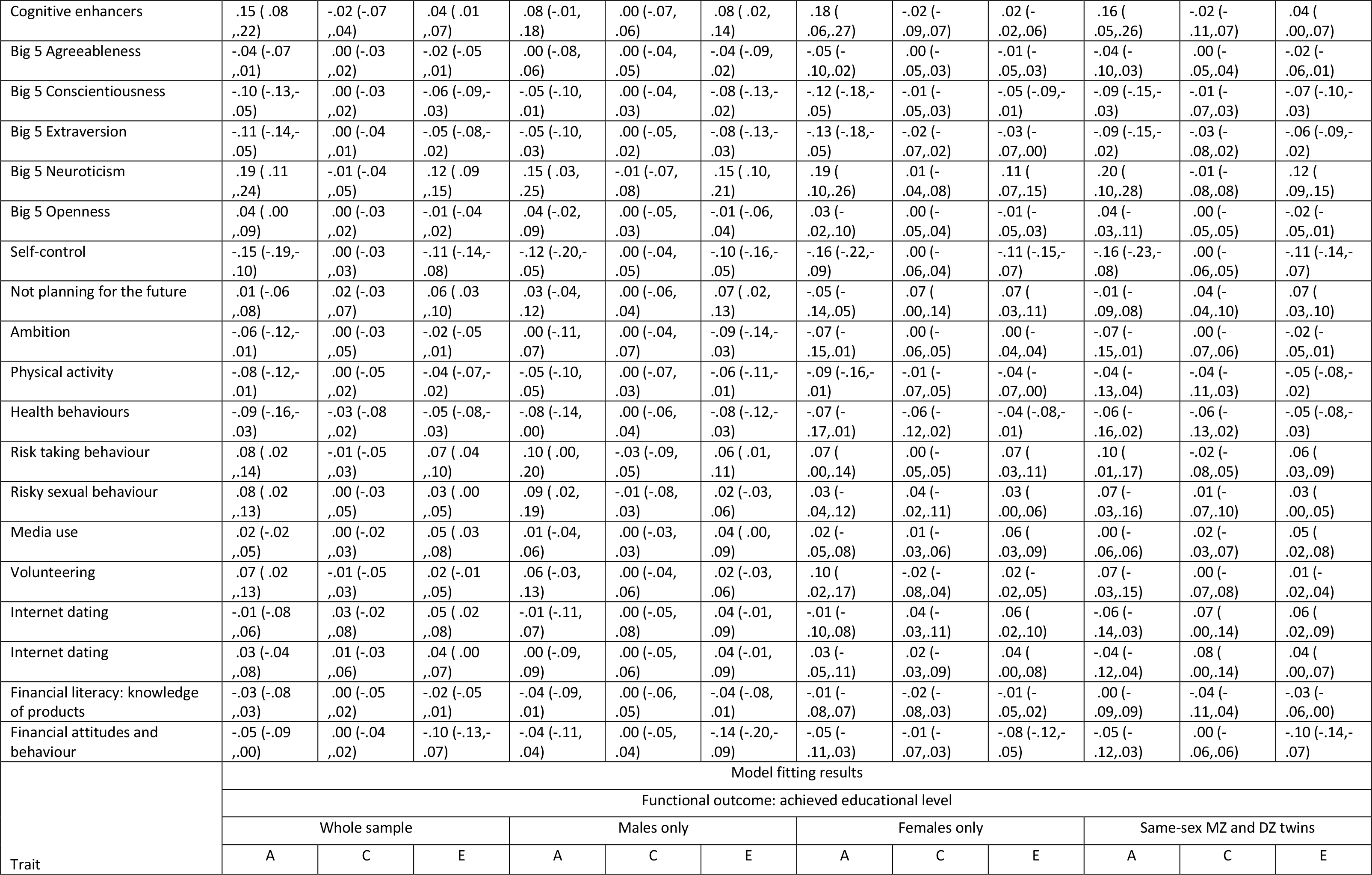
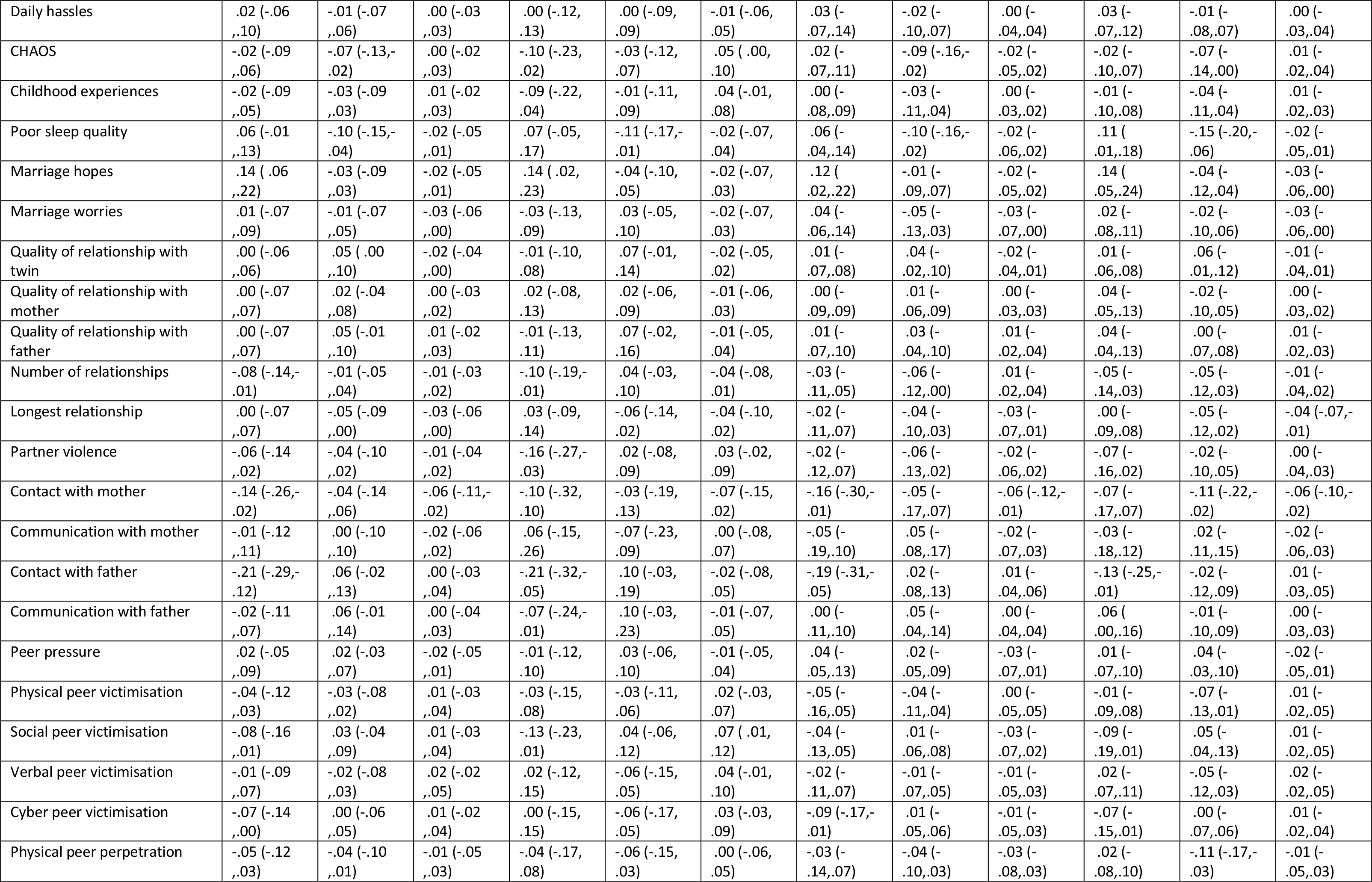
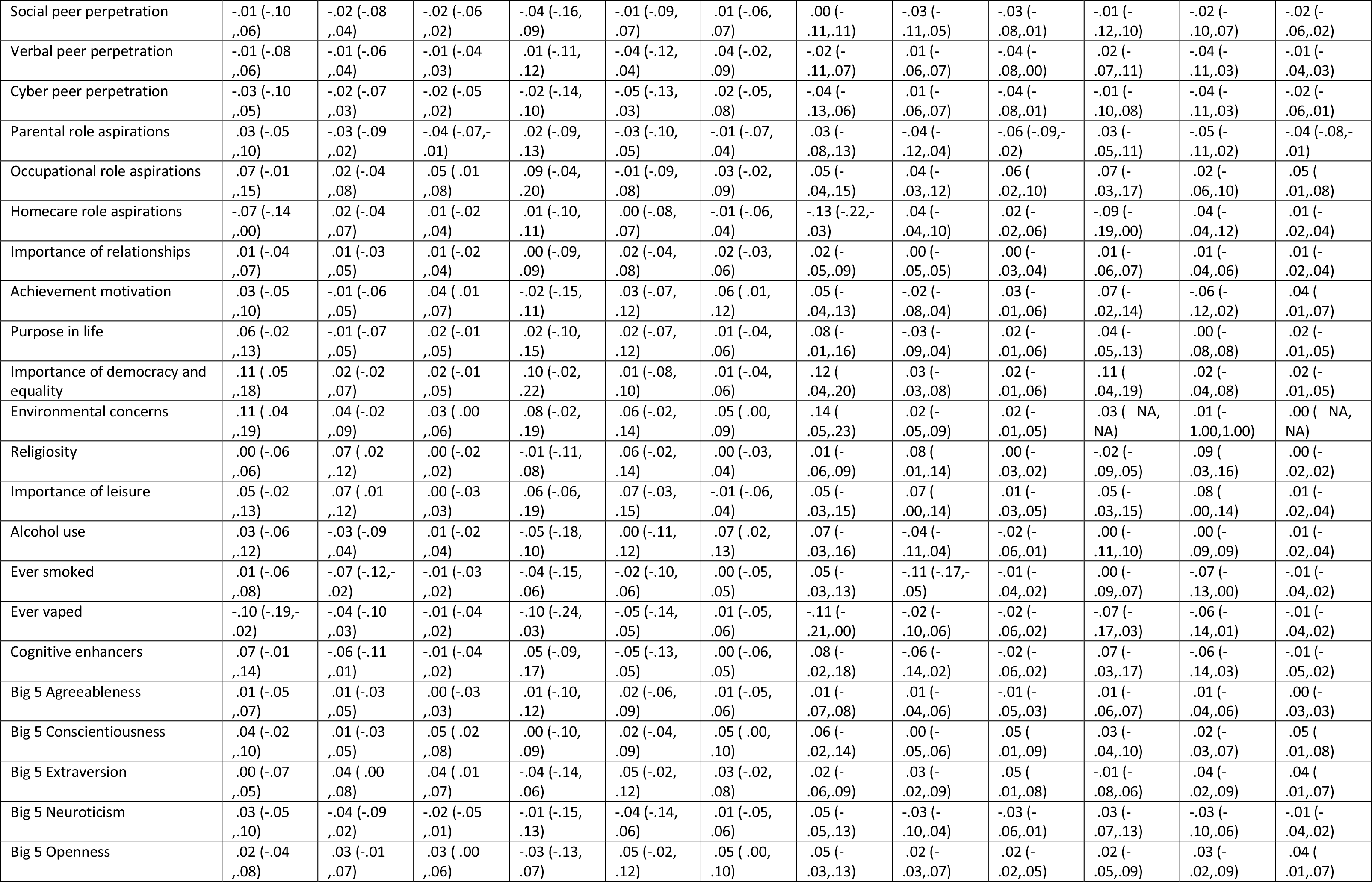
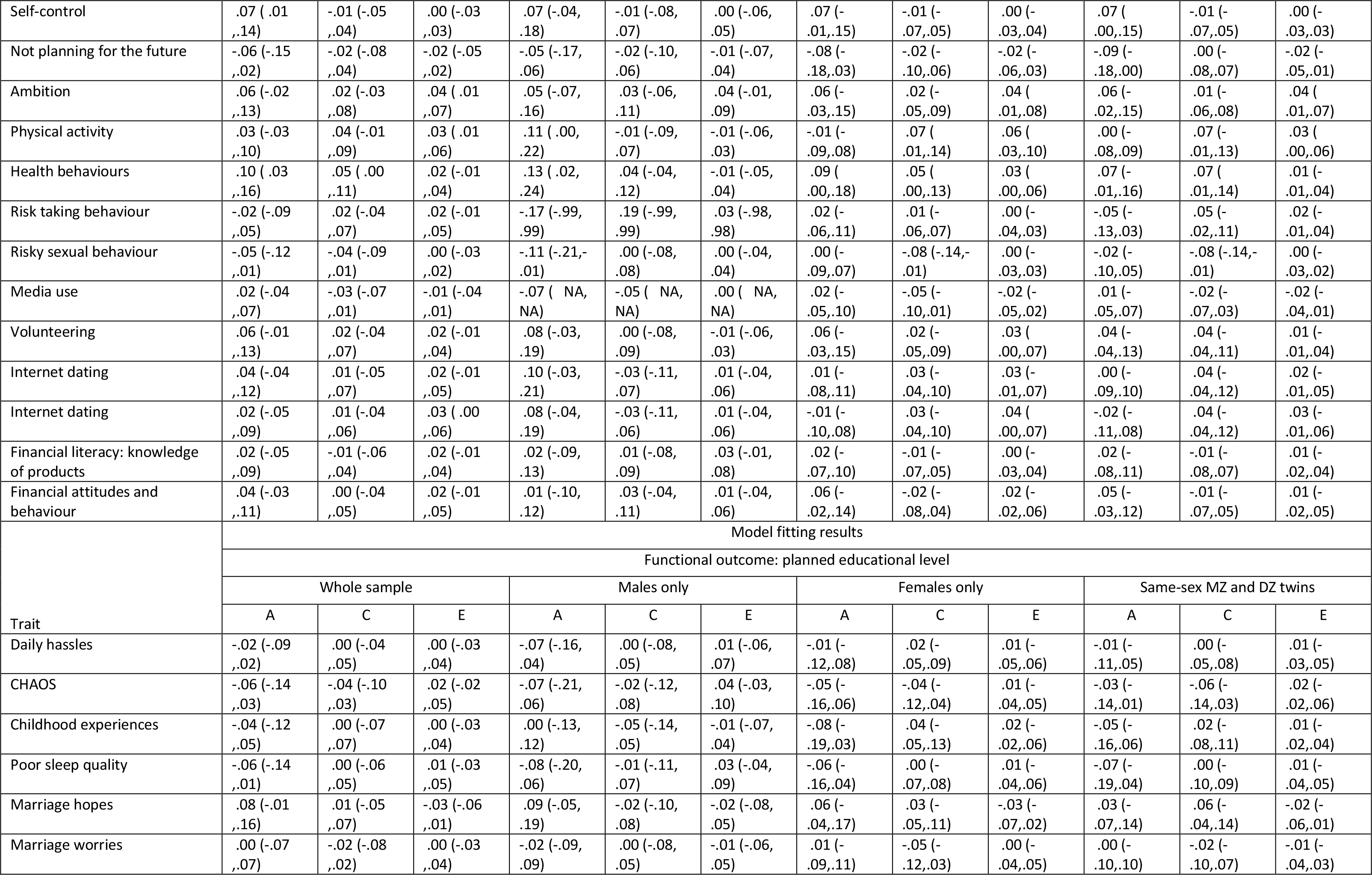
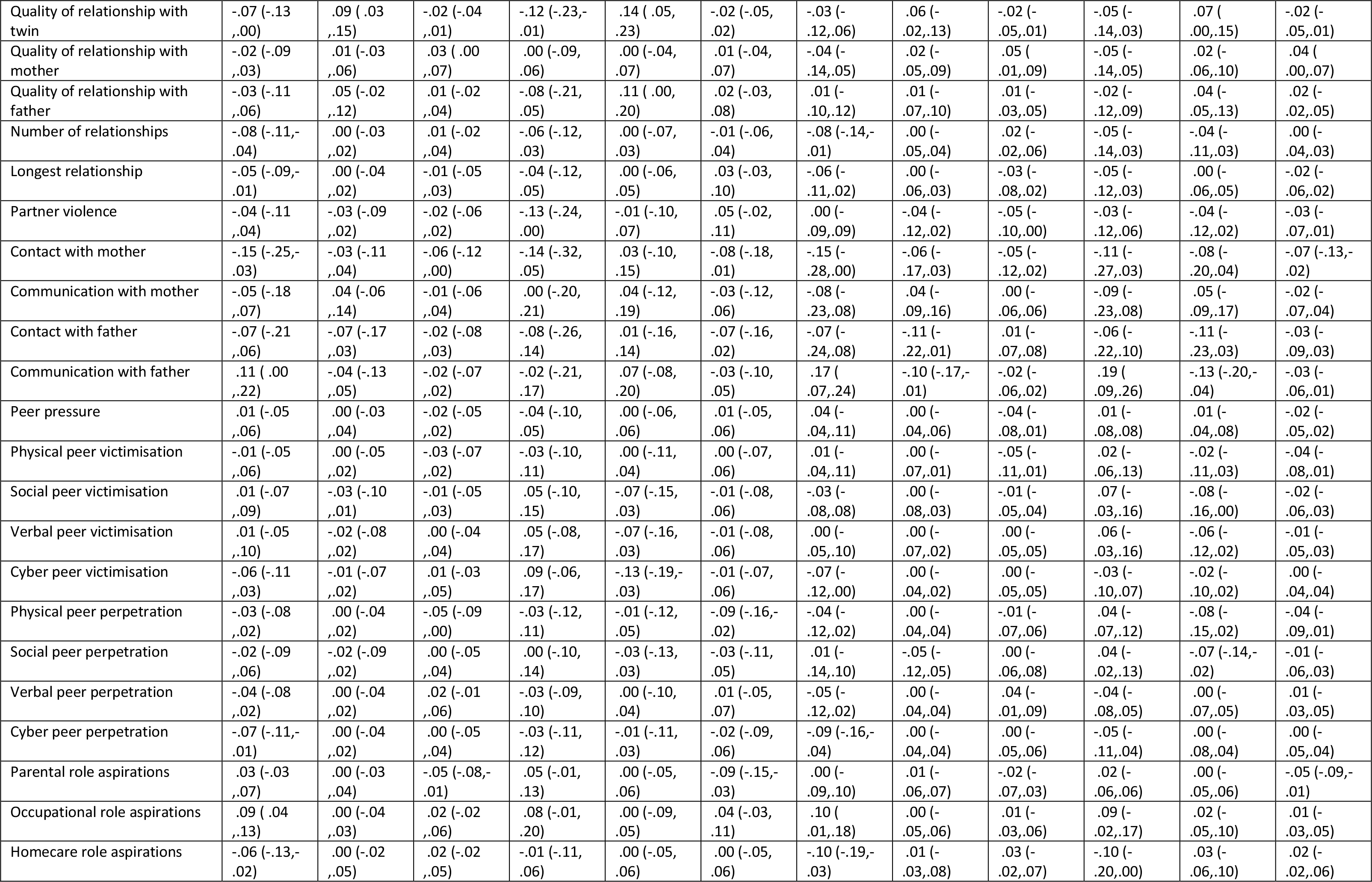
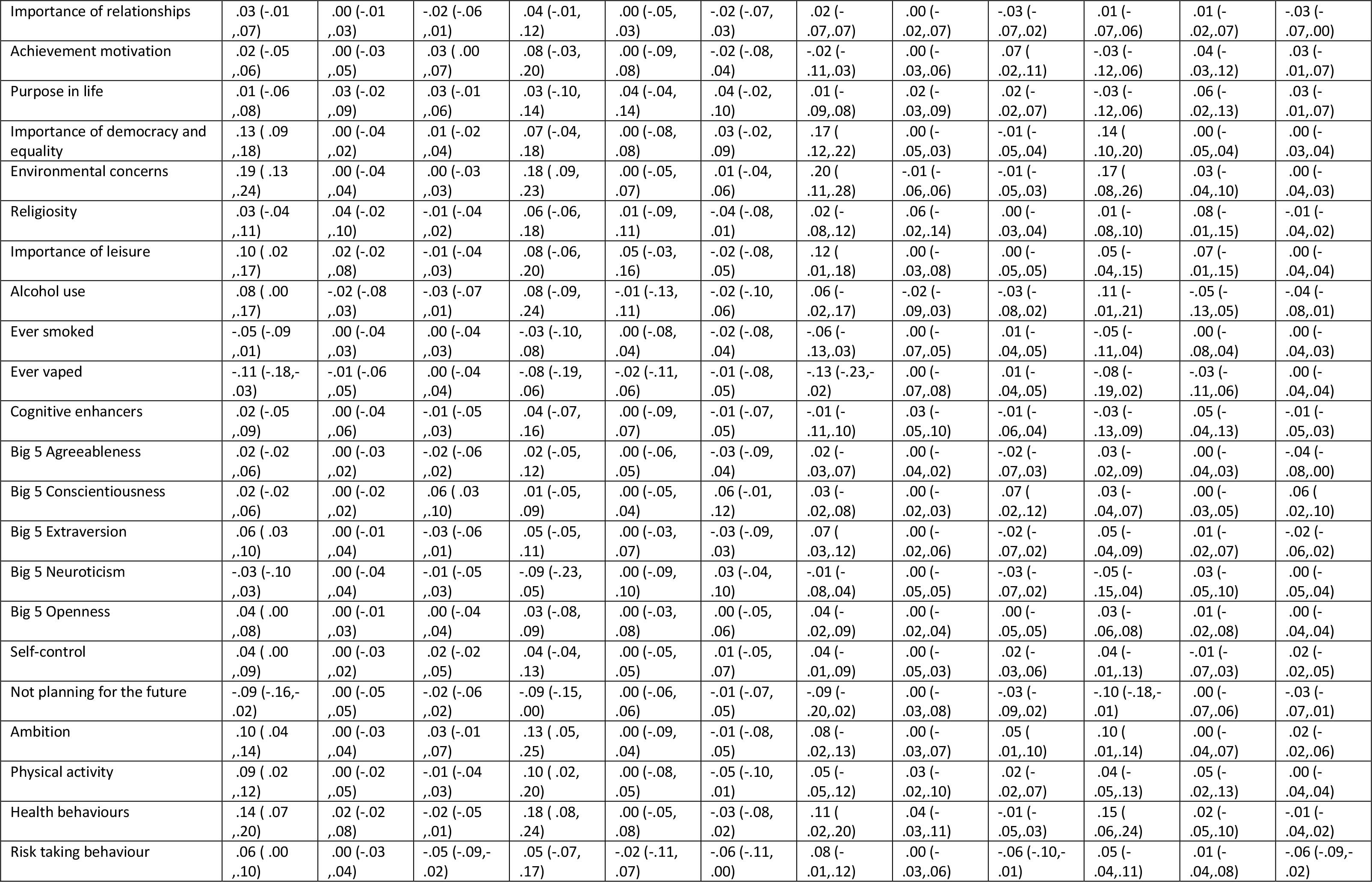
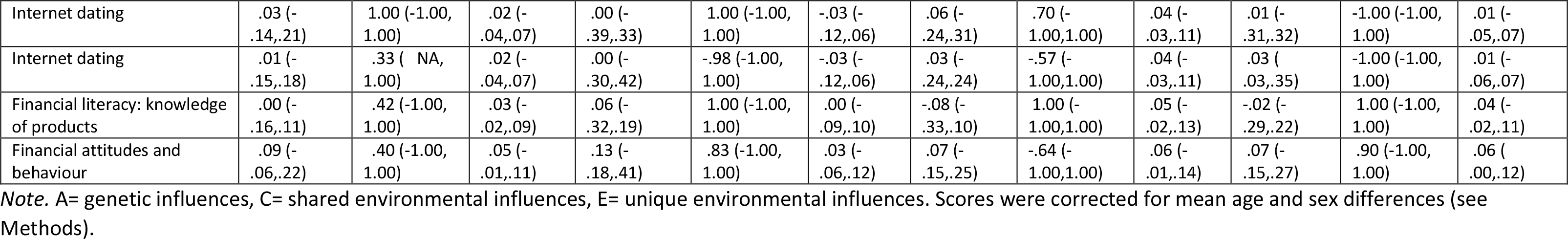
Model fitting results for bivariate analyses of the correlation between psychological traits and functional outcomes explained by genetic (A), shared environmental (C), and non-shared environmental (E) factors (95% confidence intervals are in parentheses).

**Table S13.**
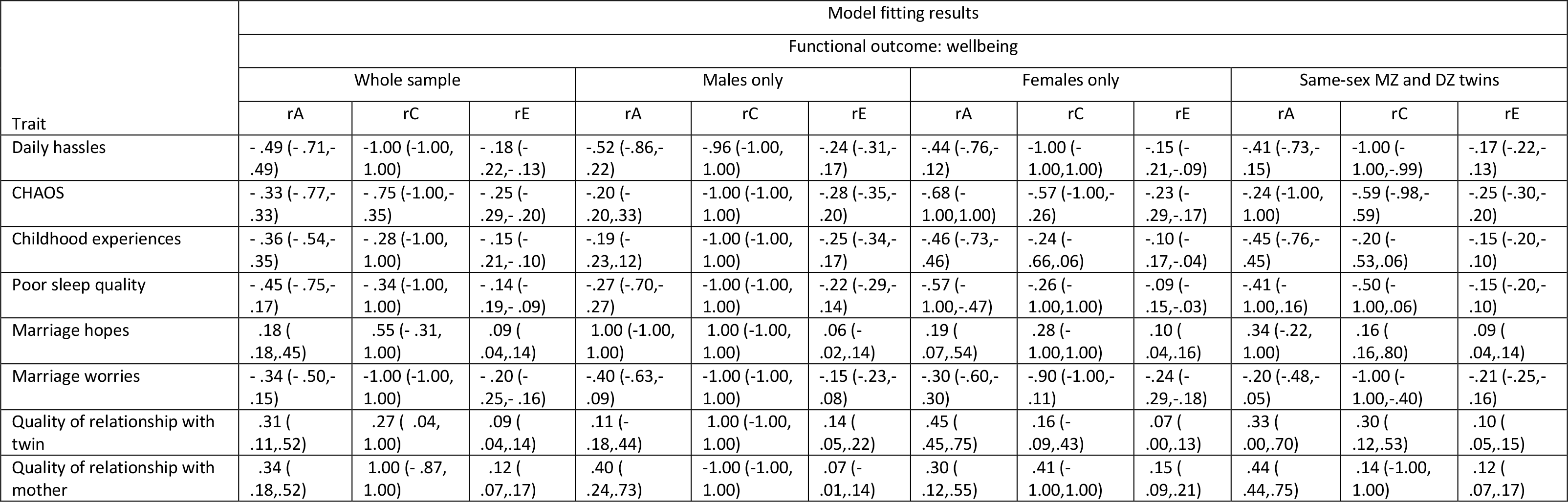
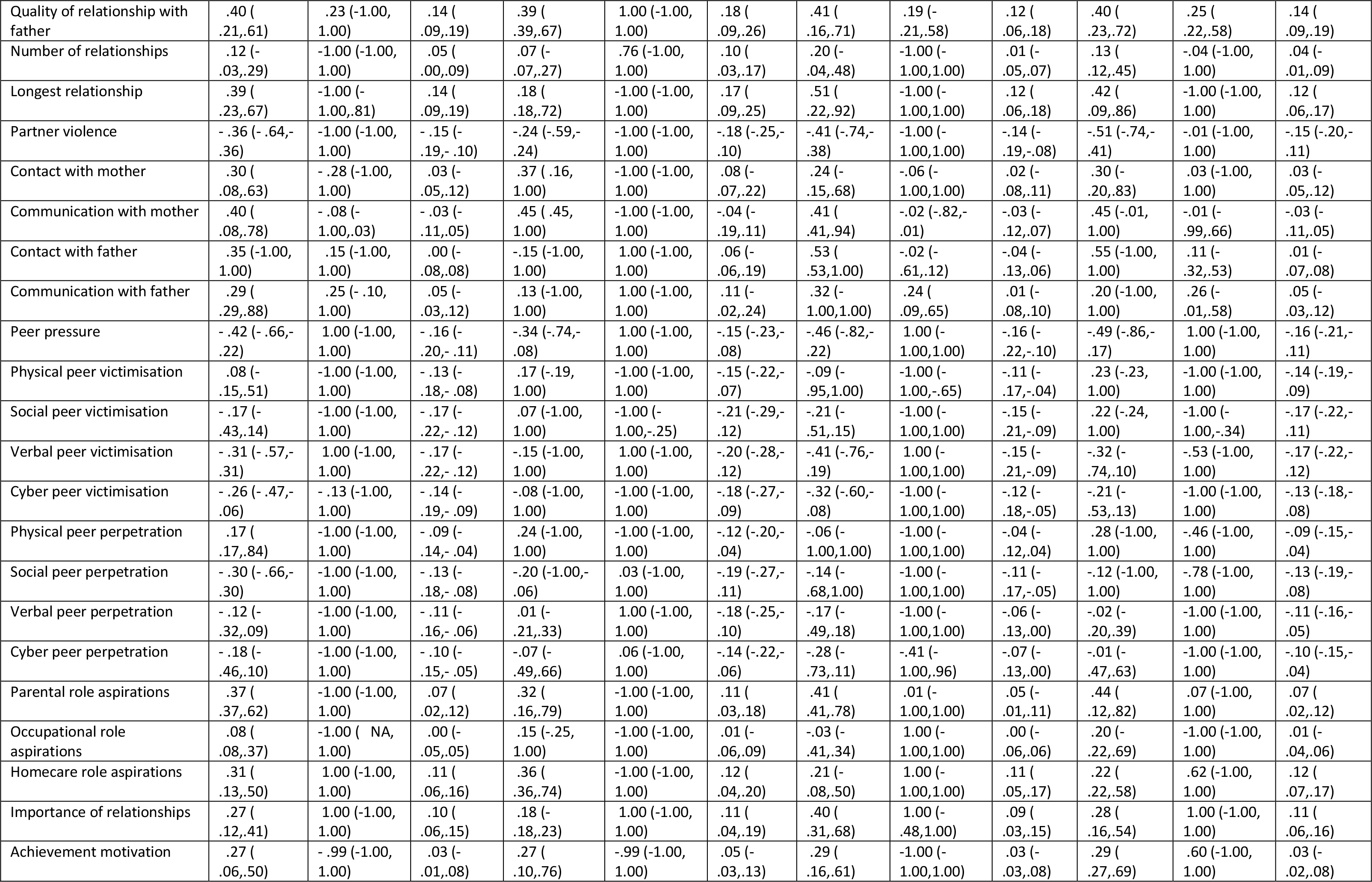
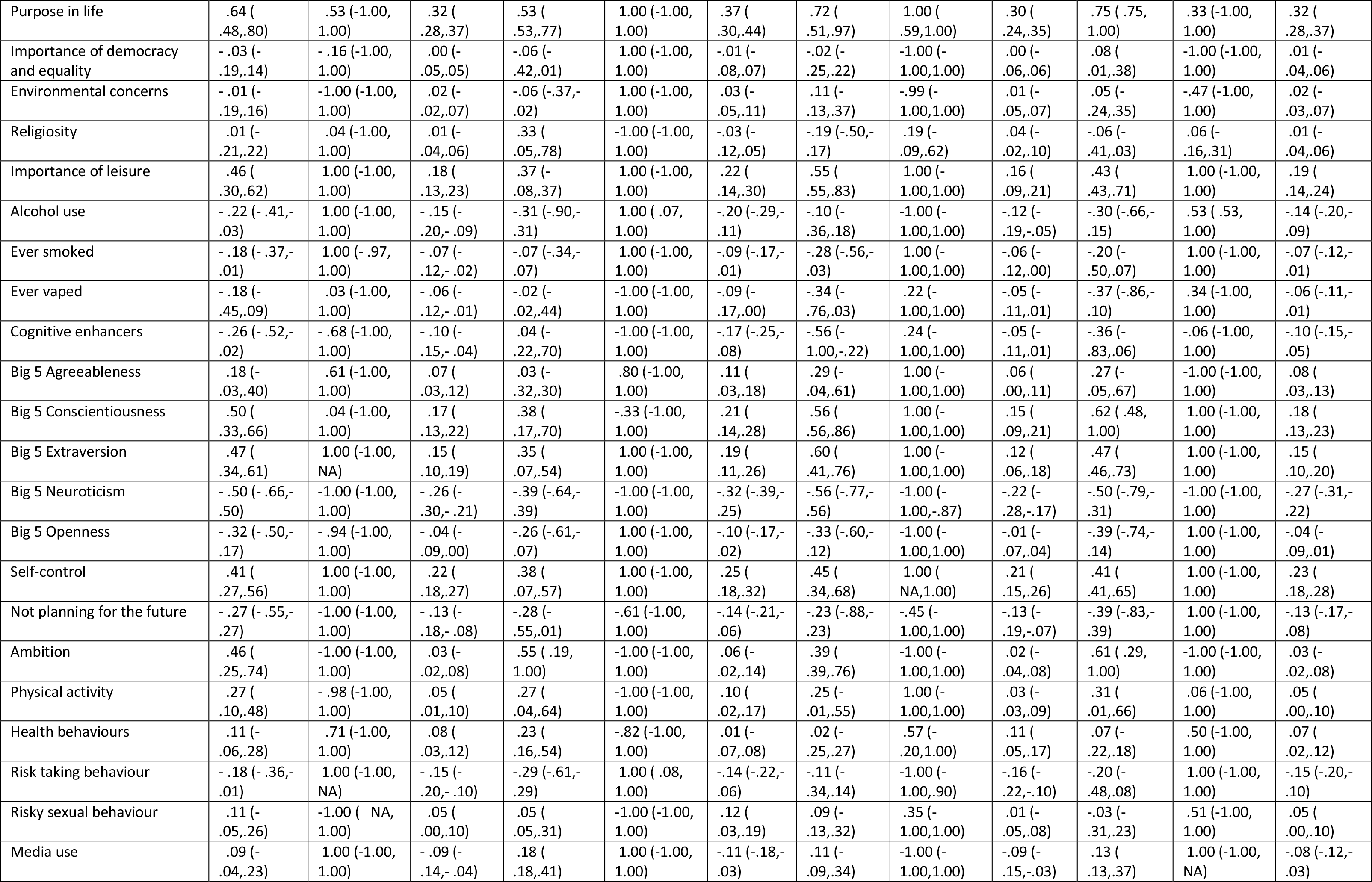
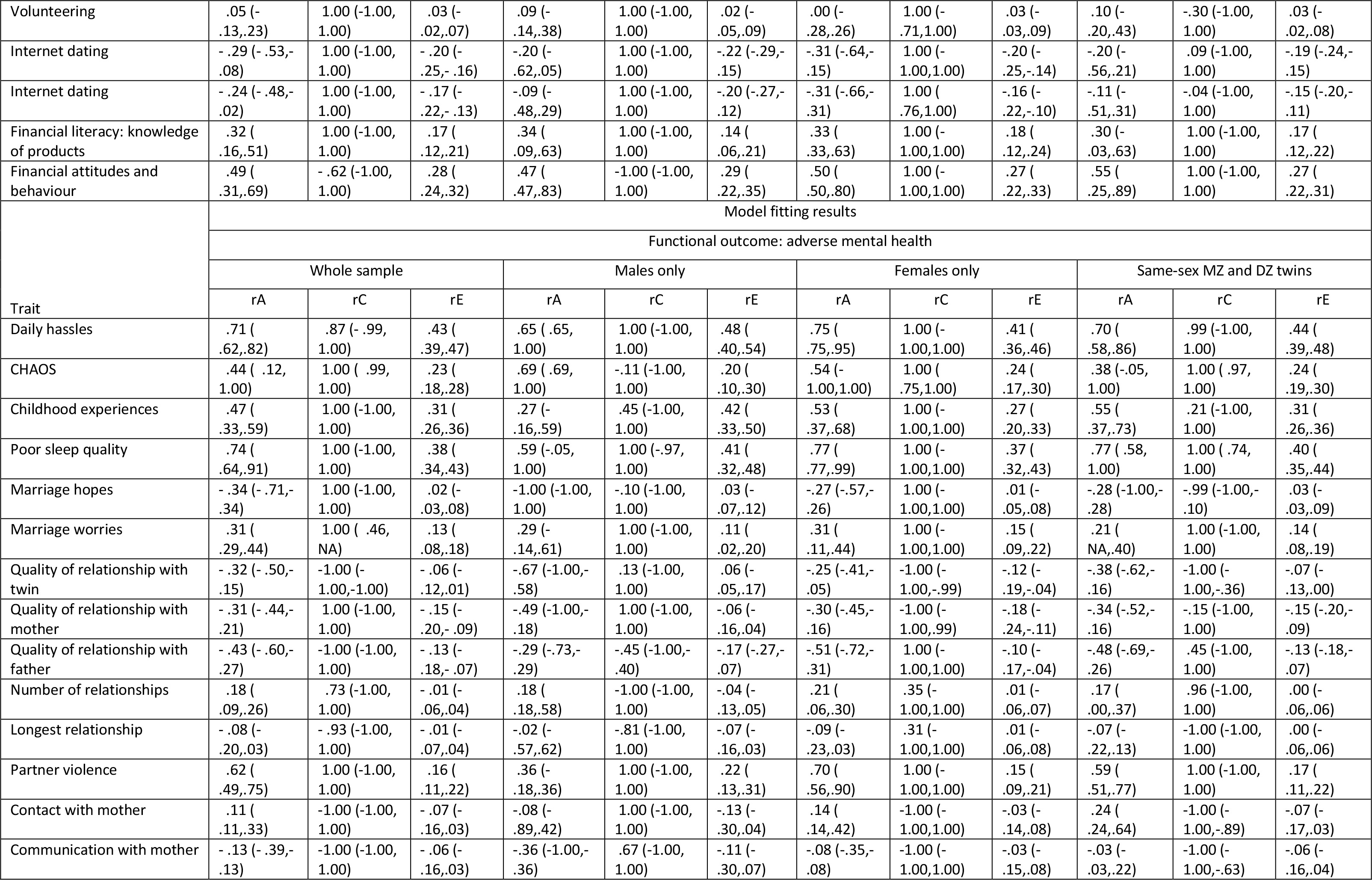
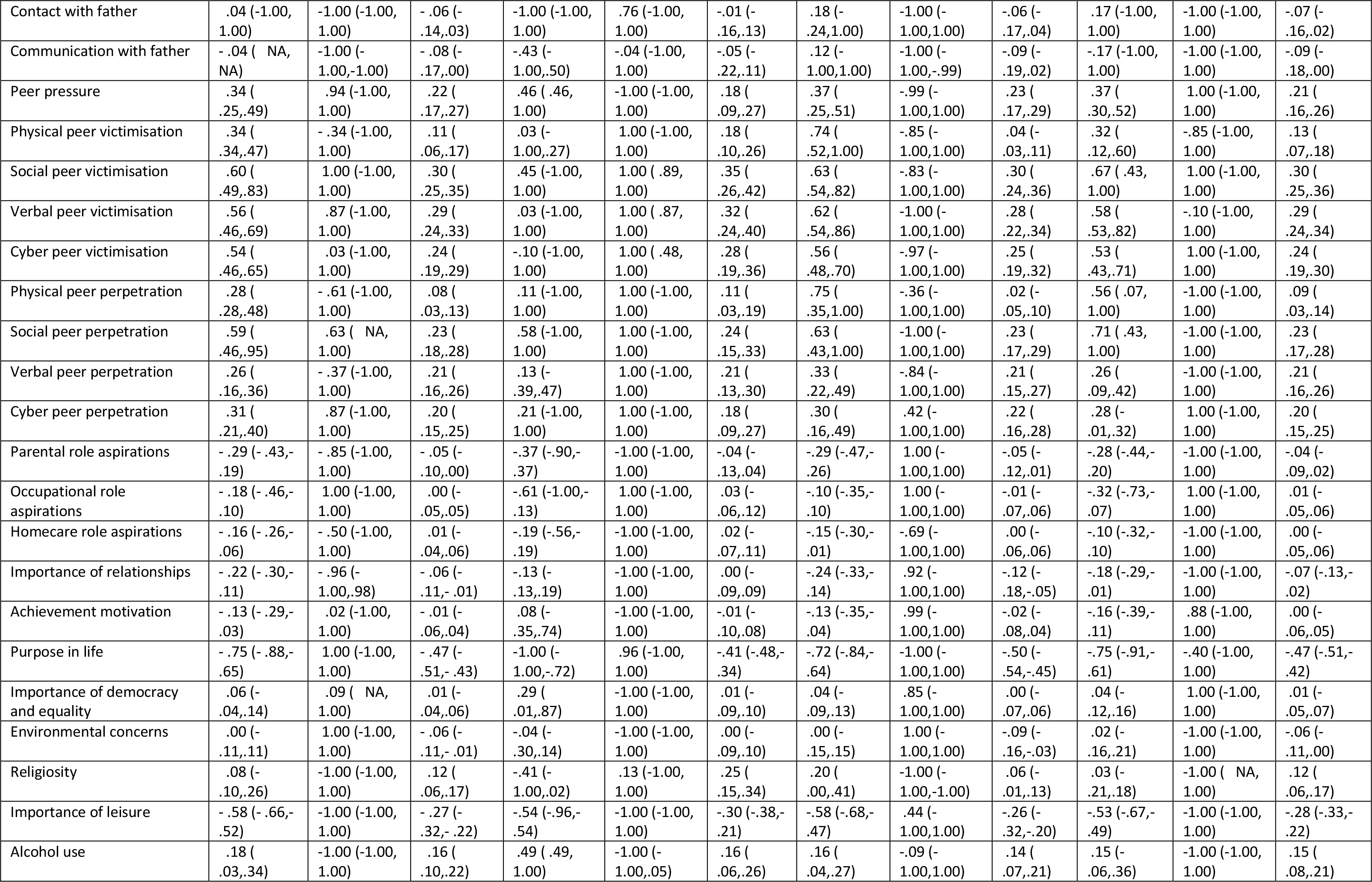
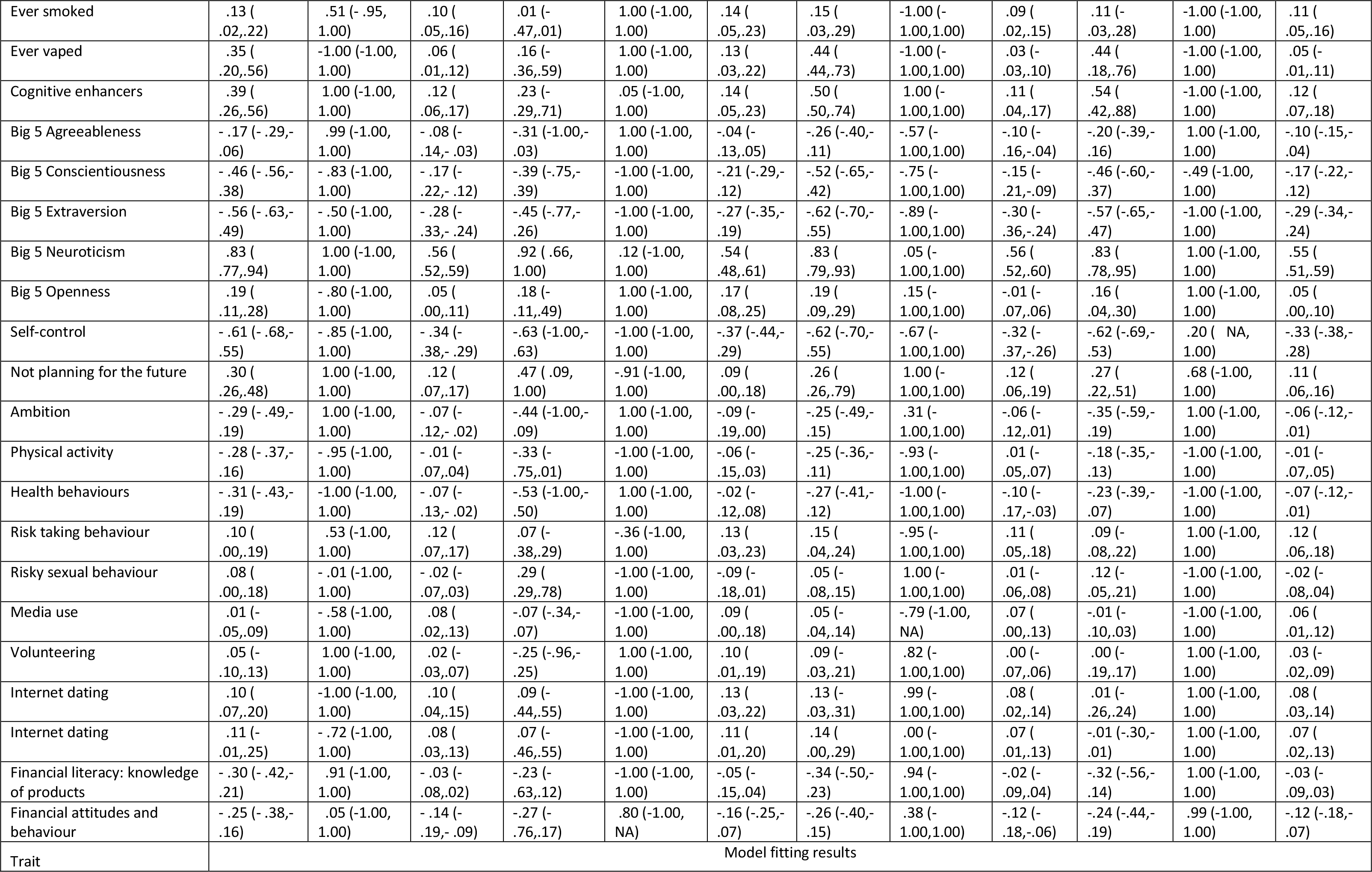
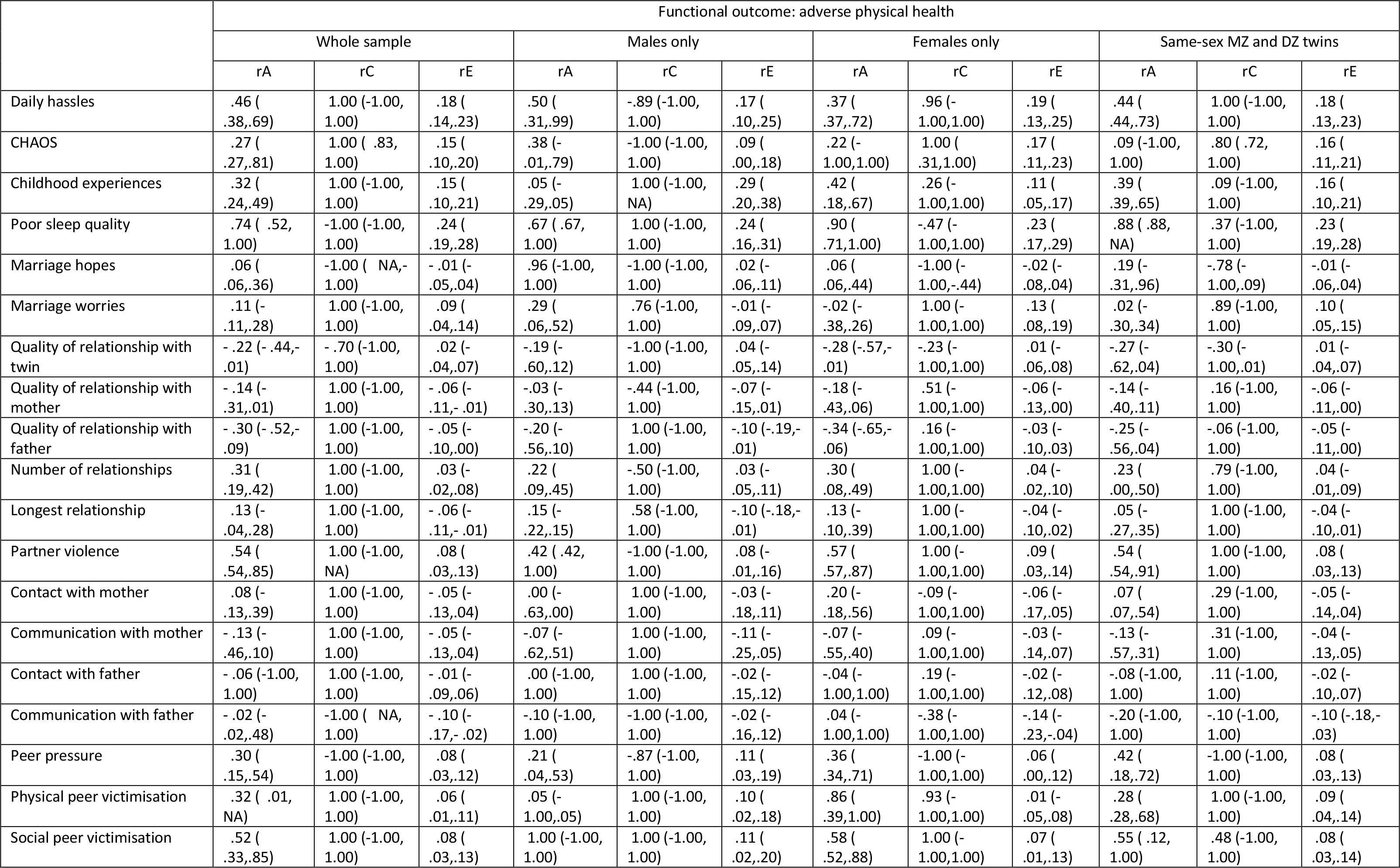
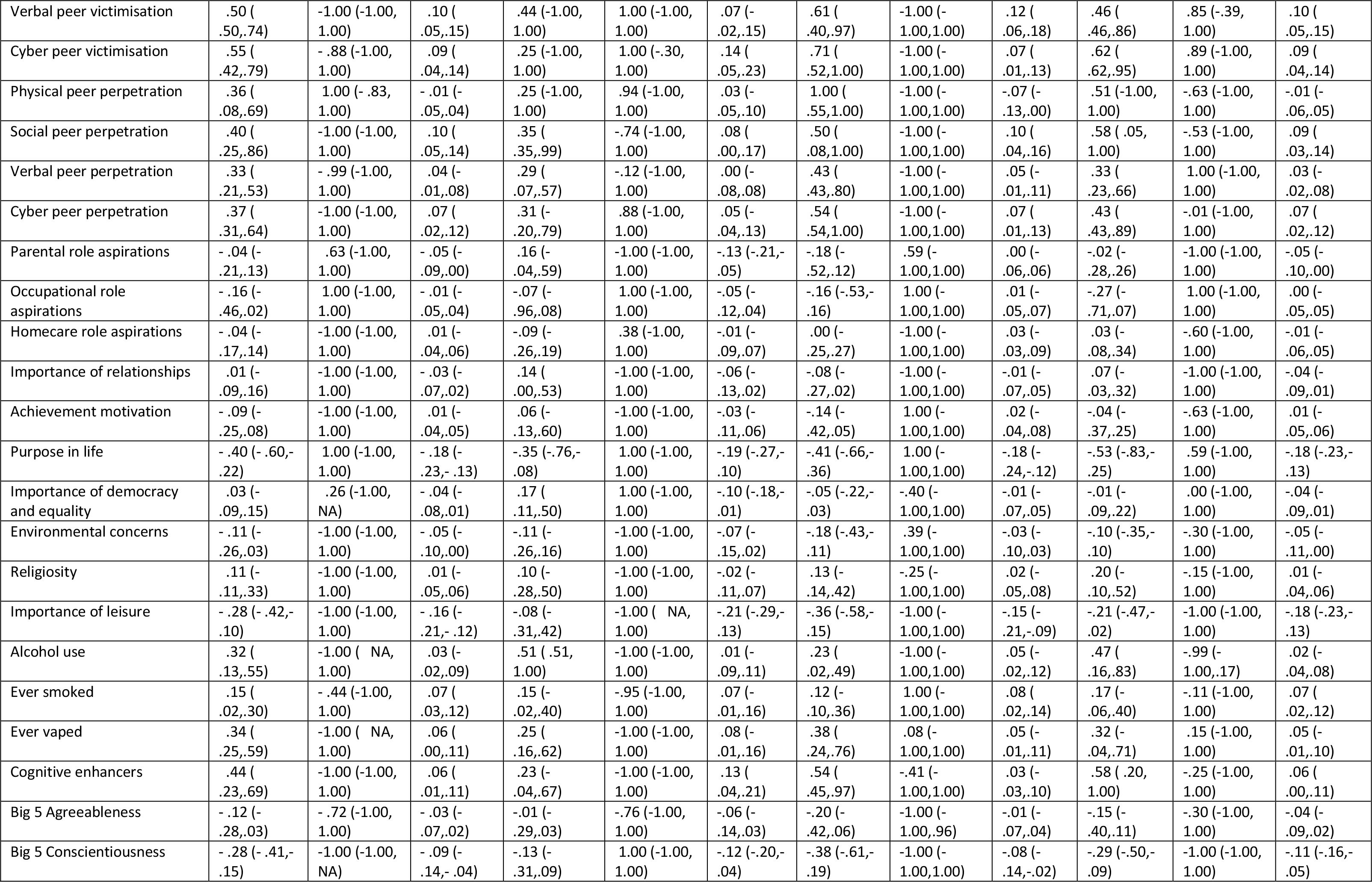
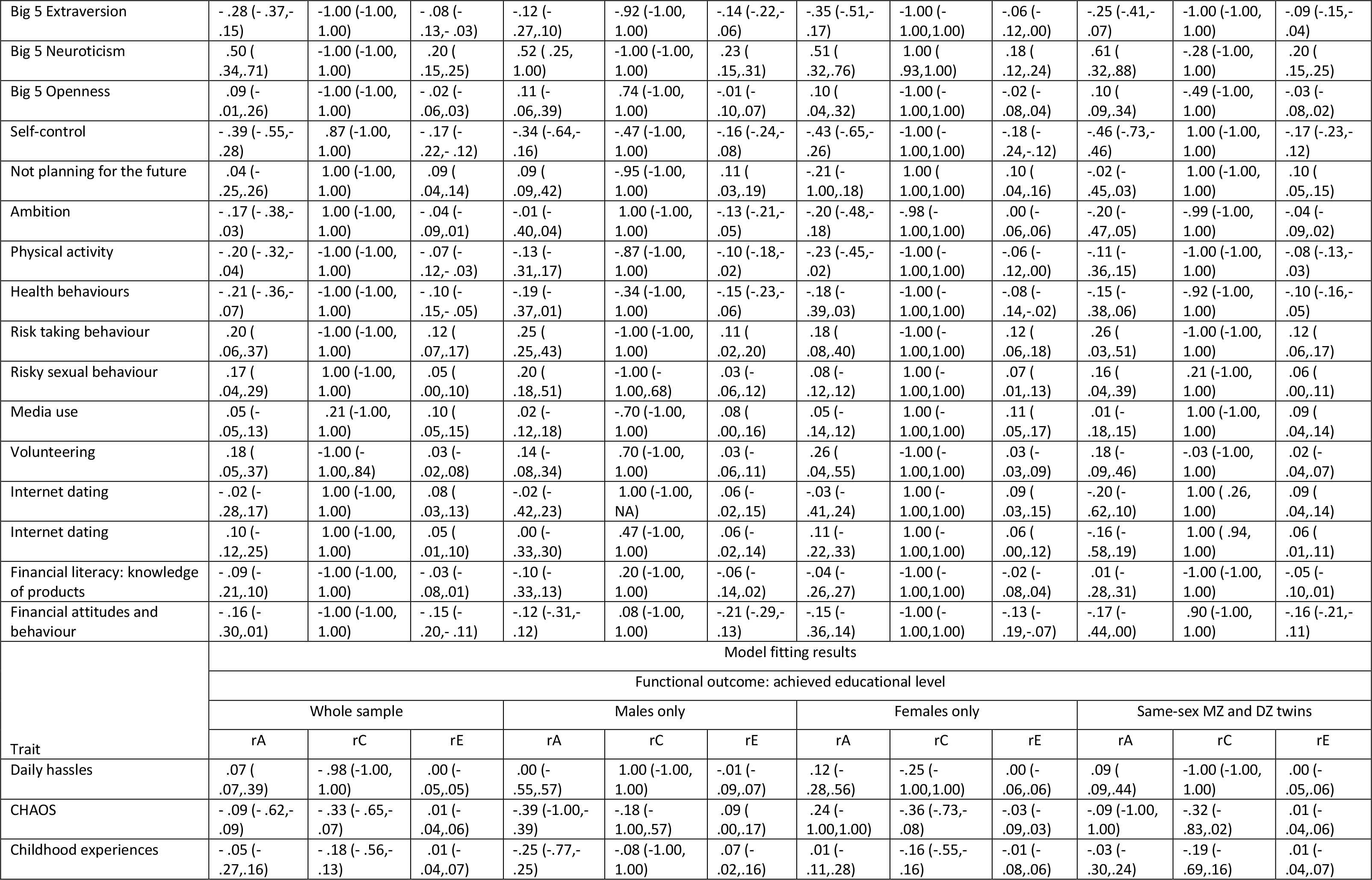
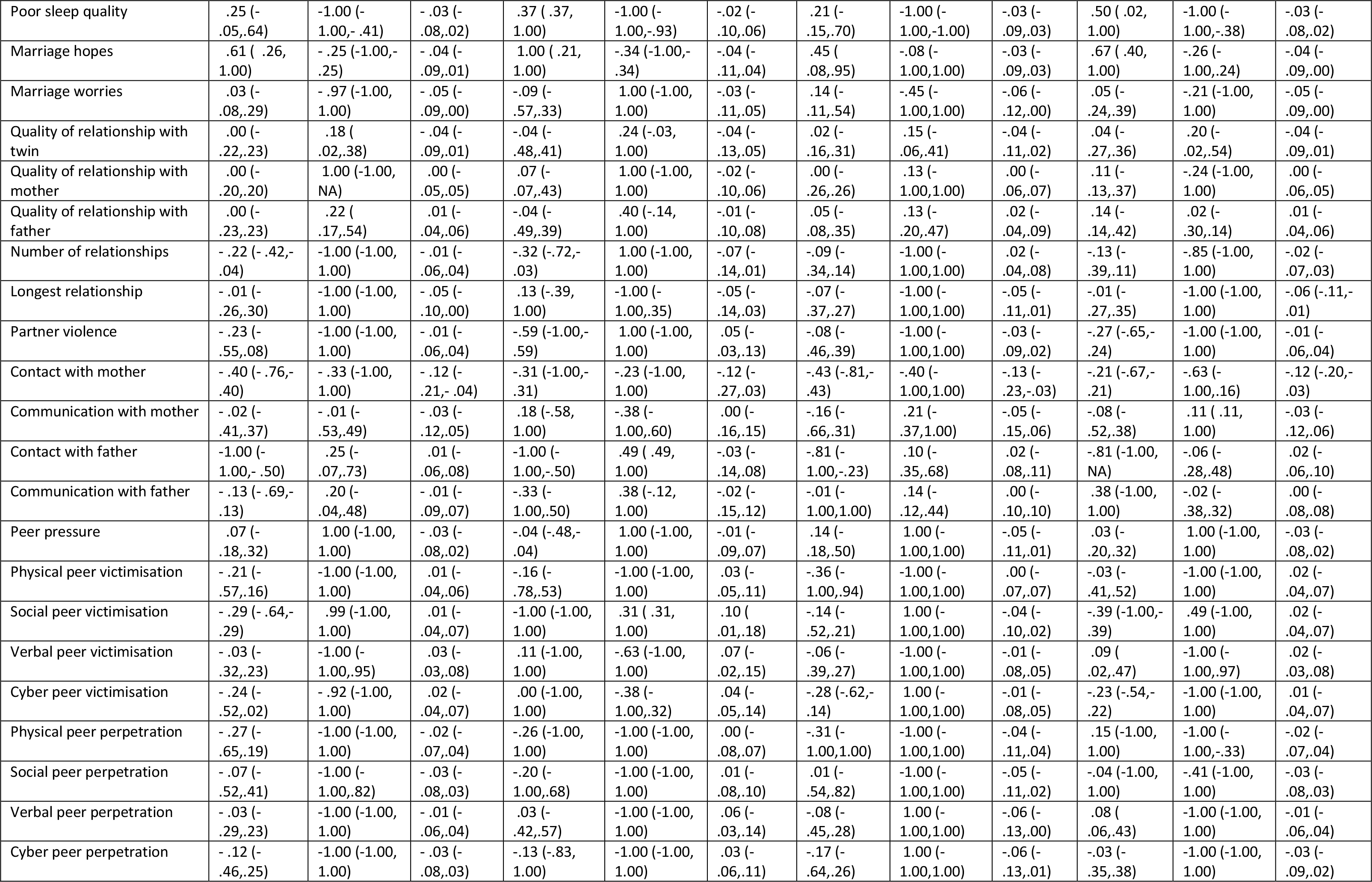
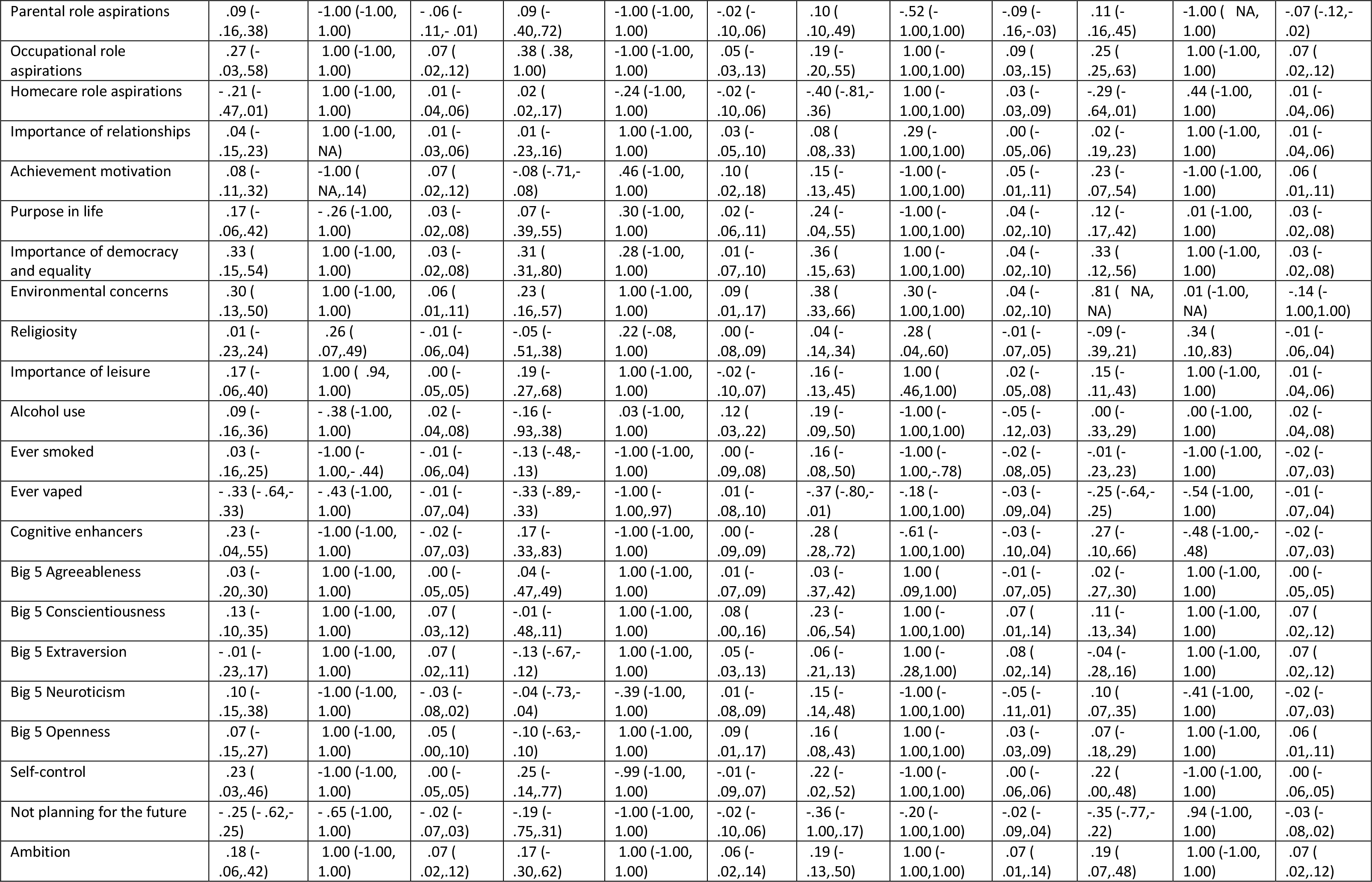
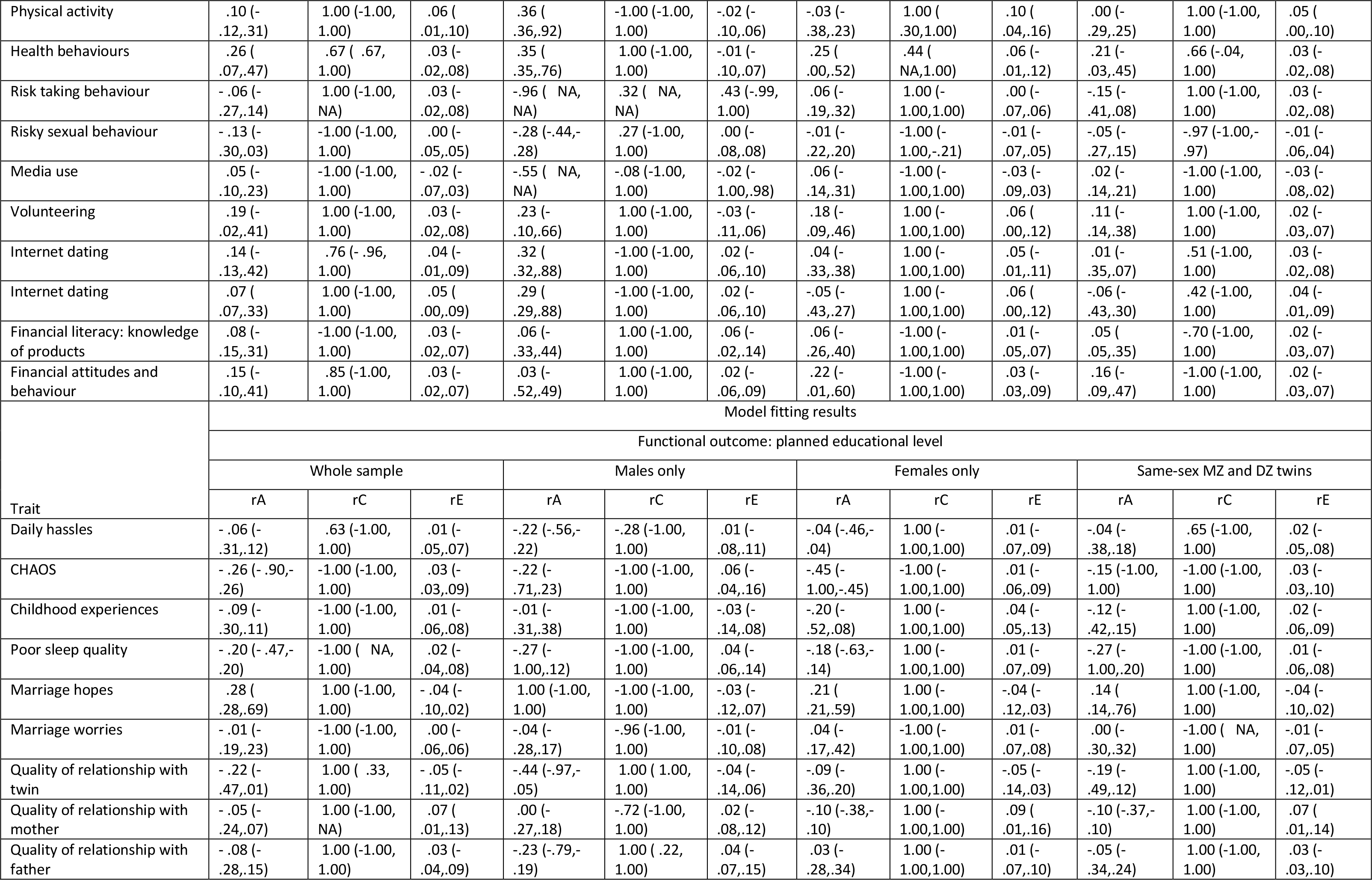
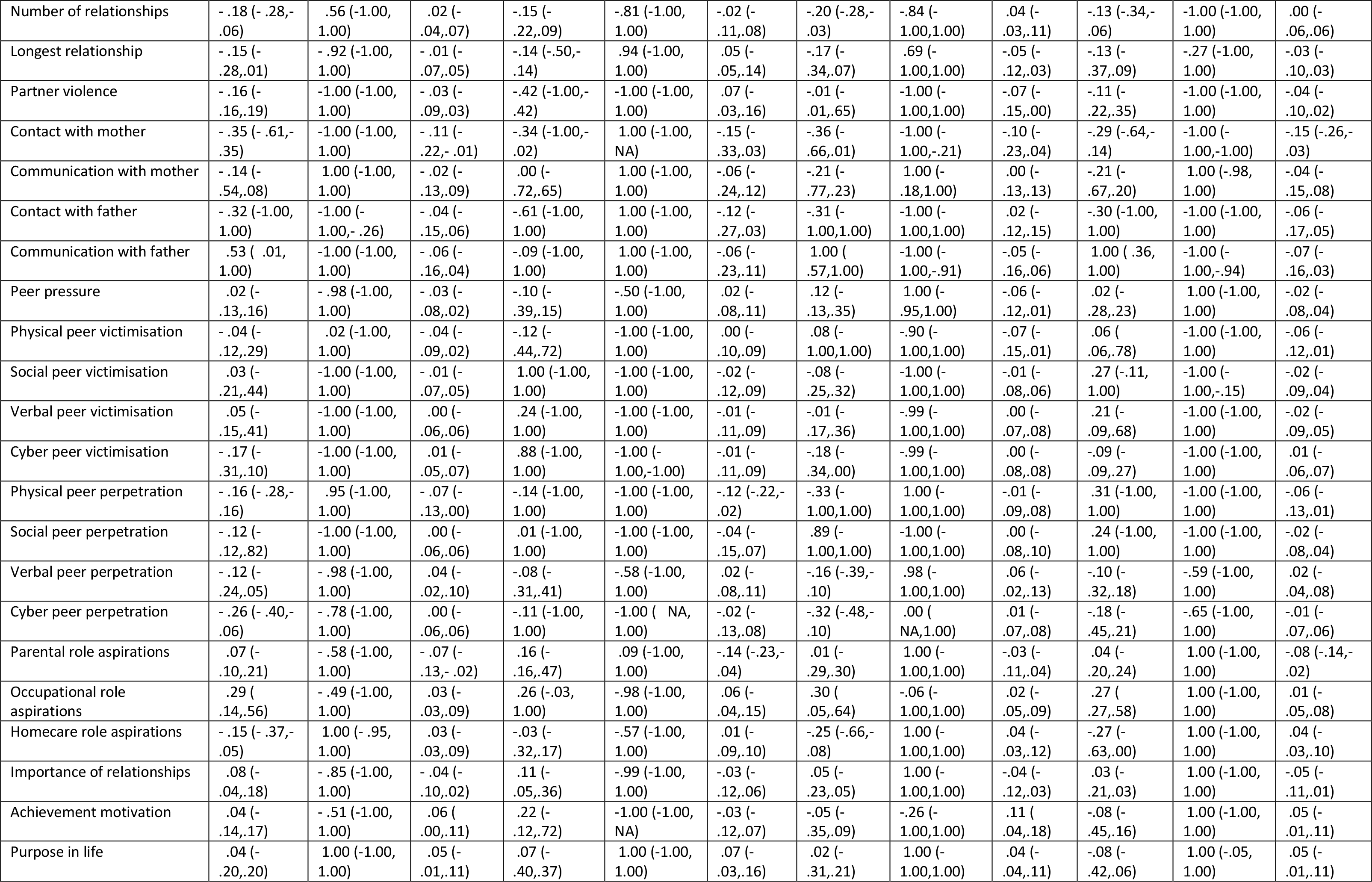
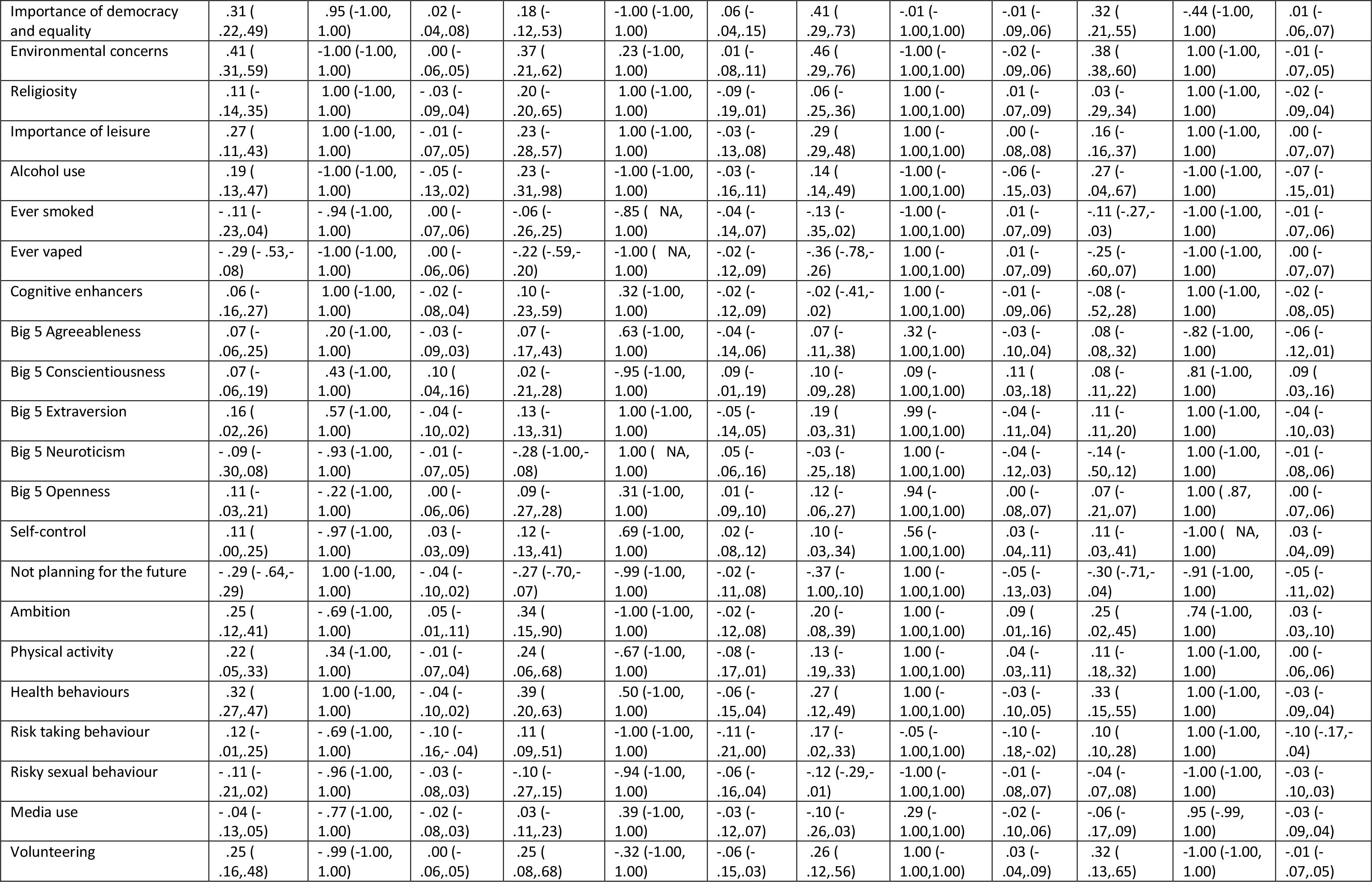
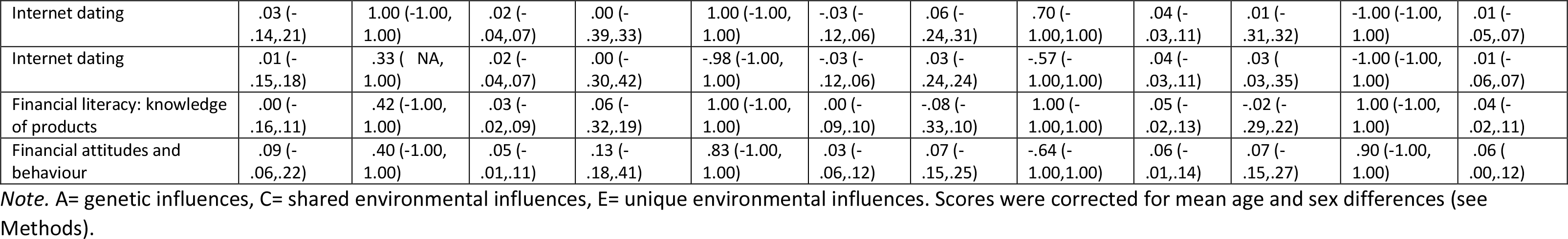
Model fitting results for bivariate analyses of genetic correlation (rA), shared environmental correlation (rC), and non-shared environmental correlation (rE) for psychological traits and functional outcomes (95% confidence intervals are in parentheses).

**Table S14.** Univariate GREML estimates

| Trait | GREML results |  |  |  |  |  |  |  |
| --- | --- | --- | --- | --- | --- | --- | --- | --- |
|  | Family-based |  |  |  | Unrelated samples |  |  |  |
|  | SNP-h<sup>2</sup> | SE | LogL | N | SNP-h<sup>2</sup> | SE | LogL | N |
| Daily hassles | .05 | .07 | -2466.48 | 5115 | .17 | .10 | -1859.61 | 3770 |
| CHAOS | .12 | .07 | -271.92 | 5705 | .20 | .09 | -2026.25 | 4099 |
| Childhood experiences | .15 | .07 | -2372.97 | 5116 | .17 | .10 | -1867.48 | 3777 |
| Poor sleep quality | .13 | .07 | -2388.89 | 5026 | .07 | .10 | -1812.23 | 3713 |
| Marriage hopes | .03 | .06 | -2784.46 | 5622 | .13 | .09 | -2042.02 | 4069 |
| Marriage worries | .10 | .07 | -2798.85 | 5621 | .06 | .09 | -2044.96 | 4068 |
| Quality of relationship with twin | .11 | .08 | -2522.54 | 5533 | .26 | .09 | -2023.64 | 4026 |
| Quality of relationship with mother | .11 | .07 | -2646.08 | 5484 | .09 | .09 | -1991.28 | 3998 |
| Quality of relationship with father | .10 | .07 | -2473.04 | 5381 | .17 | .09 | -1967.84 | 3937 |
| Number of relationships | .14 | .07 | -2671.69 | 5620 | .23 | .09 | -1954.13 | 4069 |
| Longest relationship | .05 | .08 | -2327.97 | 4803 | .05 | .10 | -1803.40 | 3672 |
| Partner violence | .10 | .07 | -270.17 | 5453 | .19 | .09 | -203.17 | 3992 |
| Contact with mother | .10 | .15 | -113.90 | 2393 | .08 | .18 | -976.90 | 2005 |
| Communication with mother | .14 | .14 | -1187.64 | 2448 | .11 | .17 | -1035.21 | 2049 |
| Contact with father | .00 | .14 | -1166.09 | 2614 | .00 | .17 | -1006.91 | 2135 |
| Communication with father | -.08 | .12 | -1206.95 | 2835 | .00 | .15 | -1102.89 | 2313 |
| Peer pressure | .02 | .06 | -281.79 | 5568 | .00 | .09 | -2081.99 | 4045 |
| Physical peer victimisation | .06 | .07 | -249.71 | 4966 | .07 | .10 | -1814.18 | 3691 |
| Social peer victimisation | .17 | .07 | -2462.50 | 4966 | .16 | .10 | -1836.95 | 3691 |
| Verbal peer victimisation | .14 | .07 | -2556.27 | 4964 | .10 | .10 | -1927.58 | 3690 |
| Cyber peer victimisation | .16 | .07 | -2488.27 | 4963 | .18 | .10 | -1932.56 | 3689 |
| Physical peer perpetration | .00 | .07 | -2533.76 | 4965 | .04 | .10 | -1957.28 | 3690 |
| Social peer perpetration | .12 | .07 | -2313.53 | 4964 | .09 | .10 | -1661.46 | 3689 |
| Verbal peer perpetration | .00 | .07 | -2566.01 | 4964 | .00 | .10 | -1884.04 | 3689 |
| Cyber peer perpetration | .16 | .08 | -253.70 | 4964 | .25 | .10 | -199.04 | 3689 |
| Parental role aspirations | .04 | .07 | -2651.00 | 5121 | .00 | .09 | -1913.54 | 3781 |
| Occupational role aspirations | .00 | .07 | -2545.07 | 5124 | .00 | .10 | -184.20 | 3782 |
| Homecare role aspirations | .10 | .07 | -253.68 | 5124 | .13 | .10 | -1841.62 | 3782 |
| Importance of relationships | .06 | .06 | -2795.22 | 5670 | .08 | .09 | -2006.33 | 4112 |
| Achievement motivation | .04 | .07 | -2838.80 | 5669 | .12 | .09 | -2037.22 | 4111 |
| Purpose in life | .09 | .07 | -2781.45 | 5667 | .14 | .09 | -207.67 | 4110 |
| Importance of democracy and equality | .05 | .07 | -2813.41 | 5657 | .07 | .09 | -2046.04 | 4105 |
| Environmental concerns | .19 | .07 | -2788.12 | 5663 | .24 | .09 | -2054.45 | 4109 |
| Religiosity | .00 | .06 | -215.37 | 5571 | .00 | .09 | -1688.22 | 4047 |
| Importance of leisure | .20 | .07 | -2463.74 | 5026 | .26 | .10 | -1842.84 | 3714 |
| Alcohol use | .01 | .08 | -214.11 | 4635 | .05 | .10 | -1616.28 | 3503 |
| Ever smoked | .04 | .07 | -2466.61 | 5043 | .00 | .09 | -1842.05 | 3728 |
| Ever vaped | .09 | .07 | -2427.37 | 4985 | .07 | .10 | -1914.77 | 3730 |
| Cognitive enhancers | .05 | .07 | -275.32 | 5508 | .08 | .10 | -184.88 | 3703 |
| Big 5 Agreeableness | .12 | .08 | -2521.59 | 5040 | .00 | .09 | -1998.17 | 4043 |
| Big 5 Conscientiousness | .00 | .06 | -2699.13 | 5500 | .01 | .09 | -197.73 | 4038 |
| Big 5 Extraversion | .13 | .07 | -2709.25 | 5528 | .21 | .09 | -1934.41 | 4059 |
| Big 5 Neuroticism | .06 | .07 | -2702.47 | 5534 | .05 | .09 | -2005.99 | 4063 |
| Big 5 Openness | .16 | .07 | -2723.37 | 5519 | .25 | .09 | -1985.46 | 4053 |
| Self-control | .02 | .06 | -2716.50 | 5498 | .00 | .08 | -1944.17 | 4037 |
| Not planning for the future | .06 | .07 | -2678.72 | 5493 | .19 | .09 | -1946.78 | 4033 |
| Ambition | .03 | .07 | -2572.27 | 5117 | .11 | .10 | -1878.06 | 3771 |
| Physical activity | .04 | .07 | -2754.09 | 5548 | .00 | .09 | -2033.70 | 4031 |
| Health behaviours | .12 | .07 | -2734.04 | 5571 | .11 | .09 | -2047.28 | 4039 |
| Risk taking behaviour | .05 | .07 | -2624.36 | 5493 | .08 | .09 | -1981.79 | 4034 |
| Risky sexual behaviour | .17 | .07 | -2606.46 | 5397 | .22 | .09 | -1963.86 | 3955 |
| Media use | .13 | .07 | -2625.49 | 5526 | .04 | .09 | -1957.01 | 4022 |
| Volunteering | .06 | .07 | -2737.51 | 5577 | .17 | .09 | -2003.49 | 4050 |
| Internet dating (scale) | .03 | .07 | -2711.68 | 5506 | .00 | .09 | -1984.05 | 4013 |
| Internet dating (y/n) | .05 | .07 | -2755.54 | 5524 | .00 | .09 | -2009.21 | 4020 |
| Financial literacy: knowledge of products | .15 | .07 | -2647.35 | 5440 | .13 | .09 | -1994.02 | 3988 |
| Financial attitudes and behaviour | .06 | .07 | -2695.96 | 5439 | .00 | .09 | -1961.40 | 3987 |
| Functional outcome | Family-based |  |  |  | Unrelated samples |  |  |  |
|  | SNP-h<sup>2</sup> | SE | SNP-h<sup>2</sup> | SE | SNP-h<sup>2</sup> | SE | SNP-h<sup>2</sup> | SE |
| Wellbeing factor (1st PC) | .04 | .13 | -1359.09 | 2703 | .04 | .13 | -1358.65 | 2701 |
| P factor (1st PC) | .20 | .17 | -1097.11 | 2142 | .20 | .17 | -1096.09 | 2139 |
| Health factor (1st PC) | .00 | .16 | -1232.74 | 2292 | .00 | .16 | -1232.50 | 2289 |
| Achieved educational level | .01 | .07 | -2607.07 | 5546 | .09 | .09 | -1979.12 | 4028 |
| Planned educational level | .14 | .10 | -1642.97 | 3641 | .24 | .12 | -1409.25 | 2956 |

Note. SE= standard error; LogL= log likelihood; N= sample size.

**Table S15.**
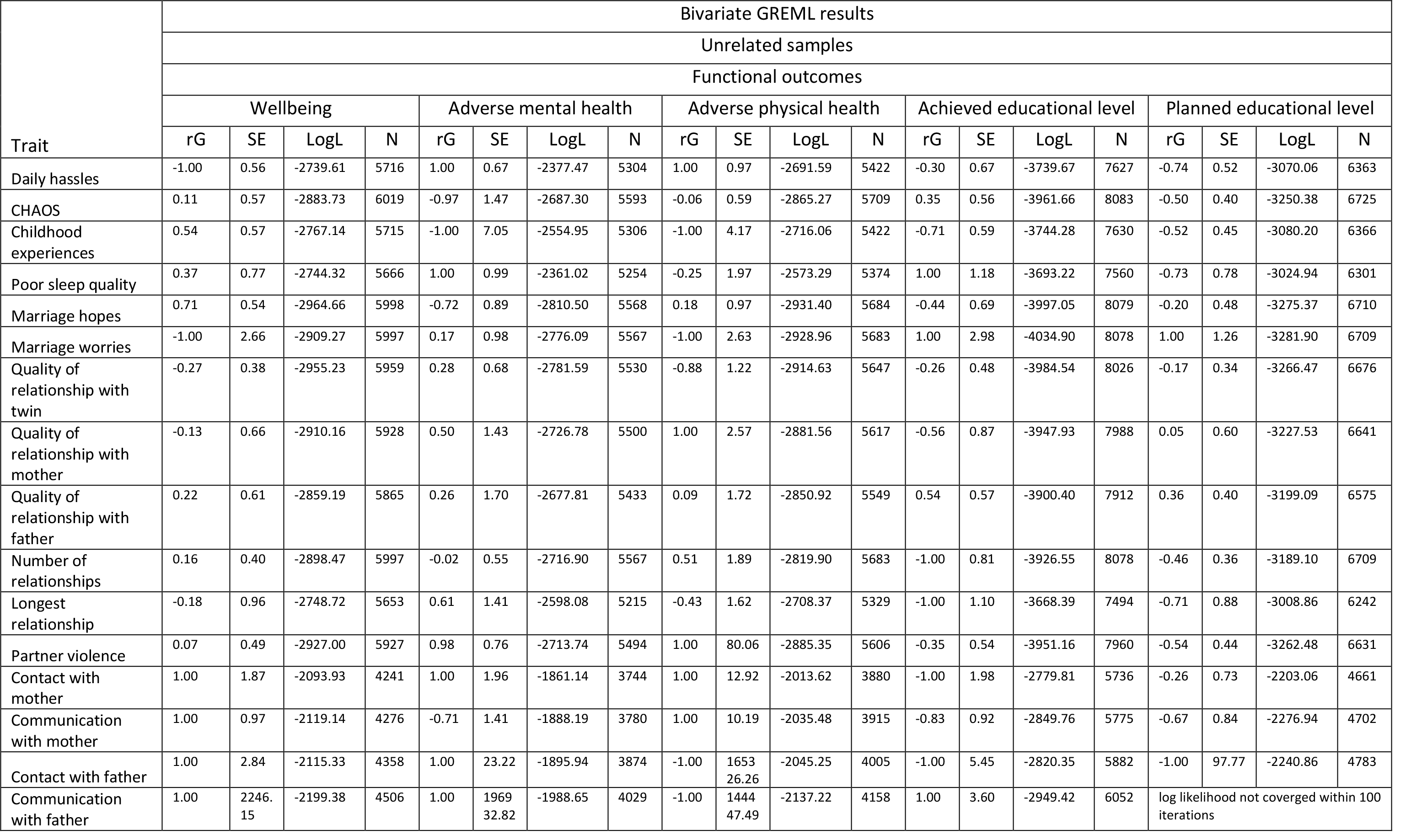
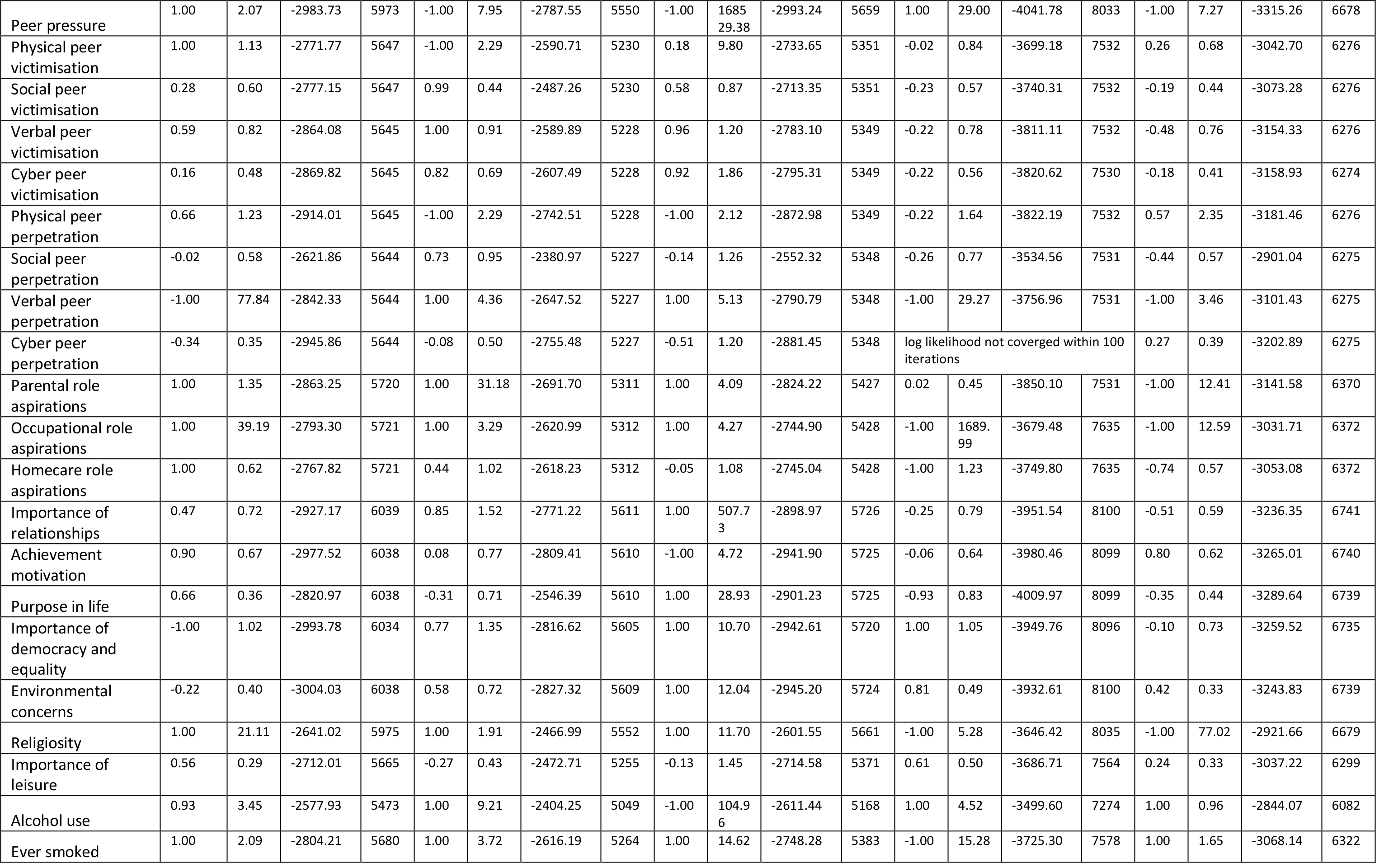
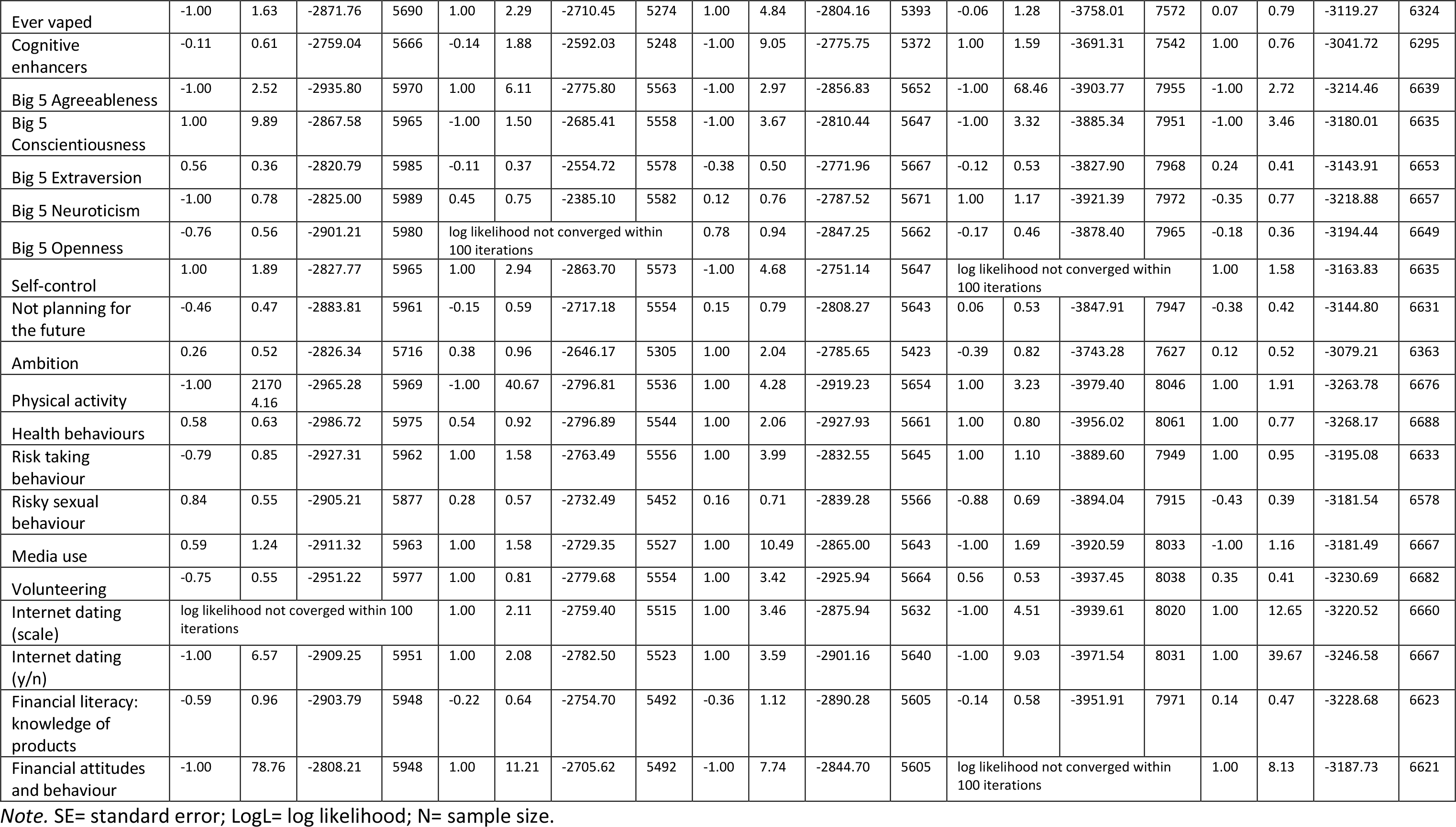
Bivariate GREML results.

#### Supplementary Figures

**Figure S1.**
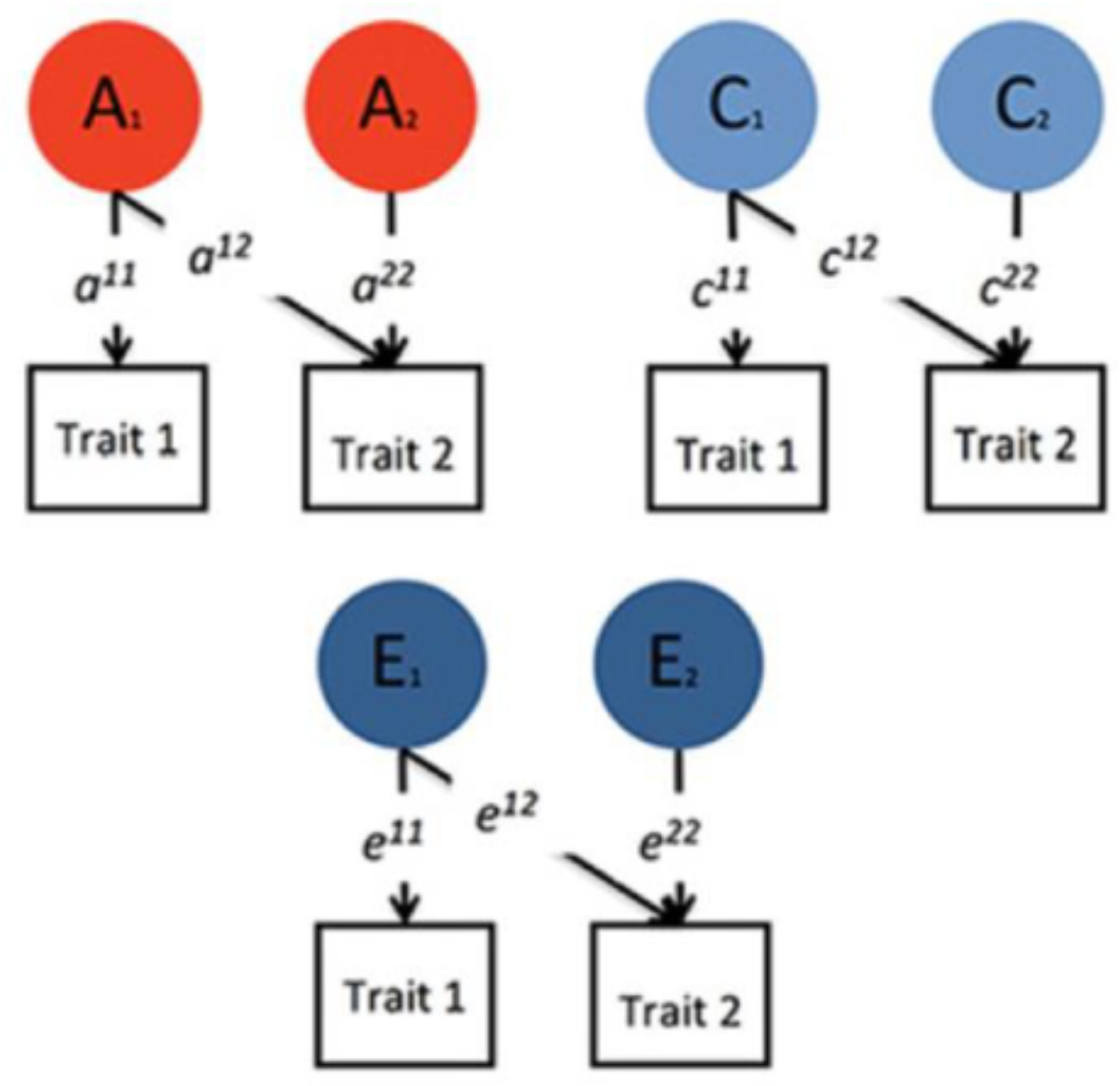
The Bivariate Cholesky Decomposition.

**Figure S2.**
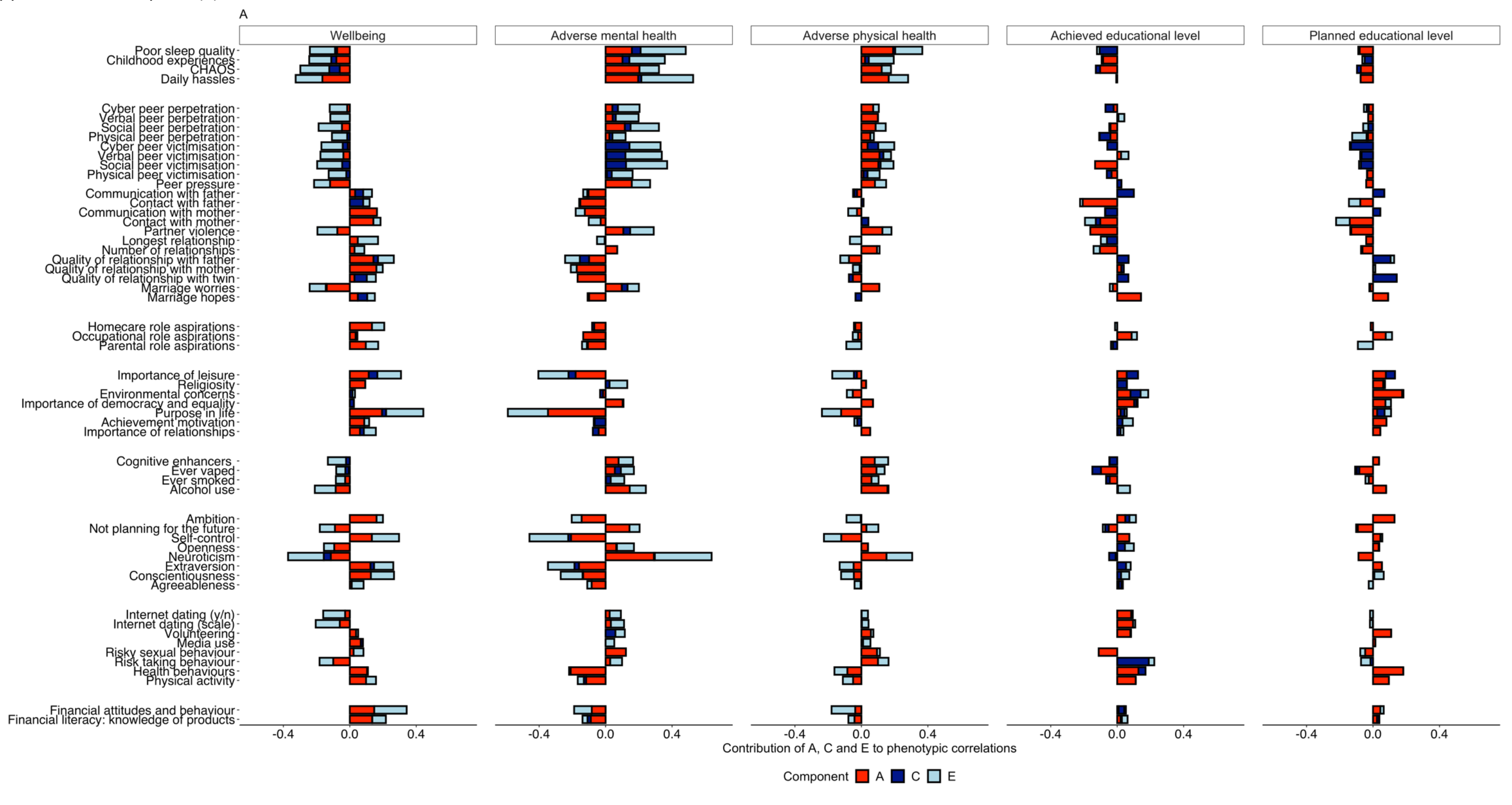
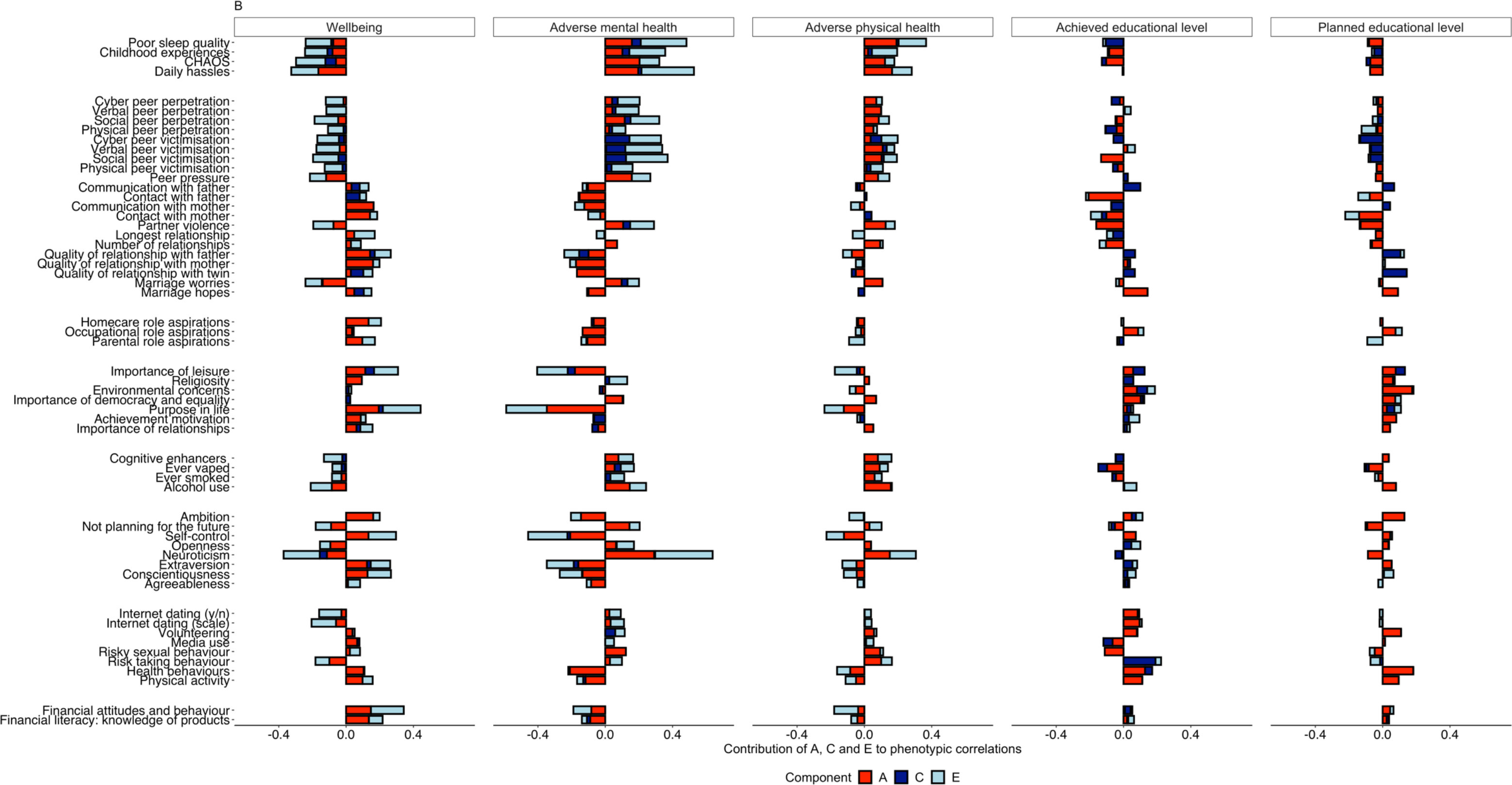
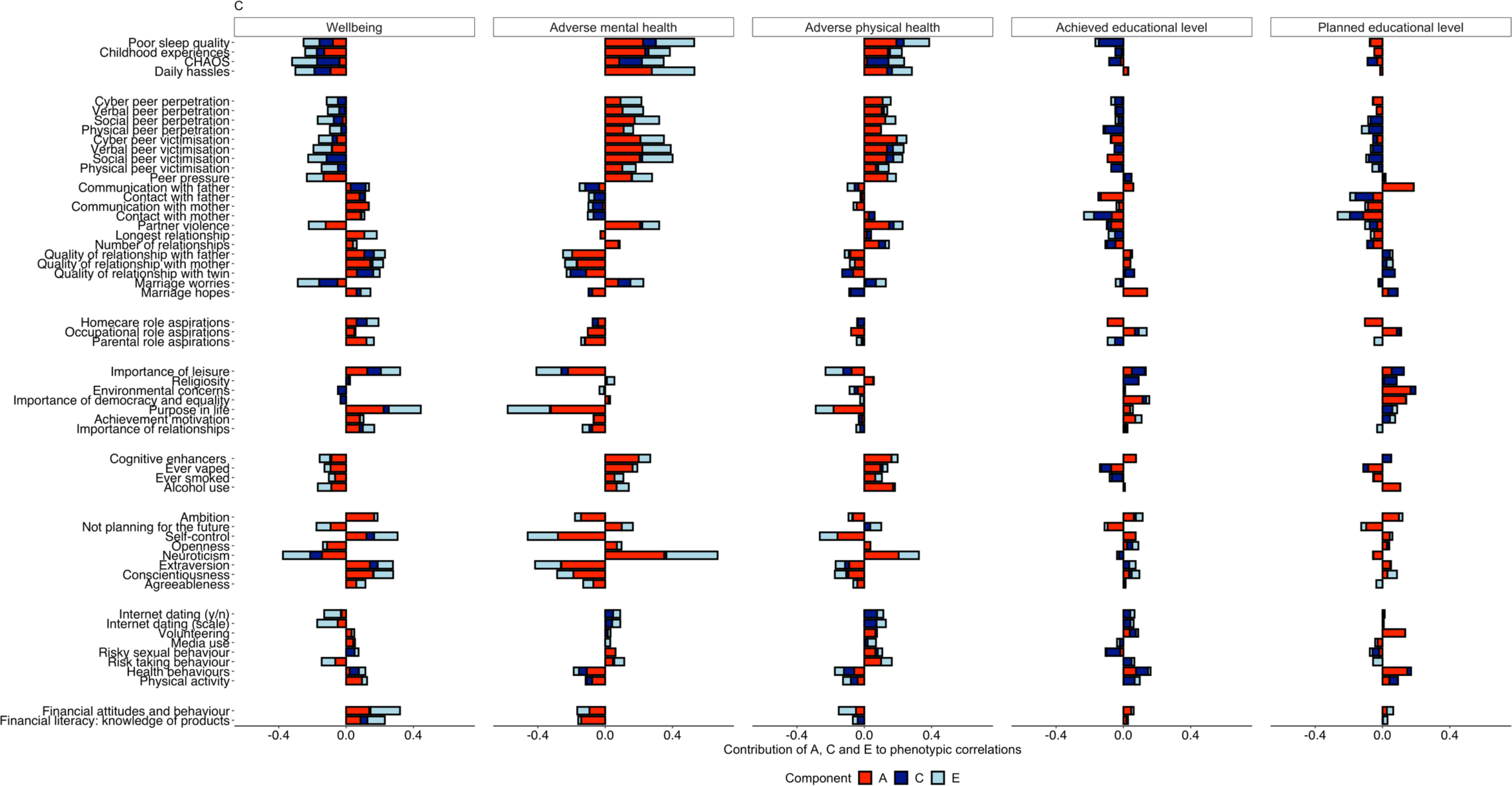
Phenotypic correlations between key outcomes and emerging adulthood variables (indicated by the total length of the bar); bivariate genetic estimates for additive genetic (A), shared environmental (C) and non-shared environmental (E) contributions to these correlations for males (a), females (b) and when excluding DZ opposite sex twin pairs (c).

**Figure S3.**
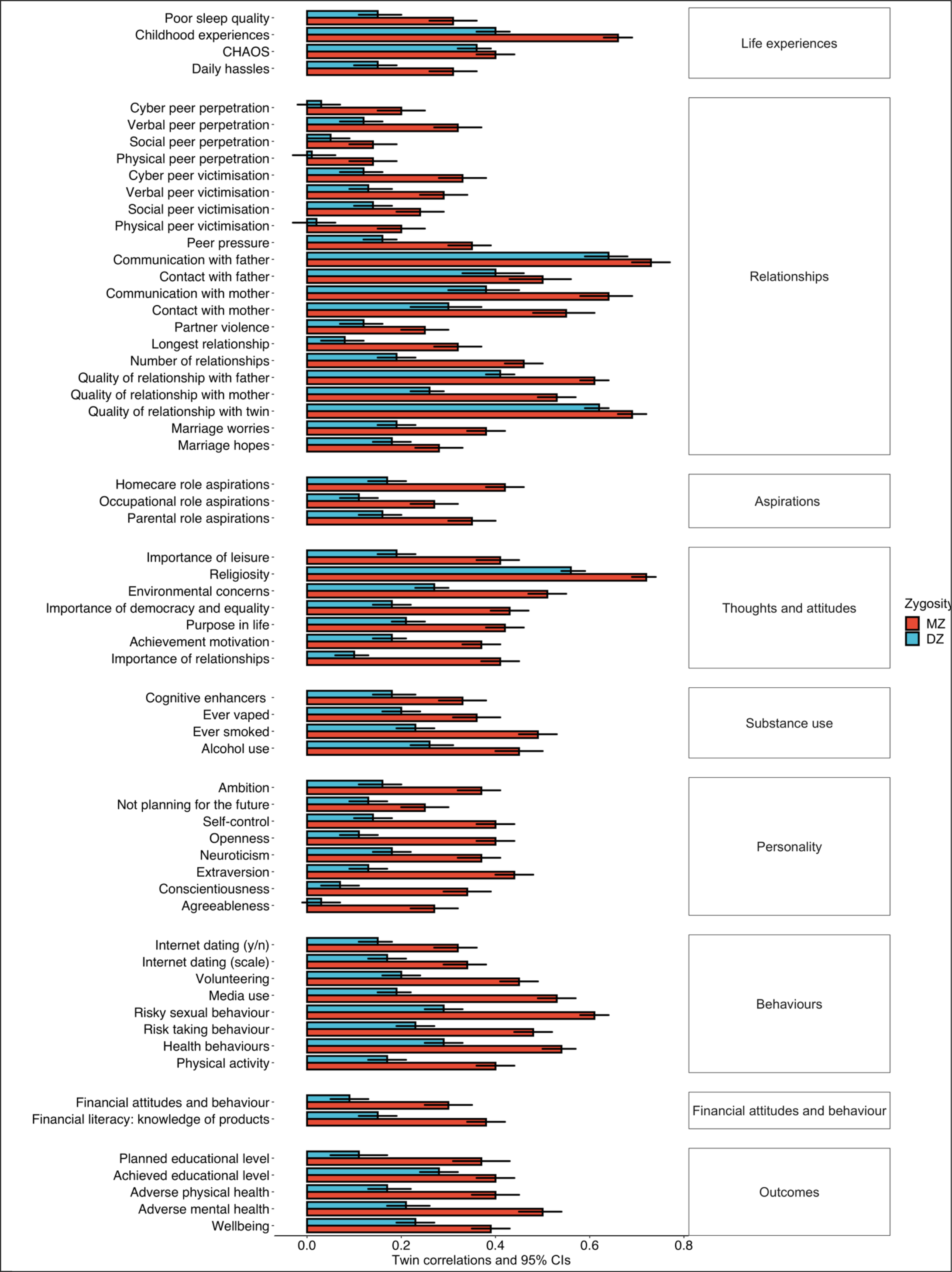
Twin correlations for psychological and behavioural traits and key functional outcomes for the whole sample.

**Figure S4.**
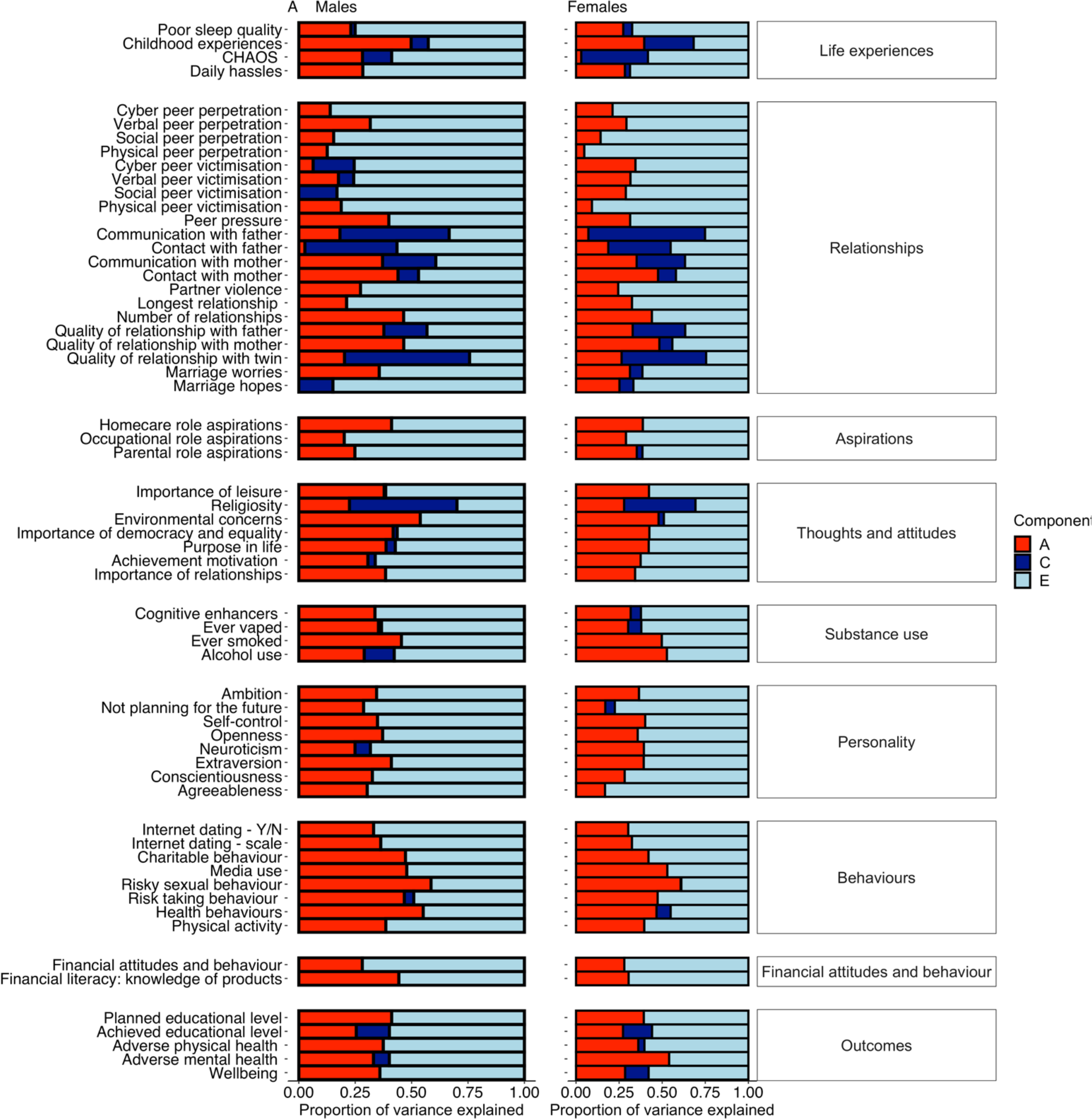
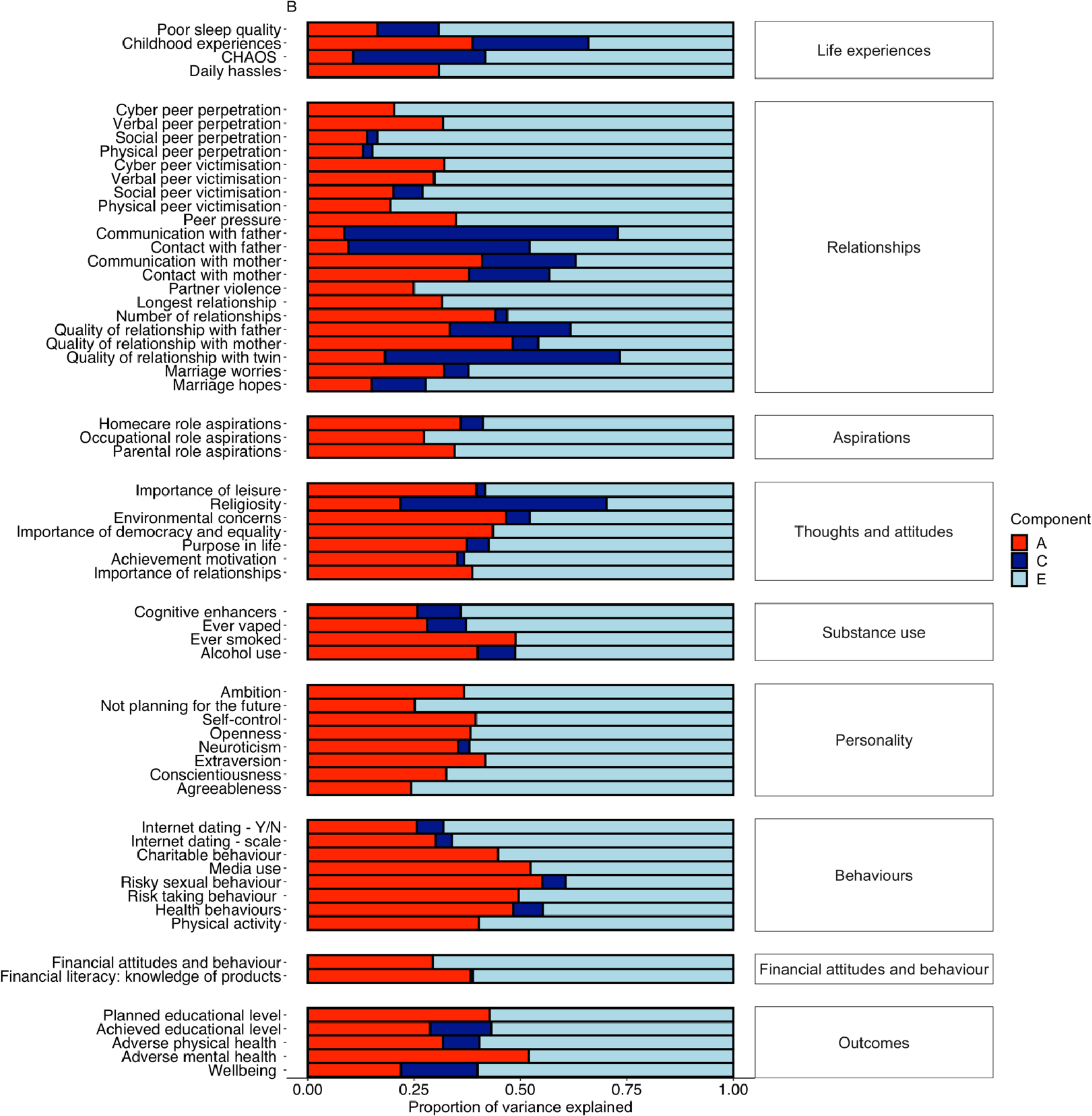
Univariate analyses of additive genetic (A), shared environmental (C), and non-shared environmental (E) components of variance for variables for psychological traits and functional outcomes for when calculated for males and females separately (a) and for the whole sample when opposite sex DZ twin pairs were excluded (b).

**Figure S5.**
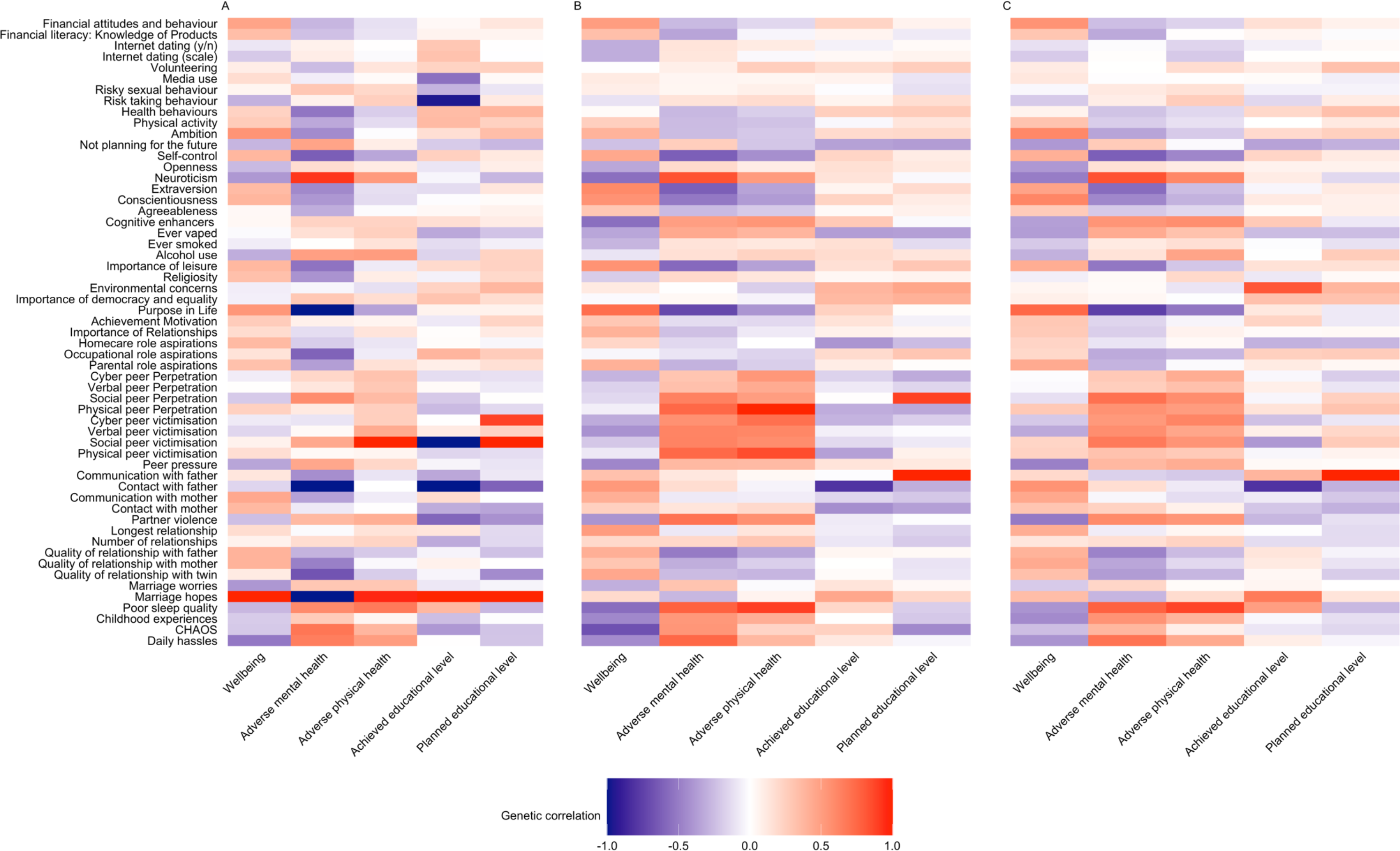
Genetic correlations between psychological and behavioural traits and key functional outcomes for the males (a), females (b) and for the whole sample when opposite sex DZ twin were excluded (c)

**Figure S6.**
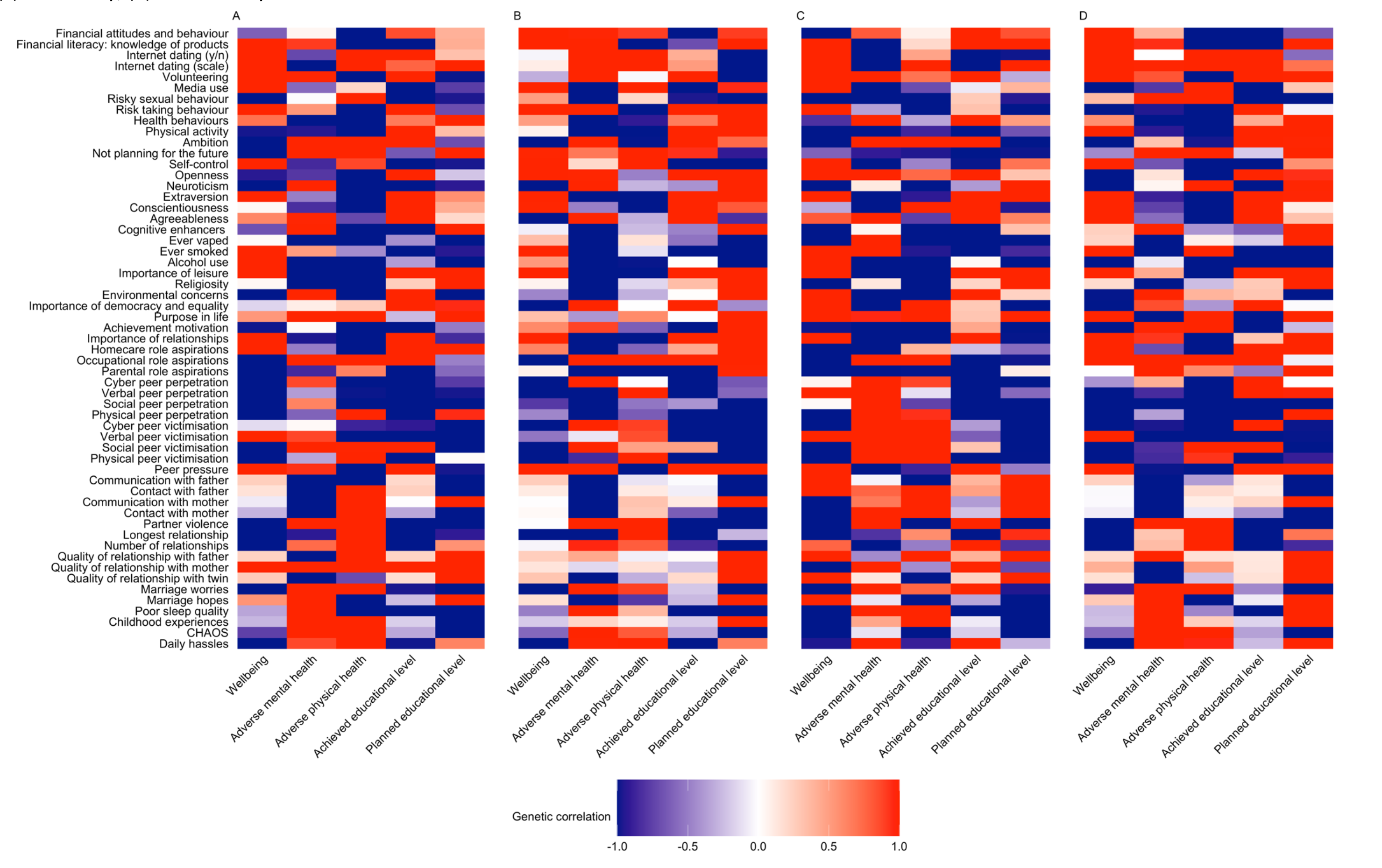
Shared environmental correlations between psychological and behavioural traits and key functional outcomes (a) for the whole sample; (b) MZ and DZ same sex; (c) males only; (d) females only.

**Figure S7.**
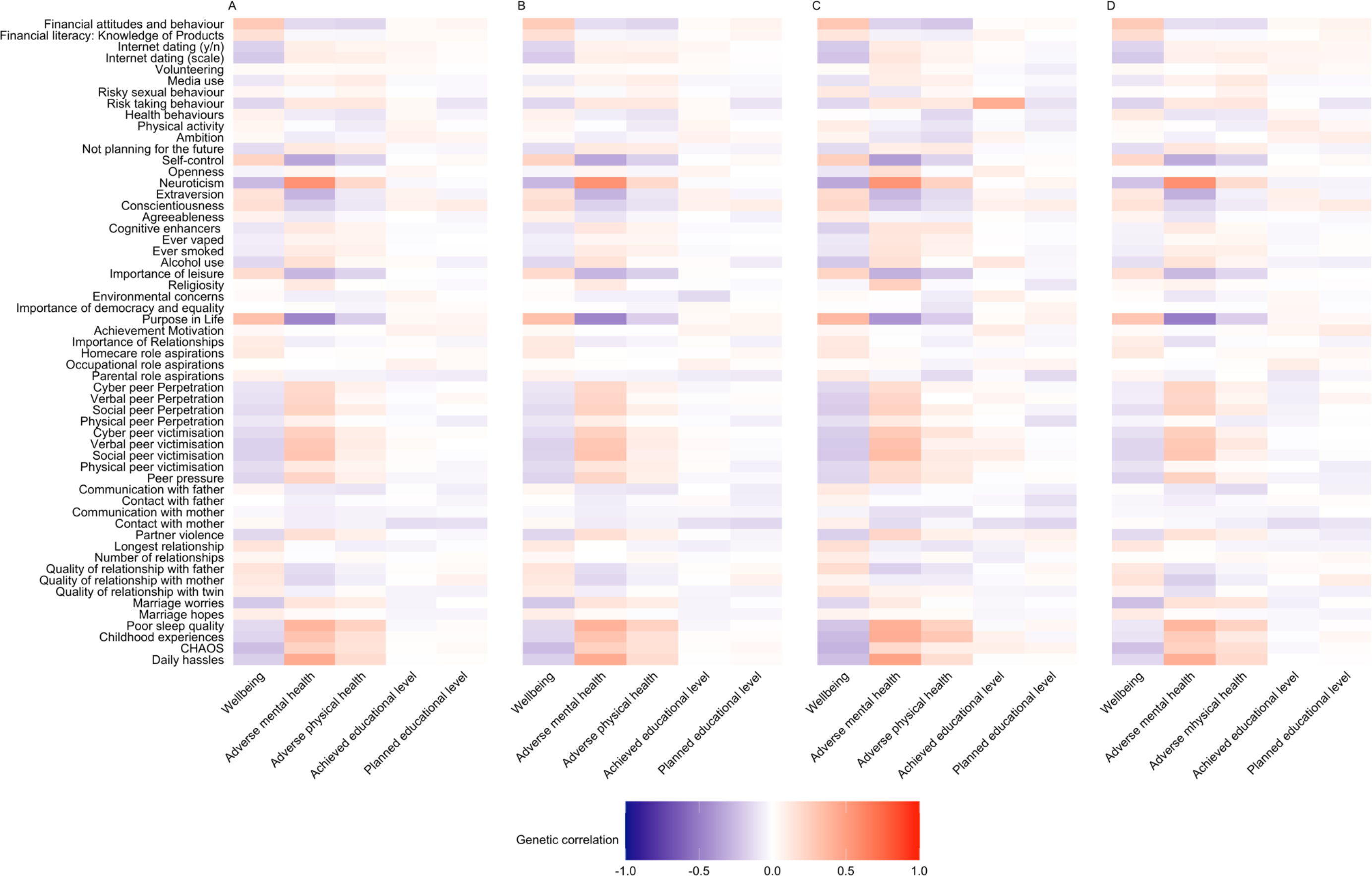
Non-shared environmental correlations between psychological and behavioural traits and key functional outcomes (a) for the whole sample; (b) MZ and DZ same sex twin pairs only; (c) males only; (d) females only

**Figure S8.**
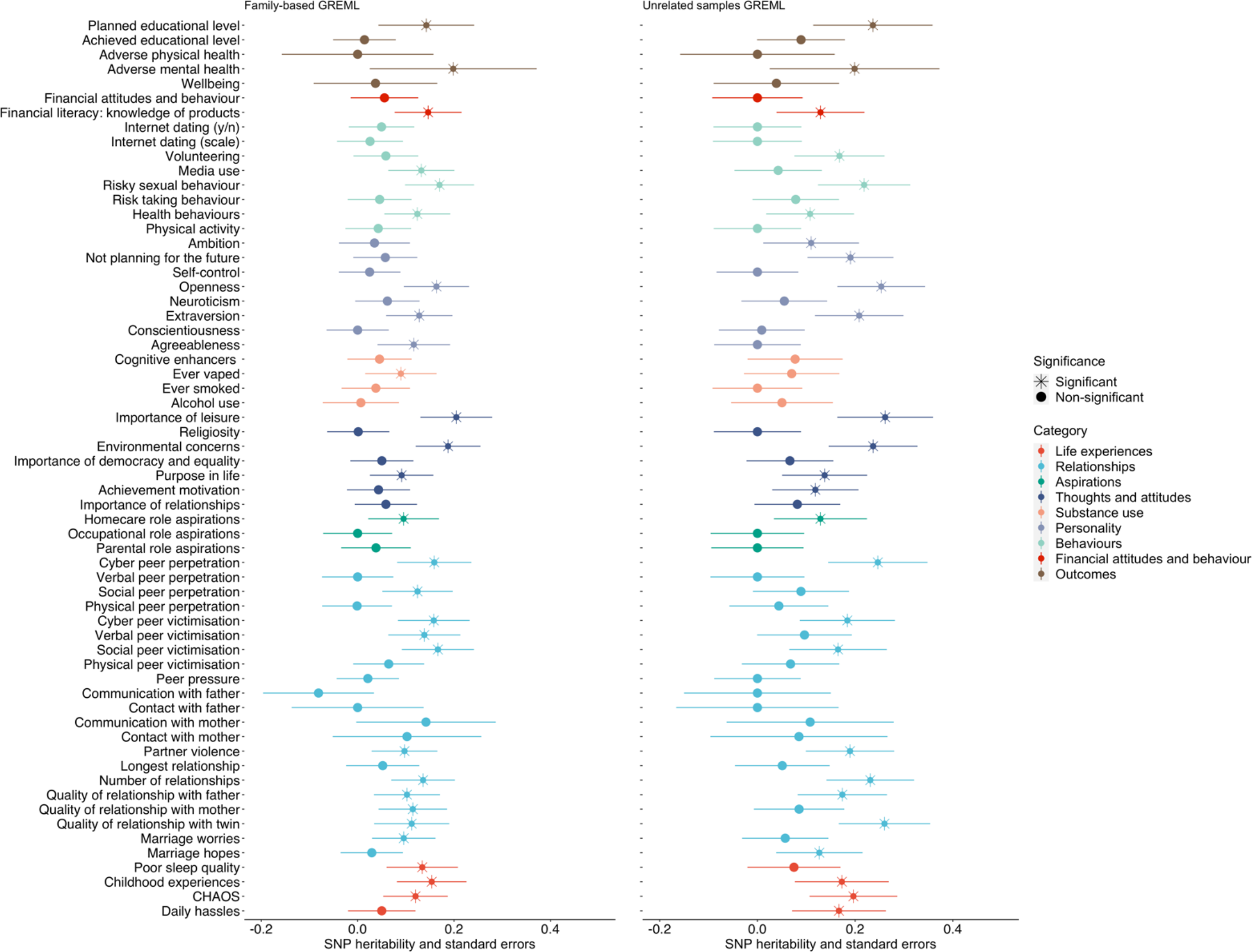
Univariate GREML analyses; SNP heritabilities for psychological traits and functional outcomes for the whole sample (standard errors represented in error bars; reml-no-constrain option indicated by the point estimates below 0).

**Figure S9.**
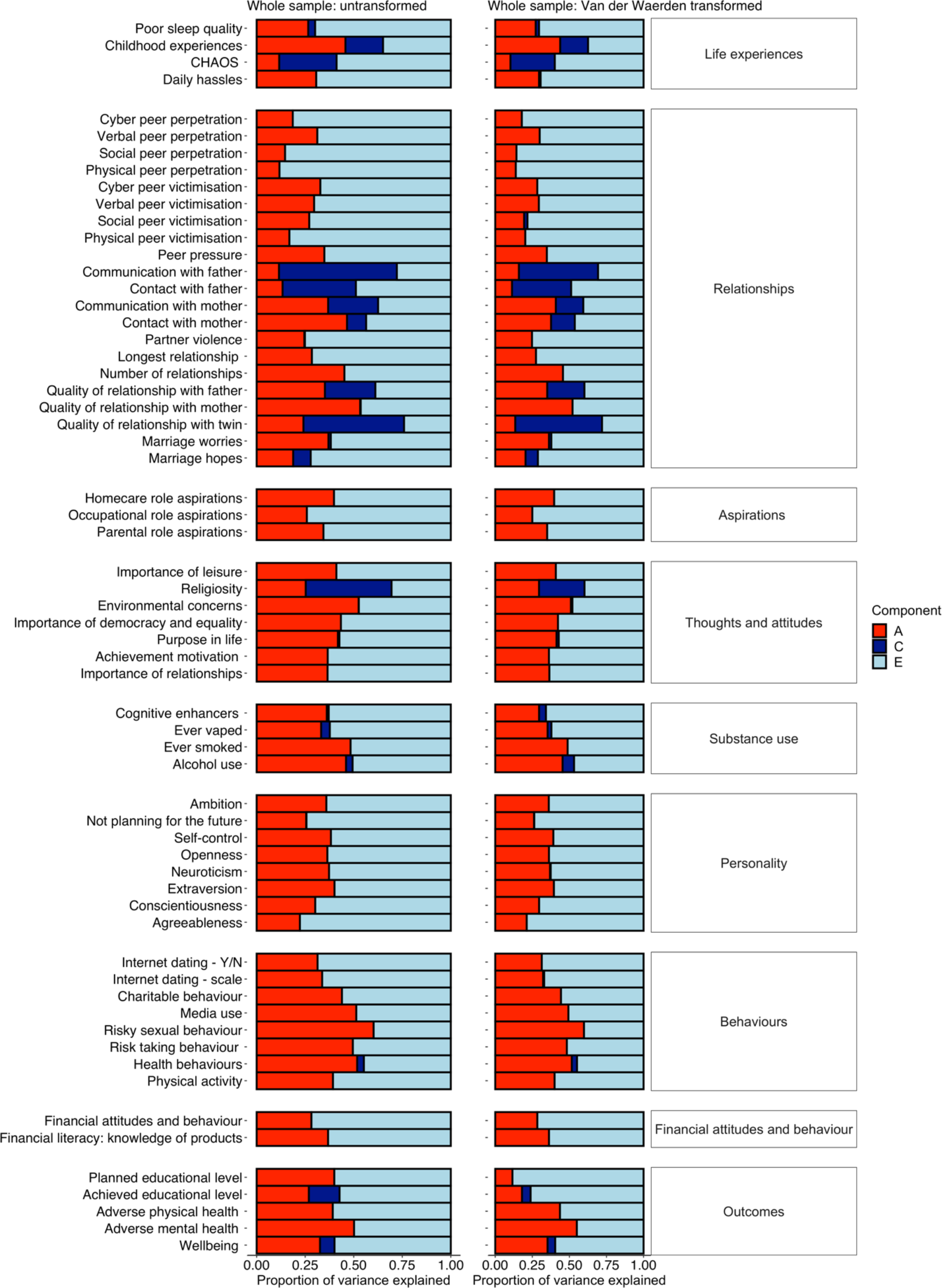
Univariate twin analyses of additive genetic (A), shared environmental (C), and non-shared environmental (E) components of variance for variables for psychological traits and functional outcomes for the whole sample for untransformed data and when using Van der Waerden transformation

**Figure S10.**
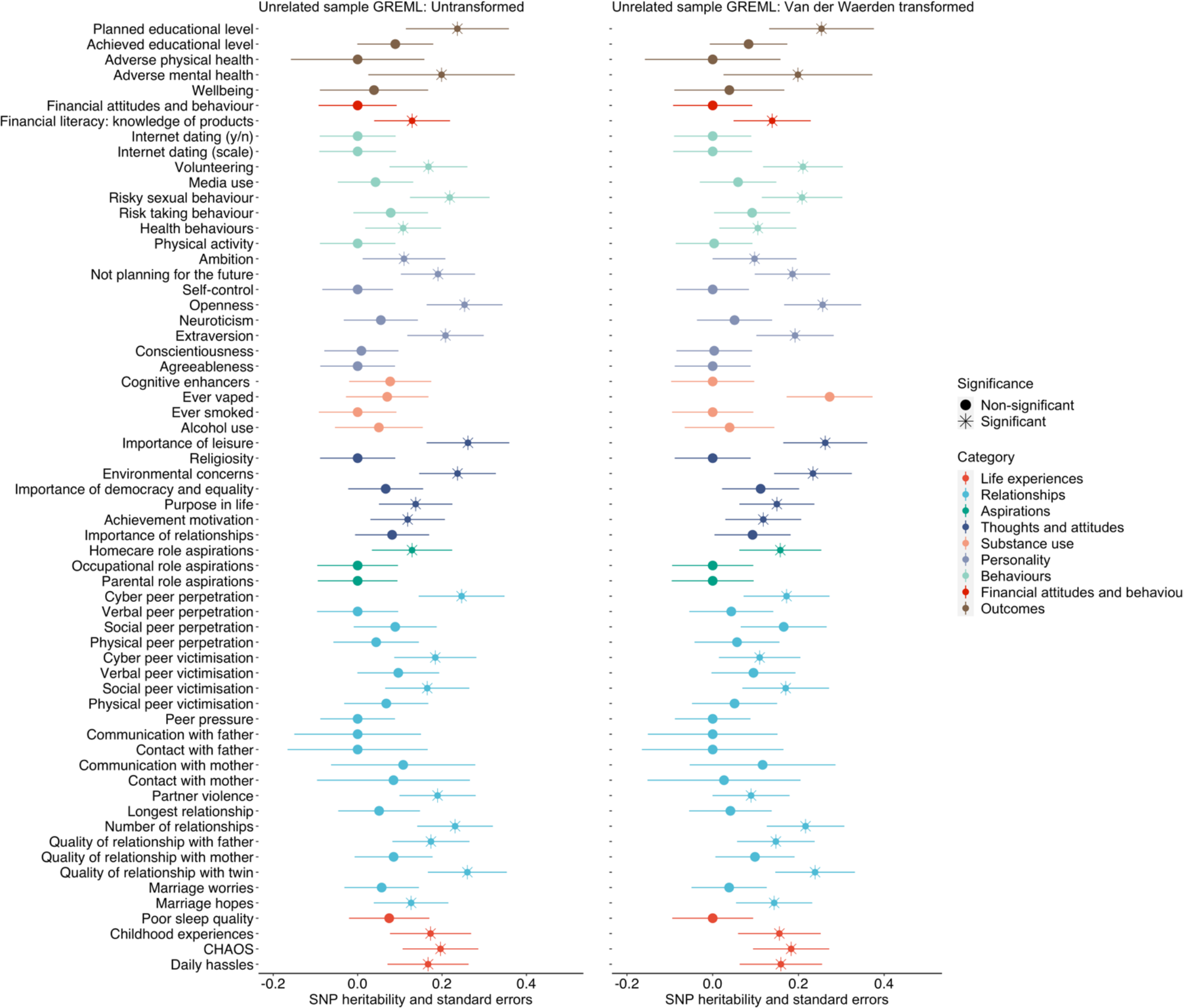
Univariate GREML analyses; SNP heritabilities for psychological traits and functional outcomes for the whole sample for untransformed data and when using Van der Waerden transformation (standard errors represented in error bars).

## References

1. Arnett, J. J. (2004). Emerging adulthood: The winding road from the late teens through the twenties. In Emerging Adulthood: The Winding Road from the Late Teens through the Twenties. https://doi.org/10.1093/acprof:oso/9780195309379.001.0001

2. Arnett, J. J. (2015). Emerging adulthood: The winding road from the kate teens through the twenties. In Emerging Adulthood: The Winding Road from the Late Teens through the Twenties. Oxford University Press, Oxford, England.

3. Arnett, J. J. (2016). Does emerging adulthood theory apply across social classes? National data on a persistent question. Emerging Adulthood, 4(4), 227–235. https://doi.org/10.1177/2167696815613000

4. Arnett, J. J., & Schwab, J. (2012). The Clark University poll of emerging adults: Thriving, struggling, & hopeful. (December), 22. Retrieved from clarku.edu/poll

5. Bergen, S. E., Gardner, C. O., & Kendler, K. S. (2007). Age-related changes in heritability of behavioral phenotypes over adolescence and young adulthood: A meta-analysis. Twin Research and Human Genetics. https://doi.org/10.1375/twin.10.3.423

6. Boker, S., Neale, M., Maes, H., Wilde, M., Spiegel, M., Brick, T., … Fox, J. (2011). OpenMx: an open source extended structural equation modeling framework. Psychometrika, 76, 306–317. https://doi.org/10.1007/s11336-010-9200-6

7. Bonnie, R. J., Stroud, C., & Breiner, H. (2015). Investing in the health and well-being of young adults. In Investing in the Health and Well-Being of Young Adults. https://doi.org/10.17226/18869

8. Briley, D. a, & Tucker-Drob, E. M. (2013). Explaining the increasing heritability of cognitive ability across development: a meta-analysis of longitudinal twin and adoption studies. Psychological Science, 24, 1704–1713. https://doi.org/10.1177/0956797613478618

9. Cutler, D. M., & Lleras-Muney, A. (2012). Education and health: insights from international comparisons. In NBER Working Papers (No. 17738).

10. Harris, K. M., Gordon-Larsen, P., Chantala, K., & Udry, J. R. (2006). Longitudinal trends in race/ethnic disparities in leading health indicators from adolescence to young adulthood. Archives of Pediatrics and Adolescent Medicine, 160(1), 74–81. https://doi.org/10.1001/archpedi.160.1.74

11. Hatemi, P. K., & McDermott, R. (2012). The genetics of politics: Discovery, challenges, and progress. Trends in Genetics. https://doi.org/10.1016/j.tig.2012.07.004

12. Hufer, A., Kornadt, A. E., Kandler, C., & Riemann, R. (2020). Genetic and environmental variation in political orientation in adolescence and early adulthood: A nuclear twin family analysis. Journal of Personality and Social Psychology. https://doi.org/10.1037/pspp0000258

13. Johnson, M. K., Crosnoe, R., & Elder, G. H. (2011). Insights on adolescence from a life course perspective. Journal of Research on Adolescence, 21(1), 273–280. https://doi.org/10.1111/j.1532-7795.2010.00728.x

14. Kessler, R. C., Berglund, P., Demler, O., Jin, R., Merikangas, K. R., & Walters, E. E. (2005). Lifetime prevalence and age-of-onset distributions of DSM-IV disorders in the national comorbidity survey replication. Archives of General Psychiatry, 62(6), 593–602. https://doi.org/10.1001/archpsyc.62.6.593

15. Knopik, V. S., Neiderhiser, J. M., DeFries, J. C., & Plomin, R. (2017). Behavioral Genetics. 7th ed. Worth Publishers, New York.

16. Koenig, L. B., McGue, M., & Iacono, W. G. (2008). Stability and change in religiousness during emerging adulthood. Developmental Psychology, 44(2), 532. https://doi.org/10.1037/0012-1649.44.2.532

17. Kornadt, A. E., Hufer, A., Kandler, C., & Riemann, R. (2018). On the genetic and environmental sources of social and political participation in adolescence and early adulthood. PLoS ONE, 13(8), e0202518. https://doi.org/10.1371/journal.pone.0202518

18. Krapohl, E., Rimfeld, K., Shakeshaft, N. G., Trzaskowski, M., McMillan, A., Pingault, J.-B., … Plomin, R. (2014). The high heritability of educational achievement reflects many genetically influenced traits, not just intelligence. Proceedings of the National Academy of Sciences of the United States of America, 111(42), 15273–15278. https://doi.org/10.1073/pnas.1408777111

19. Lee, S. H., Yang, J., Goddard, M. E., Visscher, P. M., & Wray, N. R. (2012). Estimation of pleiotropy between complex diseases using single-nucleotide polymorphism-derived genomic relationships and restricted maximum likelihood. Bioinformatics, 28, 2540–2542. https://doi.org/10.1093/bioinformatics/bts474

20. McGue, M., & Bouchard, T. J. (1984). Adjustment of twin data for the effects of age and sex. Behavior Genetics, 14, 325–343. https://doi.org/10.1007/BF01080045

21. Medland, S. E. (2004). Alternate parameterization for scalar and non-scalar sex-limitation models in Mx. Twin Research : The Official Journal of the International Society for Twin Studies, 7(3), 299–305. https://doi.org/10.1375/twin.7.3.299

22. Mehta, C. M., Arnett, J. J., Palmer, C. G., & Nelson, L. J. (2020). Established adulthood: A new conception of ages 30 to 45. American Psychologist, 75(4), 431. https://doi.org/10.1037/amp0000600

23. Munafò, M. R., Tilling, K., Taylor, A. E., Evans, D. M., & Smith, G. D. (2018). Collider scope: When selection bias can substantially influence observed associations. International Journal of Epidemiology, 47(1), 226–235. https://doi.org/10.1093/ije/dyx206

24. Neale, M. C., Boker, S. M., Bergeman, C. S., & Maes, H. H. (2005). The utility of genetically informative data in the study of development. In Methodological Issues in Aging Research (pp. 269–327). https://doi.org/10.4324/9781315820989-14

25. Ong, A. D., & Bergeman, C. S. (2004). The Complexity of Emotions in Later Life. Journals of Gerontology - Series B Psychological Sciences and Social Sciences. https://doi.org/10.1093/geronb/59.3.P117

26. Ong, A. D., Bergeman, C. S., Bisconti, T. L., & Wallace, K. A. (2006). Psychological resilience, positive emotions, and successful adaptation to stress in later life. Journal of Personality and Social Psychology, 91(4), 730. https://doi.org/10.1037/0022-3514.91.4.730

27. Pain, O., Dudbridge, F., & Ronald, A. (2018). Are your covariates under control? How normalization can re- introduce covariate effects. European Journal of Human Genetics, 26(8), 1194–1201. https://doi.org/10.1038/s41431-018-0159-6

28. Park, M. J., Paul Mulye, T., Adams, S. H., Brindis, C. D., & Irwin, C. E. (2006). The health status of young adults in the United States. Journal of Adolescent Health, 39(3), 305–317. https://doi.org/10.1016/j.jadohealth.2006.04.017

29. Plomin, R., DeFries, J. C., Knopik, V. S., & Neiderhiser, J. M. (2016). Top 10 replicated findings from behavioral genetics. Perspectives on Psychological Science, 11(1), 3–23. https://doi.org/10.1177/1745691615617439

30. Polderman, T. J. C., Benyamin, B., De Leeuw, C. A., Sullivan, P. F., Van Bochoven, A., Visscher, P. M., & Posthuma, D. (2015). Meta-analysis of the heritability of human traits based on fifty years of twin studies. Nature Genetics, 47(7), 702–709. https://doi.org/10.1038/ng.3285

31. Price, T. S., Freeman, B., Craig, I., Petrill, S. A., Ebersole, L., & Plomin, R. (2000). Infant zygosity can be assigned by parental report questionnaire data. Twin Research : The Official Journal of the International Society for Twin Studies, 3, 129–133. https://doi.org/10.1375/twin.3.3.129

32. Rijsdijk, F. V, & Sham, P. C. (2002). Analytic approaches to twin data using structural equation models. Briefings in Bioinformatics, 3(2), 119–133. https://doi.org/10.1093/bib/3.2.119

33. Rimfeld, K., Kovas, Y., Dale, P. S., & Plomin, R. (2016). True grit and genetics: Predicting academic achievement from personality. Journal of Personality and Social Psychology, 111(5), 780–789. https://doi.org/10.1037/pspp0000089

34. Rimfeld, K., Malanchini, M., Spargo, T., Spickernell, G., Selzam, S., McMillan, A., … Plomin, R. (2019). Twins Early Development Study: A genetically sensitive investigation into behavioral and cognitive development from Infancy to emerging adulthood. Twin Research and Human Genetics, 22(6), 508– 513. https://doi.org/10.1017/thg.2019.56

35. Rimfeld, K., Malancini, M., Allegrini, A., Packer, A., McMillan, A., Ogden, R., … Plomin, R. (2020). Genetic correlates of psychological responses to the COVID-19 crisis in young adult twins in Great Britain. Research Square, 1–28. https://doi.org/10.21203/rs.3.rs-31853/v1

36. Selzam, S., McAdams, T. A., Coleman, J. R. I., Carnell, S., O’Reilly, P. F., Plomin, R., & Llewellyn, C. H. (2018). Evidence for gene-environment correlation in child feeding: Links between common genetic variation for BMI in children and parental feeding practices. PLoS Genetics, 14(11), e1007757. https://doi.org/10.1371/journal.pgen.1007757

37. Tanner, J. L. (2016). Mental health in emerging adulthood. In The Oxford Handbook of Emerging Adulthood.

38. Turkheimer, E., Pettersson, E., & Horn, E. E. (2014). A phenotypic null hypothesis for the genetics of personality. Annual Review of Psychology, 65, 515–540. https://doi.org/10.1146/annurev-psych-113011-143752

39. van de Weijer, M. P., de Vries, L. P., & Bartels, M. (2020). Happiness and Wellbeing; the value and findings from genetic studies. PsyArXiv.

40. Veenhoven, R. (2008). Healthy happiness: Effects of happiness on physical health and the consequences for preventive health care. Journal of Happiness Studies, 9(3), 449–469. https://doi.org/10.1007/s10902-006-9042-1

41. Yaffe, K., Vittinghoff, E., Pletcher, M. J., Hoang, T. D., Launer, L. J., Whitmer, R., … Sidney, S. (2014). Early adult to midlife cardiovascular risk factors and cognitive function. Circulation, 129(15), 1560–1567. https://doi.org/10.1161/CIRCULATIONAHA.113.004798

42. Yang, J., Lee, S. H., Goddard, M. E., & Visscher, P. M. (2013). Genome-wide complex trait analysis (GCTA): Methods, data analyses, and interpretations. Methods in Molecular Biology, 1019, 215–236. https://doi.org/10.1007/978-1-62703-447-0-9

